# Machine learning uncovers independently regulated modules in the *Bacillus subtilis* transcriptome

**DOI:** 10.1101/2020.04.26.062638

**Authors:** Kevin Rychel, Anand V. Sastry, Bernhard O. Palsson

## Abstract

The transcriptional regulatory network (TRN) of *Bacillus subtilis* coordinates cellular functions of fundamental interest, including metabolism, biofilm formation, and sporulation. Here, we use unsupervised machine learning to modularize the transcriptome and quantitatively describe regulatory activity under diverse conditions, creating an unbiased summary of gene expression. We obtain 83 independently modulated gene sets that explain most of the variance in expression, and demonstrate that 76% of them represent the effects of known regulators. The TRN structure and its condition-dependent activity uncover novel or recently discovered roles for at least 5 regulons, such as a relationship between histidine utilization and quorum sensing. The TRN also facilitates quantification of population-level sporulation states, revealing a putative anaerobic metabolism role for SigG. As this TRN covers the majority of the transcriptome and concisely characterizes the global expression state, it could inform research on nearly every aspect of transcriptional regulation in *B. subtilis*.

## Introduction

Cells interpret dynamic environmental signals to govern gene expression through a complex transcriptional regulatory network (TRN). *Bacillus subtilis*, a model gram-positive soil and gut bacterium, is one of the most widely studied species in microbiology, providing a rich background for understanding its TRN. This generalist organism is a model for processes such as sporulation^1^, biofilm formation^2^, and competence^3^ – all of which are key to understanding pathogenesis in other bacteria, such as *Staphylococcus aureus* and *Clostridium difficile. B. subtilis* is also commonly engineered for industrial production purposes^4^, which creates demand for practical knowledge about how it responds to stimuli and alters its gene expression.

In 2012, Nicolas, *et al*., generated a transcriptomic microarray dataset of *B. subtilis* with 269 expression profiles under 104 conditions^5^, which included growth over time in various media, carbon source transitions, biofilms, swarming, various nutritional supplements, a variety of stressors, and a time-course for sporulation. The wide scope and high quality of this dataset has led to its broad adoption. It is now the expression compendium featured on *Subti*Wiki, an online resource for *B. subtilis* that is one of the most widely used and complete databases for any organism^6^. *Subti*Wiki contains detailed biological descriptions and binding sites for hundreds of transcriptional regulators; however, binding sites alone cannot explain the condition-specific transcriptomic responses of bacteria to dynamic environmental conditions^7,8^.

Independent Component Analysis (ICA) is an unsupervised statistical learning algorithm that was developed to isolate statistically independent voices from a collection of mixed signals^9^. ICA applied to transcriptomic matrices simultaneously computes independently modulated sets of genes (termed “i-modulons”) and their corresponding activity levels in each experimental condition^10^. I-modulons can be interpreted as data-driven regulons, though they rely on observed expression changes instead of transcription factor binding sites. The condition-dependent activity level of i-modulons indicates how active the underlying regulator is. Since the number of i-modulons is substantially fewer than the number of genes, they are a significantly easier way to analyze systems-level cell behavior.

ICA has been shown to extract biologically relevant transcriptional modules for a variety of transcriptomic datasets, especially in yeast and human cancer^11–15^. It was the best out of 42 methods at recovering known co-regulated gene modules in a comprehensive examination of TRN inference methods^16^. ICA also obtained the most robust modules across datasets compared to similar factorization algorithms^17^. We previously applied this approach to a large, high quality Escherichia coli RNA-seq compendium and extracted 92 i-modulons, two thirds of which exhibited high overlap with known regulons^10^. This analysis provided many novel insights into the E. coli TRN, including the addition of genes to known regulons (validated through ChIP-exo), bifurcation of the purine synthesis regulon, the characterization of new regulons, and identification of clear associations between regulator mutations and activities. We have also applied ICA to transcriptomics of evolved strains to understand evolutionary trade-offs and regulatory adaptations in naphthoquinone-based aerobic respiration^18^, and to characterize the function of the transcription factor OxyR, which responds to peroxide^19^.

Given our success with ICA applied to RNA-seq data from a model gram-negative bacterium, we were sought to determine what it can uncover about a microarray dataset from a model gram-positive bacterium. Using the wealth of TRN knowledge and high quality expression data available on *Subti*Wiki, this analysis uncovers many novel insights. We determine the main functions and regulators that control a large fraction of the transcriptome, and we characterize the i-modulon accuracy in relation to the known TRN. I-modulon activities reveal relationships and stimuli that have been present in the data but never specifically investigated; it is therefore a powerful hypothesis-generating tool. We specifically present five unexpected i-modulon activations and hypotheses about their mechanisms. We characterize sporulation, which led us to the identification of three major transcriptomic stages in the process, including i-modulons for the known sigma factor cascade. Finally, we present three transcriptional units with little prior characterization that warrant further study.

## Results

### Independent component analysis reveals the structure of the B. subtilis transcriptome

We performed ICA on the Nicolas, *et al*., 2012 dataset^5^ (Methods, Dataset S1-2) and obtained 83 robust i-modulons (Dataset S3-6). These 83 i-modulons constitute the statistically independent gene expression signals found across the conditions used in the generation of this data. Together, they explain 72% of the variance in gene expression (Supplementary Methods, Supplementary Fig. S1B). Unlike regulons, which are sets of co-regulated genes based on experimentally confirmed DNA binding sites of transcriptional regulators, i-modulons are derived solely from the measured transcriptome – we did not provide this untargeted statistical approach with any information about the known regulon structure of the TRN (Fig. 1a). However, that structure is largely recapitulated in the results. 63 of the 83 i-modulons were successfully mapped to a known regulator, and an additional 3 are likely to be co-regulated by unknown mechanisms. The TRN covers 2,235 gene/i-modulon relationships, of which 1,536 are known gene/regulator interactions and 699 are new (Dataset S8). Our TRN structure contained seven i-modulons that exhibited perfect overlap with annotated regulons and whose activity levels match expectations, such as MalR (Supplementary Results, Supplementary Fig. S2). This illustrates that independent signals such as transcription factor binding, which dictate gene expression, lead to observable signals in the TRN from condition to condition, and ICA was able to identify them.

**Fig. 1.**
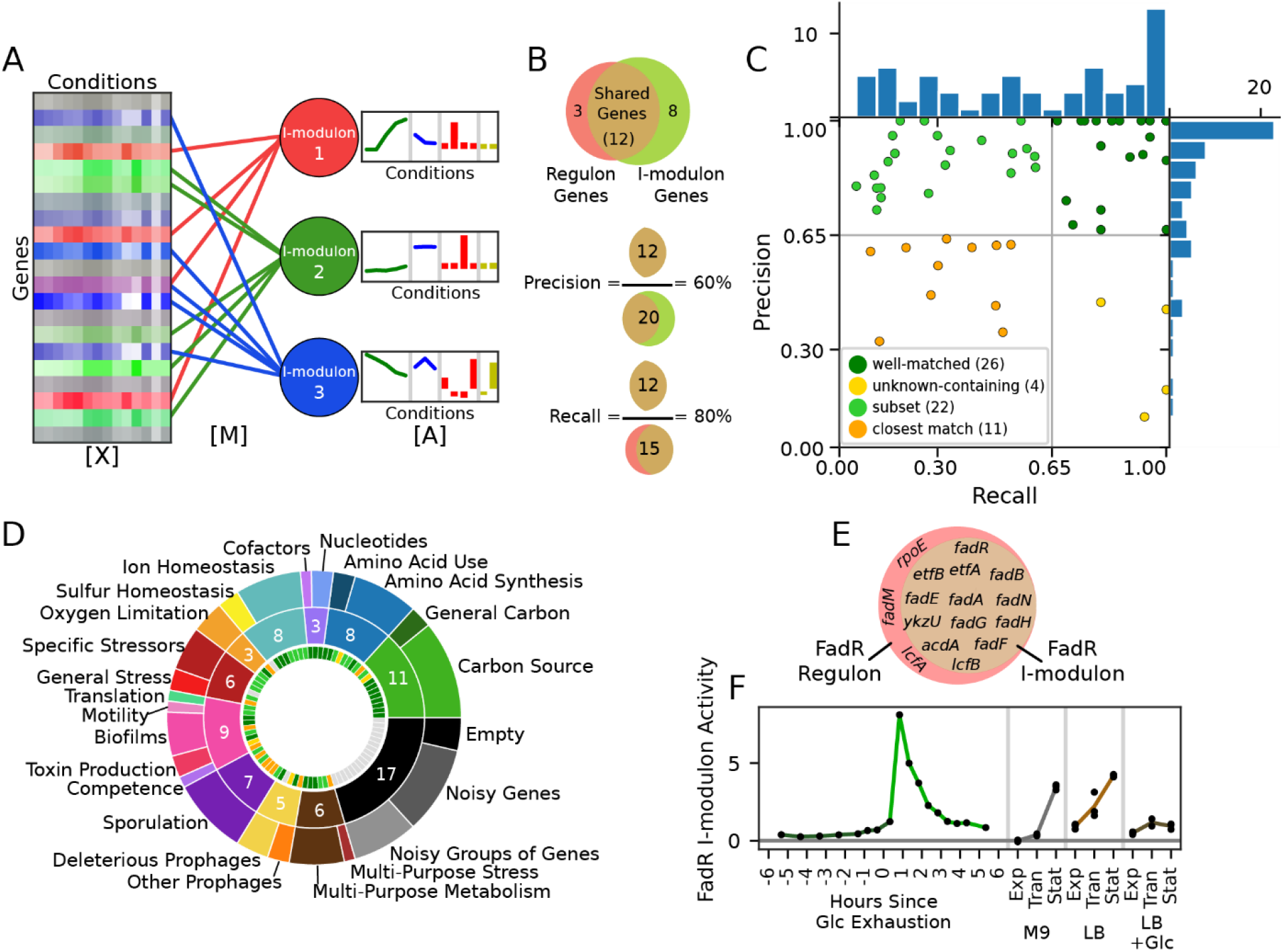
Independent Component Analysis (ICA) extracts regulatory signals from a compendium of transcriptomic data. **a**. Given a matrix of gene expression data, **X** (Dataset S2), ICA identifies independently modulated sets of genes (i-modulons) in the transcriptome which are linked to genes through the matrix **M** (Dataset S3). Three i-modulons are symbolically represented; the red i-modulon consists of 4 genes, and the green and blue i-modulons consist of 5 genes. The condition-dependent activities of the i-modulons are stored in the matrix **A** (Dataset S4). The bar chart indicates the activity levels of the i-modulons under different conditions, where the colors indicate different experiments. The three matrices are related as **X**=**M**\***A. b**. Graphical representation of the definitions of precision and recall of a given i-modulon and the corresponding regulon (example numbers shown). **c**. Scatter plot of precision and recall of the enrichments for the 63 (out of 83) i-modulons that were matched to a regulon. Histograms in the margins demonstrate the high precision of most enrichments (see Dataset S7, Supplementary Fig. S1C for more details). **d**. Donut chart of i-modulon functions. The outermost ring lists specific functions and the center ring lists broad functions, with the number of i-modulons in the broad category shown in white. The innermost ring shows the regulon confidence quadrant of the corresponding i-modulon, as defined in **c. e-f**. An example i-modulon that was enriched for FadR. **e**. Venn diagram of the FadR i-modulon genes and the FadR regulon (non-coding RNAs have been omitted). **f**. Activity level found in a row of A for four experiments (separated by vertical gray lines) from the dataset. Activity levels increase during growth in the absence of glucose (M9 media, gray; LB media, light brown), remain low during growth in the presence of glucose (dark green, dark brown) and spike upon glucose starvation (green). ‘Exp’, ‘Tran’ and ‘Stat’ refer to exponential, transition, and stationary phase, respectively. See Dataset S1 for detailed growth conditions.

I-modulons are given a short name, usually based on their enriched regulator. If multiple regulators control an i-modulon, their names are separated by ‘+’ to indicate the intersection of the regulons, or ‘/’ to indicate the union of the regulons. In some cases, a different name was chosen based on the primary regulator, gene prefix, or most representative gene in the set (Dataset S7).

The relationship between i-modulons and regulators can be characterized by two measures: 1) precision (the fraction of i-modulon genes captured by the enriched regulon) and 2) recall (the fraction of the regulon contained in the i-modulon) (Fig. 1b). These two measures can be used to classify i-modulons into six groups (Fig. 1c). 1) The ‘well-matched’ group (n = 26) has precision and recall greater than 0.65. It includes several regulons associated with specific metabolites. 2) The ‘subset’ i-modulons (n = 22) exhibit high precision and low recall. They contain only part of their enriched regulon, perhaps because the regulon is very large and only the genes with the most transcriptional changes are captured. This group contains global metabolic regulators such as CcpA and CodY, as well as the stress sigma factors. 3) A third group, deemed ‘unknown-containing’ (n = 4), has low precision but high recall. These i- modulons contain some co-regulated genes along with unannotated genes which may have as-yet-undiscovered relationships to the enriched regulators (Dataset S8), or at least be co-stimulated by the conditions in the dataset. 4) The remaining enriched i-modulons are called ‘closest match’ (n = 11) because neither their precision nor recall met the cutoff, but the grouping had statistically significant enrichment levels and appropriate activity profiles. The difference in gene membership between these i-modulons and their regulons provide excellent targets for discovery. The i-modulons with no enrichments comprise the last two groups: 5) ‘new regulons’ (n = 3) are likely to be real regulons with unexplored transcriptional mechanisms, while 6) the remaining ‘uncharacterized’ i-modulons were likely to be noise due to large variance within conditions or the fact that they contain one or fewer genes.

Functional categorization of i-modulons provides a systems-level perspective on the transcriptome (Fig. 1d). Metabolic needs account for approximately one-third of the i-modulons, while comparatively fewer i-modulons deal with stressors, lifestyle choices such as biofilm formation and sporulation, and mobile genetic elements like prophages. Some i-modulons have multiple biological functions, such as one which synthesizes both nicotinamide and biotin. These i-modulons may result from co-stimulation of the different functions by all conditions probed in the dataset (e.g., both nicotinamide and biotin synthesis were always stimulated together by minimal media, so the algorithm could not separate them into unique signals).

The FadR i-modulon provides an example of the information encoded by the i-modulon gene membership (Fig. 1e) and activities (Fig. 1f). All genes within this i-modulon are regulated by FadR, so this enrichment has 100% precision. Three genes that are annotated as belonging to the FadR regulon were not captured in the i-modulon – *lcfA, rpoE*, and *fadM*. However, all three have additional regulation separate from that of FadR^20,21^, which may lead them to have divergent expression from the rest of the i-modulon. The activity levels (Fig. 1f) reflect expectations: FadR genes are repressed by FadR in the presence of long chain acyl-coA, and FadR itself is repressed by CcpA in the presence of fructose-1,6-bisphosphate^21^, which causes the expression to rise as nutrients (specifically sugars and fats) are depleted, and to be particularly strong immediately following glucose exhaustion.

### I-modulons generate novel hypotheses

I-modulon activities can often be explained by prior knowledge, as was the case with FadR. However, they can also present surprising relationships that lead to the generation of new hypotheses or strengthen arguments for recently proposed mechanisms. Here, we list five such examples, and more are provided in the supplementary results.

#### Ethanol stimulates tryptophan synthesis

The tryptophan synthesis i-modulon (*trpEDCFB*) was strongly activated under ethanol stress (Fig. 2a), a response which has not been previously documented in bacteria. This i-modulon is regulated by the *trp* attenuation protein (TRAP), which represses its genes in the presence of tryptophan^22^. Therefore, this activation indicates that ethanol is probably depleting intracellular tryptophan concentrations. Exploring the tryptophan synthesis pathway reveals a hypothetical mechanism for this depletion: flux from the precursor chorismate may be redirected to replenish folate that has been damaged by ethanol oxidation byproducts^23^ (Supplementary Fig. S4A). If this hypothesis is accurate, it may inform research on the tryptophan deficiency and neurotransmitter metabolism problems observed in human alcoholic patients^24,25^, especially given that *B. subtilis* is an important folate producer in the gut microbiome^26,27^.

**Fig 2.**
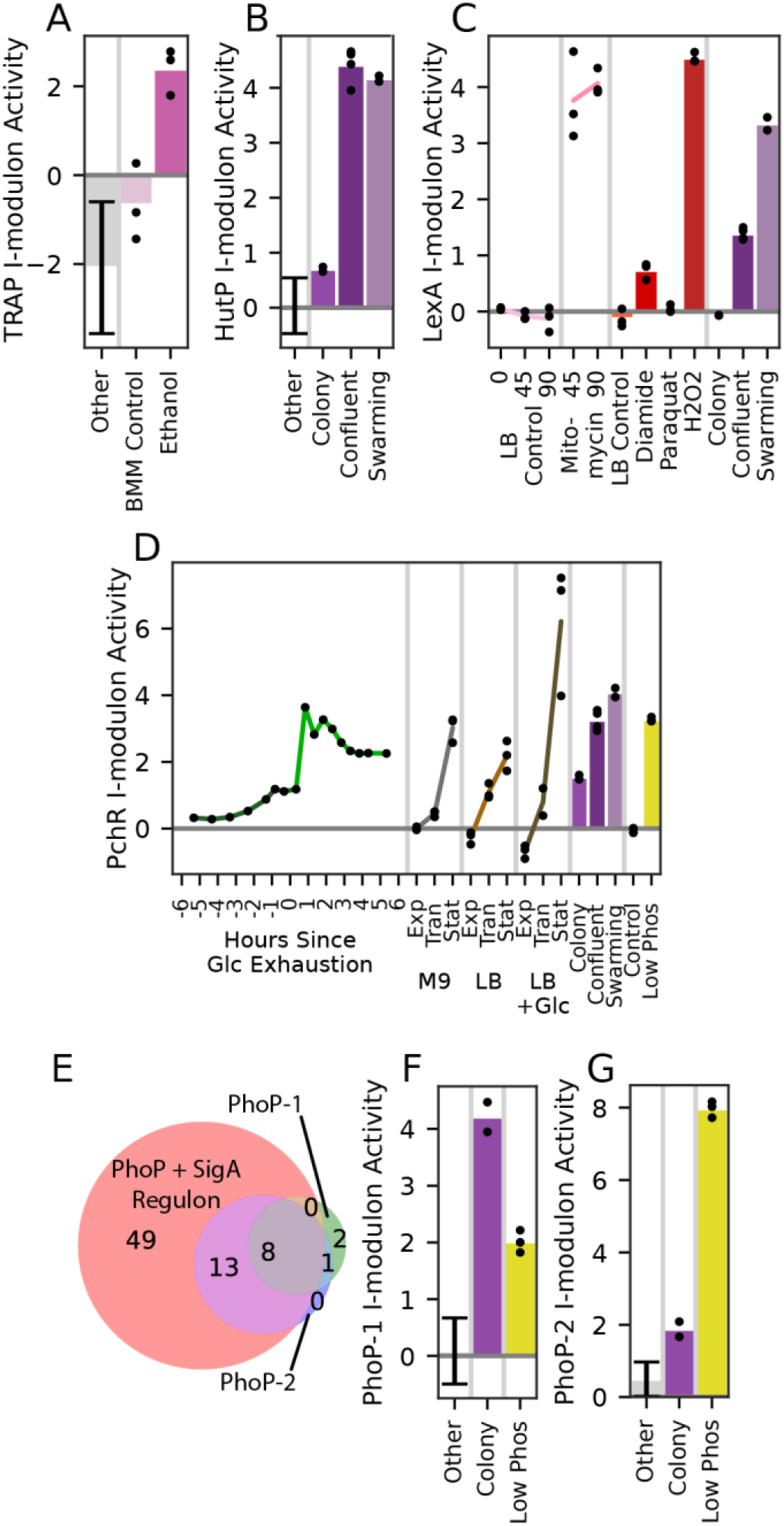
I-modulons provide a range of new insights. Error bars: mean ± standard deviation; black dots indicate separate samples; vertical gray lines separate different experiments in the dataset. ‘Other’ category includes all conditions except sporulation and those shown. **a**. Tryptophan synthesis (TRAP) i-modulon activity, which is unexpectedly elevated by ethanol (Supplementary Fig. S4A). The experiment was carried out in Belitsky minimal medium (BMM). **b**. Histidine utilization (HutP) i-modulon activity, which is strongest in quorum conditions. **c**. LexA i-modulon activity is elevated by DNA damage (mitomycin and peroxide) and in swarming (Supplementary Fig. S4B) **d**. Pulcherrimin (PchR) i-modulon activity increases when growth is expected to slow. **e**. Venn diagram of gene presence in the PhoP regulon and i-modulons. Numbers indicate the amount of genes or non-coding RNAs in each subset. **f-g**. Bar graphs of PhoP i-modulon activity demonstrating the use of PhoP-1 for early biofilm growth (‘Colony’ refers to individual colonies on a plate after 16 hours) and PhoP-2 for extreme phosphate starvation (‘Low Phos’ indicates phosphate starvation for 3 hours). ‘Other’ refers to all conditions besides those shown.

#### Histidine is utilized by quorums

The HutP i-modulon for histidine utilization (*hutHUIGM*) is controlled by an antiterminator that derepresses it in the presence of excess histidine, as well as by the master regulators CcpA and CodY; therefore, its activation indicates that histidine is plentiful while other amino acids are not, and that carbon sources are poor^28^. Surprisingly, it was by far most strongly activated in confluent biofilms and swarming cells (Fig. 2b). Independent colonies from the same experiment do not exhibit activation, which leads us to rule out the media composition as the reason for these activity levels. The connection between these lifestyle conditions and histidine metabolism has not been studied in *B. subtilis*, but it has been observed in *A. baumannii*, where histidine degradation was shown to be upregulated in proteomic studies of biofilms, and histidine supplementation stimulated increased biofilm production^29^. Two recent studies discovered that biofilm-inhibiting antimicrobials worked by suppressing histidine synthesis in *Staphylococcus xylosus*^30,31^. One proposed mechanism implicated the production of extracellular DNA, which is an important component of both *A. baumannii* and *B. subtilis* biofilms^32^. Given that this i-modulon is also activated by swarming cells, an alternative hypothesis may be that HutP is involved with quorum sensing or surfactant production: both activating conditions have a quorum and high surfactant production, while independent colonies do not.

#### DNA damage stimulates swarming

The LexA i-modulon regulates the SOS response for DNA protection and repair. It is strongly activated by three conditions (Fig. 2c). LexA stimulation by mitomycin and hydrogen peroxide is expected since those conditions damage DNA^33,34^. Unexpectedly, this i-modulon is also activated in swarming cells despite a lack of DNA damaging agents in that condition. We propose a potential mechanism for this activation: recent research has indicated that certain cells in a culture will tend to accumulate reactive oxygen species and DNA damage. Those cells will produce Sda (a developmental checkpoint protein), and form a subpopulation separate from those that produce biofilm^35^. The LexA+, biofilm– population would no longer be producing EpsE, which catalyzes a step in the biofilm synthesis process and also suppresses swarming^36^. Therefore, we predict that DNA damage encourages swarming motility based on i-modulon activation and this mechanism (Supplementary Fig. S4B).

#### An iron chelator signals the stationary phase

The PchR i-modulon produces, extrudes, and imports pulcherrimin, an iron chelator^37^. Over all of the exponential to stationary phase growth experiments, we observe increases in PchR activation (Fig. 2d). We also see PchR activation in late stage biofilm, glucose exhaustion, and phosphate starvation experiments. These results agree with a recent study that found pulcherrimin to be an important intercellular signal for the stationary phase that also helps exclude competing bacteria from established biofilms^38^. The regulation mechanisms of i-modulons like this one can be the subject of future research.

#### Phosphate limitation stimulates tiers of regulation

The PhoP regulon controls phosphate homeostasis. It appears as two separate i-modulons (Fig. 2e-g). PhoP-1 encodes high-affinity phosphate uptake transporters. Phosphate is used to produce (and is effectively stored in) teichoic acid, which is a major component of the cell wall. As a colony grows, it must uptake phosphate to produce more cell walls – indeed, teichoic acid intermediates are the major stimulus for PhoP activity^39^. It is therefore unsurprising that PhoP-1 is strongly activated in independent colonies, which are exponentially growing in close quarters with low local free phosphate concentrations. PhoP-2 contains PhoP-1 as well as 13 other genes which encode more extreme phosphate recovery strategies: *phoABD*, which salvages phosphate monoesters but produces reactive alcohols, *glpQ*, which degrades extracellular teichoic acid, and *tuaBCDEFGH*, which replaces teichoic acid with phosphate-free teichuronic acid. PhoP-2 is only active under phosphate starvation, consistent with the extreme strategy it encodes. Perhaps the affinities of the promoters of the PhoP-2 specific genes are lower than that of the PhoP-1 genes, which could lead to this graded response.

### Six i-modulons capture the major transcriptional steps of sporulation

The dataset we analyzed contained an eight-hour sporulation time course, which yielded six major sporulation i- modulons that were activated sequentially over the first six hours (Fig. 3a). The identification of these gene sets by ICA indicates coherent expression across the transcriptome, and more dramatic transcriptional variation compared to excluded genes. The conclusions drawn from these i-modulons are limited by the complexity of sporulation^1,40^ and the stochasticity of its onset^41^. Because of this, we observe many genes shared between consecutive i-modulons (Supplementary Fig. S7a). Nonetheless, the following analysis demonstrates that they still provide valuable information, including identifying 20 unknown proteins that could now be classified as putatively sporulation-related (Supplementary Fig. S7b, Dataset S8).

**Fig. 3.**
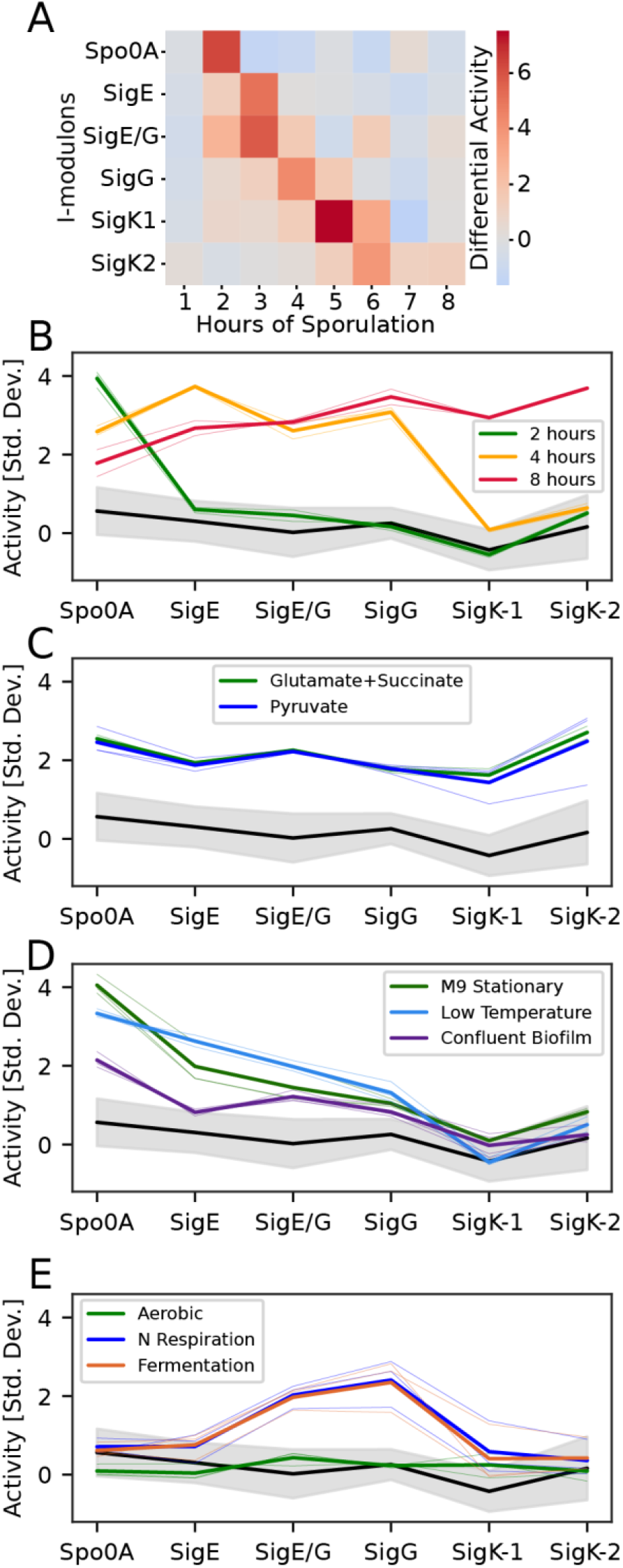
Six I-modulons (named for their enriched regulators) mark progress through sporulation. **a**. Heatmap color indicates change in i-modulon activity over the previous hour. **b-e**. Line plots of the sporulation progression for selected conditions, with thick lines indicating mean activity and thin lines indicating individual samples. Activity levels were divided by the standard deviation. The black line surrounded by a shaded gray region is the average of all conditions not shown in any plot ± standard deviation. **b**. Three time points of sporulation, showing Spo0A activation at sporulation onset (2 hours, green), cumulative expression up to the fourth step (SigG) for an intermediate time point (4 hours, orange), and expression of all stages at 8 hours (red). **c**. Minimal media supplemented with these carbon sources leads to expression of all sporulation i-modulons. **d**. Three conditions reached the intermediate steps of sporulation. **e**. Anaerobic conditions exhibit unusual activity. ‘Aerobic’ is the control condition.

The gene sets and regulators of the sporulation i-modulons roughly match the known sporulation progression (Supplementary Fig. S7c-f). The Spo0A i-modulon contains mostly genes known to be activated by high levels of Spo0A∼P, including the sigma factors for upcoming sporulation steps, chromosome preparation machinery, and septal wall formation. It is rapidly activated between hours 1 and 2 of the time course. Next, the SigE i-modulon carries out functions in the mother cell for engulfment of the forespore. Notably, SigF is the only absent sigma factor; it should be present in the forespore at this point in the sporulation process, but we believe it was not identified because its genes are expressed to a lesser degree in the overall population due to the small size of the forespore at this stage. It is replaced by a dual SigE/G i- modulon, which regulates early spore coat formation by both the mother and forespore cells. The SigG i-modulon follows; it contains more coat maturation proteins, germination receptors, metabolic enzymes, and stress resistance genes. Finally, the SigK regulon is split into two i- modulons with functions including DNA protection, coat maturation, and mother cell lysis. These functions and regulators largely match expectations based on literature, providing an *a priori* validation of the set of known sporulation steps.

The activity levels of the sporulation i-modulons can be viewed as markers of progress through sporulation: high Spo0A activity indicates that new spores are forming, and high SigK-2 indicates that some spores are completing the process. Therefore, we can understand how far along other conditions are based on their sporulation activity levels (Fig. 3b-e). Most conditions have a very low level of activation, but the “glutamate + succinate” and pyruvate supplements to minimal media conditions both have elevated expression across all sporulation i-modulons, which indicates that the poor carbon sources in these conditions stimulated sporulation (Fig. 3c). Indeed, pyruvate has been shown to regulate sporulation^42,43^. Some other conditions appear to have made it partway through the process: confluent biofilms, the stationary phase in minimal media, and growth at cold temperature all reached the third of six steps. This is appropriate for these conditions based on previous studies^44–46^ (Fig. 3d).

With one exception, the progression from one sporulation i-modulon to the next is cumulative: we do not see strong activation of step 2 unless step 1 is active, and so on. This agrees with prior observations^47^. The only exception to this rule is elevated SigG activity by cells in anaerobic conditions (Fig. 3e). This connection is also evident from gene presence: a flavohemoglobin required for anaerobic growth, hmp, is part of the i-modulon despite no known connection to SigG. Previous studies have also acknowledged that some SigG-dependent genes are required for anaerobic survival^48^. Despite that, recent research into the activation of SigG has stated that it should never be expressed during vegetative growth, and framed all activation mechanisms within the context of SigF and other sporulation-specific molecules^49^. That study did not explore anaerobic conditions. We therefore propose that there may be an anaerobic mechanism through which SigG or its known targets are activated, distinct from sporulation.

### Changes in i-modulon activity reveal global transcriptional shifts during sporulation

In complex processes such as sporulation, the entire cellular transcriptome undergoes system-wide changes beyond those directly related to the process at hand. While much effort has been put into understanding metabolic changes at the onset of sporulation^1,45,47^, metabolic and lifestyle-related regulatory activity are difficult to summarize concisely with previous methods. Because ICA provides a simple method for tracking transcriptome-wide changes, we analyzed activity level fluctuations for the sporulation time course (Fig. 4). Three major stages are involved: a self-preserving metabolic response to amino acid starvation in the first hour, a community-wide lifestyle reallocation in the second hour, and progression through sporulation in the remaining time points.

**Fig. 4.**
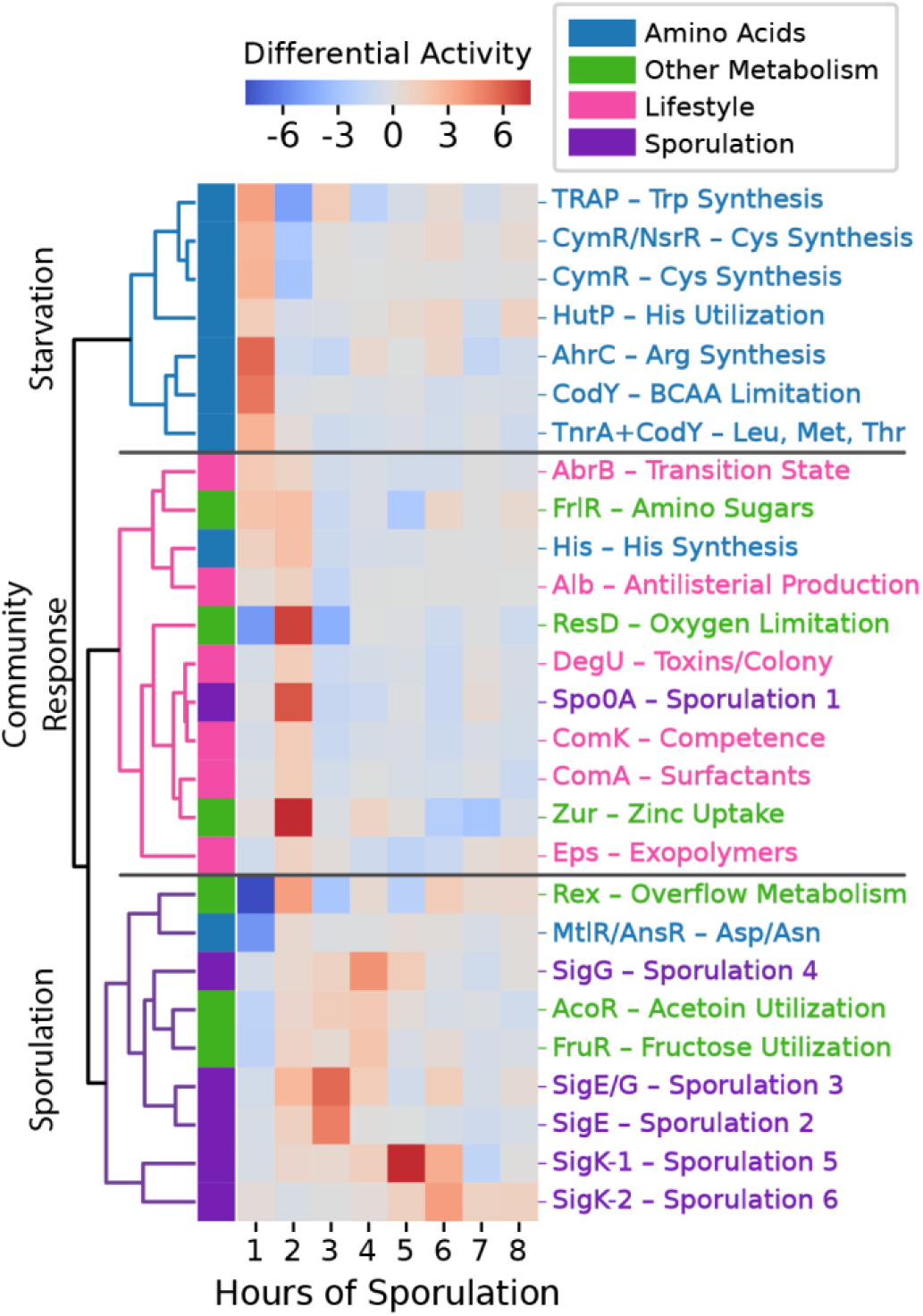
Changes in i-modulon activity reveal global transcriptional reallocation during sporulation. Heatmap color indicates change in i-modulon activity over the previous hour. Selected i-modulons were hierarchically clustered according to the Pearson R correlation between sporulation activity derivatives.

In the first hour, many amino acid synthesis i-modulons (tryptophan, cysteine, arginine, leucine, and threonine) and one amino acid utilization i-modulon (histidine) are rapidly activated. This is likely the result of amino acid starvation by the sporulation media, which derepresses these i- modulons through transcription factors including CodY. CodY also derepresses the fructosamine consumption i-modulon^50^ at this time. The AbrB i-modulon is activated as well; it responds to nutrient limitation through a variety of functions, including cannibalism^51^, that herald the stationary phase and prolong entry into sporulation.

In the second hour, Spo0A is strongly activated in a process that has been widely studied; this marks the onset of sporulation^52^. Also, the histidine utilization of the first hour is compensated by histidine synthesis in the second hour. Zinc, an important cofactor for sporulation proteins^53,54^, is taken up. Various colony, biofilm, and antimicrobial i-modulons are activated to support the forming spores (DegU, ComA, Eps, Alb). ComK, the competence i-modulon, is expressed as an alternative response to starvation. ComK’s brief activation at this time point is consistent with the short competence window observed before commitment to sporulation^3^. We also observe activation of resD, which is typically associated with anaerobic conditions^55,56^, and Rex, which regulates overflow metabolism, providing interesting connections to the potential anaerobic activity of SigG discussed in the previous section.

As sporulation continues, fewer non-sporulation i-modulons are activated. The notable exceptions are AcoR and FruR, which are both activated around the fourth hour. Both acetoin and polymeric fructose function as extracellular energy stores^57,58^, so perhaps they are used at this stage to provide a final energy source for the completion of sporulation. Overall, these results demonstrate an application of ICA for observing transcriptome-wide changes and lay out the major population dynamics and metabolic changes that underscore spore formation.

### Some poorly characterized i-modulons may perform important functions

Given the vast number of uncharacterized genes in bacterial genomes, ICA can help to narrow the search for new and important regulons by identifying groups of genes with transcriptional co-regulation (Dataset S5, Dataset S8) and their corresponding activity levels. We have identified three i- modulons that warrant further study. The first, the *ndhF-ybcCFHI* operon, may be involved in heat shock and germination (Fig. 5a, Supplementary Results). Another, the *yrkEFHI* operon, contains putative sulfur carriers that are very likely to assist in the cellular response to diamide stress (Fig. 5b, Supplementary Results).

**Fig. 5.**
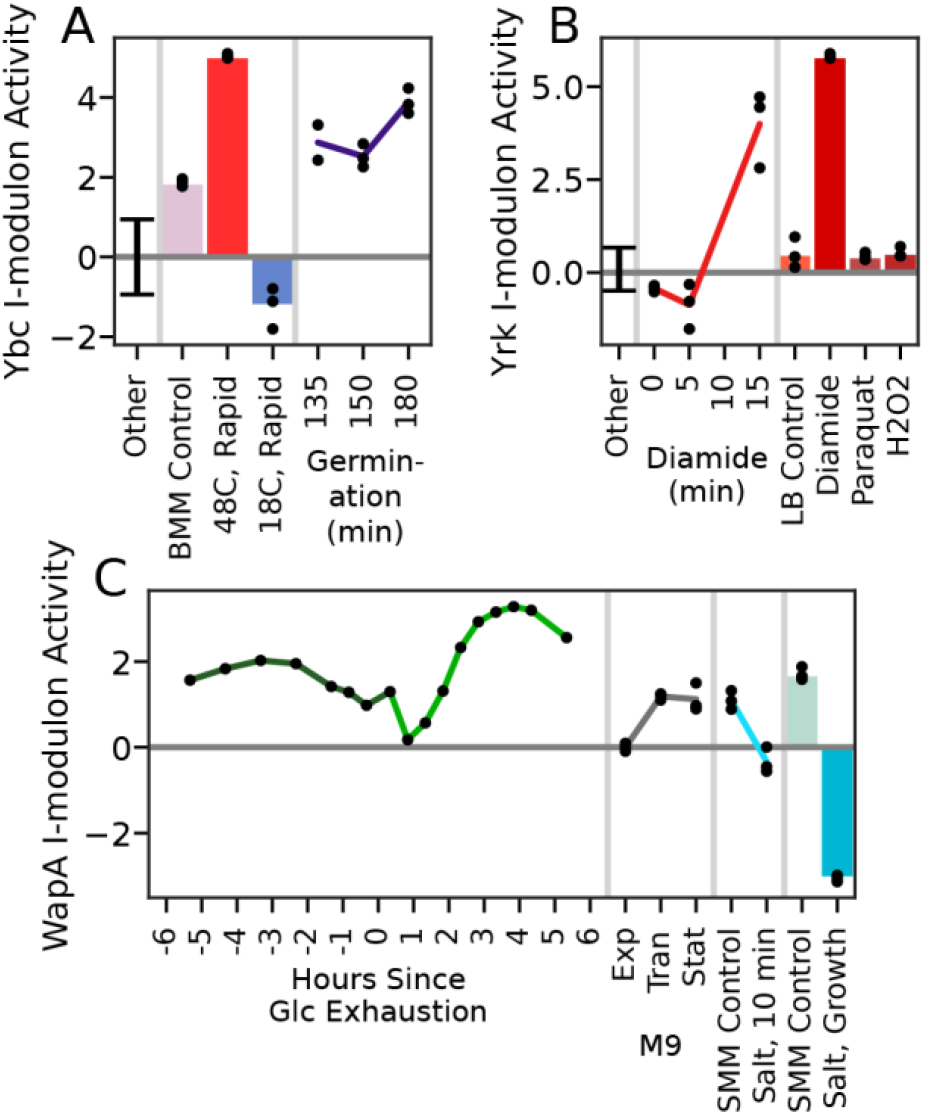
The activity levels of uncharacterized i-modulons agree with their putative functions. Bars and lines indicate means, black dots indicate individual samples, and error bars indicate one standard deviation. The ‘other’ category includes all conditions except the ones in the plot. Vertical gray lines separate different experiments in the dataset. **a**. The activity levels of the Ybc i- modulon indicate that it may be a response to heat shock or germination. **b**. The activity levels of the Yrk i-modulon (putative sulfur carriers) suggest that it is a response to diamide. The three conditions on the right were taken from LB cultures 10 minutes after exposure to the labeled stressor. **c**. The activity levels of the WapA i- modulon indicate activation by nutrient limitation (glucose exhaustion and the three growth phases of M9 media) and suppression by osmotic stress, both in the short (light blue time course) and long term (bars). ‘Exp’, ‘Tran’ and ‘Stat’ refer to exponential, transition, and stationary phase, respectively.

Also, the WapA i-modulon contains several uncharacterized genes which may be co-regulated by YvrHb, DegU, and WalR and participate in a unique, recently discovered interspecies competition mechanism^59^. This system protrudes fibers from the cell wall to deliver the WapA tRNase to enemy bacteria, potentially compromising cell wall integrity for greater nutrient availability. We observe activation of this i-modulon under starvation conditions and repression under cell wall stress (Fig. 5c), consistent with its putative function.

The other uncharacterized i-modulons which are not likely to be noise are prophage elements, whose regulatory mechanisms and effect on phenotype warrant further study. See Dataset S8 for their gene sets, Dataset S9 for summaries of their activating conditions, and Dataset S6 for graphical summaries.

## Discussion

Here, we decomposed the existing, high-quality *B. subtilis* expression dataset^5^ using ICA. This decomposition identified 83 i-modulons in the transcriptome whose overall activity can explain 72% of the variance in gene expression across the wide variety of conditions used to generate the dataset. Sixty-six of the i-modulons correspond to specific biological functions or transcriptional regulators. We analyzed the gene sets and activity levels of the i-modulons and presented findings that either agree with existing knowledge or generate new hypotheses that could be tested in future studies. The remaining 17 i-modulons are independent signals with no coherent biological meaning.

Through the application of ICA, we were able to identify well-studied gene sets with high accuracy (such as the MalR and FadR i-modulons), and uncover novel insights that suggest candidate underlying mechanisms. We discovered unexpected relationships between stress, metabolism, and lifestyle: ethanol appears to stimulates tryptophan synthesis, histidine utilization may be a feature of quorum sensing, DNA damage may induce swarming, and the iron chelator pulcherrimin could help to signal the stationary phase. The tiered response to phosphate limitation was captured as two separate i-modulons, which may provide evidence for variable promoter affinity across the known regulon. ICA accurately decomposed sporulation into a small set of steps which allow sporulation progress to be tracked; this led to the discovery of unusual activity for SigG in anaerobic conditions that may indicate a secondary function for this sporulation sigma factor. The global transcriptional response to sporulation in metabolism and lifestyle governance was summarized concisely in three stages by i-modulon activities.

Finally, three i-modulons contain mostly uncharacterized gene sets, which represent a promising area for further research. Overall, we have demonstrated that ICA produces biologically relevant i-modulons with hypothesis-generating capability from microarray data in this model gram-positive organism.

The i-modulon genes and activity profile data (Datasets S3-5), along with graphical summaries (Dataset S6) are available for examination by microbiologists with specific interests about functions in *B. subtilis* that are not detailed in this article. Code for our analysis pipeline is maintained on github (https://github.com/SBRG/precise-db). There is a strong potential for protein identification, transcription factor discovery, metabolic network insights, function assignment, and mechanism elucidation derived from this i-modulon structure of the TRN.

As with all machine learning approaches, the results from ICA improve as it is provided with more high-quality data^10^. Future research may append unique conditions to this dataset and observe the changes to the set of i-modulons it finds. Perhaps multi-purpose i-modulons will be divided into their biologically accurate building blocks, noise will be removed, and new regulons emerge as the signal-to-noise ratio improves. With enough additional data, ICA could potentially characterize the entire TRN in great detail, a goal which has been the subject of research for over half a century. Ultimately, this could be the foundation for a comprehensive, quantitative, irreducible TRN.

## Methods

### Data Acquisition and Preprocessing

We obtained normalized, log2-transformed tiling microarray expression values from Nicolas *et al*., 2012 (GEO accession number GSE27219)^5^, which span 5,875 transcribed regions (4,292 coding sequences and 1632 previously unannotated RNAs) and 269 sample profiles (104 conditions). The strain used, BSB1, is a prototrophic derivative of the popular laboratory strain, 168. Three samples (S3_3, G+S_1, and Mt0_2) were removed so that the Pearson R correlation between biological replicates was no less than 0.9, except in the case of sporulation hour 8, where n = 2 and R = 0.89 (Supplementary Fig. S1a). To obtain more easily interpretable activity levels, we centered the data by subtracting the mean in the M9 exponential growth condition from all gene values.

### Independent Component Analysis

Independent Component Analysis decomposes a transcriptomic matrix (**X**, Dataset S2) into independent components (**M**, Dataset S3) and their condition-specific activities (**A**, Dataset S4):

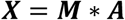

ICA was performed as previously described^10^. Note that the **M** matrix was previously called **S**; it has been changed to avoid confusion with other nomenclature. See Supplementary Methods.

We normalized each component in the **M** matrix such that the maximum absolute gene weight was 1. We performed the inverse normalization on the **A** matrix to conserve the same values. Therefore, each unit in **A** is equivalent to a unit log(TPM) change in expression if the i-modulon were to contain only one gene.

Thresholds were applied to the columns in the **M** matrix to acquire gene sets for each i-modulon (Supplemental Methods).

### Regulator Enrichment

Regulon information was obtained from *Subti*Wiki^6^. For each i- modulon, we obtained all regulators that regulate any gene in their gene sets. We also used all combinations of regulators, denoted by ‘+’ between regulator names, to capture regulons with more than one regulator. For each of those individual regulators and regulator combinations, we obtained a regulon set, a list of all genes that share that regulation. Next, we computed p-values for each regulon’s overlap with the i-modulon gene set using the two-sided Fisher’s exact test (FDR < 10^−5^)^60,61^. We also computed F1 scores, which are the harmonic averages of precision and recall.

After the sensitivity analysis determined the appropriate cutoff, significant enrichments for each i-modulon were then manually curated (Dataset S7). In most cases, the most significant enrichment was chosen. Some I-modulons appeared to be a combination of two or more significantly enriched regulons, so their assigned regulator was a union of both, denoted by ‘/’ between regulator names.

### Differential Activation Analysis

Differential activation analysis was performed as previously described^10^. We fit a log-normal distribution to the differences in i-modulon activities between biological replicates for each i-modulon. For a single comparison, we computed the absolute value of the difference in the mean i-modulon activity and compared against the i-modulon’s log-normal distribution to determine a p-value. We performed this comparison (two-tailed) for a given pair of conditions across all i-modulons at once and designated significance as FDR < 0.01.

### Data and Code Availability

All data generated or analyzed during this study are included in this published article (and its supplementary information files). The original dataset is from Nicolas *et al*., 2012 (GEO accession number GSE27219)^5^. Code for our analysis pipeline is maintained on github (https://github.com/SBRG/precise-db).

## Acknowledgements

We thank Dr. Joe Pogliano, Saugat Poudel, and Eammon Riley for helpful discussions and biological insights. This research used resources of the National Energy Research Scientific Computing Center, a DOE Office of Science User Facility supported by the Office of Science of the U.S. Department of Energy under Contract No. DE-AC02-05CH11231. This work was funded by the Novo Nordisk Foundation Center for Biosustainability (grant number NNF10CC1016517).

## Author Contributions

K.R. analyzed data and drafted the paper; A.V.S. designed research; A.V.S. and B.O.P. provided mentorship and guidance throughout. All participated in writing the paper.

