## Appendix for "Machine learning uncovers independently regulated modules in the *Bacillus subtilis* transcriptome"

### Supplementary Results

#### *The MalR i-modulon perfectly captures the corresponding regulon.*

The dataset contained carbon source transitions from glucose to malate and vice versa[1], which provided rich data for understanding the transcriptomics of malate metabolism. The MalR regulon is composed of 4 genes which are activated by extracellular malate and not directly affected by master regulators such as CcpA[2,3]. ICA produced an i-modulon composed of the same 4 genes (Supplementary Fig. S2A). The i-modulon activity rapidly increased upon malate addition to glucose media and declined over the course of glucose addition to malate media (Supplementary Fig. S2B), as would be expected. Six other i-modulons also have perfect overlap with their corresponding regulon annotations; they are mostly associated with specific carbon sources (Supplementary Fig. S1C, Dataset S7).

#### *Amino acid metabolism i-modulons exhibit generally expected behavior.*

Eight i-modulons function in amino acid metabolism, either fulfilling synthesis (n = 6) or utilization (n = 2) roles. These i-modulons exhibit expected general patterns, but some of them are highlighted in these results because they have unpredicted activity levels under stress or specific lifestyles (See Dataset S9 for a full list of expected and unexpected activity levels for all characterized i-modulons). Generally, amino acid synthesis i-modulons are less active in rich media compared to minimal media as a result of the exogenous amino acid supply. Over the course of growth in minimal media, their activity tends to decrease as intracellular amino acid stores are built up and protein synthesis for growth declines (Supplementary Fig. S3A). The opposite is true of the two utilization i-modulons (HutP, RocR/PutR).

#### *The arginine synthesis (AhrC) i-modulon.*

The arginine synthesis i-modulon provides two interesting insights. Its genes include *argGHCJBDF* and *carAB*, which are known to be repressed in the presence of arginine by AhrC[4]. The first insight is that it also contains *artPQR* (Supplementary Fig. S3B), which are arginine importers not known to be transcriptionally regulated by AhrC – given that they are part of the same independent signal in the transcriptome, they likely share this regulation. In addition, the i-modulon was unexpectedly downregulated in salt shock, but not after growth in salt (Supplementary Fig. S3B); this has not been explored in previous studies. Here, the putative mechanism is less clear. It may involve the production of osmoprotective solutes such as proline[5], which might perturb metabolic networks in such a way that arginine concentrations increase and then downregulate these genes. After proline stores have been established, arginine concentrations appear to be restored. There is also evidence of a proline/arginine metabolic link in another i-modulon: the RocR/PutR joint i-modulon combines the utilization of both amino acids into one signal. Exploration of this relationship may help to understand broader changes in amino acid metabolism and its regulation under stress conditions.

#### *The CcpA regulon is captured by two i-modulons.*

The CcpA i-modulons regulate carbon catabolites in different phases of growth (Supplementary Fig. S5), which may suggest divergent preferences for carbon catabolites determined by growth phase and starvation state; the same catabolites that are preferred during exponential growth are preferred during germination.

***The correlations between i-modulon activity and regulator expression.***
Regulatory proteins are often subject to ligand binding or kinase activity, which switches them between active and inactive states. Therefore, the gene expression of a given transcription factor does not usually correlate with the expression of its targets; this is the case with MalR, which is activated through phosphorylation by MalK only when malate is present[3]. Since i-modulons combine co-regulated gene expression into easy-to-evaluate activity levels, we can attempt to correlate regulator expression with i-modulon activity (Supplementary Methods). As expected, many of these correlations are low (see Dataset S7). For example, MalR activation occurs through a post-transcriptional binary switch, so there is no correlation between MalR gene transcription and i-modulon activity (Supplementary Fig. S6A). One major exception to this is the sigma factors, which are generally only regulated at the expression level. In these cases, we observe much higher correlations between expression and activity, such as with the motility sigma factor, SigD (Supplementary Fig. S6B). High correlations are also observed when the regulator undergoes positive feedback, in which case it is a member of its own i-modulon.

The Thi-box is a riboswitch that is conserved in all domains of life and regulates the expression of genes for thiamine synthesis and transport. In *B. subtilis*, this sequence is upstream of a transcriptional terminator, which it deactivates in the absence of thiamine[6,7]. We would therefore expect the Thi-box sequence to be constitutively expressed, and its downstream genes to respond to thiamine levels – there would be no correlation between Thi-box RNA expression and the activity levels of its genes. Instead, this relationship had a unique shape which was consistent for all 5 Thi-boxes (Supplementary Fig. S6C). We believe that this may be explained by differential degradation of the short Thi-box RNA sequence. Under minimal media conditions, the thi-box sequence does not bind thiamine, so RNA polymerase reads through it and produces a long, relatively stable RNA molecule, which is measured as both high thi-box expression and high thi-box i-modulon activity. Under rich conditions, the binding of thiamine terminates transcription, preventing thi-box i-modulon activity and producing a short, less stable RNA molecule. Interestingly, this short sequence may be degraded quickly in flasks (evidenced by a lack of apparent Thi-box expression) but appears to remain in biofilms for long enough that it could be measured in this experiment. Little is known about differential RNA degradation in biofilms, but this result motivates further study of that phenomenon.

***Two Uncharacterized I-modulons may have important functions.***
The *ndhF-ybcCFHI* operon was identified as its own i-modulon (Fig. 6A). *ndhF* is known to be a subunit of NADH dehydrogenase, but the *ybc* genes have not been characterized at all. Peptide homology (Supplementary Methods) suggests that *ybcC* may form a protein that binds to ndhF, and that *ybcF* may be carbonic anhydrase. The activity levels of this group of genes demonstrate very strong activation under heat shock, as well as repression during cold shock and unusually high germination activity. Perhaps this is a new category of heat-responsive genes; since heat shock is a complex response[8], a small operon like this may have been overlooked in previous studies. We can hypothesize mechanisms through which this operon might benefit the cell: heat shock should upregulate *ndhF* to help to power the heat stress response, *ybcC* might be a chaperone for *ndhF*, and maybe *ybcF* assists in raising the pH after a temperature increase lowers it. We propose gene knockout experiments to validate that these genes play a role in the survival of heat shock.

Another uncharacterized i-modulon is the *yrkEFHI* operon. None of these genes have been characterized, but two of them are putative sulfur carriers, and one is a putative sulfurtransferase. This i-modulon exhibits consistently low activity except in the two conditions with ten or fifteen minutes of diamide exposure (Fig. 6B). Since diamide oxidizes thiols to disulfides, it would make sense for sulfur carriers to be necessary in this condition. Future experiments can be performed to confirm this reasoning and identify transcriptional regulatory mechanisms.

### Supplementary Methods

***Obtaining robust independent components.***
ICA was performed as previously described[9].

Briefly, we processed the $log_2$-transformed, quality-checked, centered data ($\boldsymbol{X}$) with the Scikit-Learn (v0.19.0) implementation of FastICA[10] using 100 iterations, a convergence tolerance of $10^{-7}$, $\log(\cosh(x))$ as the contrast function, and parallel search. We calculated enough components to reconstruct 99% of the variance as determined by PCA.

After 100 iterations of ICA, the M matrices were pooled and clustered with Scikit-Learn DBSCAN[10] (epsilon = 0.1, minimum size = 50) in order to find robust components which appear in each random restart. Since identical components can have opposite signs, we defined distance for this algorithm using a sign-agnostic method:

$$d_{x,y} = 1 - |\rho_{x,y}|$$

where $d_{x,y}$ is the distance and $\rho_{x,y}$ is the Pearson correlation between components $x$ and $y$. Components belong to a cluster if $d_{x,y} < 0.1$ with all other components in the cluster. To ensure repeatability, all signs in a cluster were inverted if necessary so that the highest weighted gene would have a positive sign. The centroids of each cluster defined the weightings in $\boldsymbol{M}$ and were used to calculate $\boldsymbol{A}$.

This process was repeated 100 times (for a total of 10,000 ICA runs), and components that did not arise in every run were discarded. The result contained 83 robust components.

***I-modulon Threshold Determination.***
The distribution of $\boldsymbol{M}$ matrix weights of each gene for a given component consists of a large number of near-zero values along with a small number of genes at the tails. To define the gene set of the i-modulon, we need to choose a threshold value that separates the normally distributed near-zero genes from the more meaningful, non-gaussian tails. To do so, we used Scikit Learn's implementation of the D'Agostino $K^2$ test, which quantifies the skew and kurtosis of the distribution as a measure of gaussianity[10,11]. We iteratively remove the gene with the highest absolute value in the component and calculate the $K^2$ value until the value falls below a $K^2$ cutoff value (1300). All genes that were removed are members of the i-modulon gene set, and the non-removed genes are sufficiently normally distributed around zero to be considered noise. In all cases discussed in this study, all member genes have positive weights, which allows for easier representation as a set of genes and a simple interpretation of activity.

The cutoff value of 1300 was determined by a sensitivity analysis. Over a range of cutoffs (200 - 2200), we computed the top regulator enrichments and F1 scores as described in the following section. The cutoff with the highest mean F1 score was selected. In seven cases (Dataset S3), this cutoff was not appropriate because it removed all genes from the I-modulon (5/7) or captured many non-important genes (2/7), so the threshold was adjusted to 500 *or increased slightly as necessary.

***Explained Variance Calculation.***
We reconstructed the dataset using only the i-modulons. To do this, we first define a binary matrix $\boldsymbol{M_{bin}}$, [genes]x[i-modulons] (Dataset S5), whose elements are 1 if the row's corresponding gene is in the column's corresponding i-modulon, 0 otherwise. We compute $\boldsymbol{M'}$ using element-wise multiplication $\boldsymbol{M} * \boldsymbol{M_{bin}}$, which removes the effect of all non-significant genes from $\boldsymbol{M}$. We then define $\boldsymbol{X'} = \boldsymbol{M'} * \boldsymbol{A}$, using matrix multiplication to reconstruct the data. We used the scikit-learn explained variance score function[10] to obtain a total explained variance of 72% (Supplementary Fig. S1B).

***Correlation Analysis.***
I-modulon activities were compared to regulator gene expression values using scatter plots and a best fit line. The best fit lines are composed of two parts:

$y = a * c + b$ if $x < c$

$y = a * x + b$ if $x \geq c$

where $x$ is the expression level of the regulator, $y$ is the i-modulon activity level, and $a$, $b$, and $c$ are fitting parameters determined by the optimize.curve_fit function in the Python sciPy package[12]. The flat part of the curve, defined by the first equation, represents the minimal activity level required before a correlation is observed; it does not exist for all correlations. The adjusted $R^2$ value accounts for the $k$ parameters used ($k = 2$ or $k = 3$ depending on whether or not the line required both parts) based on the following equation:

$$R_{adj}^2 = 1 - (1 - R^2)(n - 1)/(n - k - 1)$$

where $R^2$ is the coefficient of determination.

***Protein Homology.***
Potential functions of uncharacterized genes were identified by hhblits[13]. Gene sequences were obtained using the genome sequence from GenBank Accession AL009126.3[14]. Gene locations were found in the *Subti*Wiki gene annotations[15], and translated into peptides using Bio Entrez[16]. Peptide sequences were compared to those in the Uniclust30 database[17] with hhblits. This produced detailed output for each gene, from which a homology probability of 95% or above was required for all stated conclusions.

**Supplementary Figures**

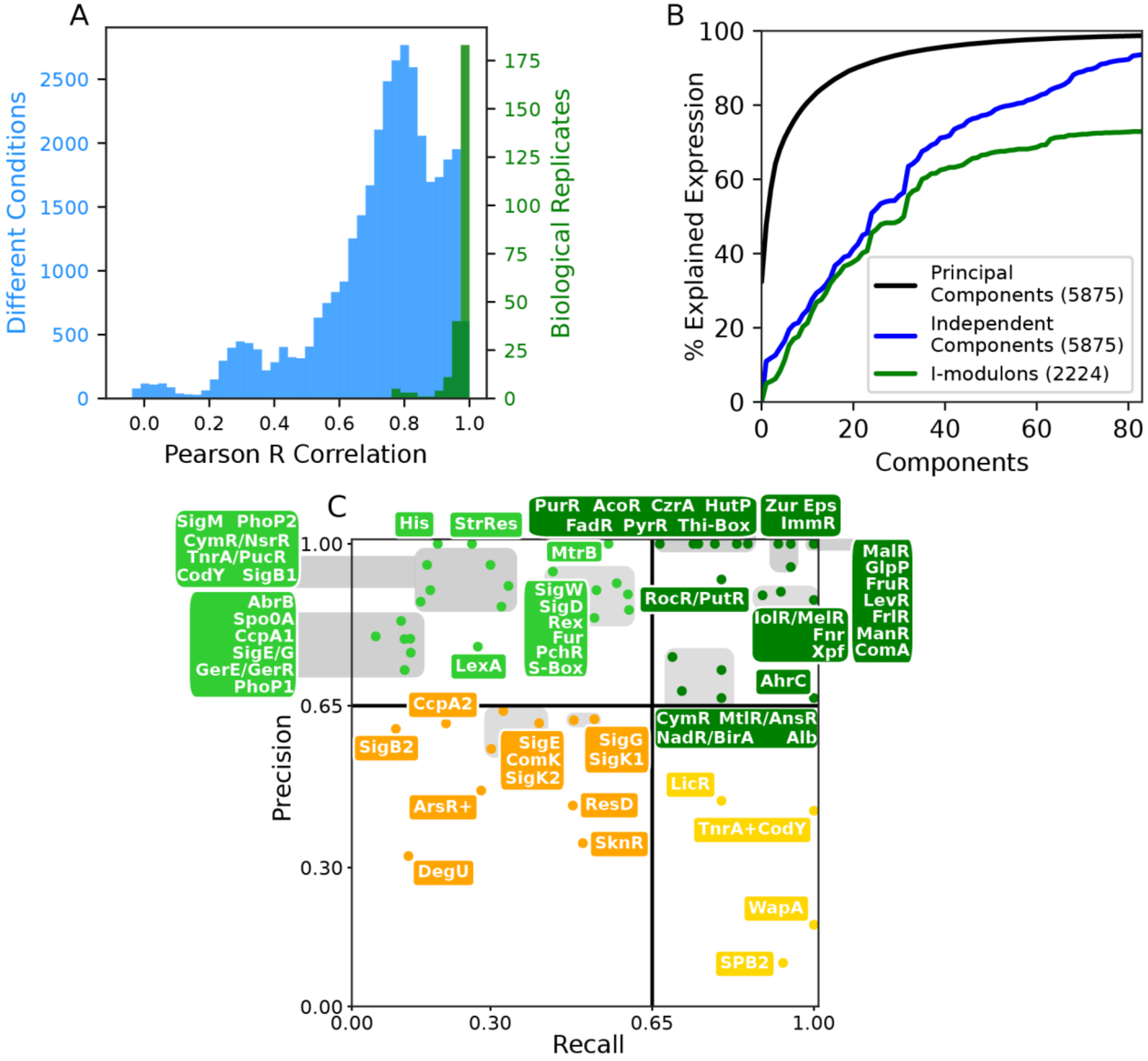

***Supplementary Fig. S1: Overview of the data***

**a.** Histogram of Pearson R correlations in gene expression between different conditions (blue) and biological replicates (green). High correlation between replicates indicates high quality data. Three noisy samples were removed from the initial dataset to enforce this behavior. **b.** Variance in the expression data using PCA, the full $\boldsymbol{M}$ and $\boldsymbol{A}$ matrices (independent components) and the thresholded $\boldsymbol{M}$ matrix (I-modulons). Parentheses in the legend indicate the number of genes used for the backprojection of the original data. While PCA explains more expression quantitatively, the components it obtains are less biologically interpretable than those of ICA. Thresholding the independent components to obtain I-modulons causes a loss of only ~20% of the explained variance, which indicates that most of the variance captured by ICA is in the i-modulon member genes. **c.** Labeled version of Main Fig. 1C, with I-modulon short names shown near their precision/recall coordinates. When space required that labels be grouped together, the relative positions of the label and point were maintained as best as possible (e.g. vertically listed names correspond to each point in order of decreasing precision).

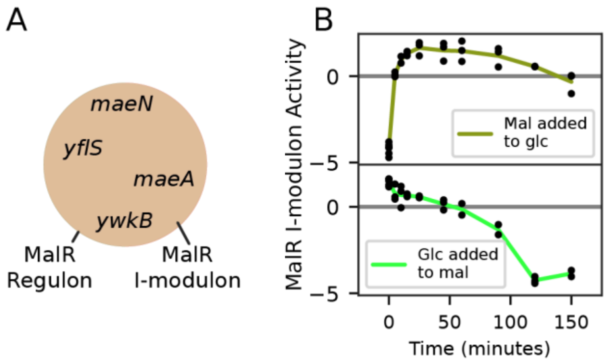

***Supplementary Fig. S2: The MalR i-modulon***
**a.** Venn diagram of the MalR i-modulon and MalR regulon. Precision and recall are 100%. **b.** MalR i-modulon activity over time for malate addition to glucose media (top) and glucose addition to malate media (bottom). Activity falls slowly after glucose addition (from approximately 0 to -5) and rises rapidly upon malate addition (from -5 to approximately 0).

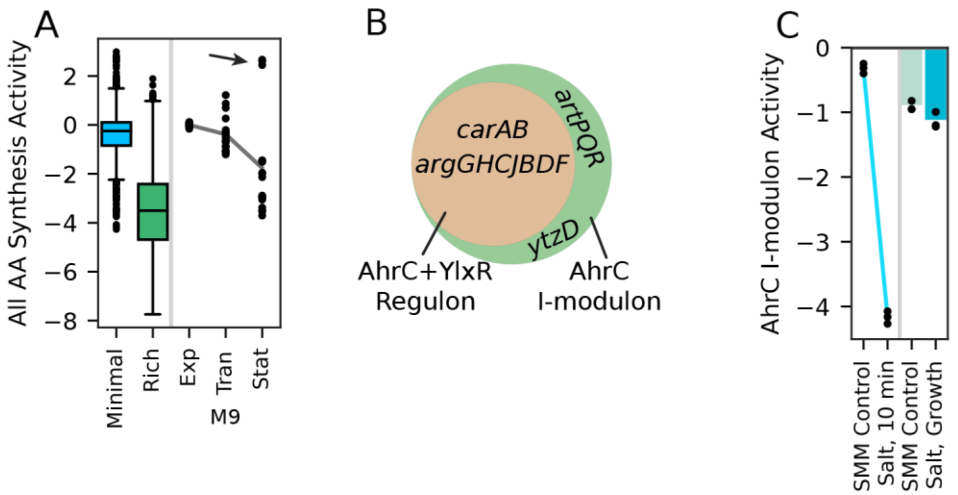

***Supplementary Fig. S3: Supplementary amino acid i-modulon plots***

**a.** Box plot and time course of all 6 amino acid synthesis i-modulons in media with and without casamino acids, and over three growth phases in M9 media. 'Exp', 'Tran' and 'Stat' refer to exponential, transition, and stationary phase, respectively. The data set is normalized to minimal media, thus the blue box centers at 0, while the rich media (green box) has a negative value, showing lower activity relative to the reference condition. Activity follows expected trends based on amino acid stimulus presence. Outliers (arrow) for M9 growth are from the CodY i-modulon, the only synthesis i-modulon which increased over this time course. **b-c.** The arginine synthesis (AhrC) i-modulon. **b.** Venn diagram of the arginine synthesis (AhrC) i-modulon and regulon; the regulon contains additional arginine-related genes (*artPQR* and *ytzD*) that are not known to be regulated by AhrC. **c.** AhrC i-modulon activity in osmotic stress conditions. This i-modulon is surprisingly downregulated by salt shock.

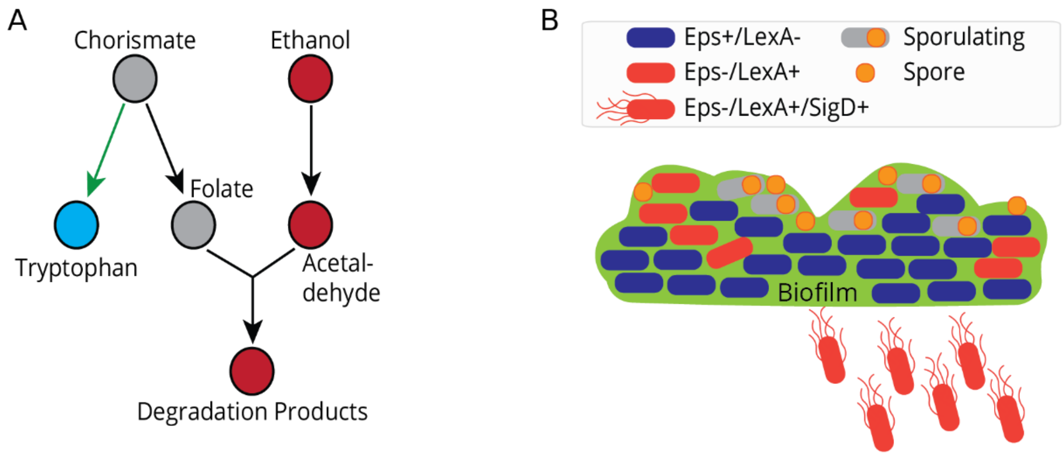

***Supplementary Fig. S4: Graphical representations of hypotheses.***
**a.** Potential pathway for tryptophan loss in the media during ethanol stress (see Main Fig. 2D), with intermediates removed. In the presence of ethanol, flux from chorismate may be diverted to replenish degraded folate. **b.** Cross-section of biofilm morphology, with the cellular switches between biofilm (Eps, exopolymeric substances) production, LexA (DNA damage response) expression, and sporulation. Swarming cells (SigD+) are likely to only arise from LexA+, Eps- cells (see Main Fig. 2H)

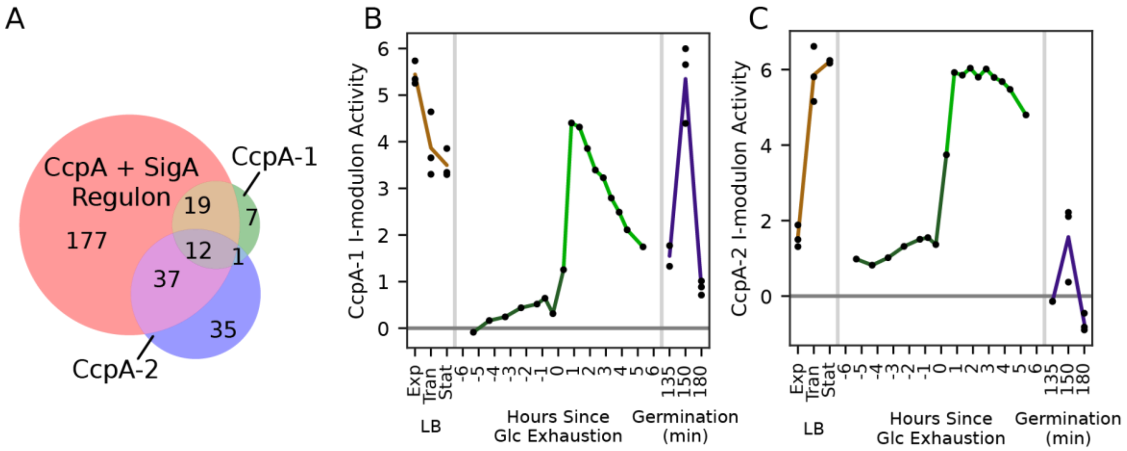

***Supplementary Fig. S5: The CcpA i-modulons***

CcpA-1 contains mostly sugar metabolism enzymes (ribose, sucrose, mannose, trehalose, lichenan, etc.) while CcpA-2 contains a mix of genes including those for inositol consumption, tricarboxylic acid permeability, and acetyl-CoA utilization (Full set of genes: Dataset S5). **a.** Venn diagram of gene membership for these I-modulons and their matched regulon. **b-c.** Activity of CcpA i-modulons for three experiments: growth in LB media (‘Exp’, ‘Tran’ and ‘Stat’ refer to exponential, transition, and stationary phase, respectively), glucose exhaustion, and germination. Dots indicate individual samples and lines pass through means. **b.** CcpA-1 is active during exponential growth and germination, but declines in stationary phase and during glucose exhaustion. **c.** CcpA-2 is active during stationary phase and throughout the first 5 hours of glucose exhaustion, and comparatively less active during germination.

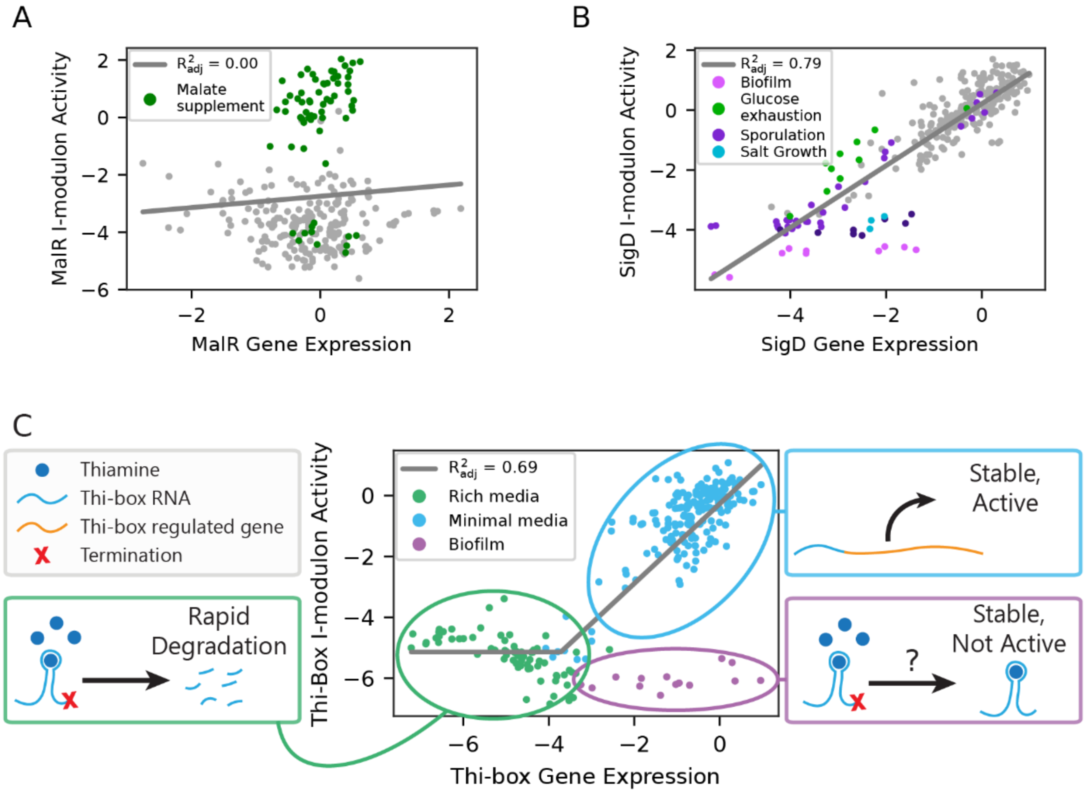

***Supplementary Fig. S6: Correlation between i-modulon activity and regulator expression***
Each plot contains an i-modulon's transcriptional regulator expression on the x axis and the corresponding activity level on the y axis. **a.** MalR is a typical transcription factor that responds to kinase activity downstream of malate binding. The activity is therefore not correlated with regulator expression but is increased with malate supplementation. **b.** SigD is the sigma factor governing motility, which is regulated at the transcriptional level, so a correlation between activity and expression is observed. Four experiments that exhibit low activity are highlighted; their activity matches expectations from literature. **c.** This Thi-box transcript precedes *thiC*, and the other 4 Thi-box transcripts exhibit similar patterns. A broken line was used for this regression (Supplementary Methods). When activity is low, the expressed RNA contains only the short thi-box sequence, which appears to be degraded quickly in the rich media condition (flasks) but not in the biofilm condition. The thi-box RNA appears to be stable in biofilms for an unknown reason.

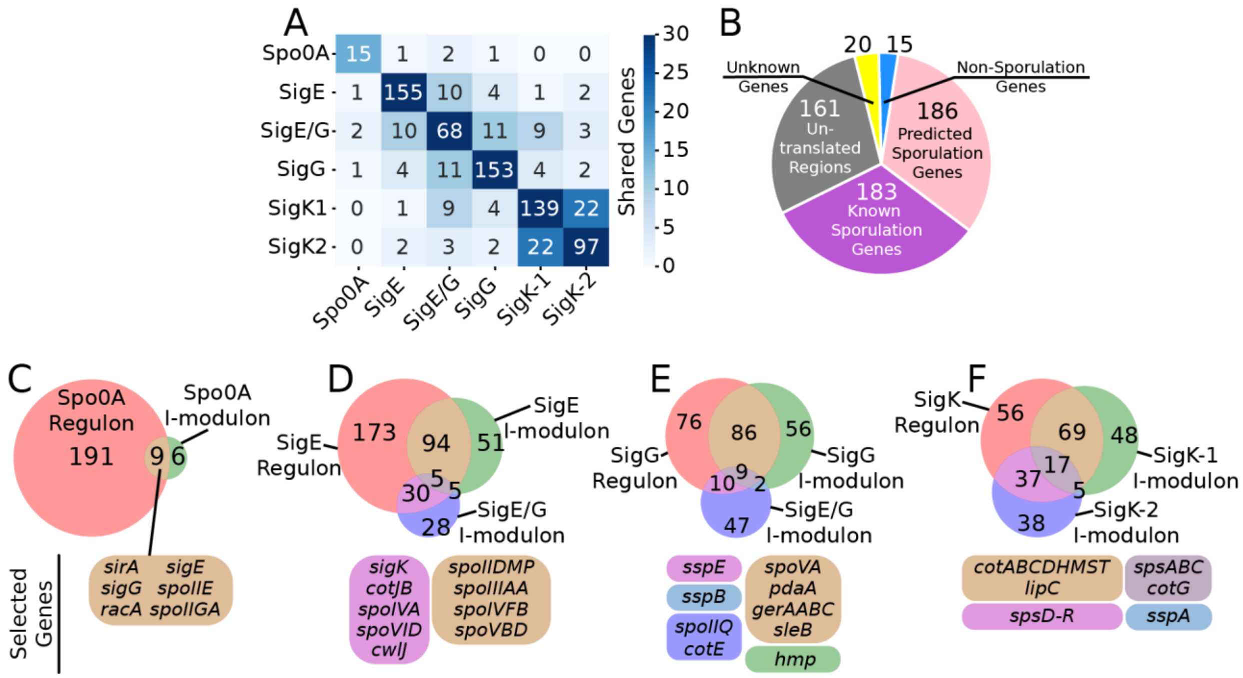

***Supplementary Fig. S7: The genes in sporulation i-modulons***
**a.** Heatmap displaying the number of shared genes (off-diagonal) and self-genes (diagonal) for each i-modulon. Many genes are shared between i-modulons. **b.** A pie chart of gene type for all transcripts in any of the sporulation i-modulons. **c-f.** Venn diagrams of known regulons and relevant i-modulons for each of the enriched regulators. Boxes list important sporulation genes that are present in the subset with a matching color. The genes listed have functions that include transcriptional/post-translational regulation, chromosome segregation, spore coat production, DNA protection, germination initiation, and mother cell lysis.

**Supplementary Dataset Legends**

***Supplementary Dataset S1: Growth Conditions***

**File:** SI_Data_S1-5.xlsx

**Sheet:** "S1 - Growth Conditions"

Dataset S1 contains a list of all conditions used, which were generated by Nicolas, *et al.*[18]. We removed three noisy samples (see Methods). The columns describe the following:

- *Sample_n:* sample number
- *Condition:* name of the condition
- *Experiment:* name of the experiment. Activity plots in the main figures contain gray lines separating unique experiments.
- *Experiment_description:* Description of the experiment and conditions, copied from the original dataset
- *Cells_collected:* description of collected cells, copied from the original dataset
- *X_label:* the label used for the condition in the activity plots in the main figures, if applicable
- *Exp_label:* the experiment label used (appears floating below the *x_label*) in the activity plots, if applicable
- *Control:* Most relevant control group for this sample, if applicable. Used to determine statistical significance in the activity plots for dataset S6.
- *Color:* Hex code for the condition's color, used for plots
- *Exp_color:* Hex code for the experiment's color, used for time course plots
- *Ref:* Reference that generated the data. Parentheses indicate specific members of the original group that rain the experiments.

***Supplementary Dataset S2: Expression Data (X)***

**File:** SI_Data_S1-5.xlsx

**Sheet:** "S2 - X - Expression Data"

For unprocessed expression data, see the original dataset[18]. Rows indicate genes and non-coding RNAs, and columns indicate samples. Each value is a log-transformed microarray expression value that has been centered such that the mean of the baseline condition (M9_exp) for all genes is 0.

***Supplementary Dataset S3: TRN Structure (M)***

**File:** SI_Data_S1-5.xlsx

**Sheet:** "S3 - M - TRN Structure"

This matrix links the genes to their corresponding i-modulons. Columns indicate i-modulons (see Dataset S7 for a description of each i-modulon), rows indicate genes or non-coding RNAs. The values represent the weight of each gene in determining the activity of each i-modulon. The top two rows are thresholds: The top row was automatically computed (Methods), and the second row was curated in seven cases to retain genes (if the threshold was decreased) or remove excessive noise from an otherwise meaningful i-modulon (if the threshold was increased). All genes with an absolute value in the column greater than the curated threshold are considered to be members of the i-modulon.

***Supplementary Dataset S4: Activity (A)***

**File:** SI_Data_S1-5.xlsx

**Sheet:** “S4 - A - Activity”

This matrix contains the condition-dependent activity of each i-modulon. Rows indicate i-modulons (Dataset S7) and columns indicate conditions (Dataset S1). Values indicate how active the i-modulon is in the condition, with zero indicating the activity in the baseline condition (M9exp) and each unit being approximately one log(expression value) if the i-modulon had only one gene.

***Supplementary Dataset S5: I-modulon Presence***

**File:** SI_Data_S1-5.xlsx

**Sheet:** “S5 - I-modulon Presence”

Rows are genes and non-coding RNAs and columns are i-modulons; a 1 indicates that the gene is in the i-modulon and a 0 indicates that it is not. This is simply a thresholded version of the **M** matrix (Dataset S3).

***Supplementary Dataset S6: I-modulon Dashboards***

**File:** SI_Data_S6_I-modulon_Dashboards.pdf

This PDF contains automatically generated summaries of each i-modulon. Note that the gene lists and counts here will include non-coding RNAs, which were usually omitted for simplicity in the main text.

- *Title:* n - Short name - long name. N corresponds to the i-modulon number in datasets S3-S5
- *Biological function:* brief description of the function of the i-modulon’s genes
- *Regulon:* includes the category that the i-modulon falls into (see Main Fig. 1C) and the string of regulators. The regulator may be a boolean combination with ‘/’ denoting union of regulons and ‘+’ denoting intersection.
- *Plot 1:* Scatter plot of mean gene expression (dataset S2) and i-modulon gene weight (dataset S3), with horizontal lines indicating the weight threshold and colors indicating gene category annotations from *Subti*Wiki. Gene categories of i-modulon member genes are listed in the legend, with the number of member genes in each category in parentheses.
- *Plot 2:* Semi-log histogram of gene weights. Regulated genes are colored as shown in the legend, and member genes are listed above the appropriate bars.
- *Plot 3:* Activity level of the i-modulon across all conditions (mean ± standard deviation). Stars indicate statistically significant conditions relative to their matched control (FDR < 0.05, dataset S7, Methods), and shaded backgrounds indicate significant correlation with time (Pearson R > 0.8, FDR < 0.05).
- *Plot 4:* Venn diagram of the known regulon (red), the annotated i-modulon genes (green), and the unannotated i-modulon genes (blue). Numbers indicate the size of the subset.
- *Plot 5:* Scatter plot(s) of regulator expression and i-modulon activity (see Figure 3D-F). A best fit line and adjusted $R^2$ value is shown. If there are multiple regulators, the adjusted $R^2$ was computed for all of them and the top 3 regulators are shown, sorted with the highest correlation on the left. Colors of points match those in plot 3 and are listed in dataset S1.

***Supplementary Dataset S7: Summary table of i-modulons***

**File:** SI_Data_S7.xlsx

Numbering is consistent with datasets S3-S6.

- *Regulator(s):* All regulator annotations were downloaded from *Subti*Wiki[15]. Where multiple regulators are listed, ‘+’ indicates the intersection of regulons, and ‘/’ with brackets indicates the union of separate regulons.
- Top Activity-TF $R^2_{adj}$: The $R^2$ correlation between the i-modulon activity and its matched regulator, adjusted for the minimum activity level before a correlation is observed, is given. If there are multiple regulators, all $R^2_{adj}$ were computed and the maximum is reported. See Supplementary Methods.

***Supplementary Dataset S8: Uncharacterized and divergently regulated genes***

**File:** SI_Data_S8.xlsx

Genes were included if they are coding sequences with their description, product, or function annotated as “unknown” on *Subti*Wiki, or if they are not known to be regulated by the I-modulon’s regulator. These genes and gene/regulator relationships are opportunities for discovery. For the complete list of genes in any i-modulon, use the i-modulon presence matrix, dataset S5.

***Supplementary Dataset S9: Activating Conditions***

**File:** SI_Data_S9.xlsx

Expected and unexpected activating conditions for characterized i-modulons. Expectations are based on the known mechanism of the enriched regulator combination and the expected function of the i-modulon. Elucidating the mechanisms of unexpected activation is a potential path for discovery.
