## Supplemental Dataset S6 for "Machine learning uncovers independently regulated modules in the *Bacillus subtilis* transcriptome"

### 0 - FadR – Fatty Acids

Biological Function:
Fatty acid degradation

Well-defined regulon:
FadR

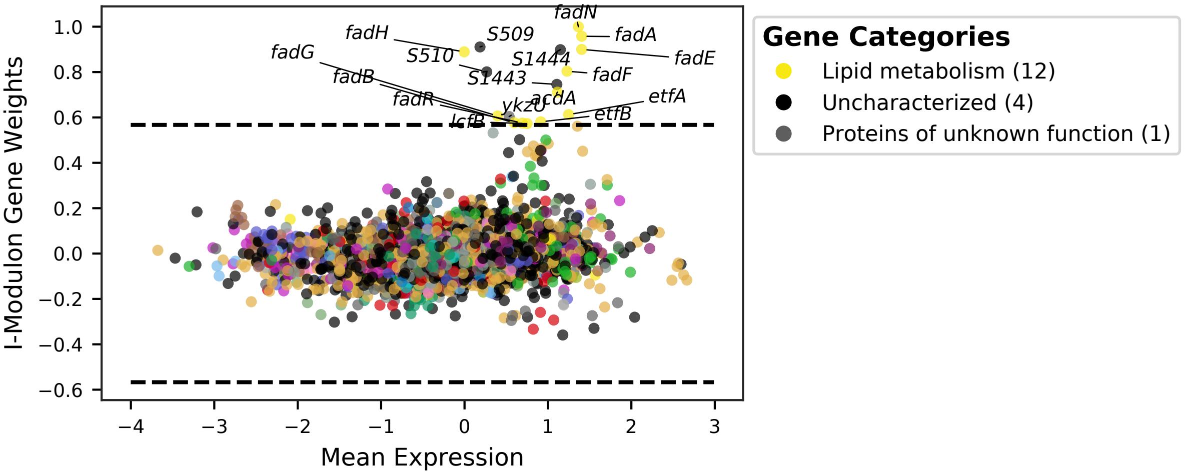


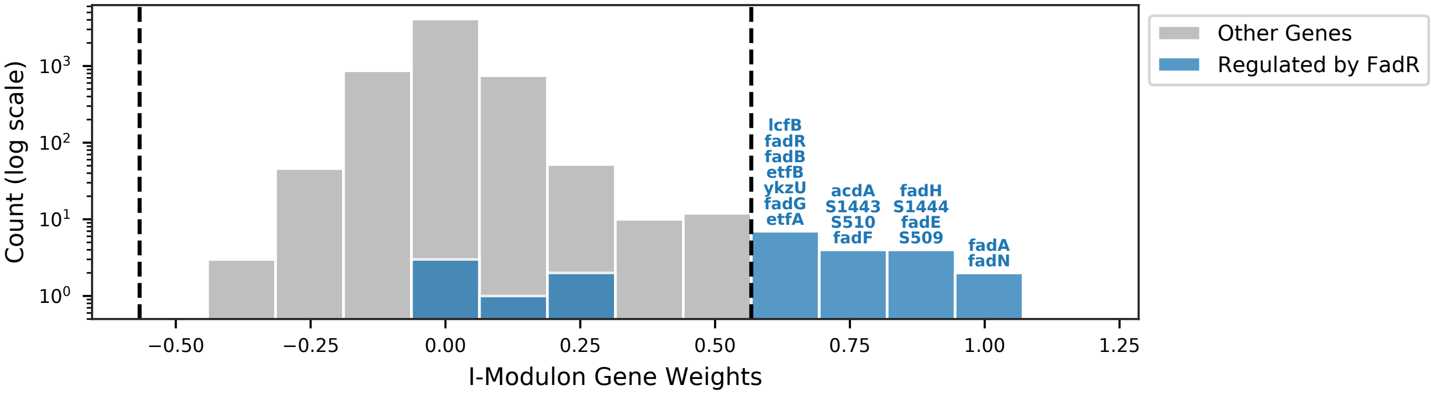


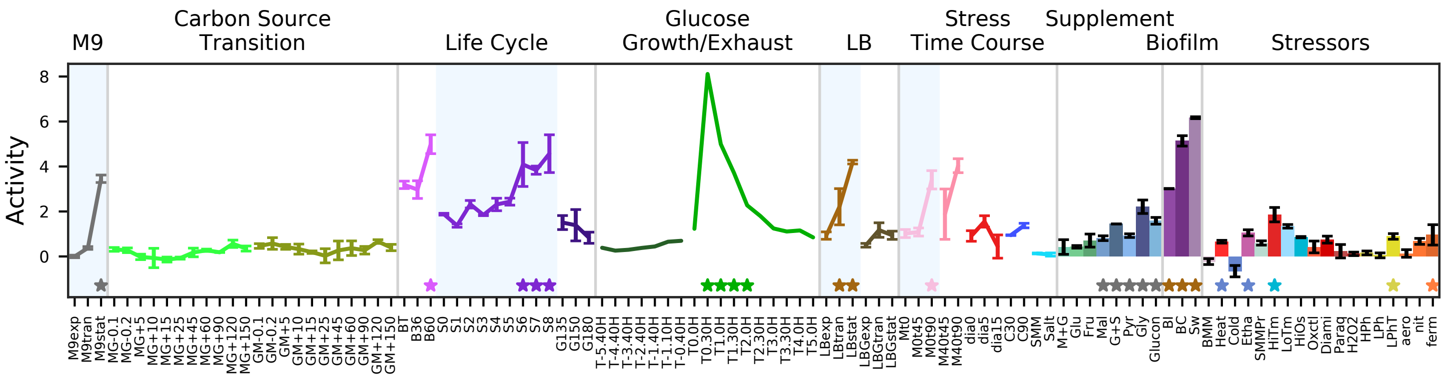


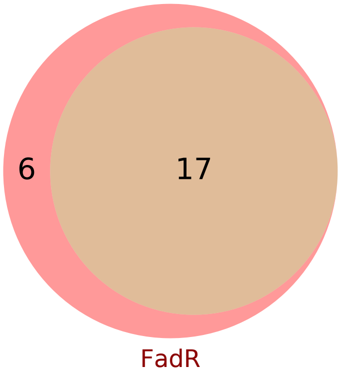


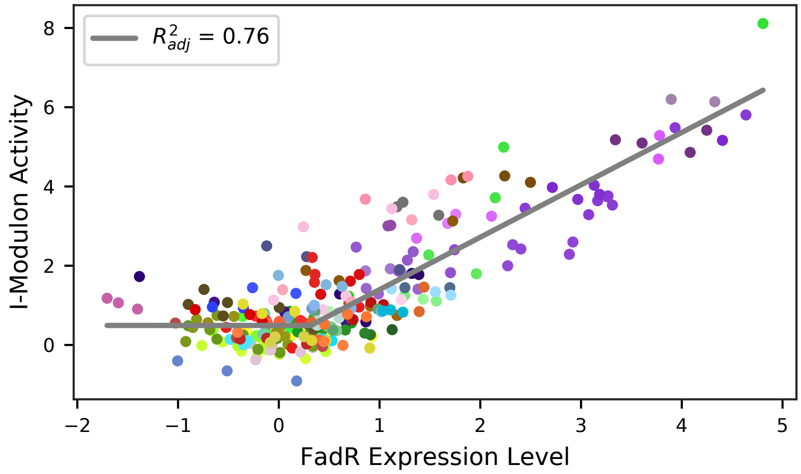

### 1 - MalR - Malate

Biological Function:
Malate uptake and utilization

Well-defined regulon:
MalR

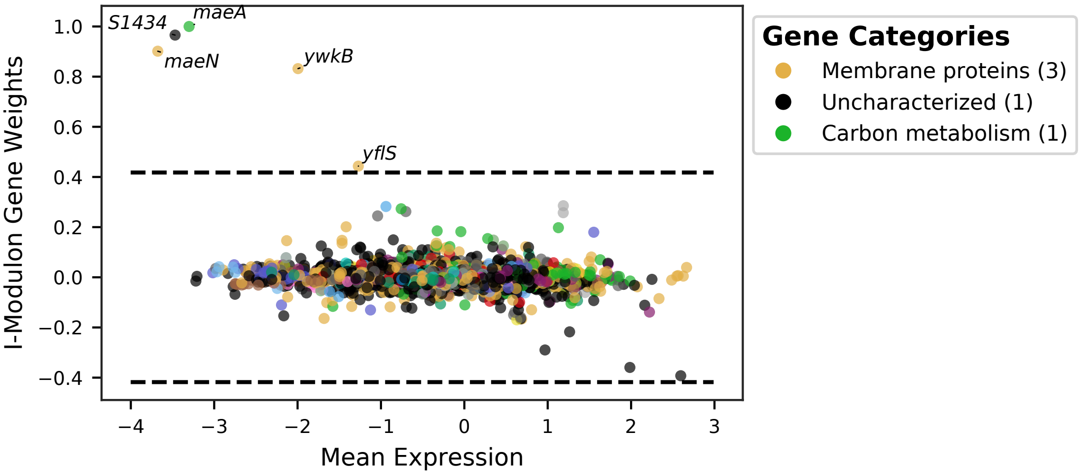


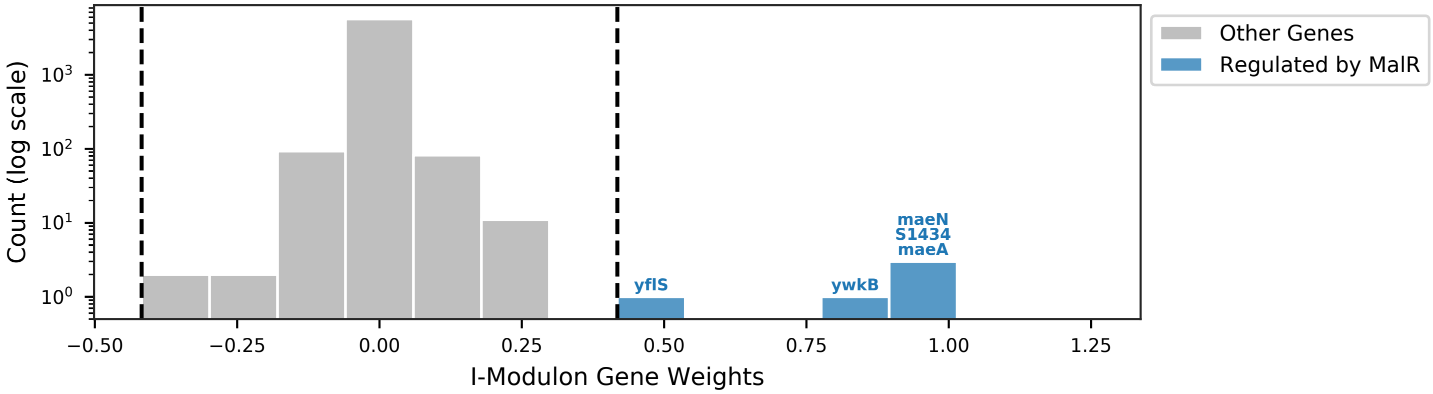


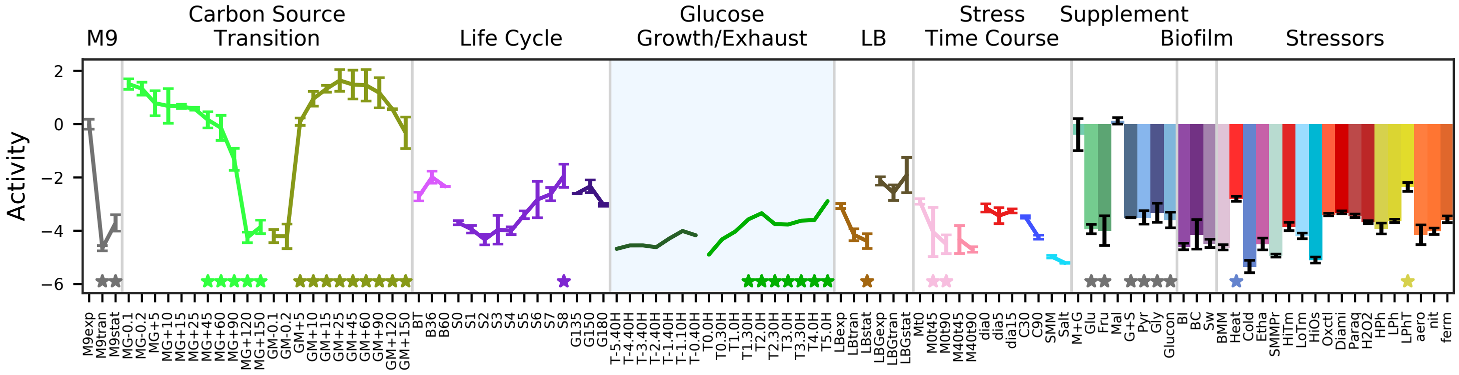


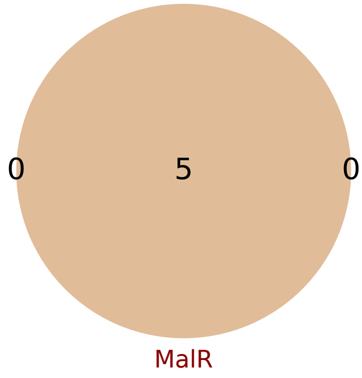


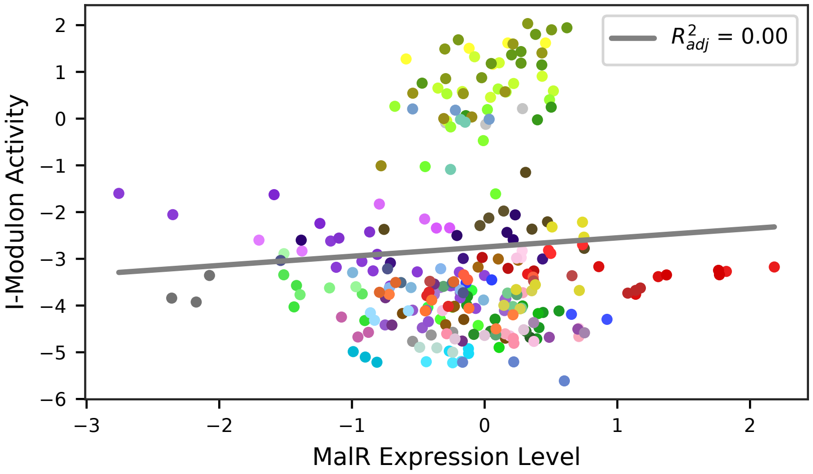

### 2 - GlpP – Glycerol

Biological Function:
Glycerol uptake and utilization

Well-defined regulon:
GlpP

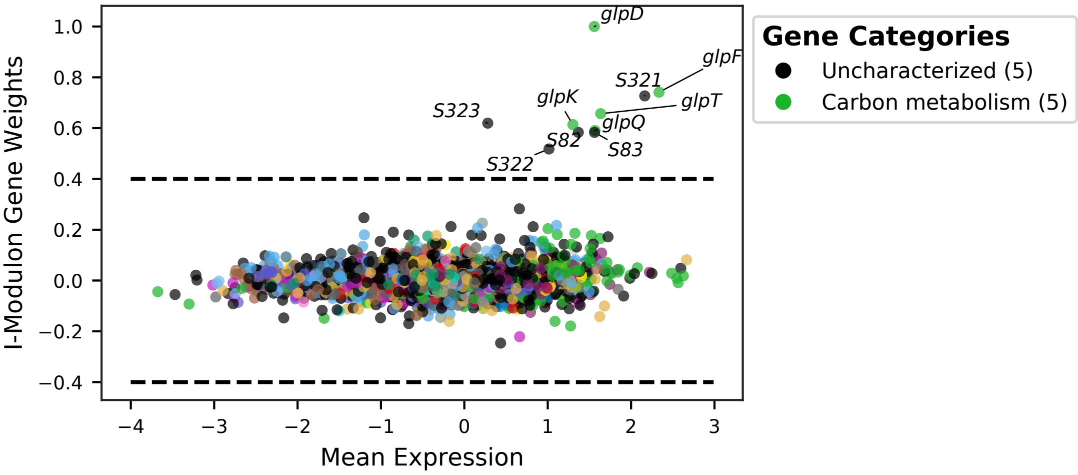


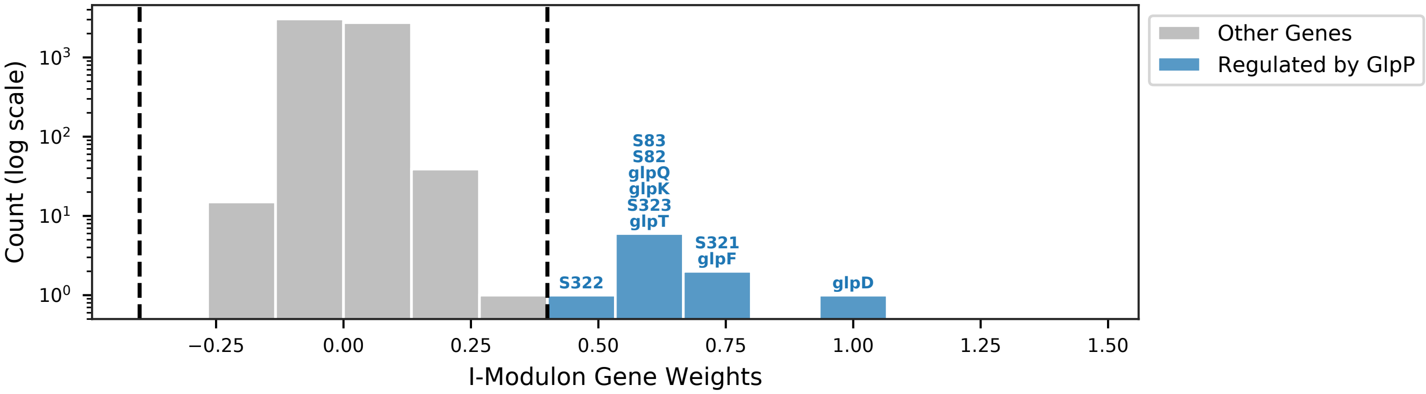


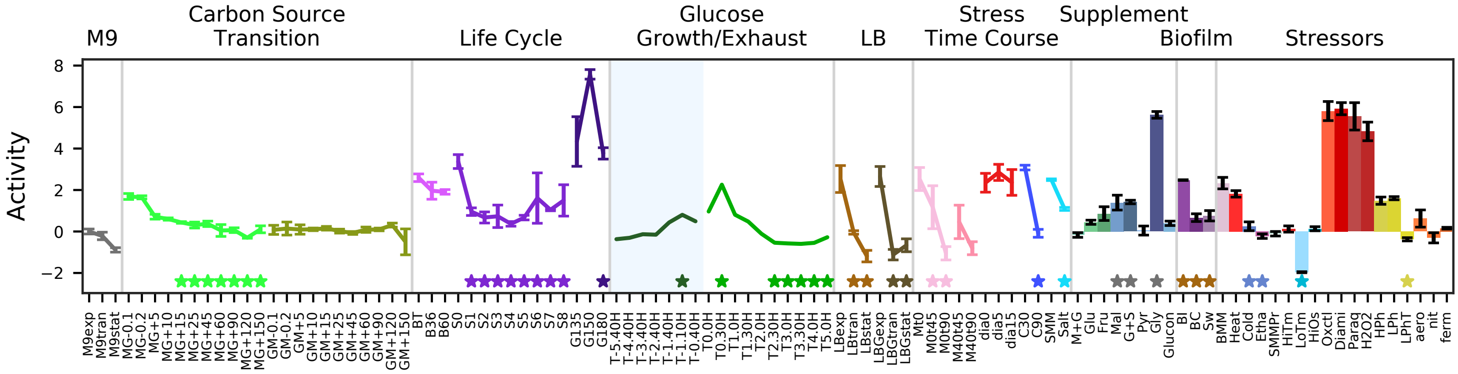


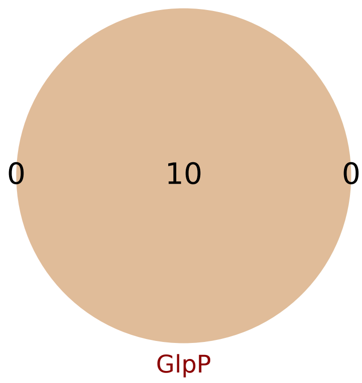


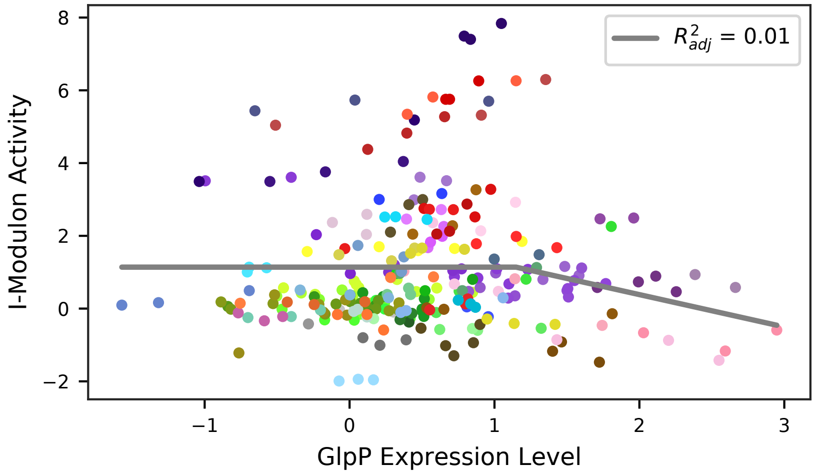

### 3 - FruR – Fructose

Biological Function:
Fructose uptake and utilization

Well-defined regulon:
FruR

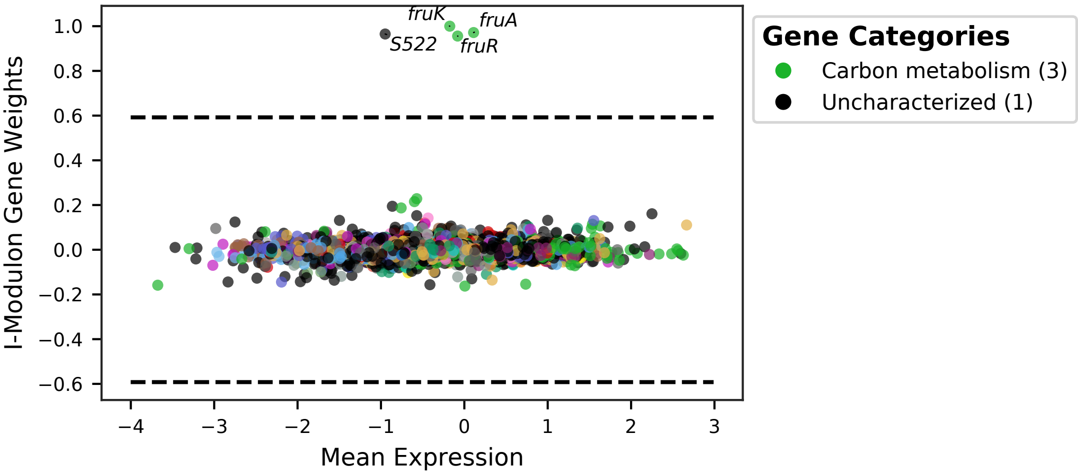


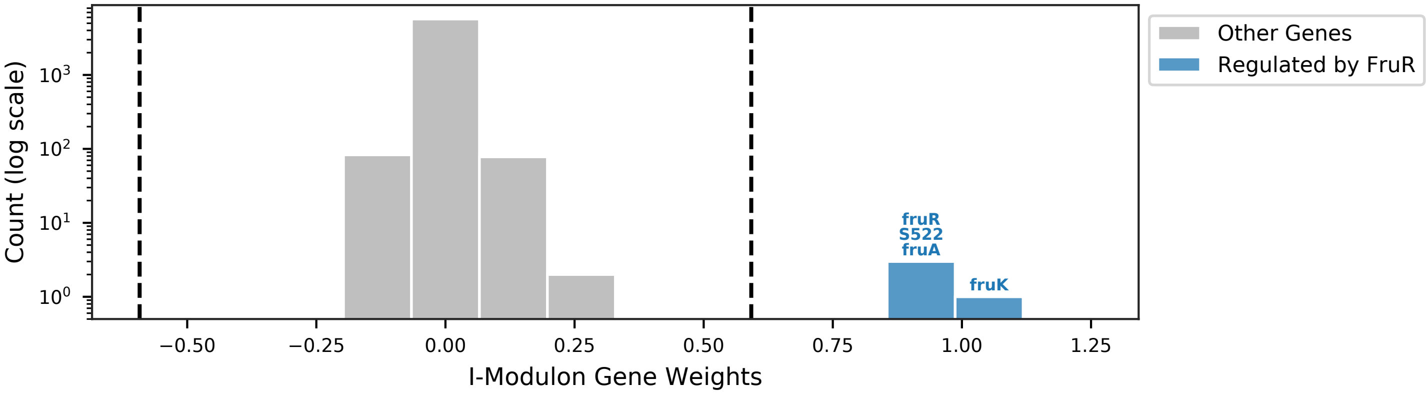


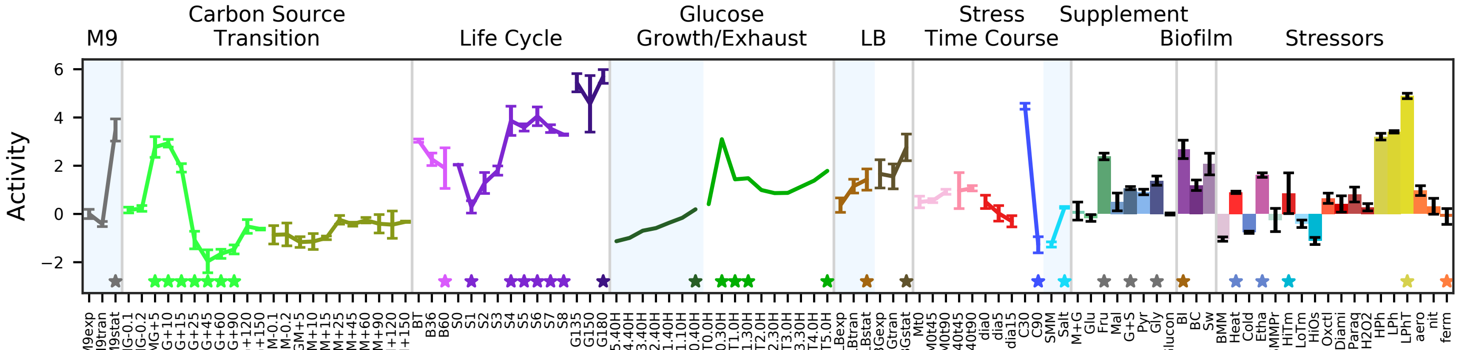


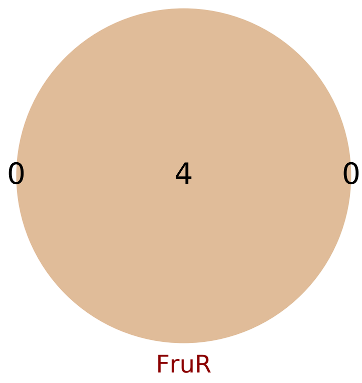


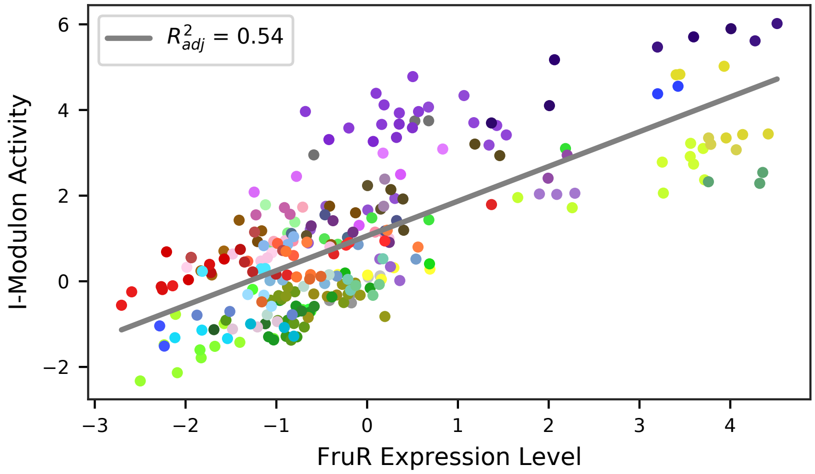

### 4 - LevR – Levan and Fructose

Biological Function:
Fuctose uptake and phosphorylation, degradation of the exopolysaccharide levan into fructose

Well-defined regulon:
LevR

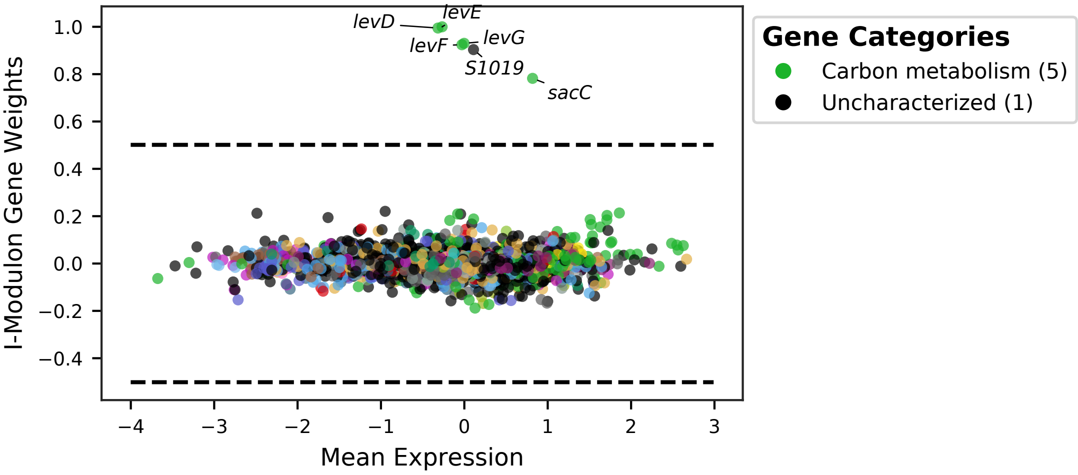


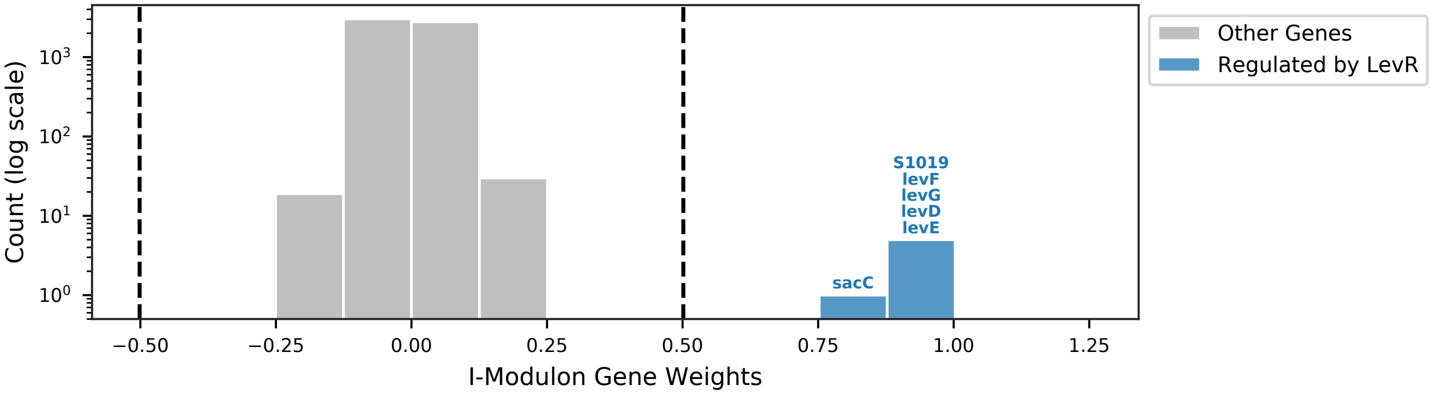


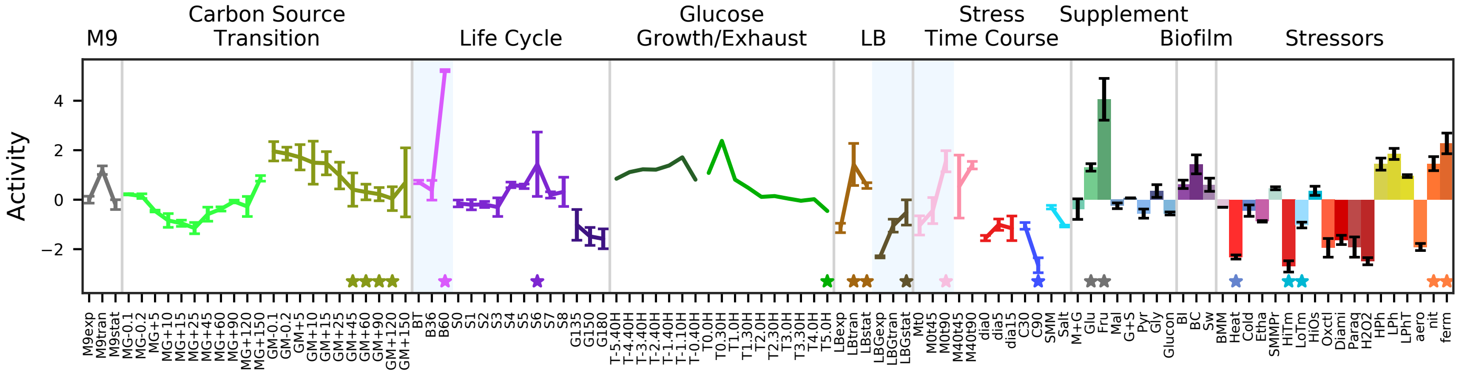


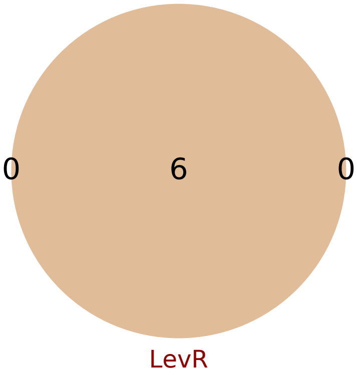


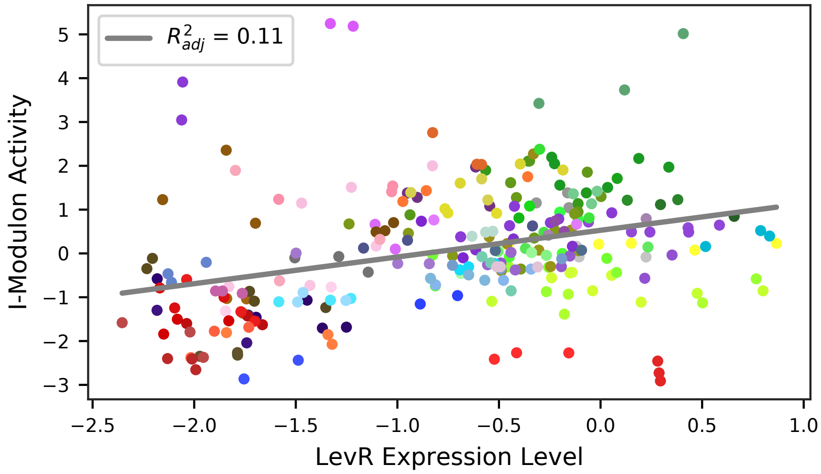

### 5 - FrlR – Amino Sugars

Biological Function:
Amino sugar uptake and metabolism

Well-defined regulon:
FrlR

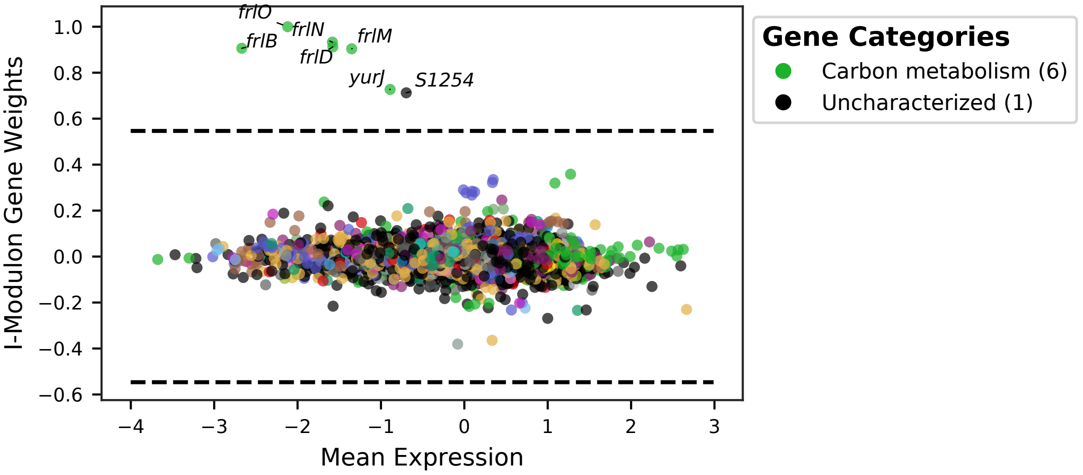


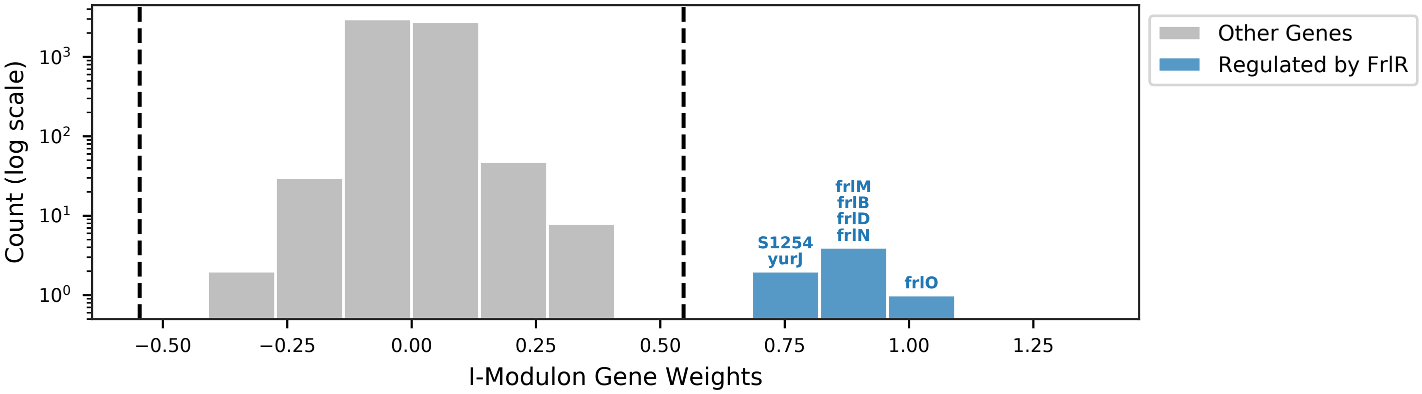


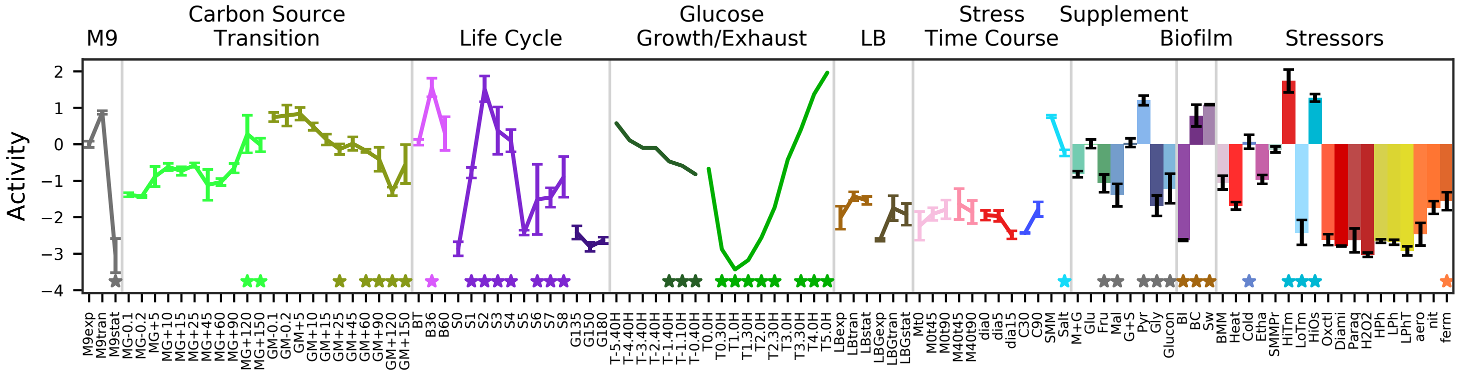


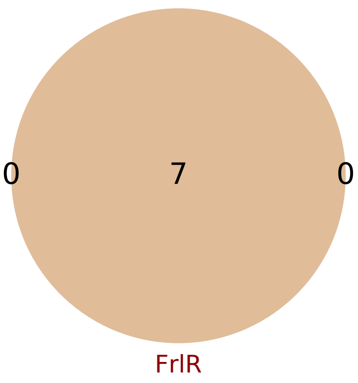


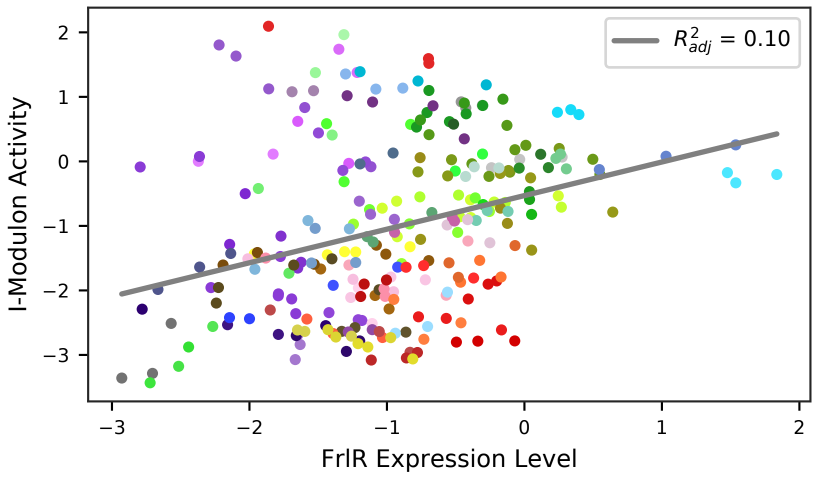

### 6 - ManR – Mannose

Biological Function:
Utilization of mannose for cell wall synthesis. Needed for exponential growth in LB media (absent glucose) or in the presence of fructose

Well-defined regulon:
ManR

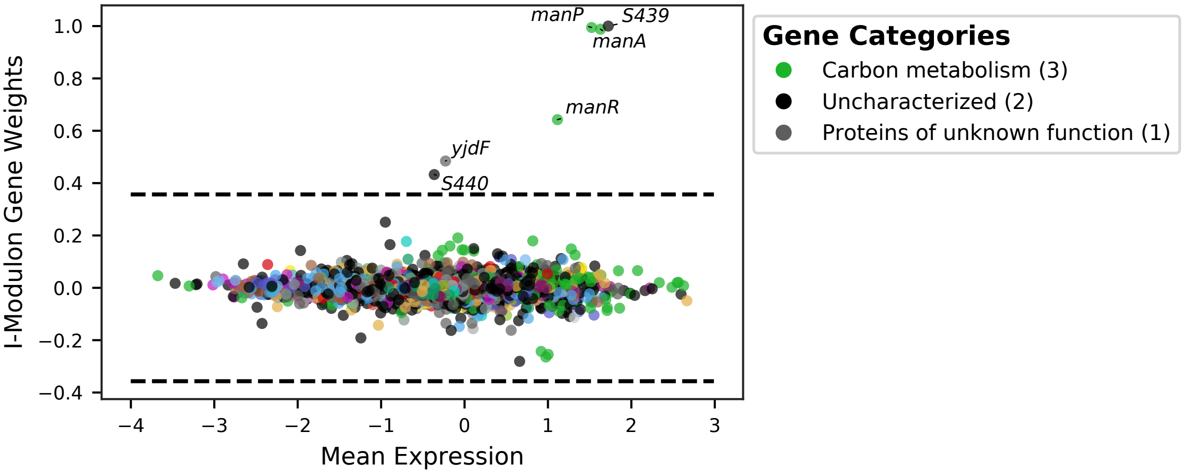


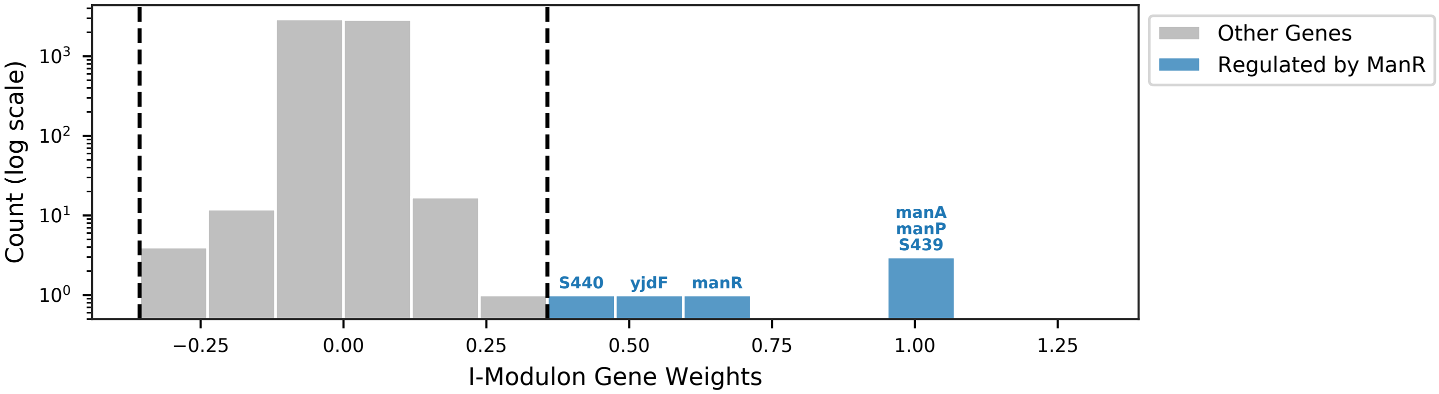


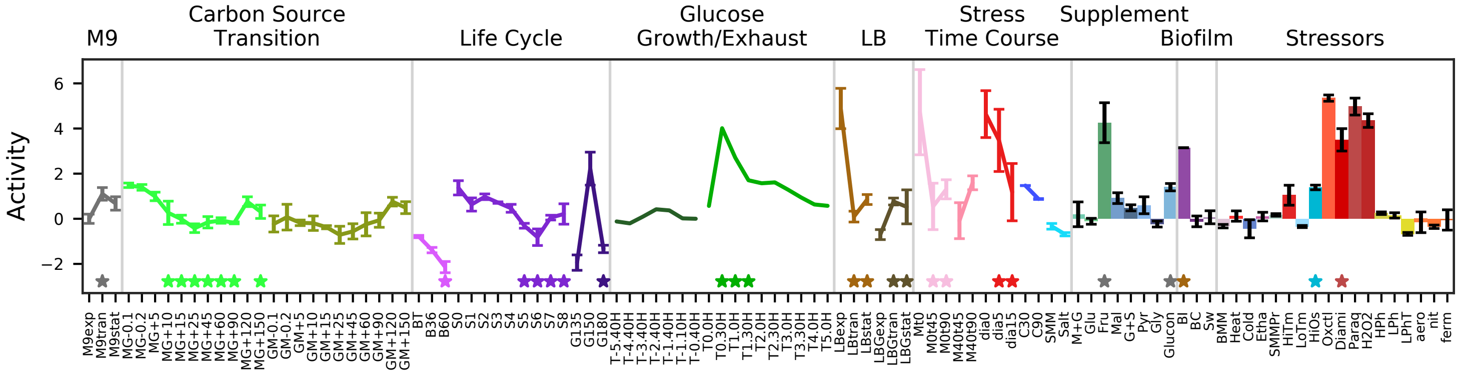


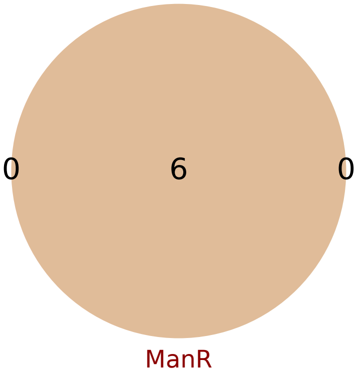


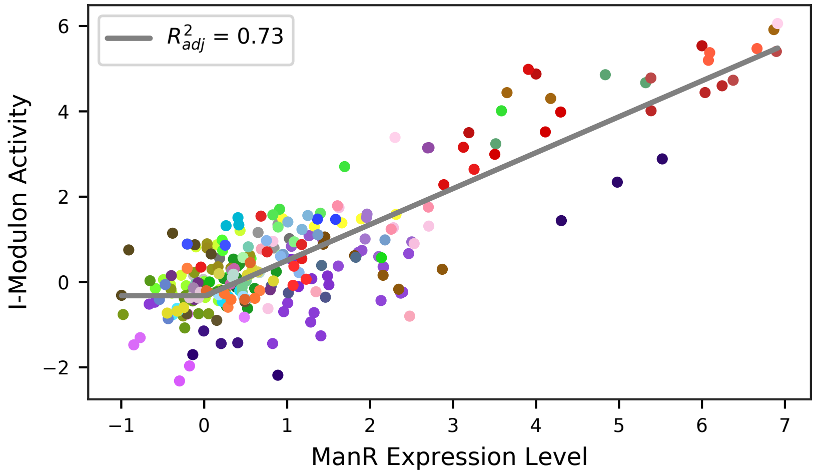

### 7 - AcoR – Acetoin

Biological Function:
Utilization of acetoin, extracellular energy storage

Well-defined regulon:
AcoR

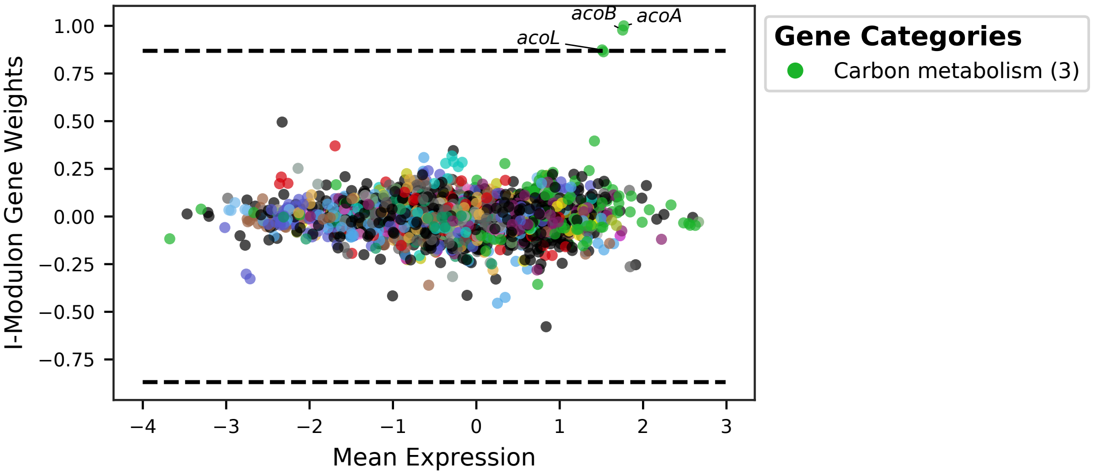


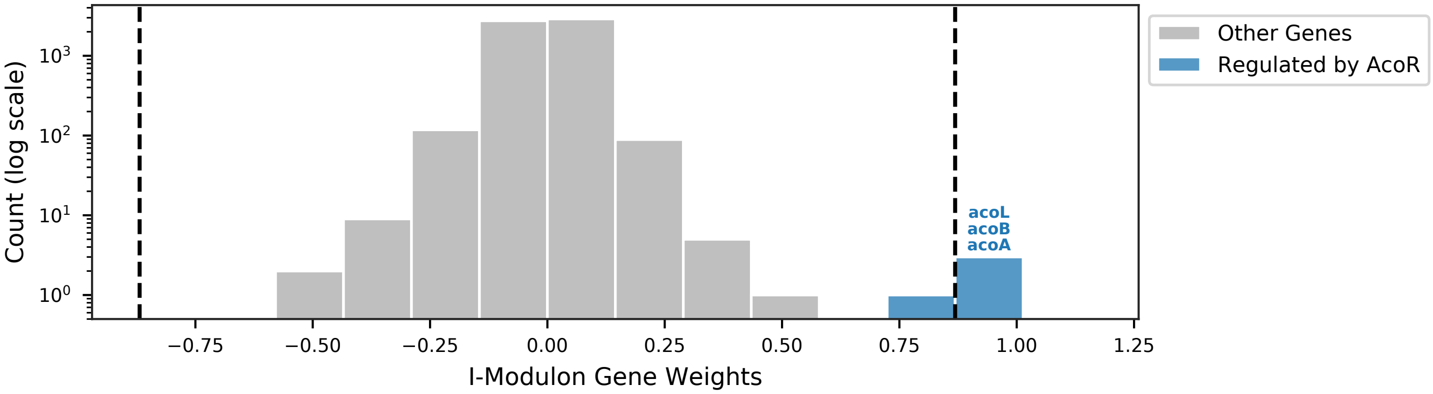


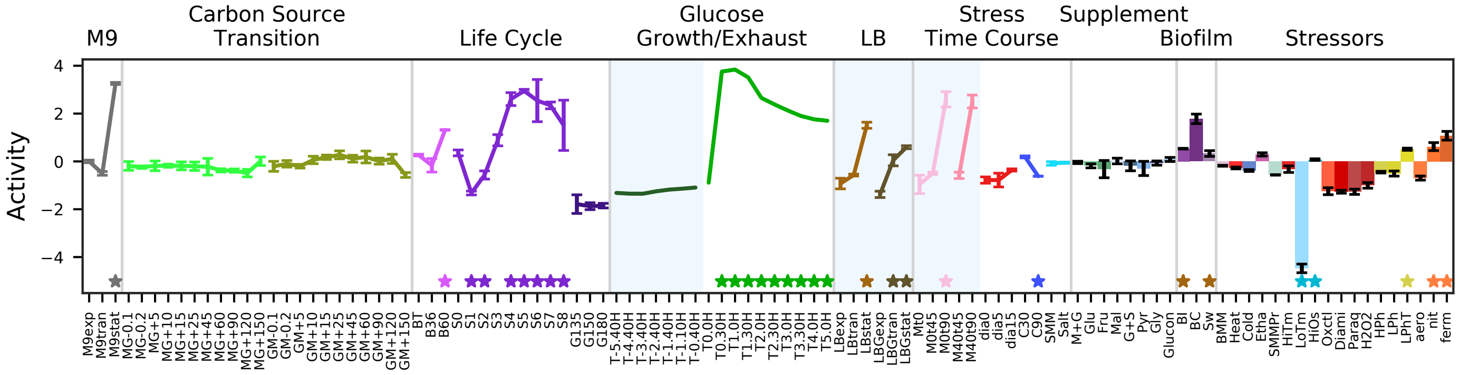


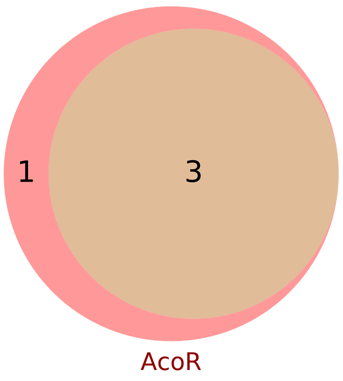


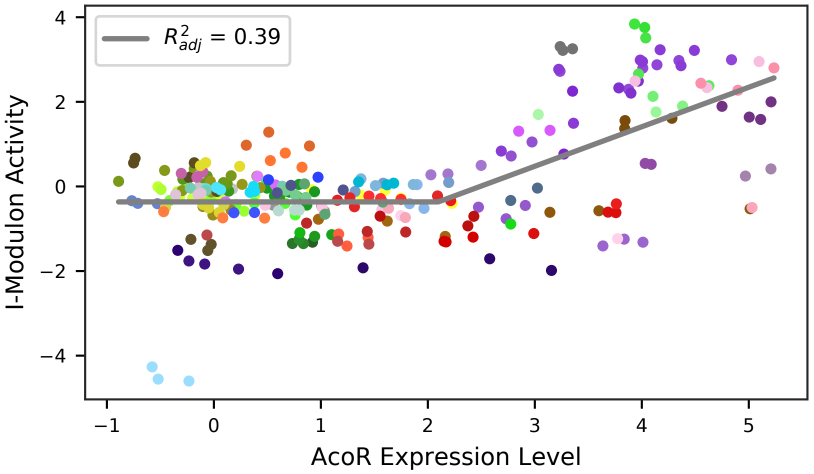

### 8 - LicR – Lichenan

Biological Function:
Uptake, phosphorylation, and utilization of lechenin, a glucan found in soil

Contains unknown genes and known regulon:
LicR

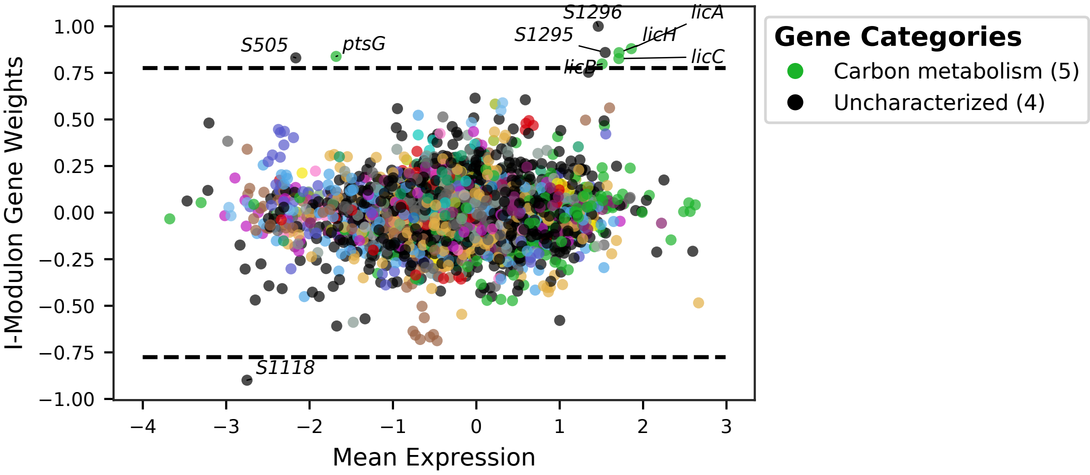


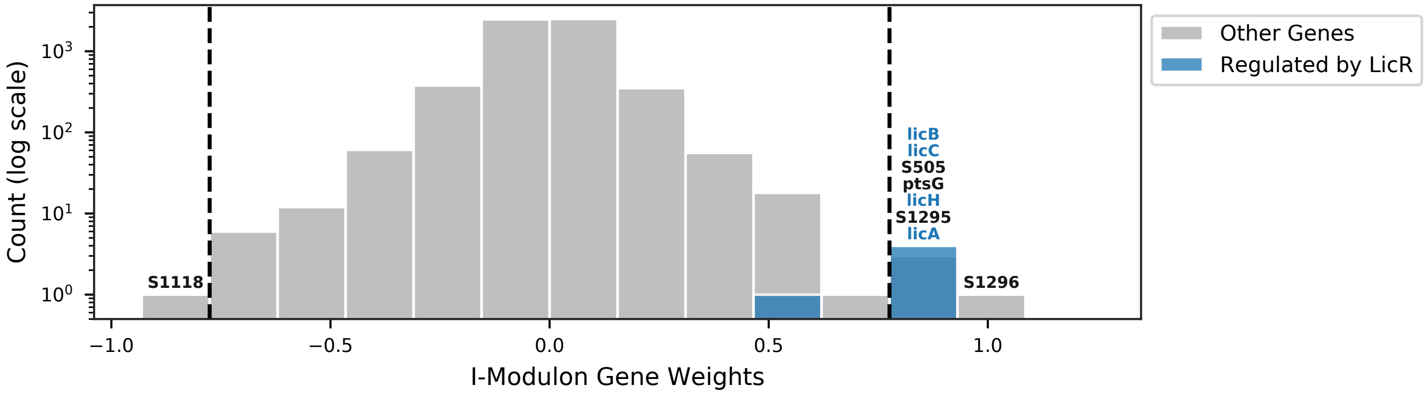


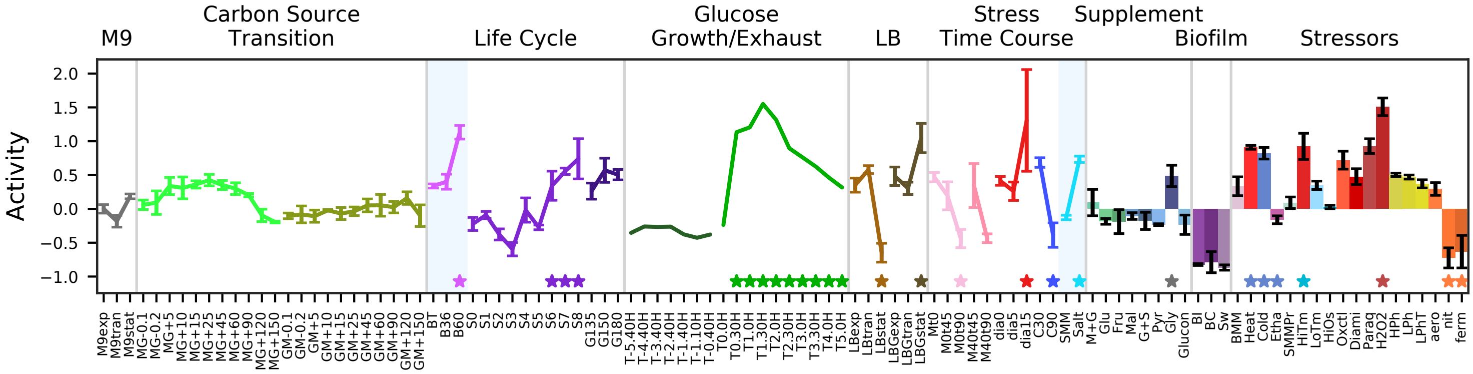


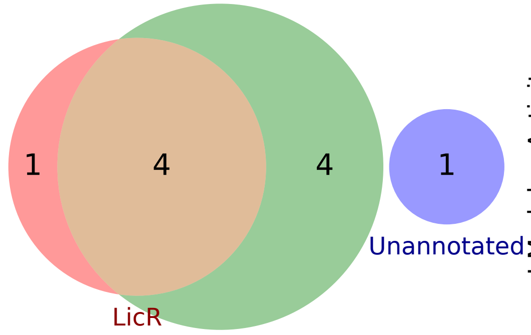


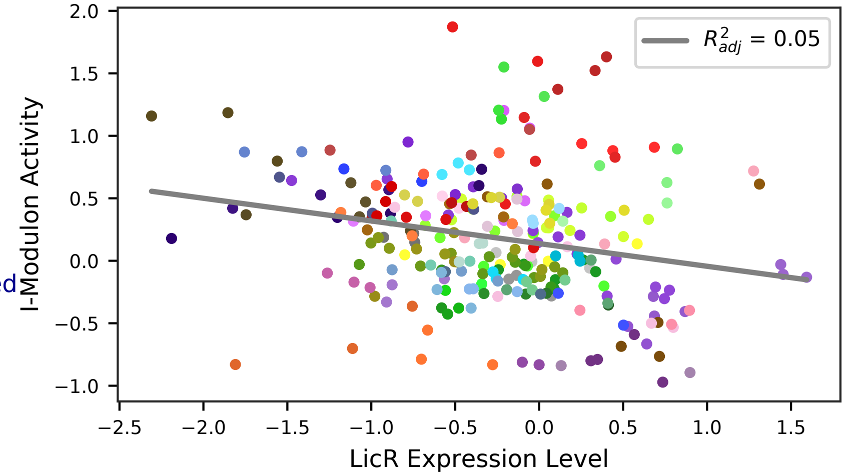

### 9 - CcpA-1 – Low Glucose 1

Biological Function:
Uptake and utilization of alternative carbon sources: ribose, sucrose, salicin, beta-glucosides, lichenan, glucomannon, trehalose, G3P, etc. High expression in exponential phase on LB media

Subset of known regulon:
CcpA + SigA

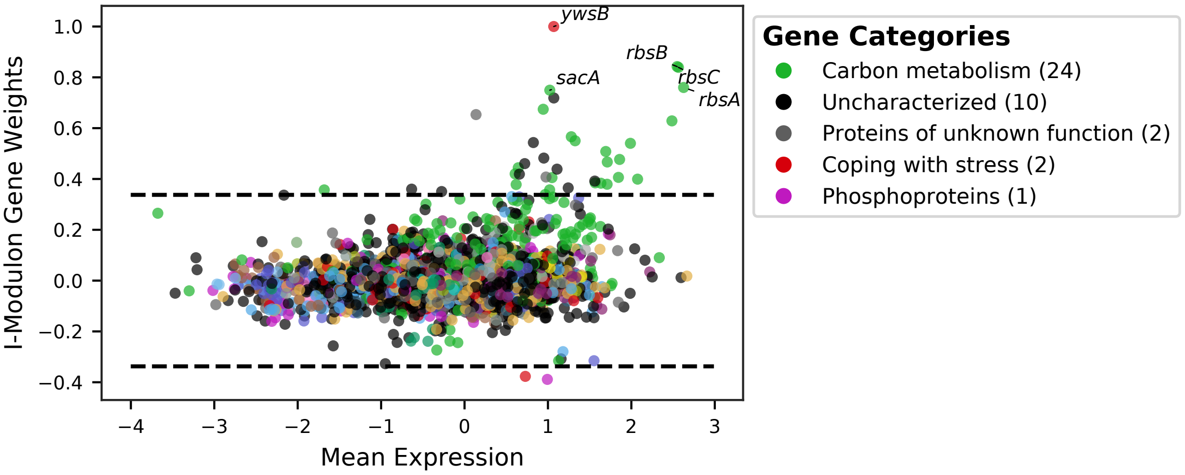


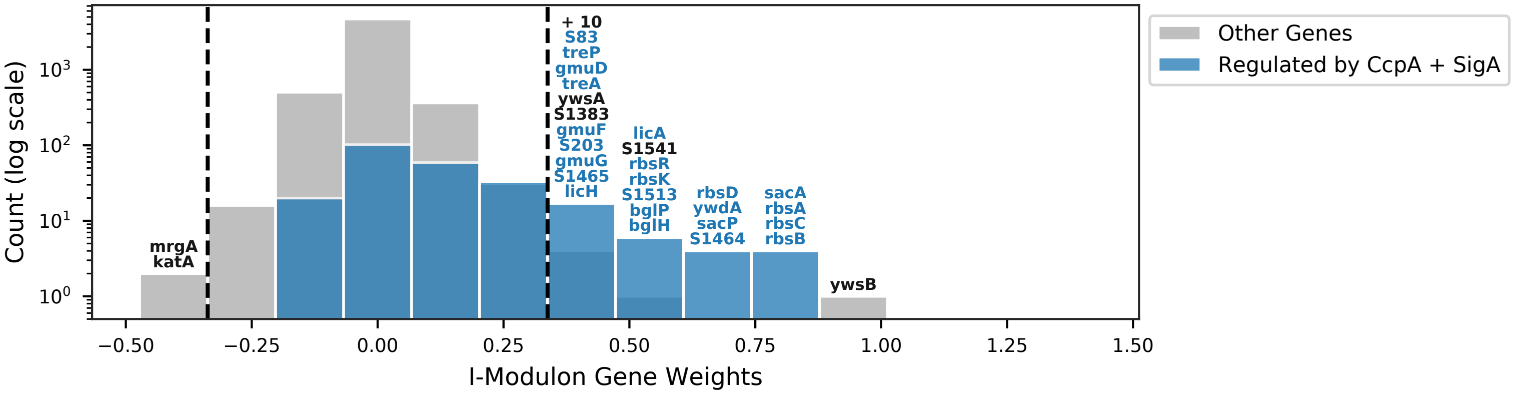


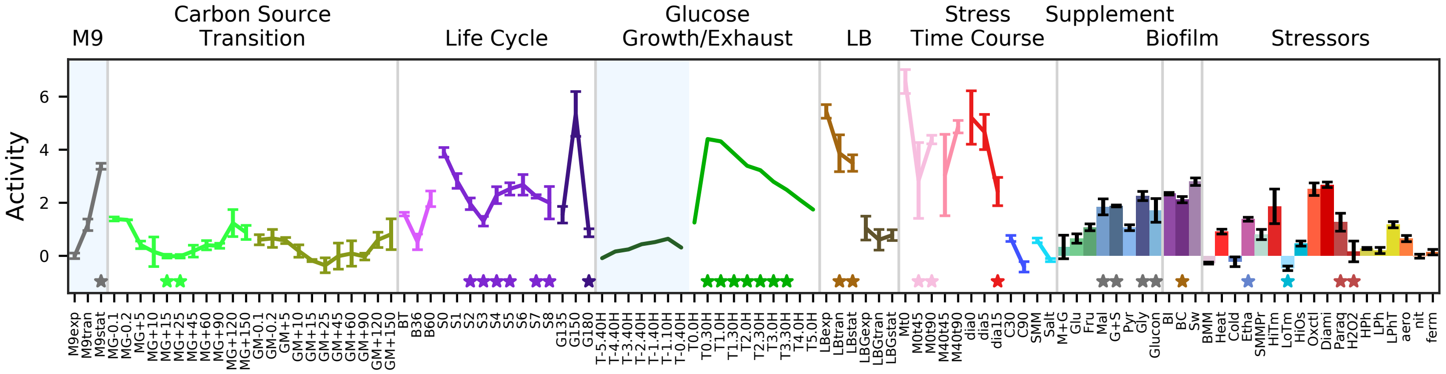


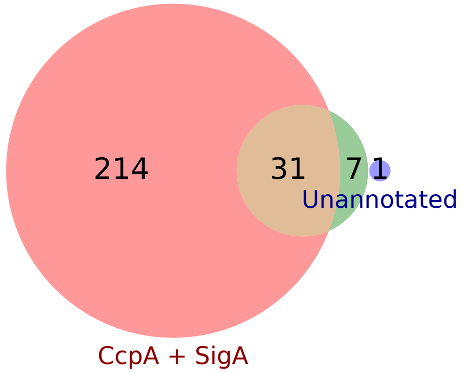


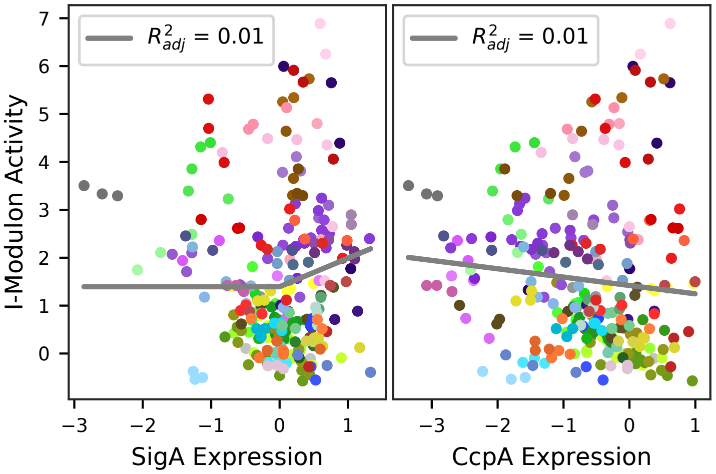

### 10 - CcpA-2 – Low Glucose 2

Biological Function:
Uptake, utilization, and regulation of alternative carbon sources: TCA intermediates, ribose, maltodextrin, galacto-oligosaccharides, acetoin, gluconate, etc. Suppression of glucose pathways. Stationary phase.

Enriched for known regulon:
[CcpA + SigA] / [CcpN]

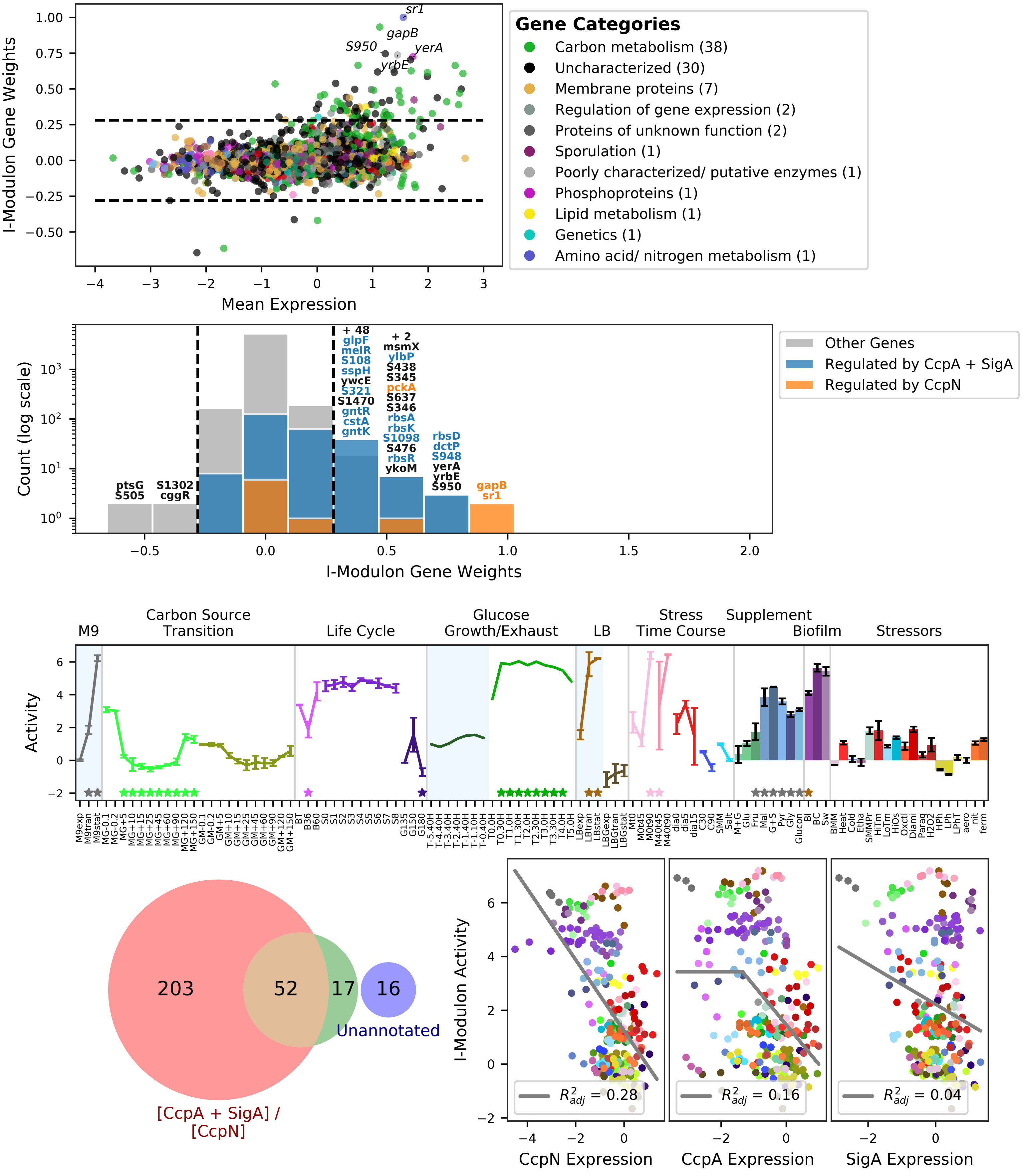

### 11 - His – Histidine Synthesis

Biological Function:
Synthesis of histidine

Subset (single operon) of known regulon:
YlxR

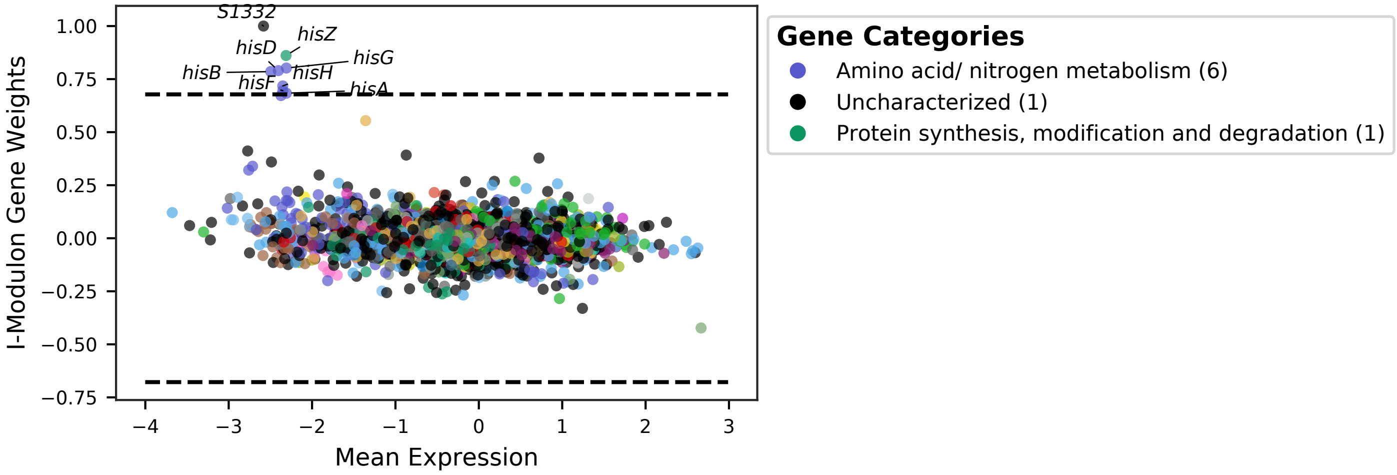


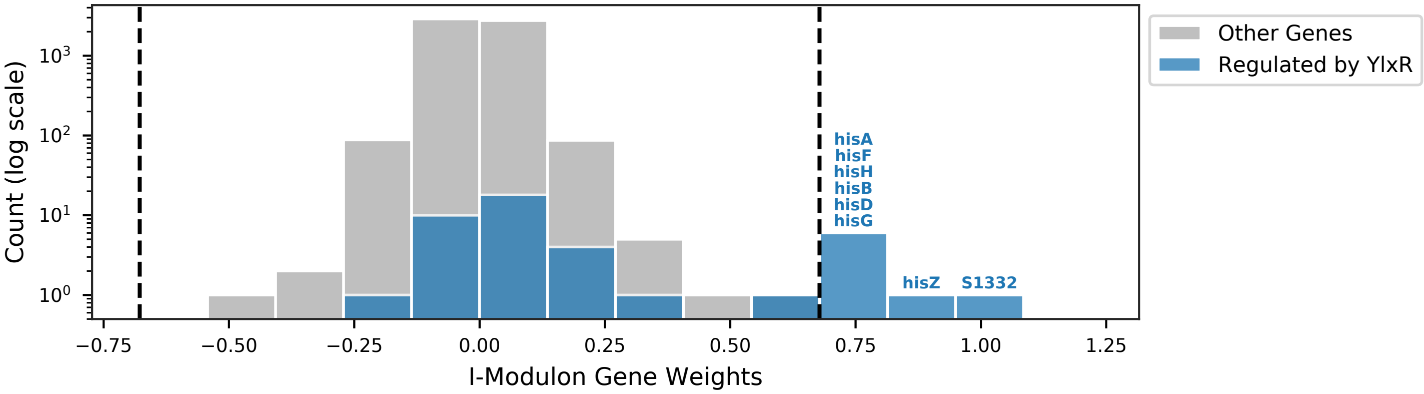


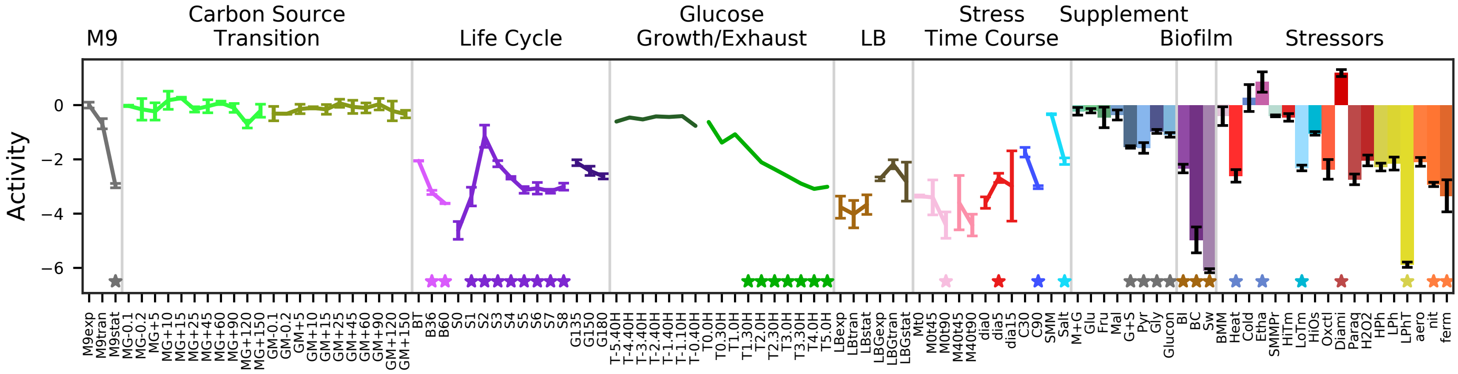


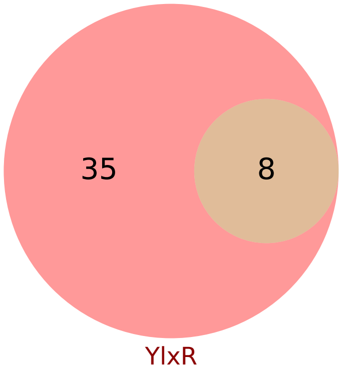


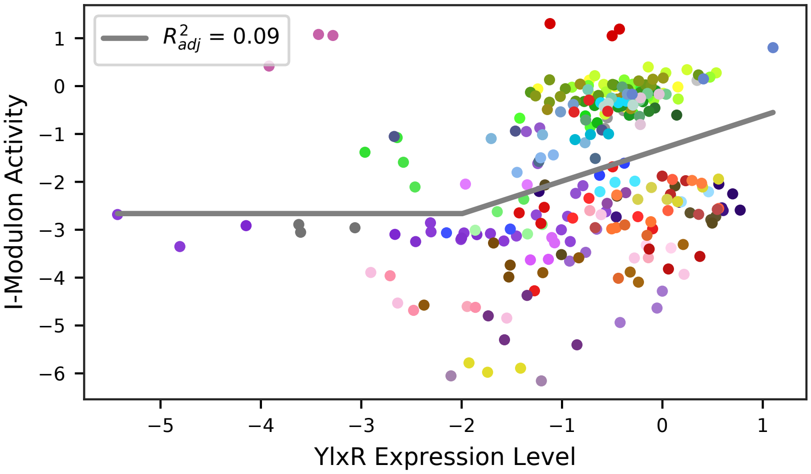

### 12 - AhrC – Arginine

Biological Function:
Uptake and biosynthesis of arginine

Well-defined regulon:
AhrC + YlxR

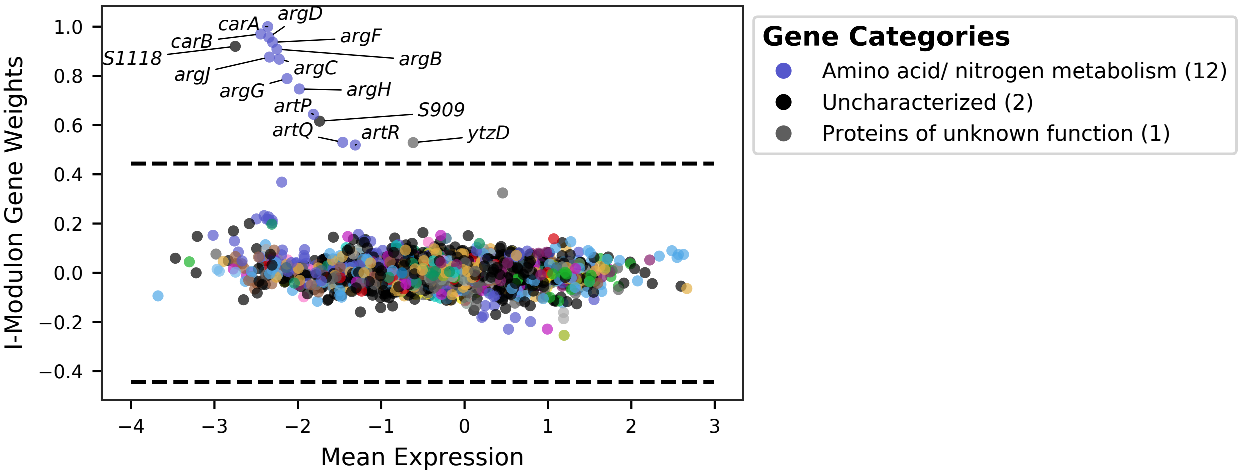


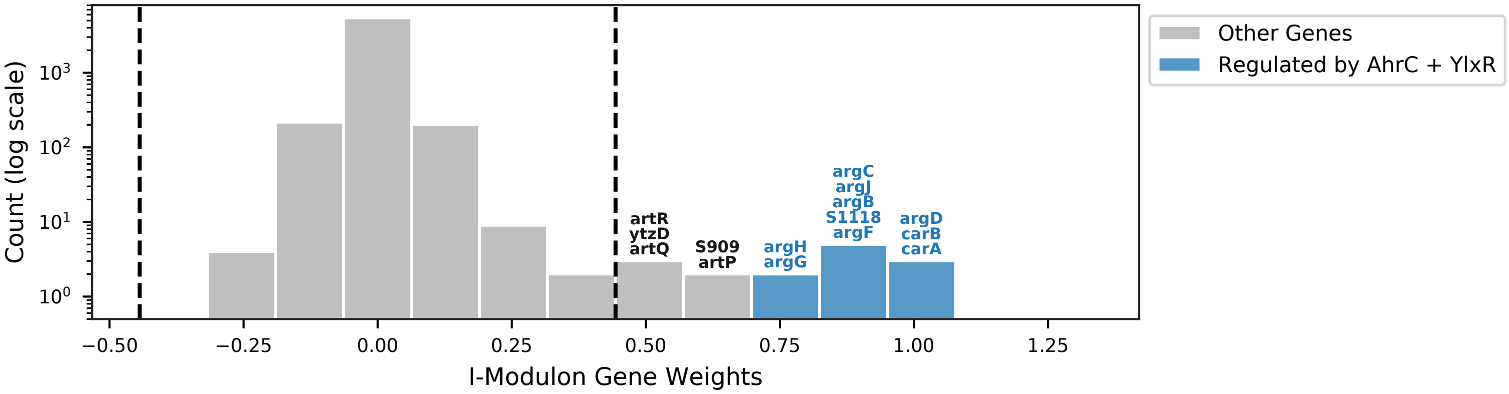


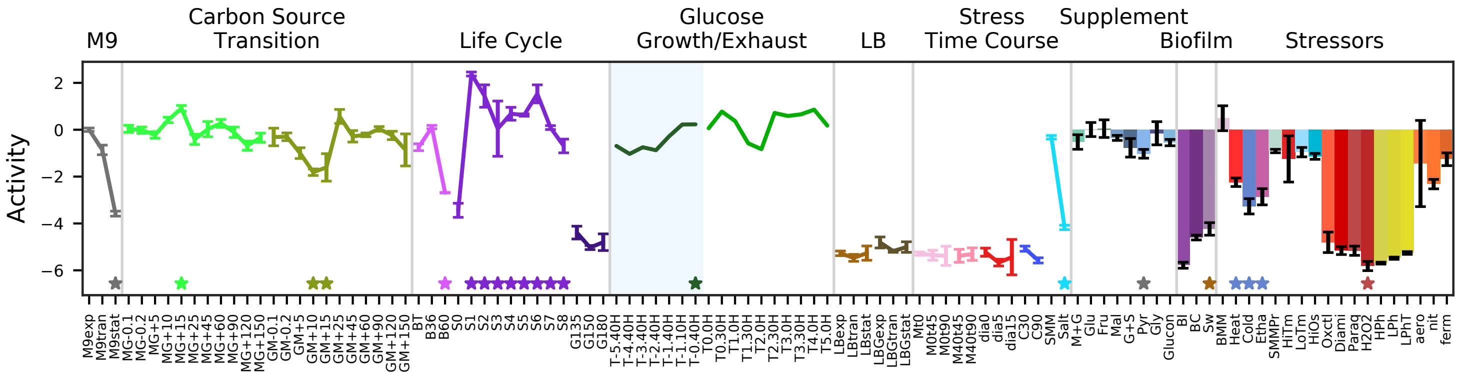


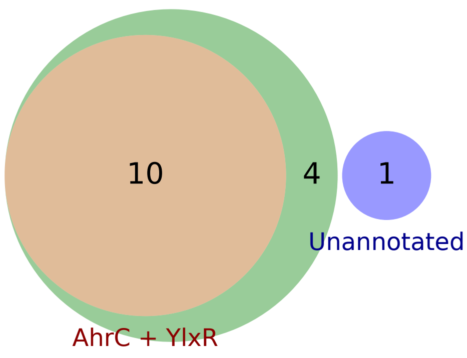


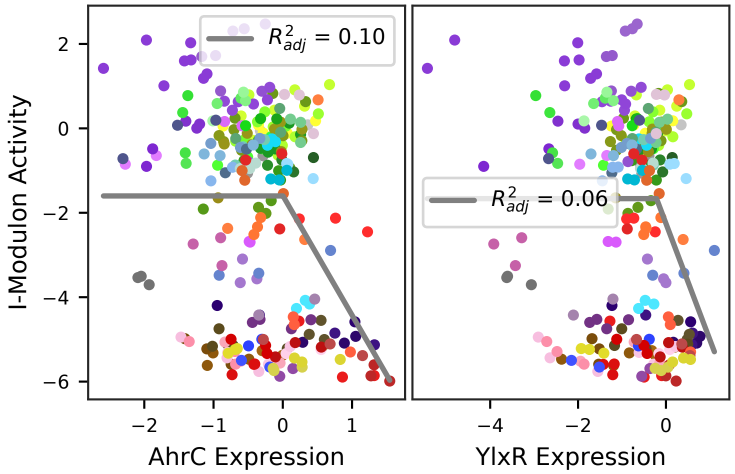

### 13 - MtrB – Tryptophan

Biological Function:
Biosynthesis of tryptophan

Subset (single operon) of known regulon:
MtrB

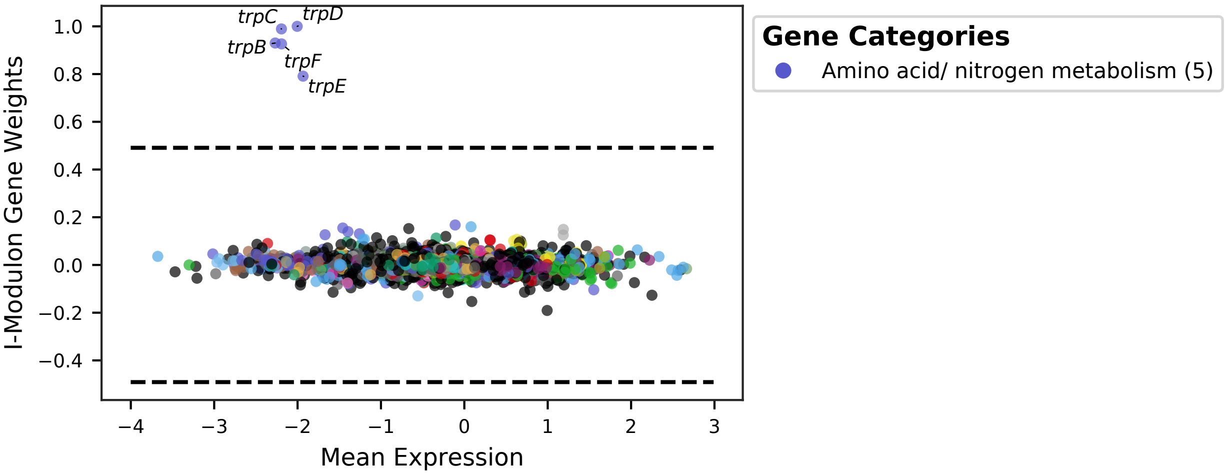


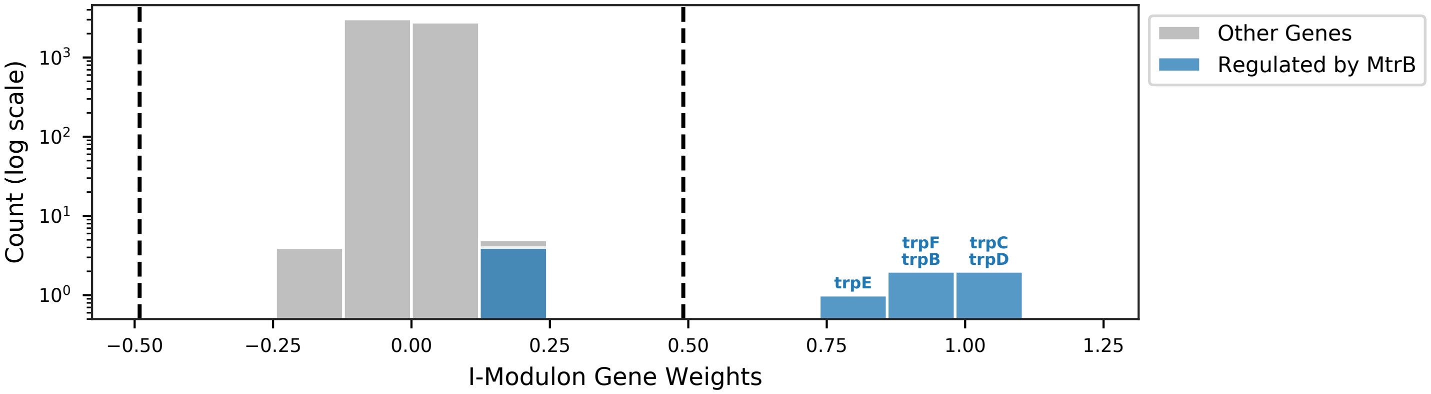


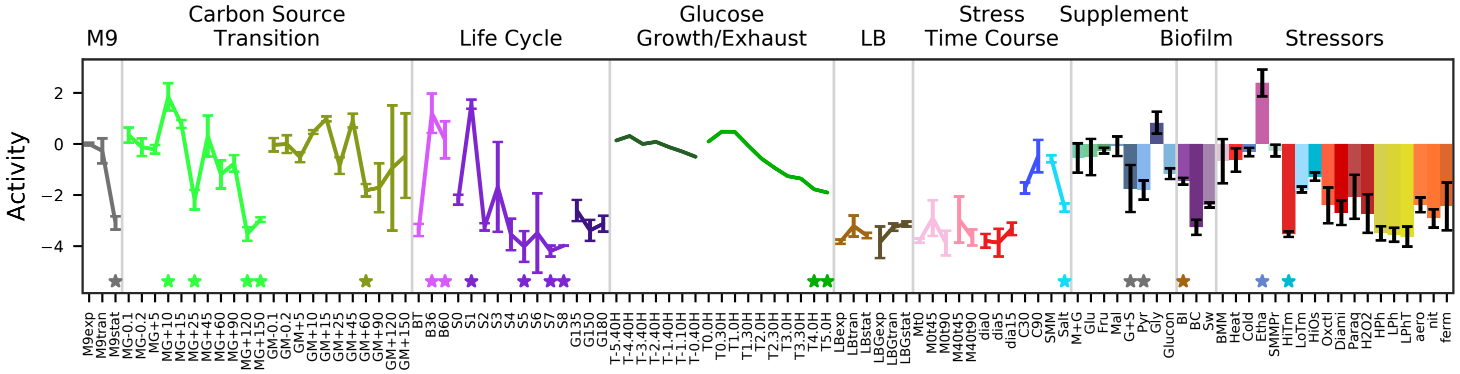


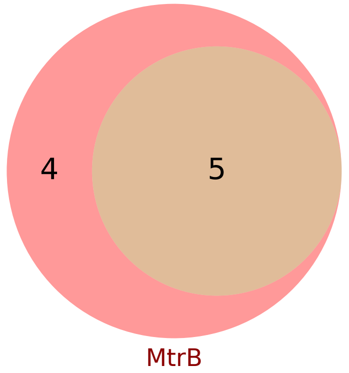


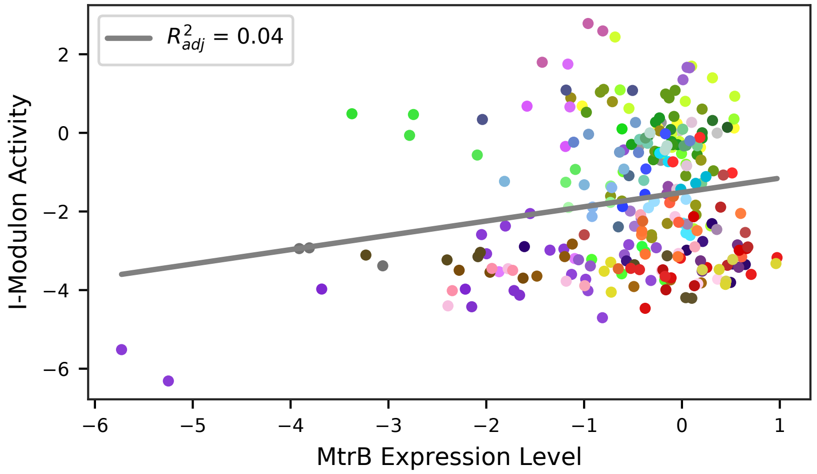

### 14 - S-Box – Methionine

Biological Function:
Methionine salvage and biosynthesis

Subset of known regulon:
S-box

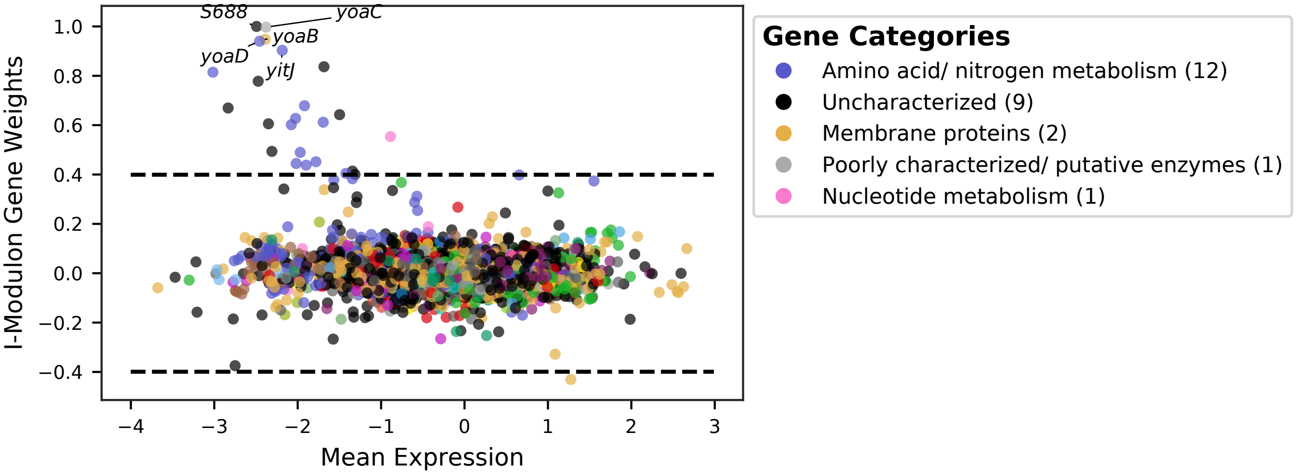


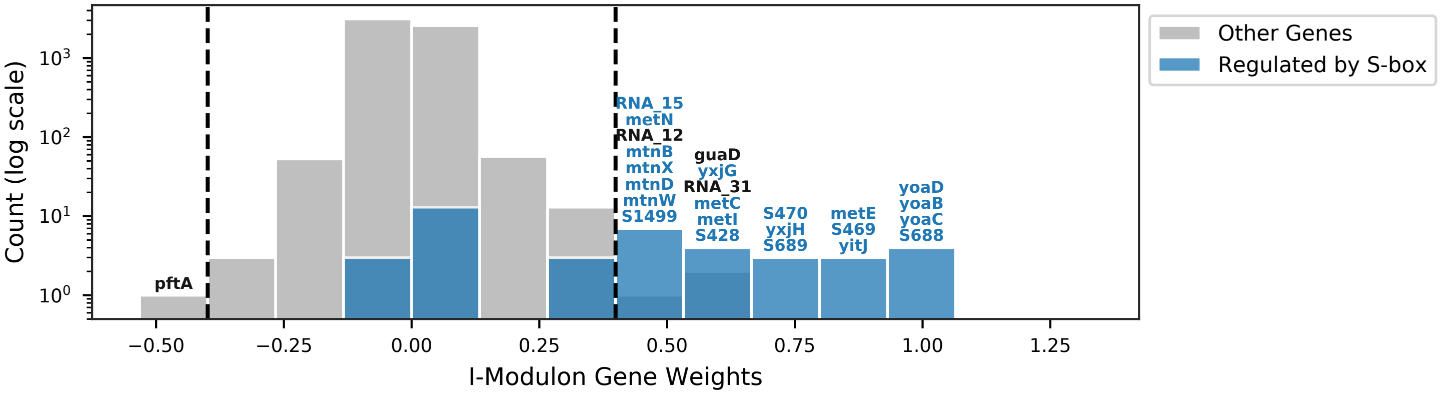


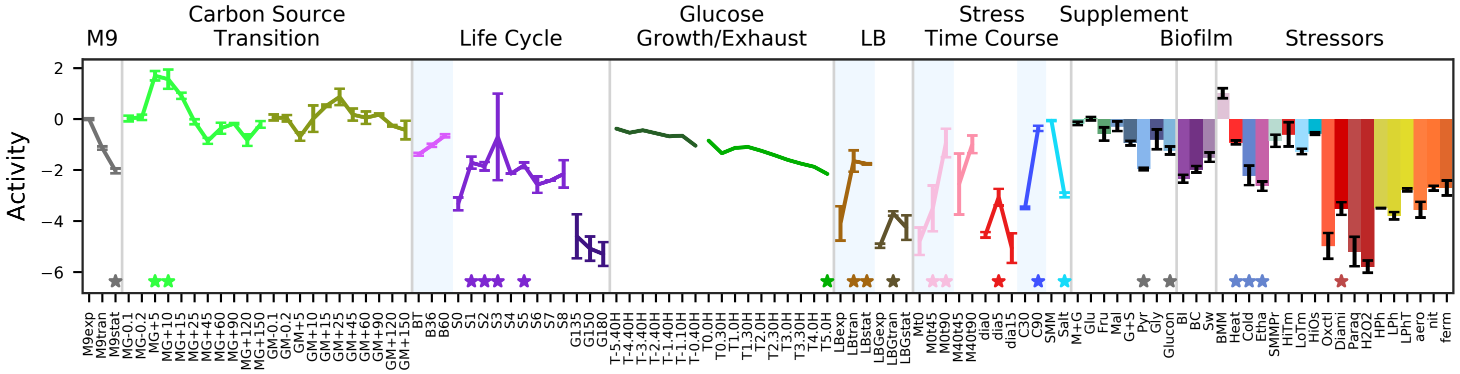


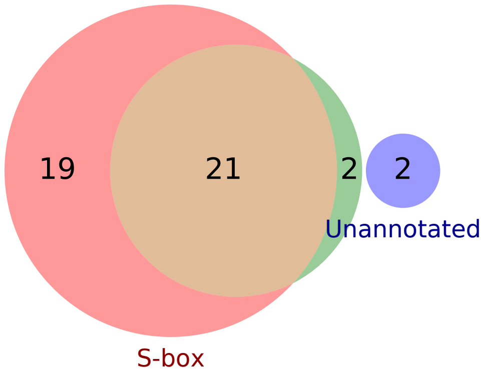

### 15 - TnrA + CodY – Leucine, Methionine, and Threonine

Biological Function:
Synthesis and salvage of leucine, methionine, threonine

Contains unknown genes and known regulon:
[TnrA + CodY + ThrR] / [TnrA + CodY + T-box + CcpA + SigA]

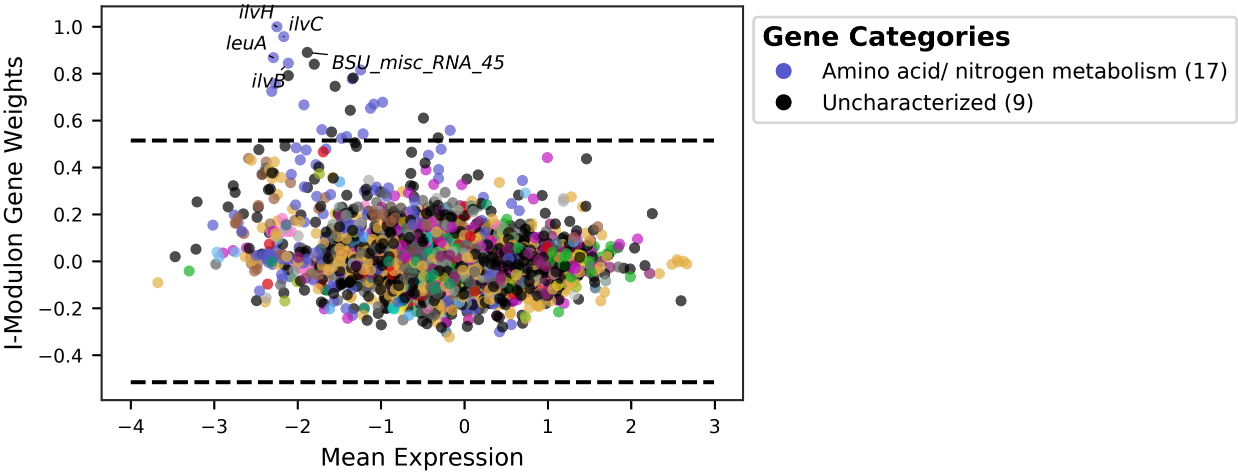


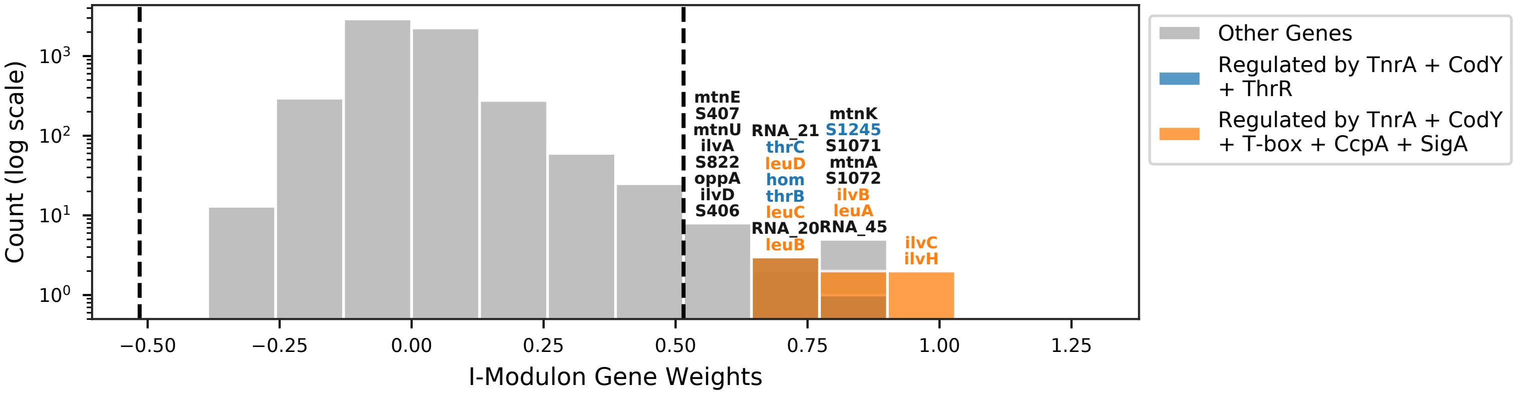


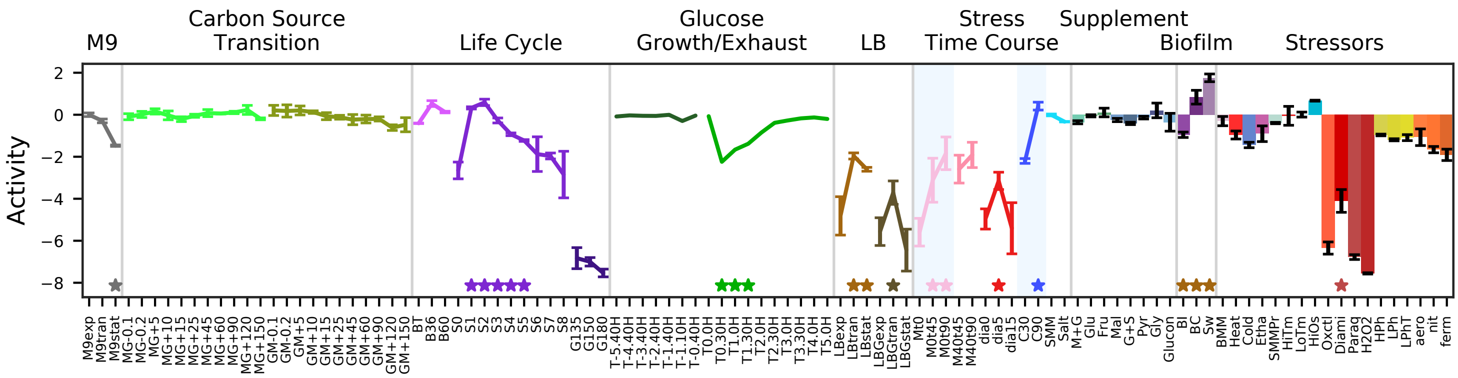


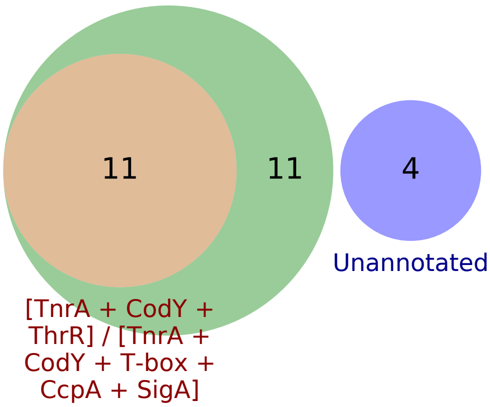


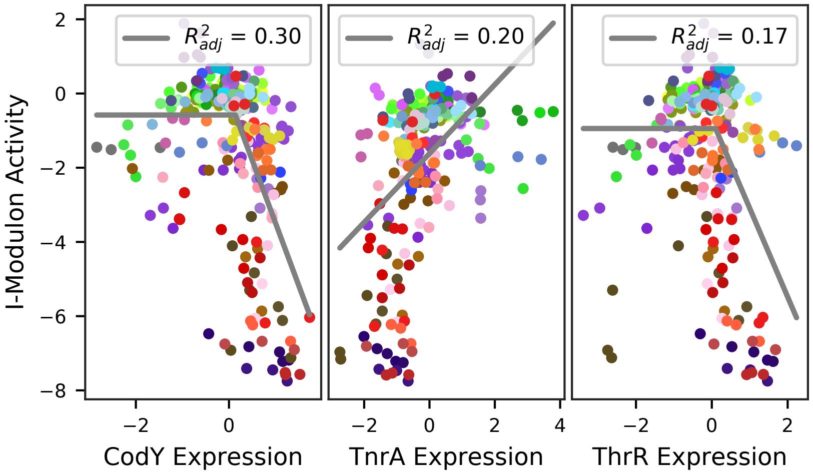

### 16 - CodY – BCAA Limitation

Biological Function:
Uptake of di-/oligopeptides, uptake of guanosine, cell wall metabolism

Subset of known regulon:
CodY

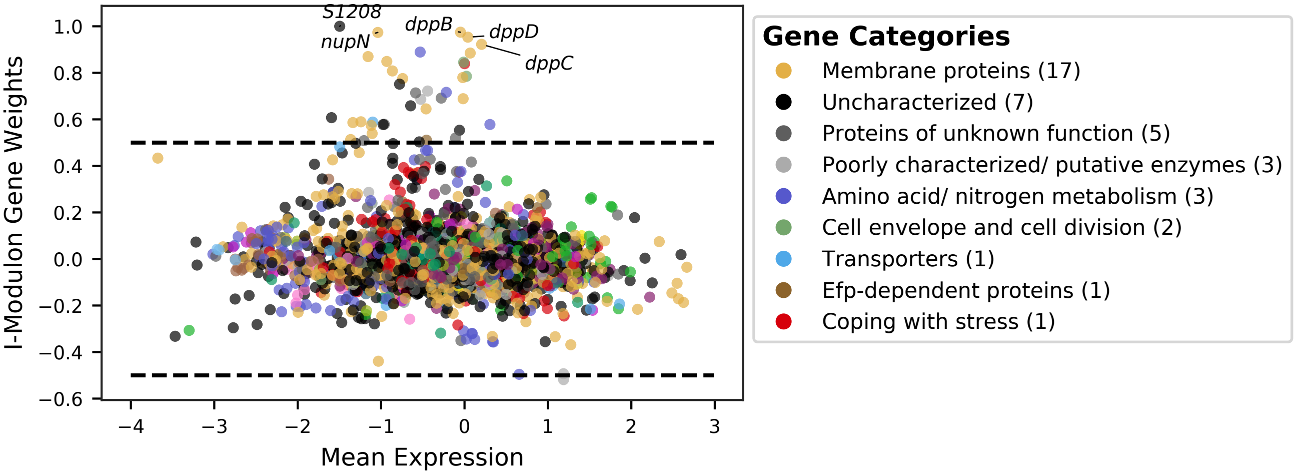


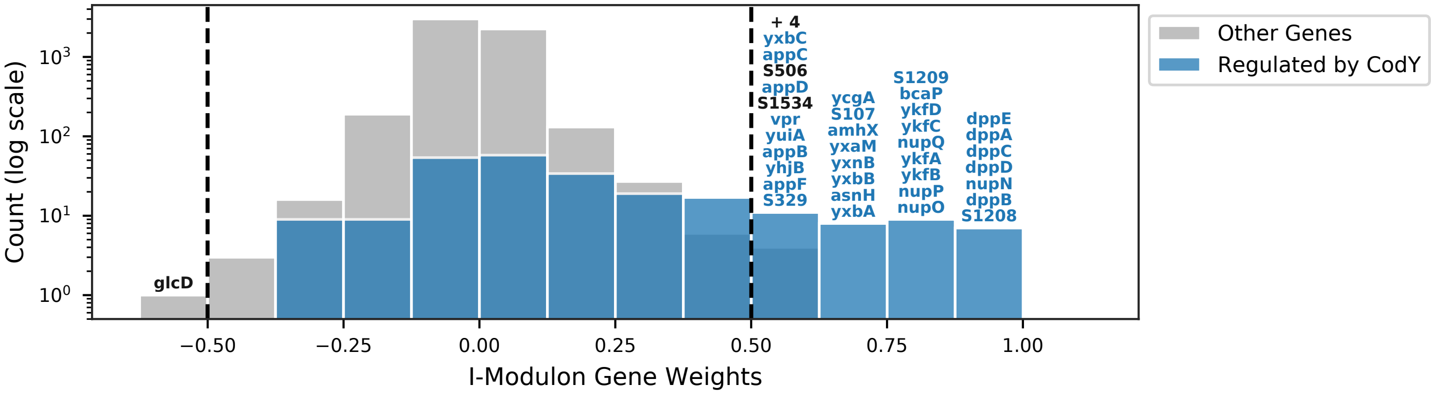


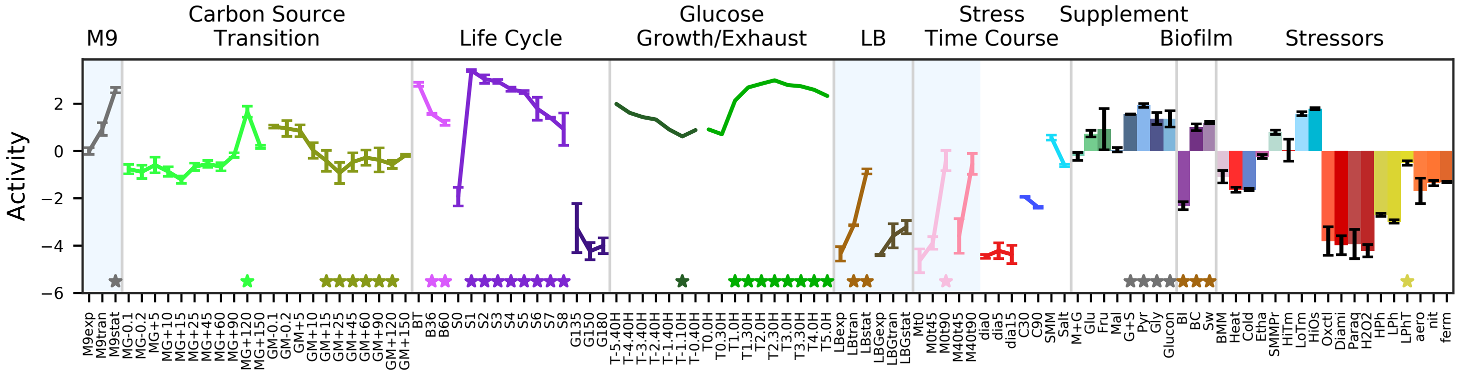


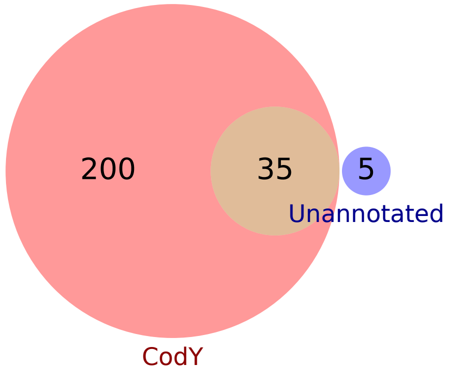


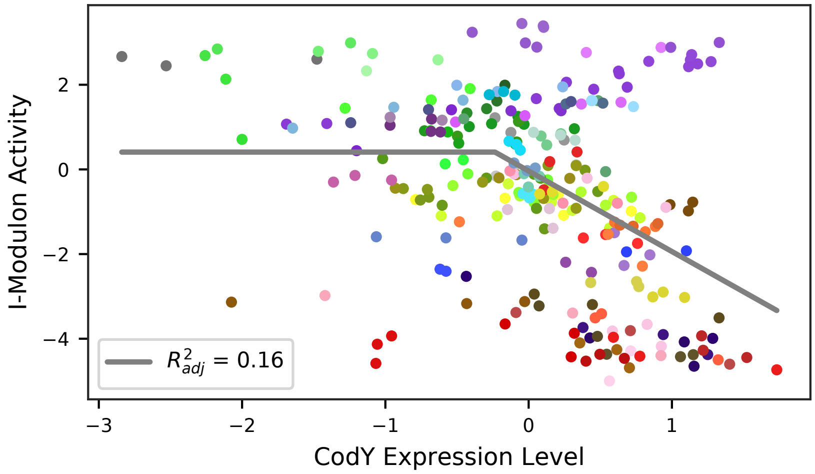

### 17 - HutP – Histidine Utilization

Biological Function:
Histidine uptake and utilization

Well-defined regulon:
HutP

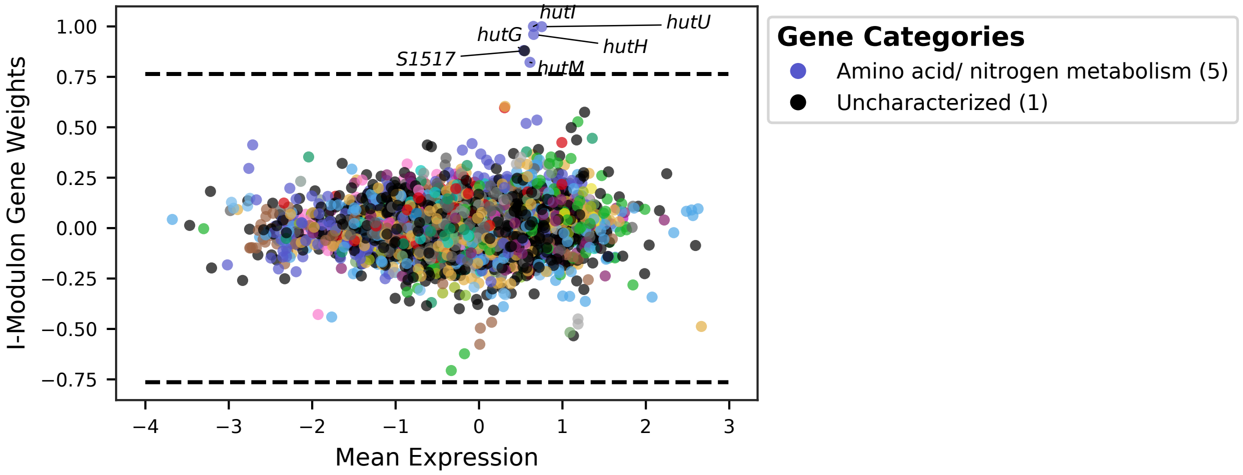


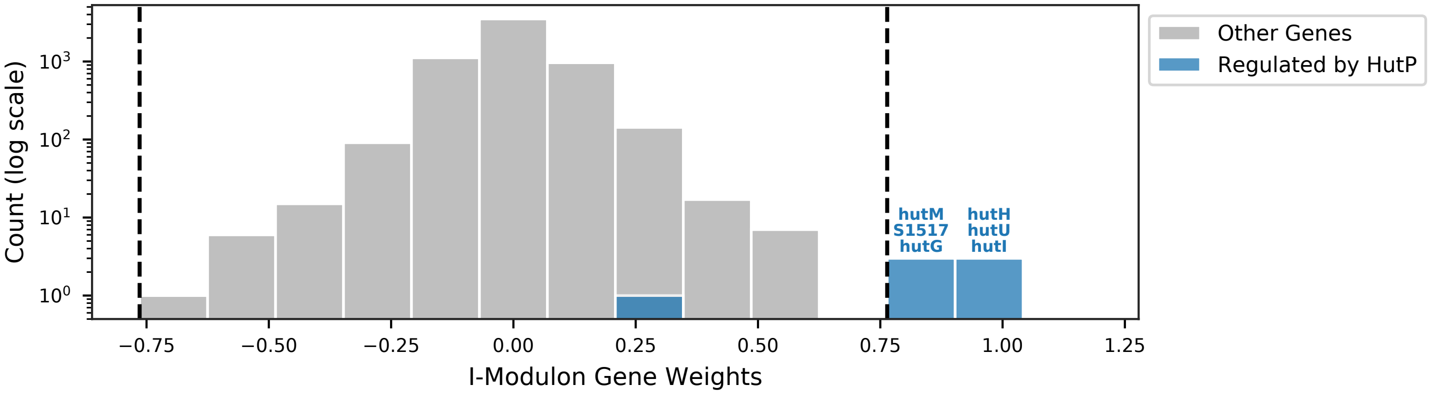


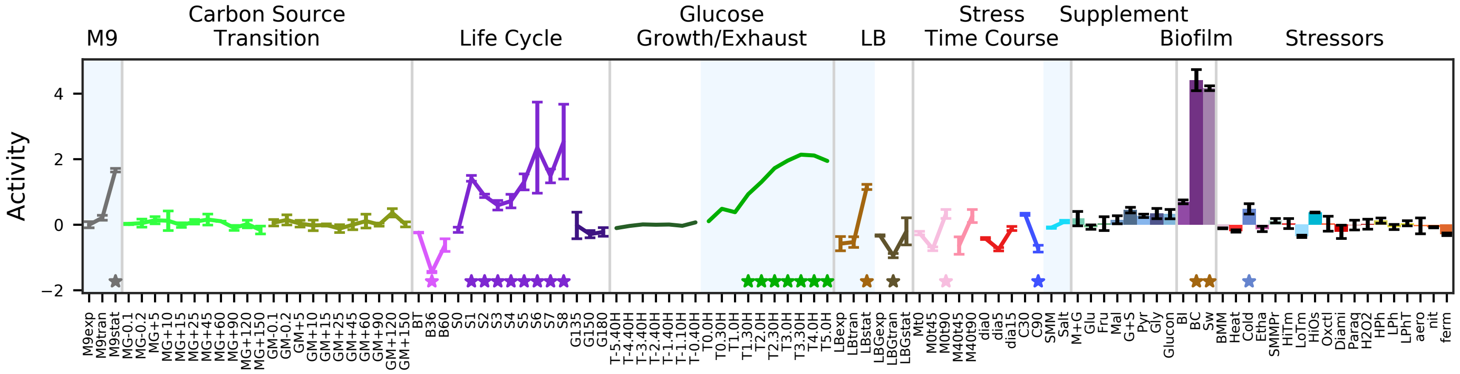


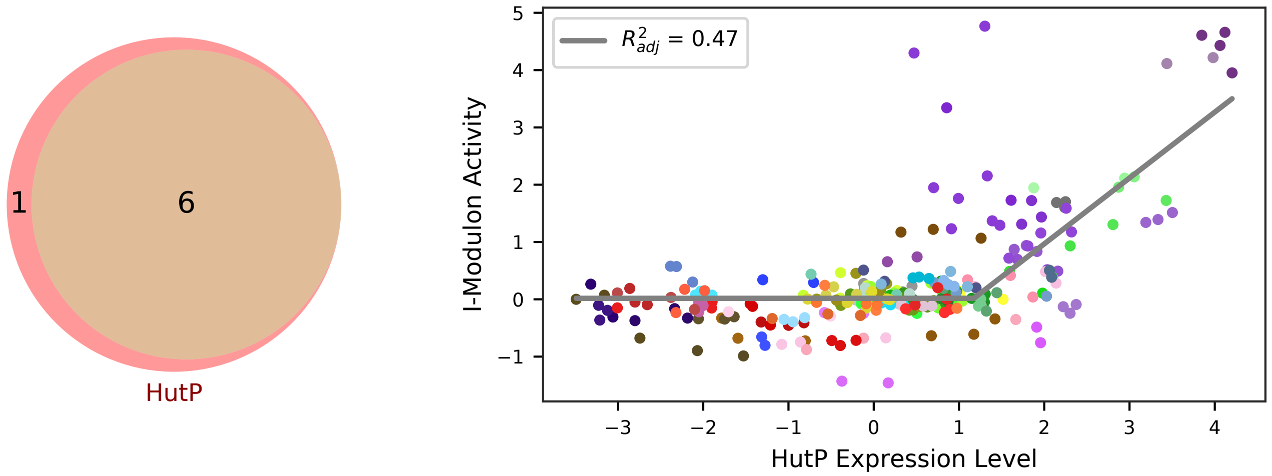

### 18 - RocR / PutR – Arginine and Proline Utilization

Biological Function:
Utilization of arginine, ornithine, citrulline and proline as carbon and energy sources

Well-defined regulon:
[SigL + AhrC + RocR] / [Spo0A + SigA + CodY + PutR]

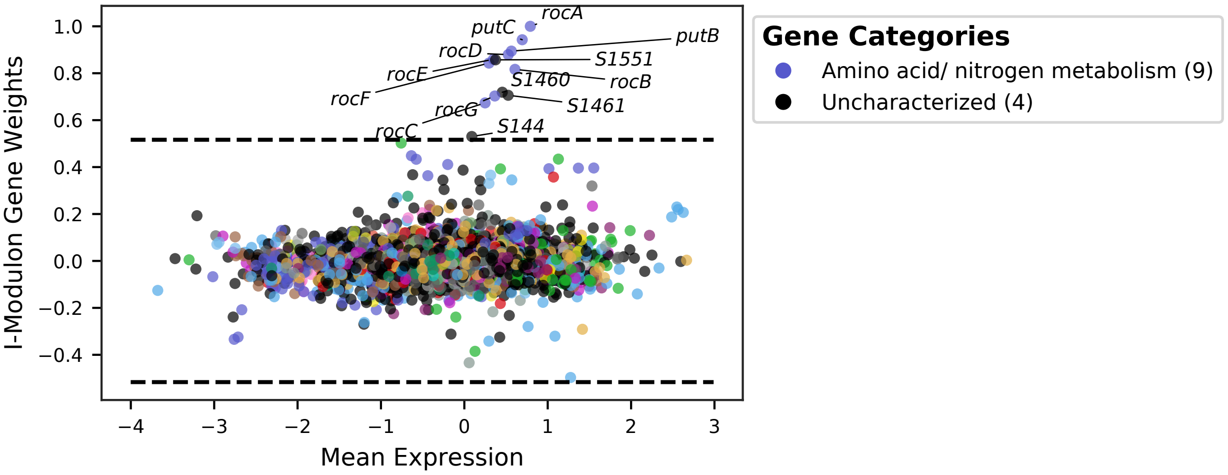


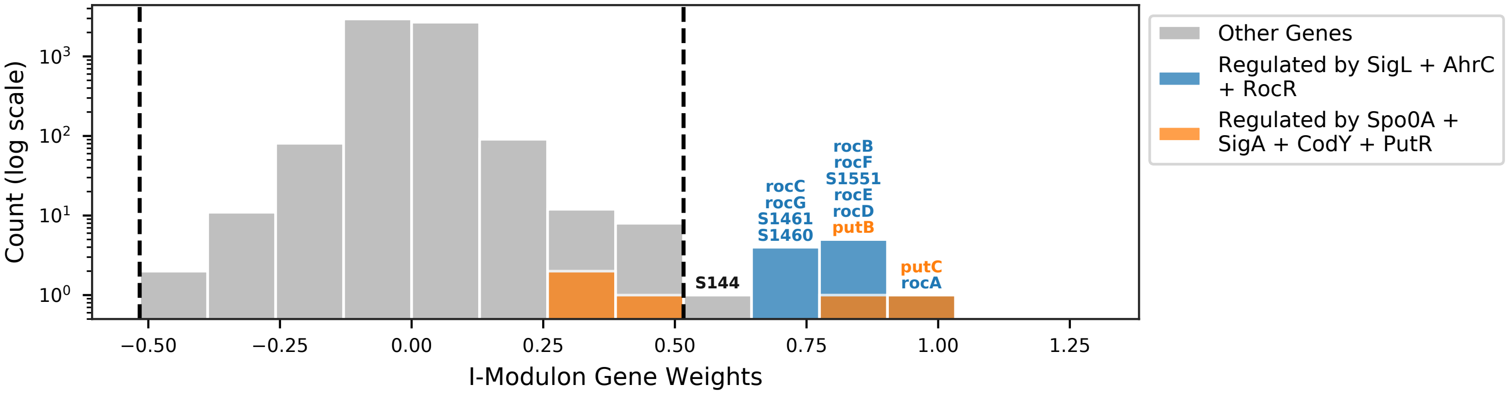


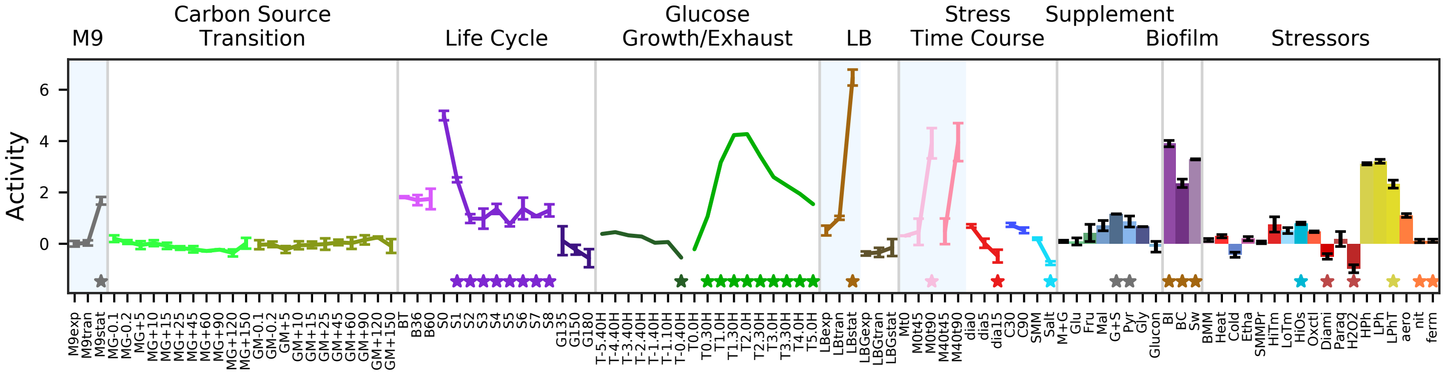


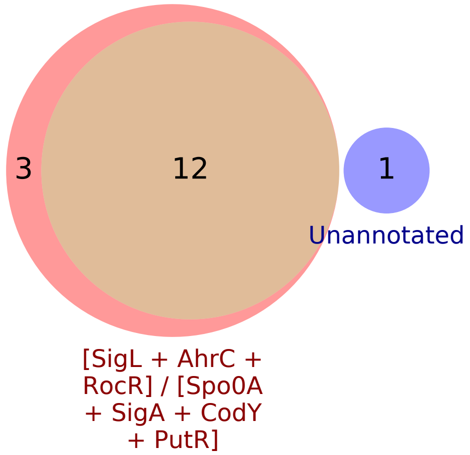


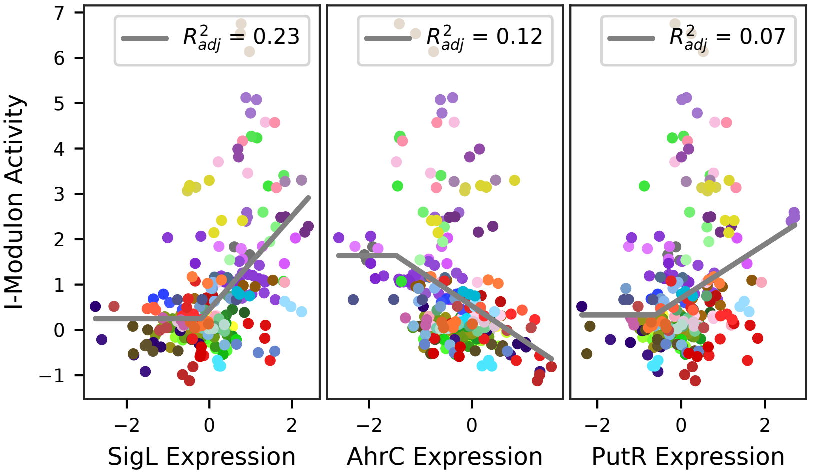

### 19 - PyrR – Pyrimidines

Biological Function:
Synthesis of pyrimidines

Well-defined regulon:
PyrR

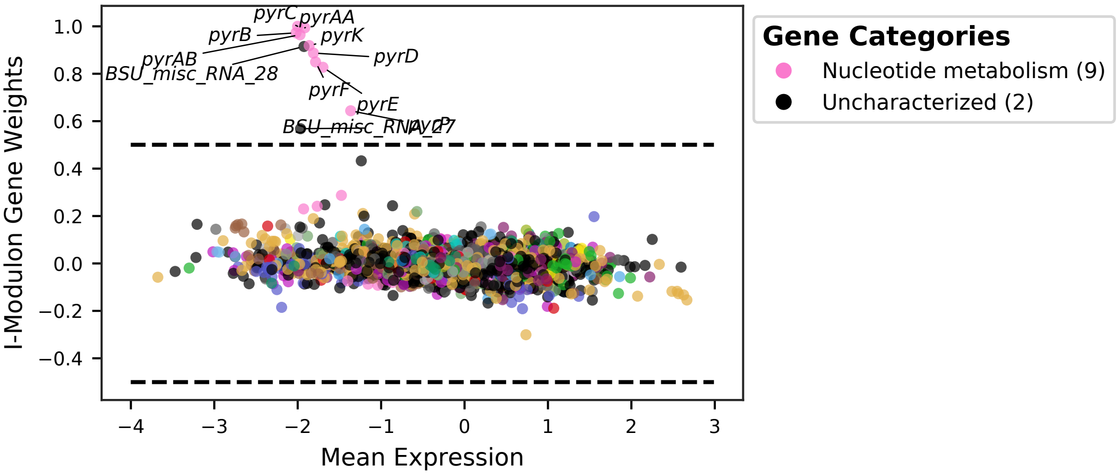


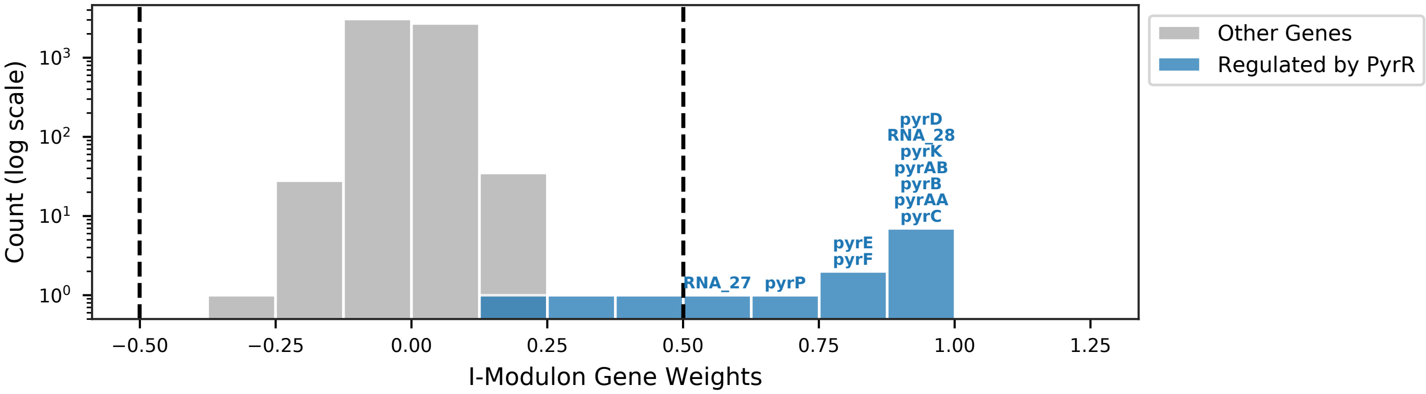


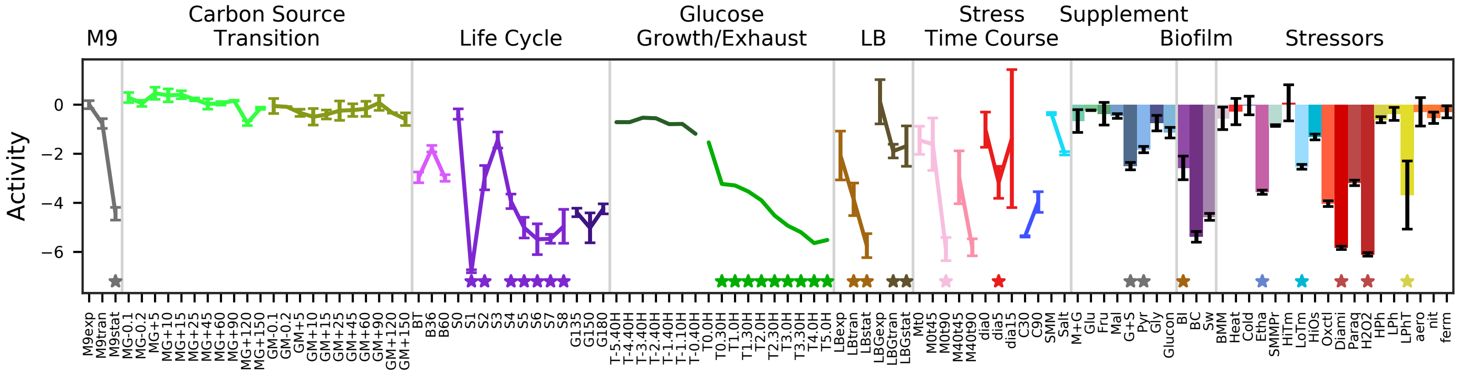


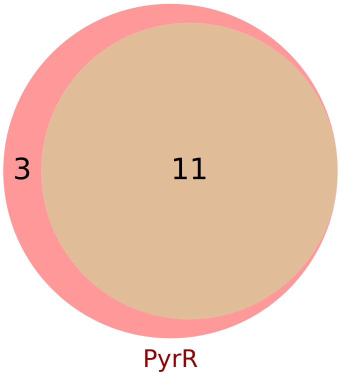


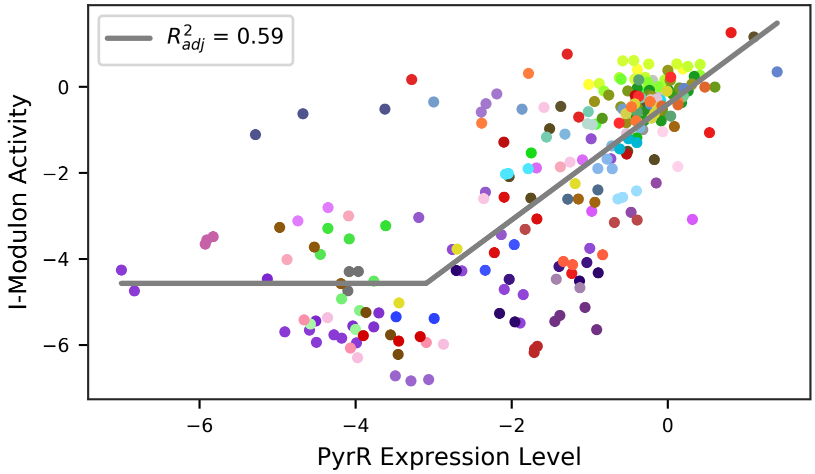

### 20 - PurR – Purines

Biological Function:
Synthesis of purine, xanthine and guanine uptake and conversion into purines

Well-defined regulon:
PurR

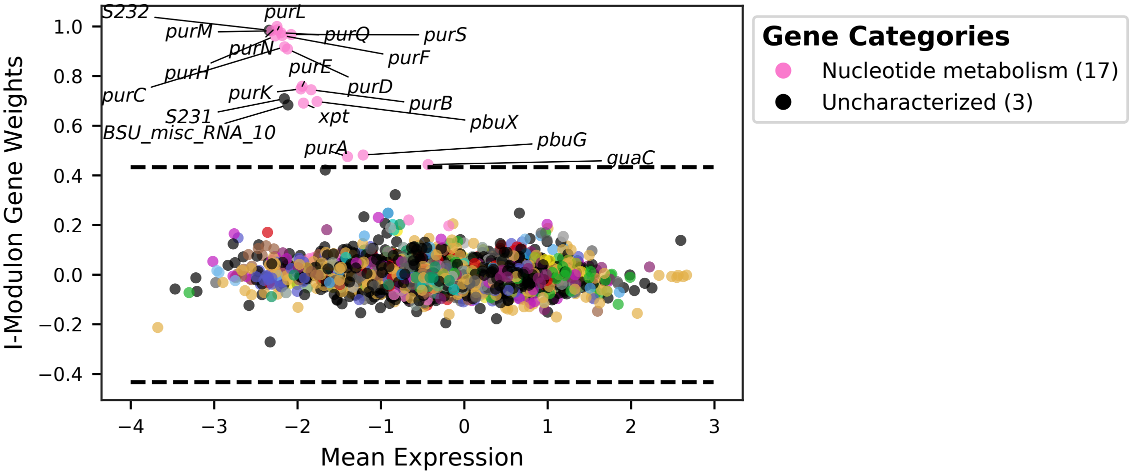


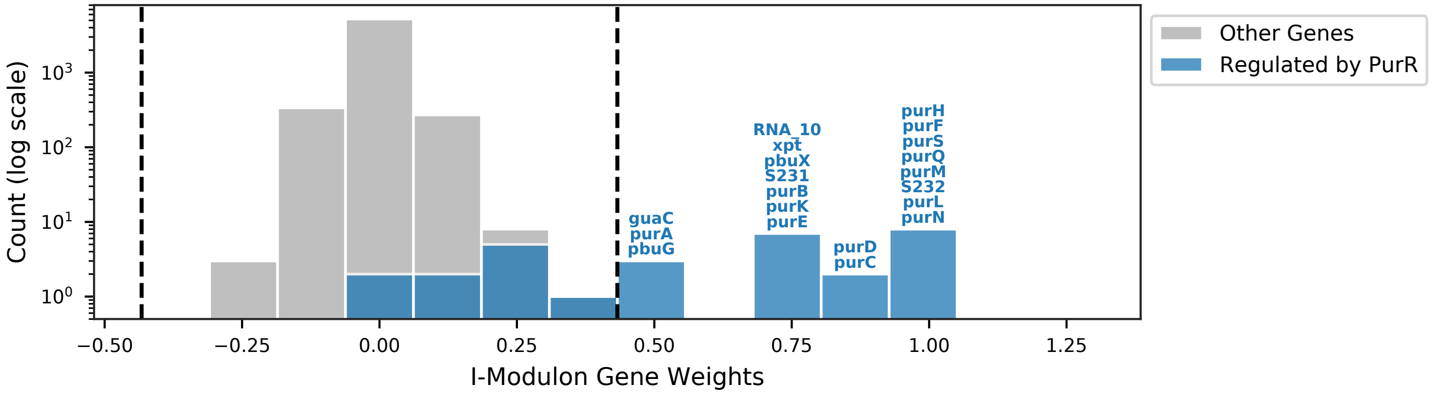


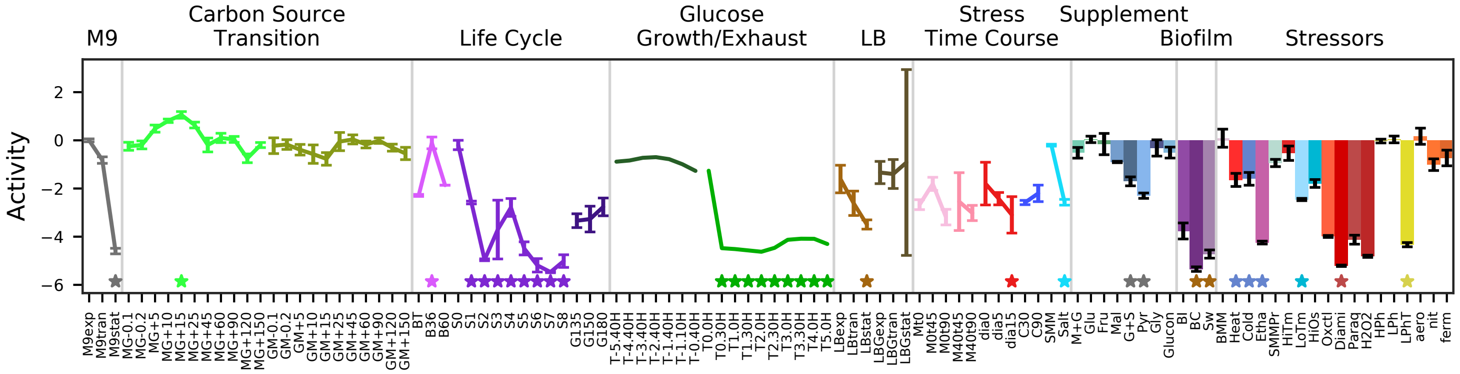


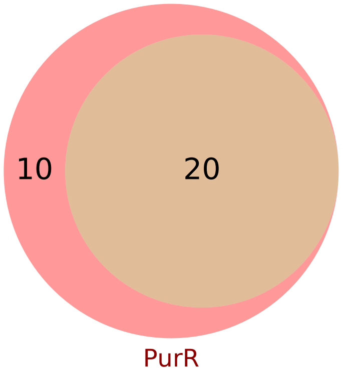


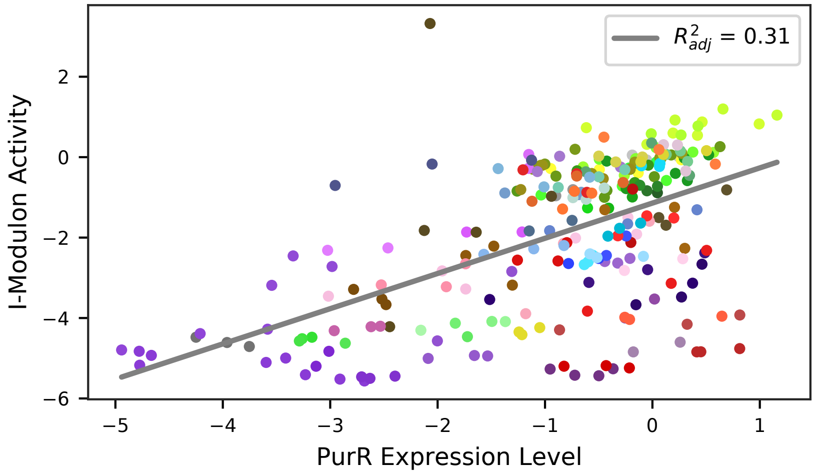

### 21 - Thi-Box – Thiamine

Biological Function:
Synthesis and uptake of thiamine

Well-defined regulon:
Thi-box

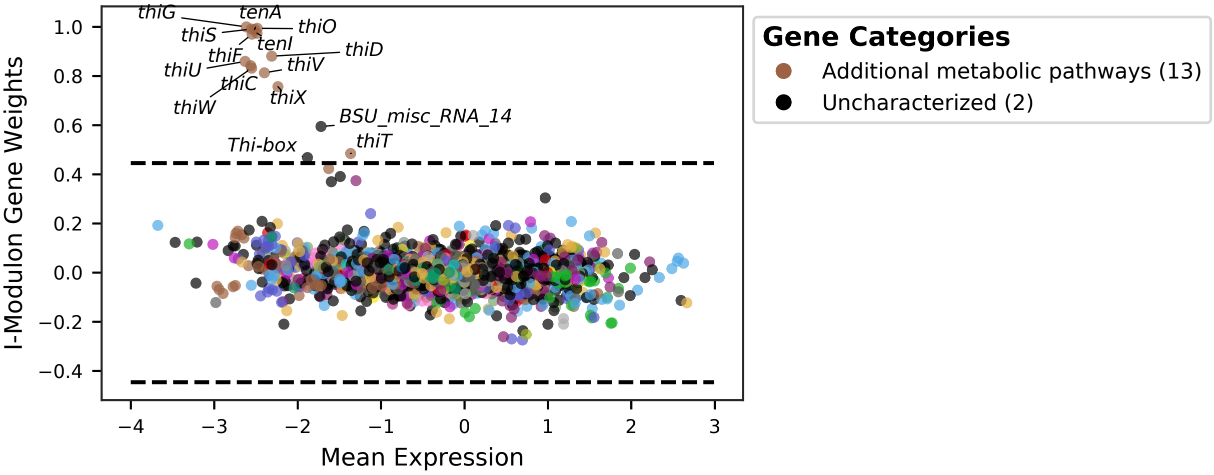


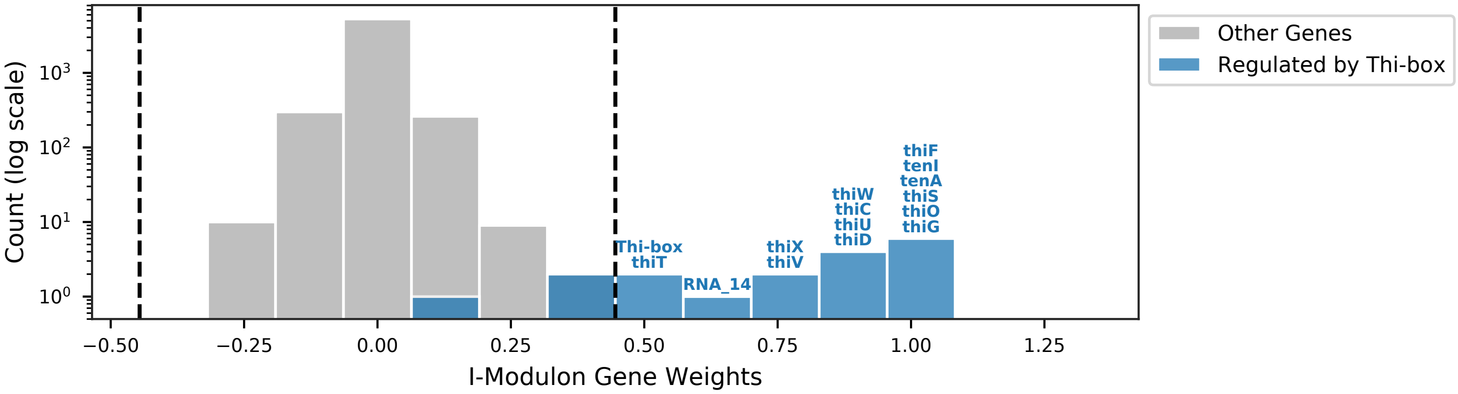


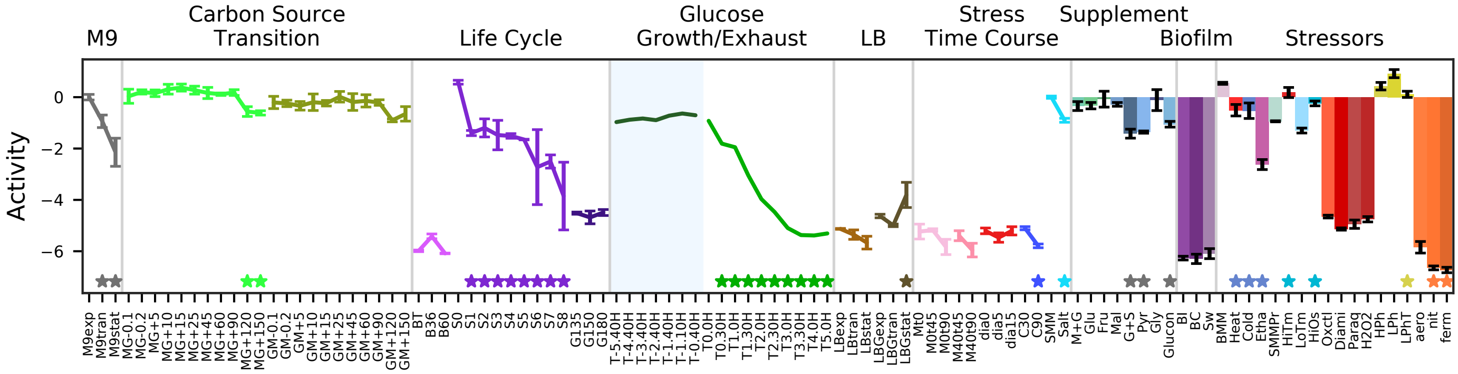


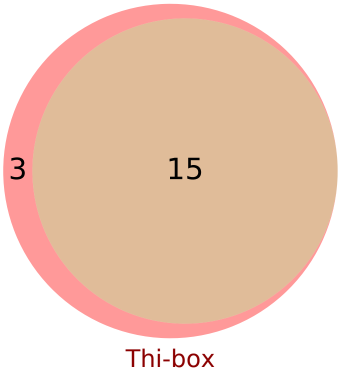


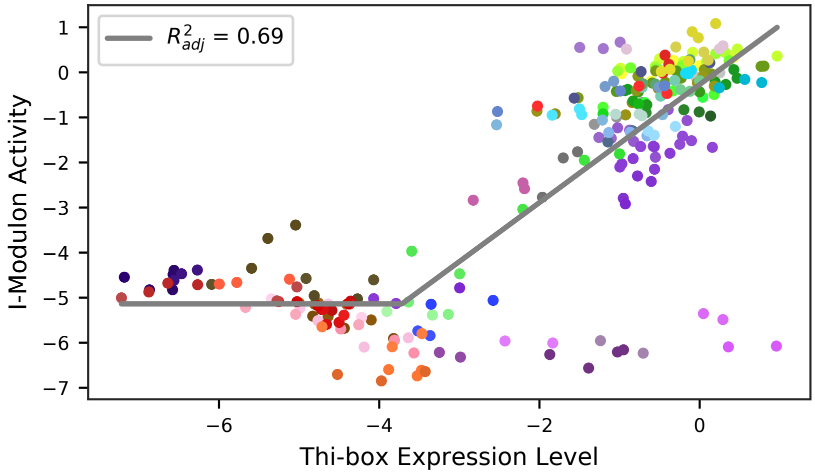

### 22 - PchR – Pulcherrimin

Biological Function:
Extracellular synthesis and import of pulcherrimin for iron acquisition

Subset of known regulon:
PchR

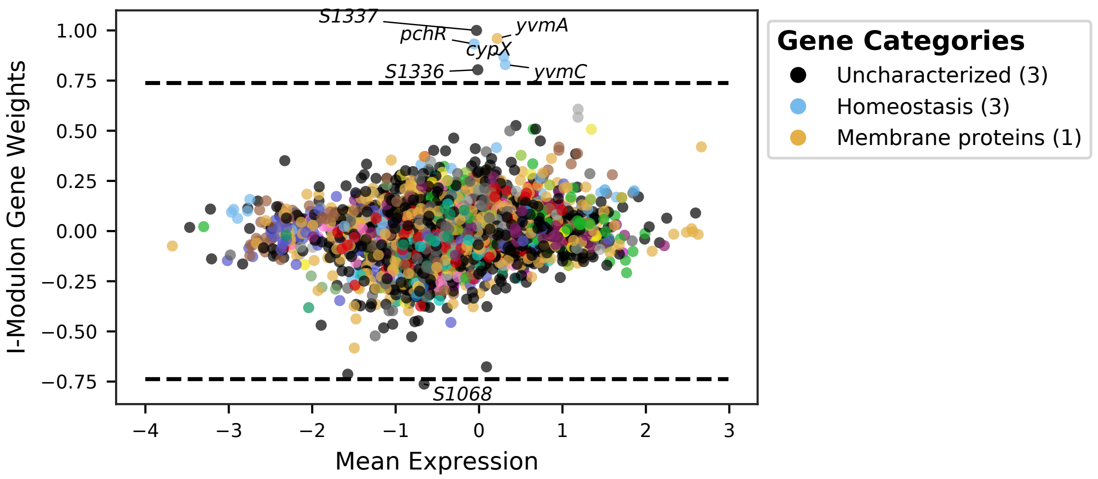


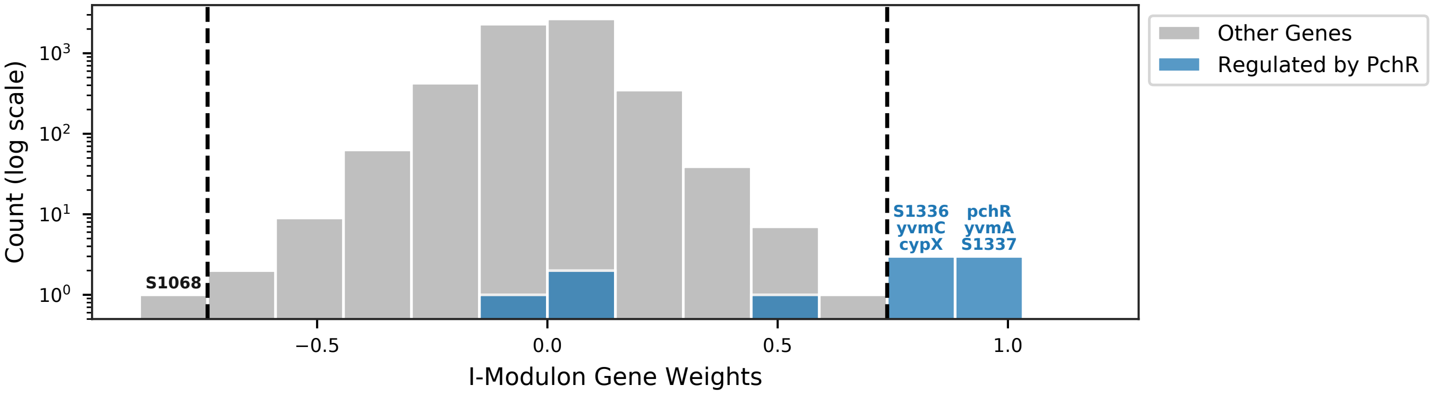


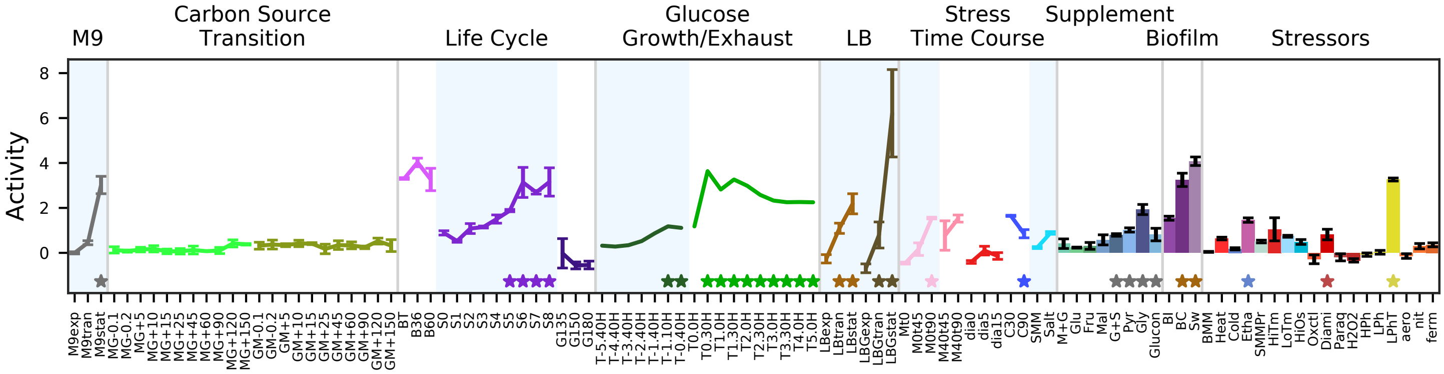


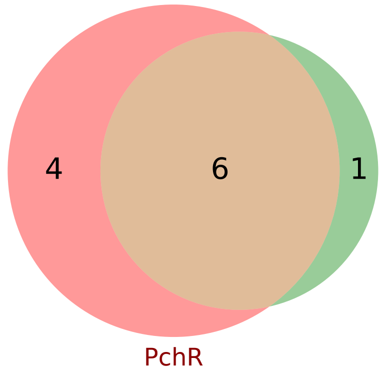


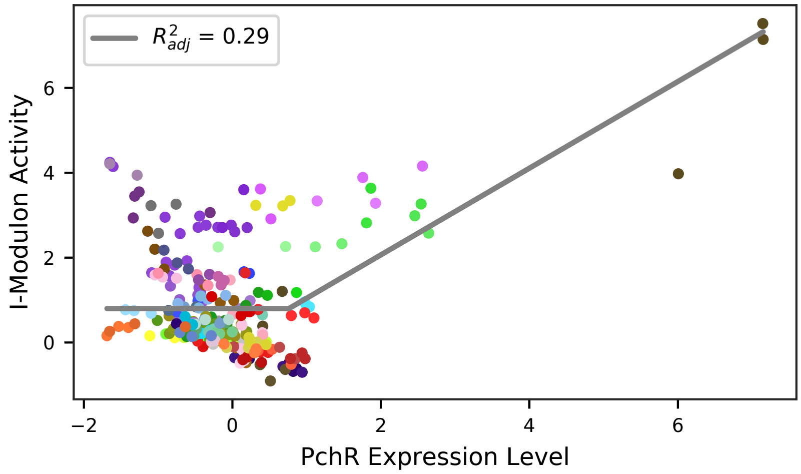

### 23 - Fur – Iron Limitation

Biological Function:
Production and transport of siderophores for iron chelation, replacement of ferredoxins in the ETC with flavodoxins, RNA chaperones for iron-limited conditions, heme degradation

Subset of known regulon:
Fur

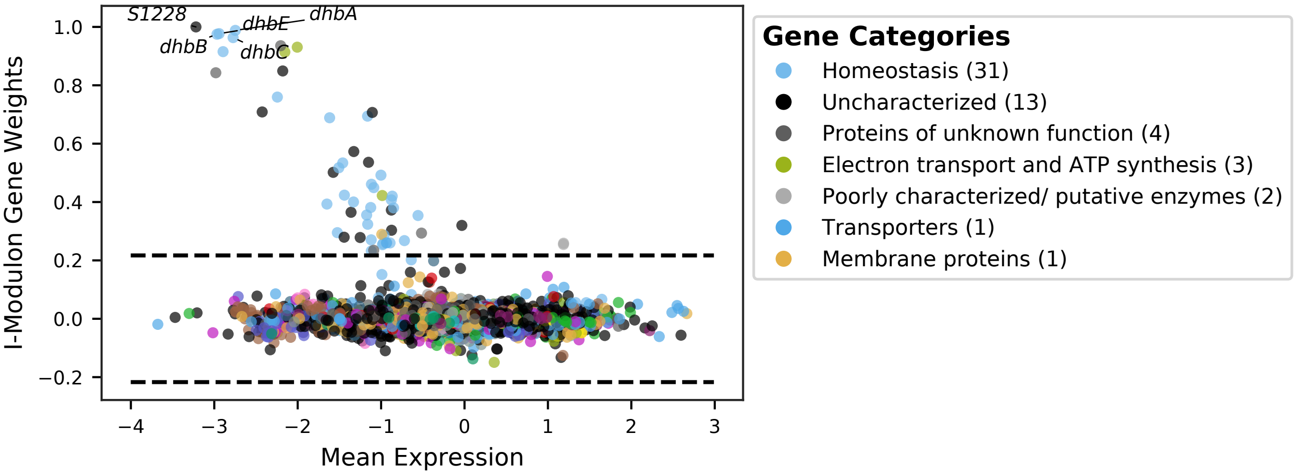


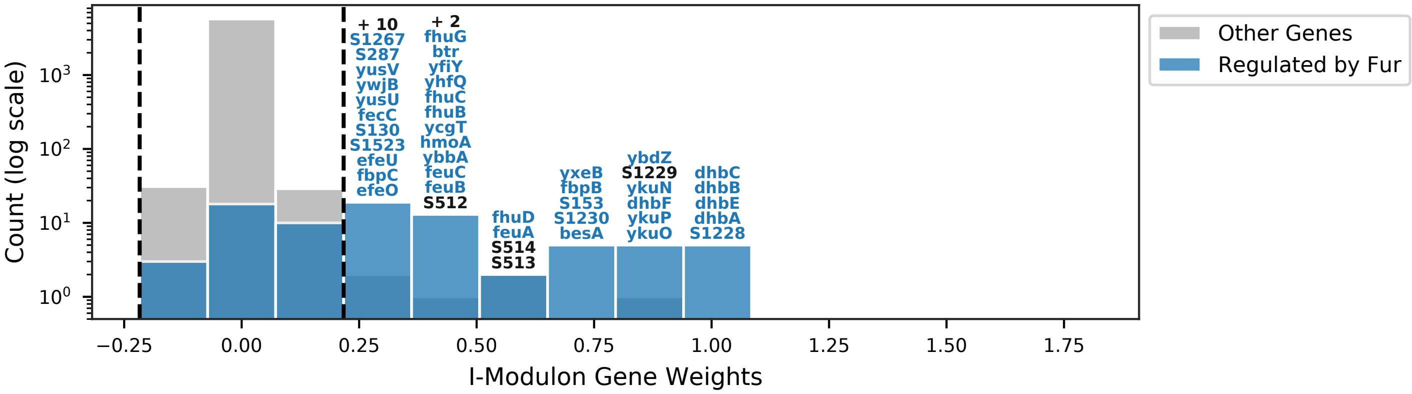


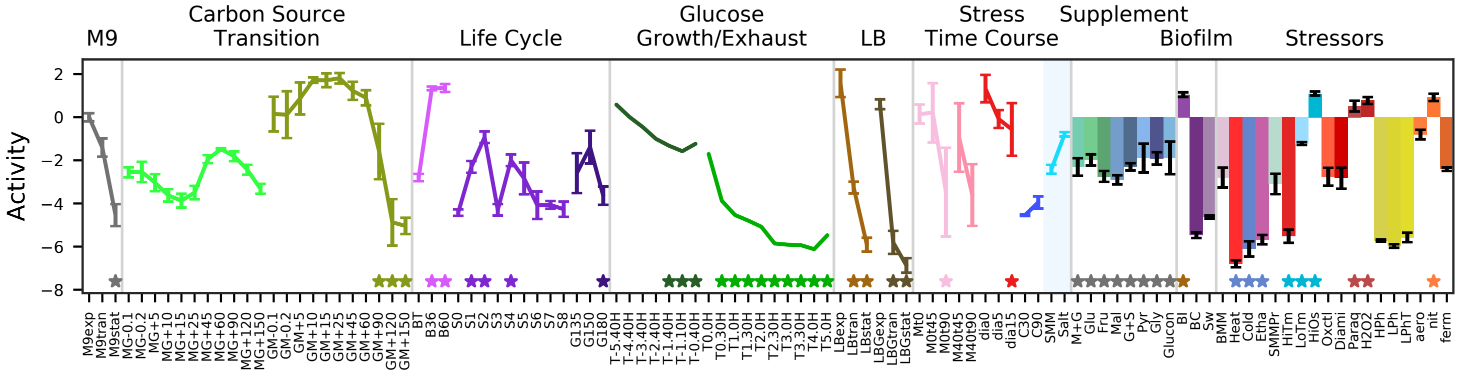


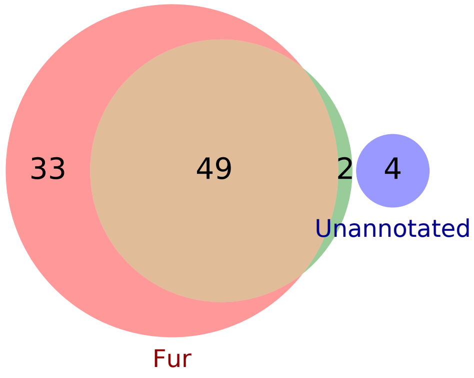


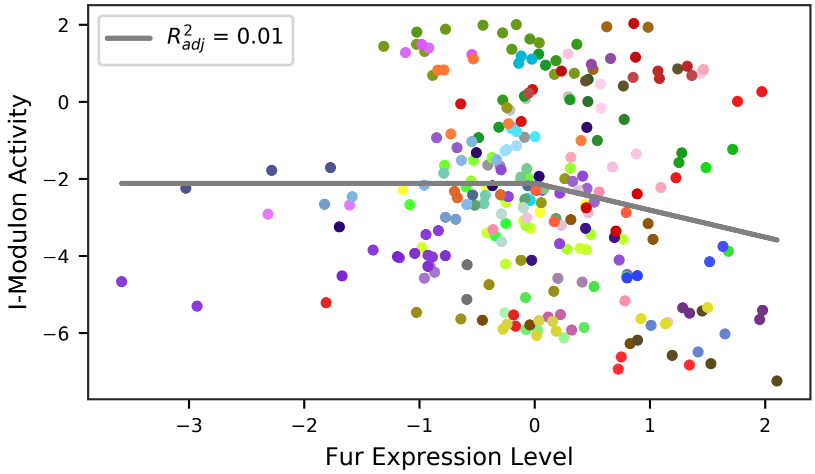

### 24 - PhoP-1 – Phosphate Limitation 1

Biological Function:
High-affinity phosphate uptake, teichuronic acid production pseudogene.

Subset of known regulon:
PhoP + SigA

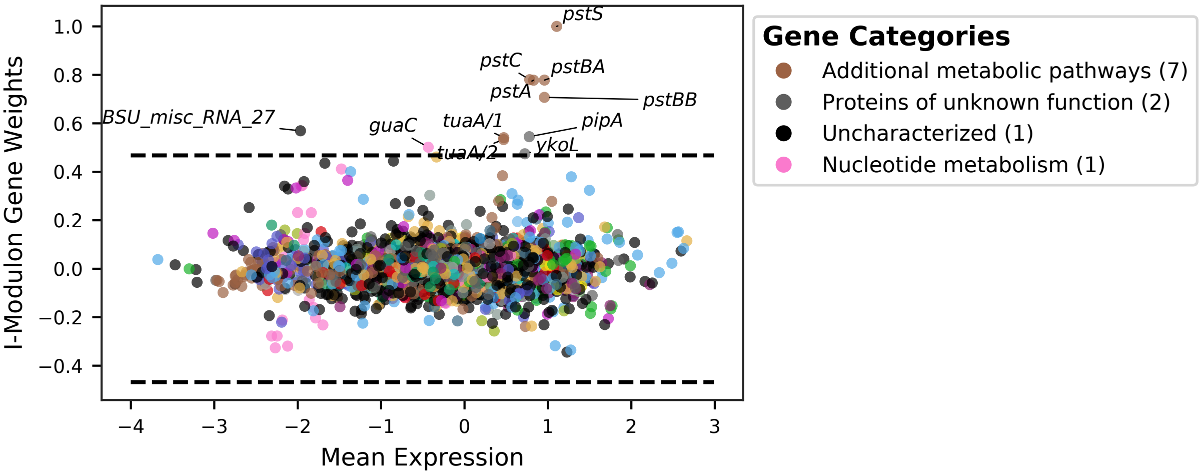


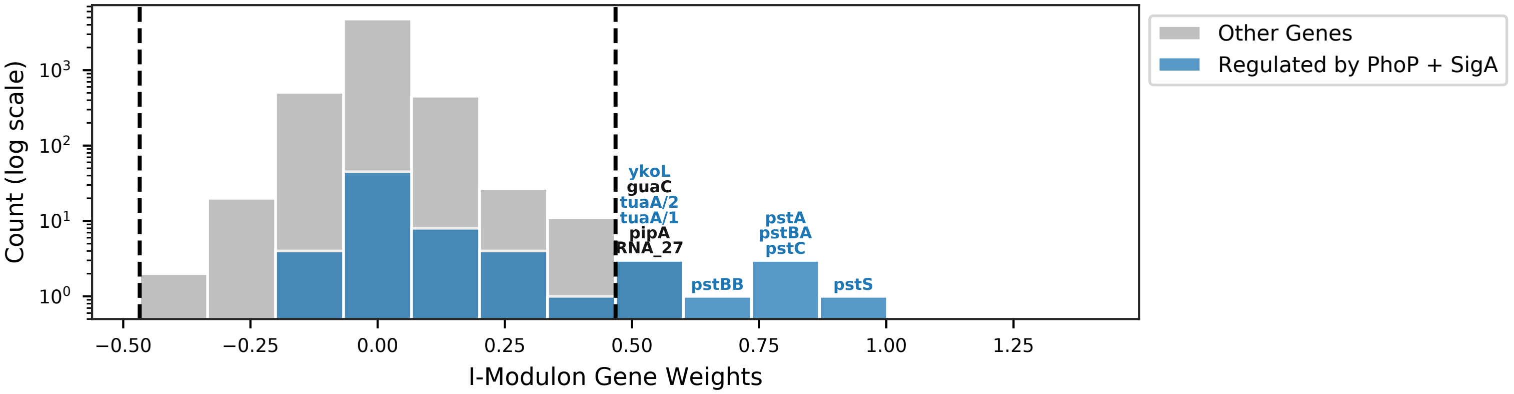


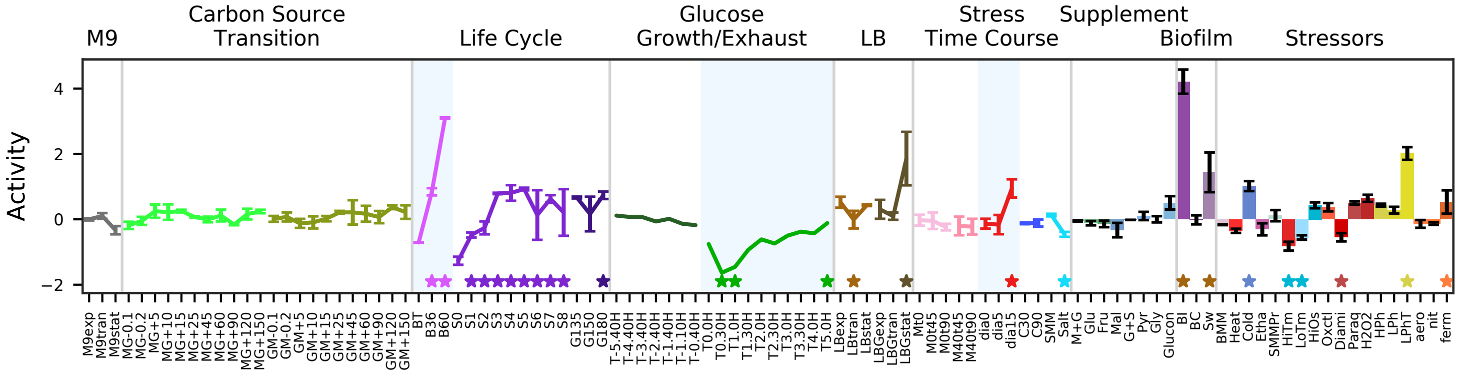


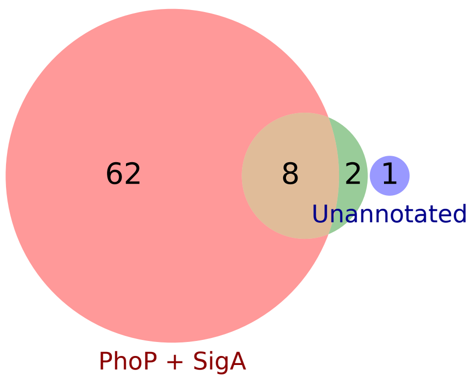


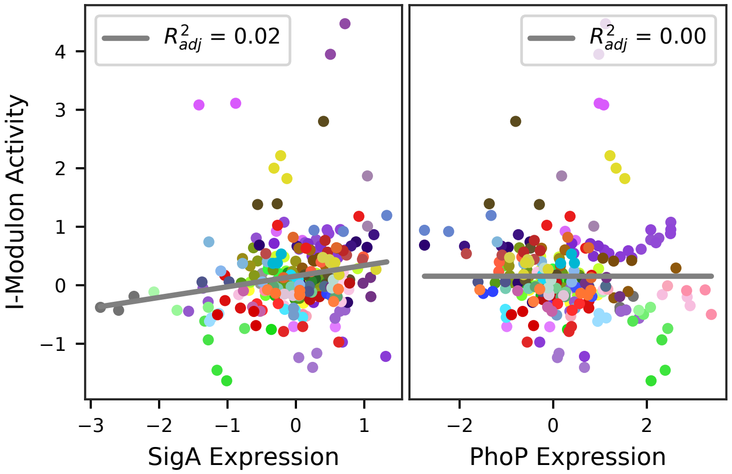

### 25 - PhoP-2 – Phosphate Limitation 2

Biological Function:
Degradation of wall teichoic acid to salvage phosphate, synthesis of teichuronic acid to replace wall, high-affinity phosphate uptake.

Subset of known regulon:
PhoP + SigA

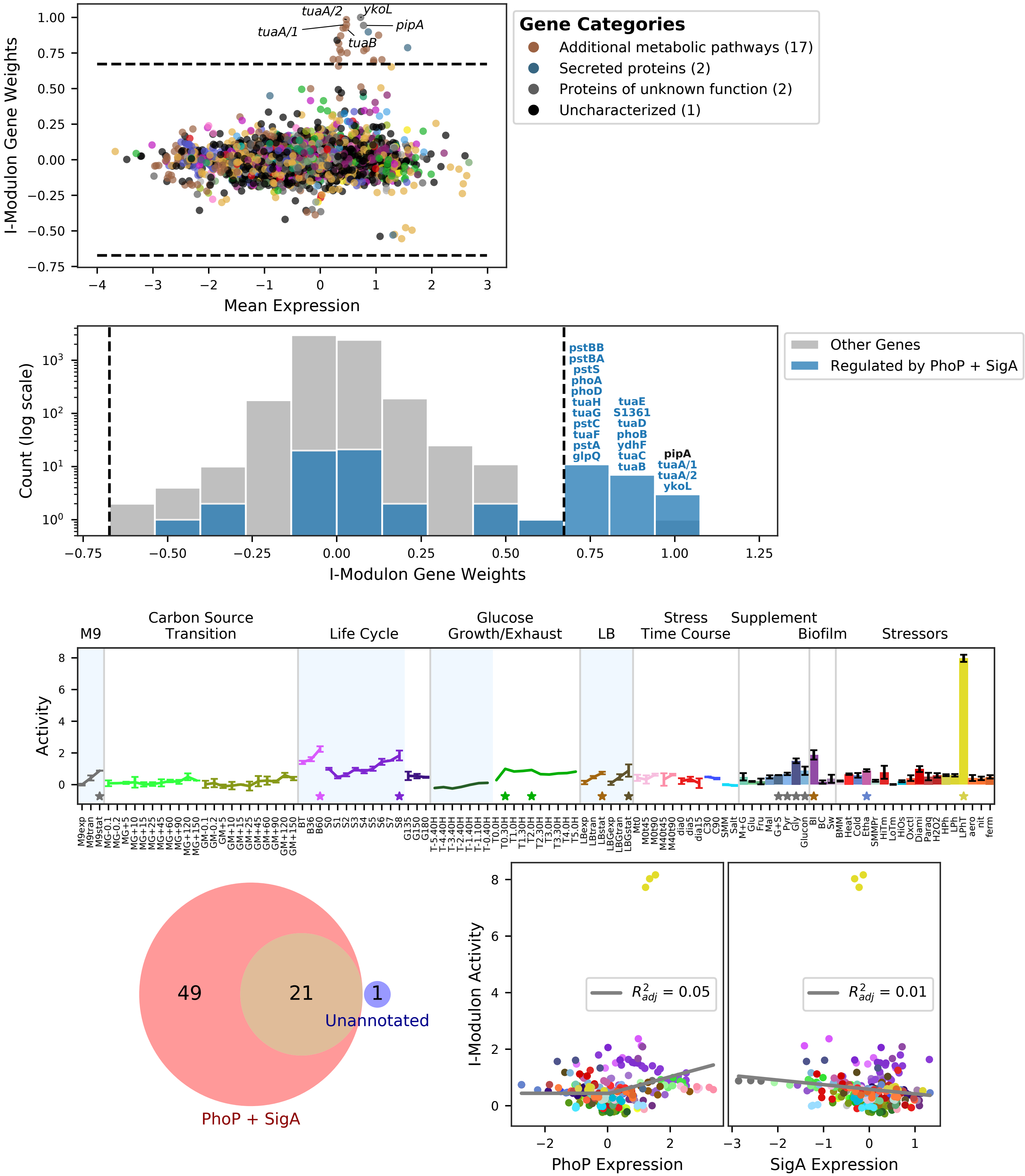

### 26 - Zur – Zinc Limitation

Biological Function:
Zinc uptake, zinc metallochaperones, alternatives to zinc-containing proteins

Well-defined regulon:
Zur

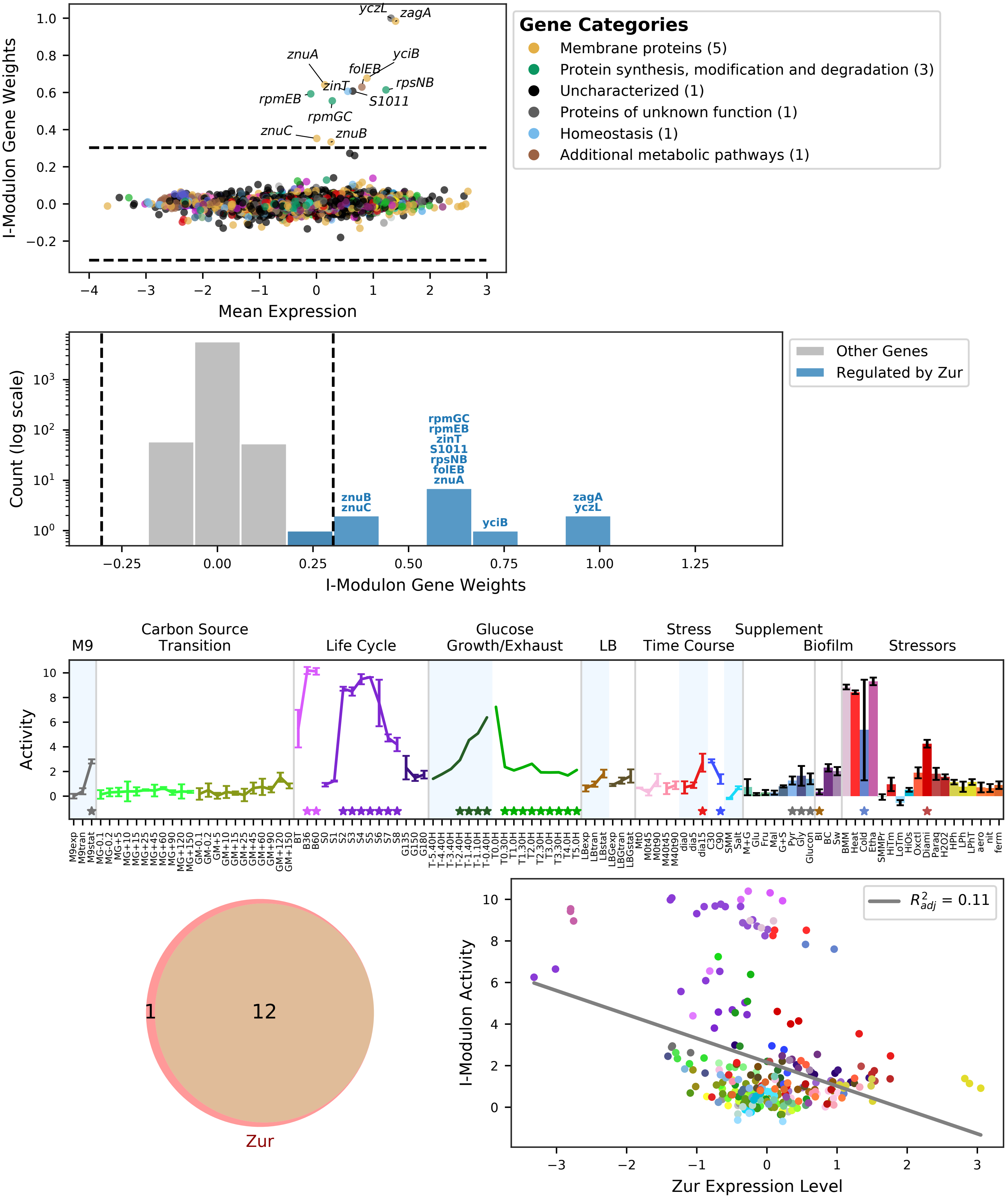

### 27 - CzrA – Toxic Cation Export

Biological Function:
Resistance to Zn, Cu, Co, Ni, and Cd

Well-defined regulon:
CzrA

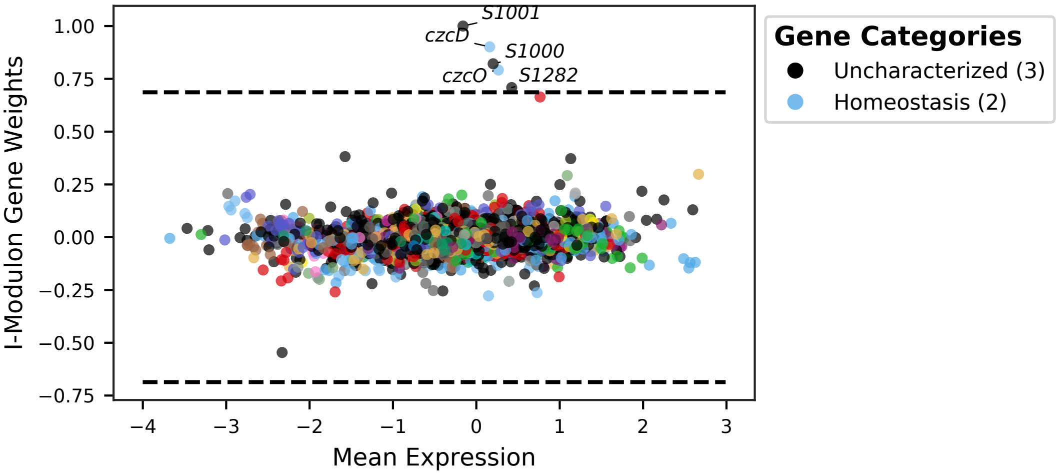


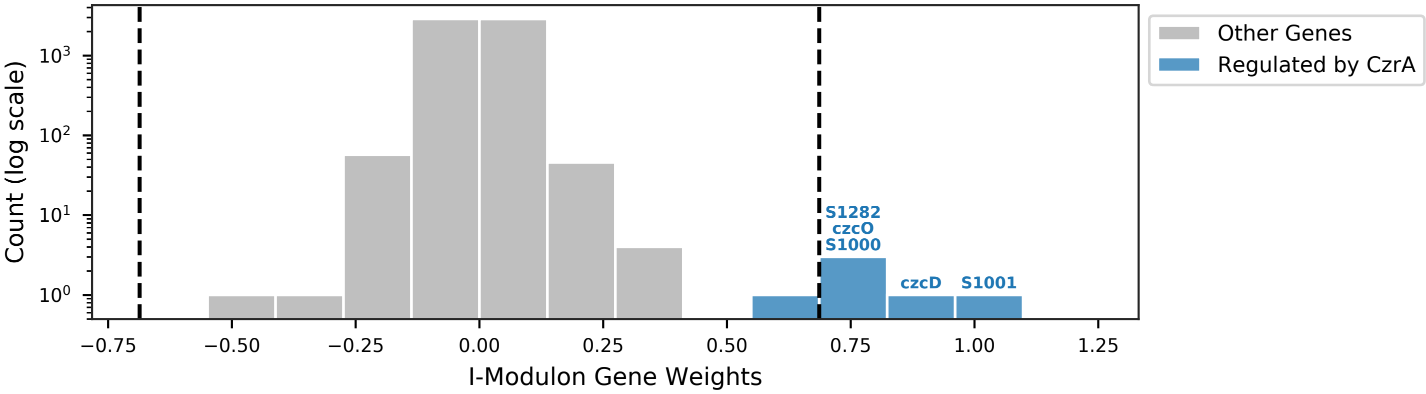


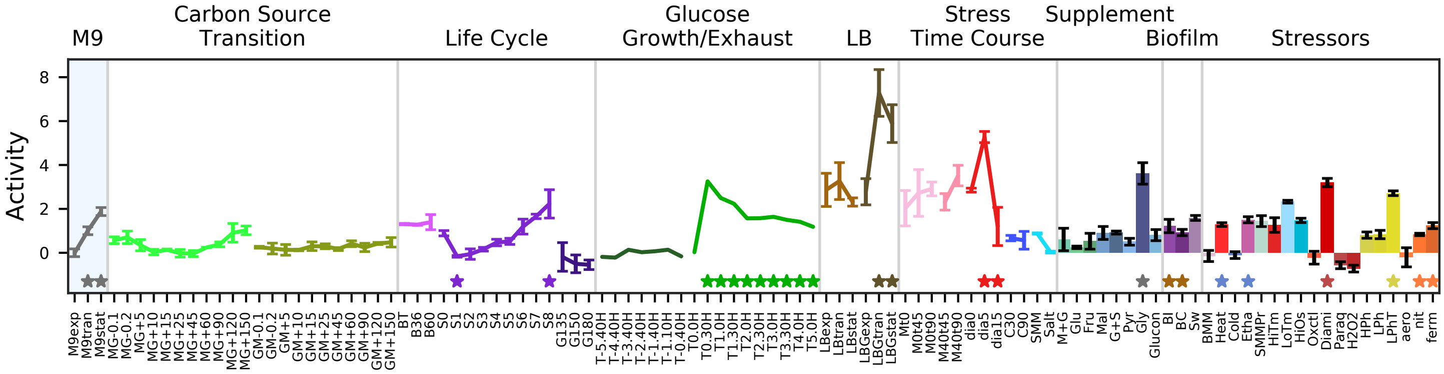


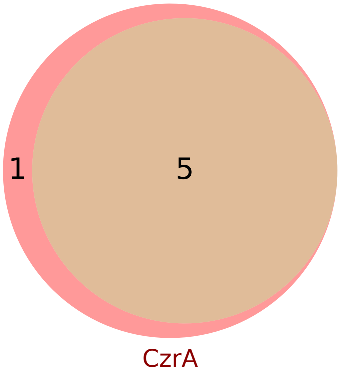


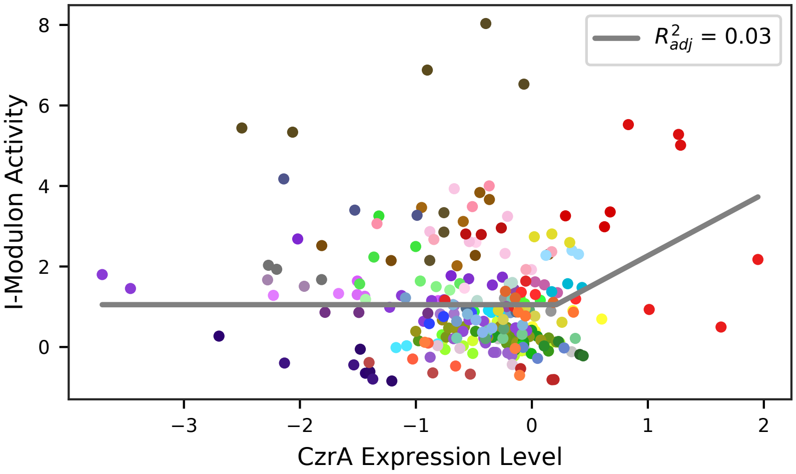

### 28 - CymR – Sulfur

Biological Function:
Sulfur metabolism in response to oxidative stress. Utilization and detoxification of S-(2-succino)cysteine, methionine-cysteine conversion, siroheme biosynthesis

Well-defined regulon:
CymR

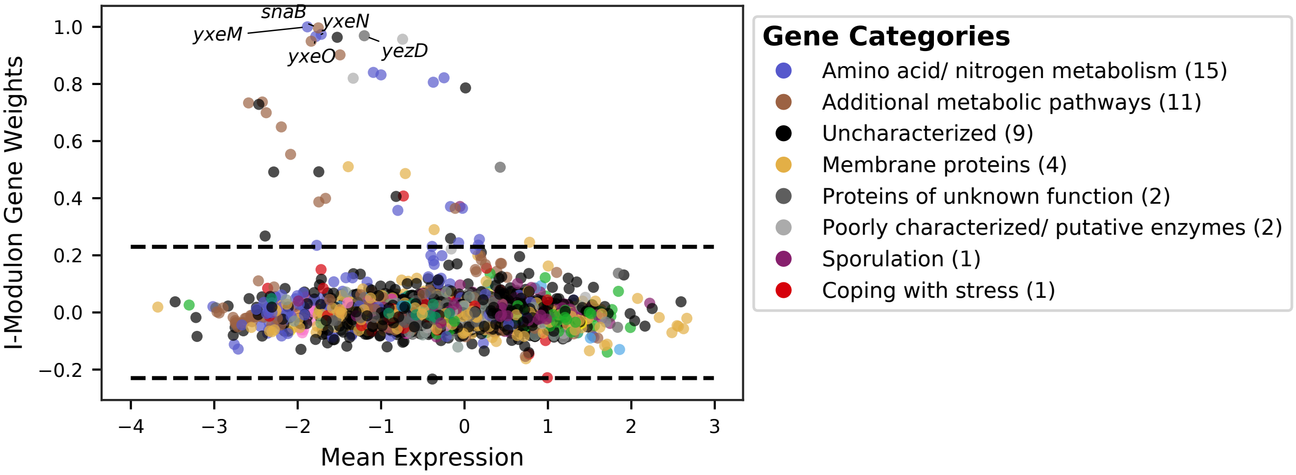


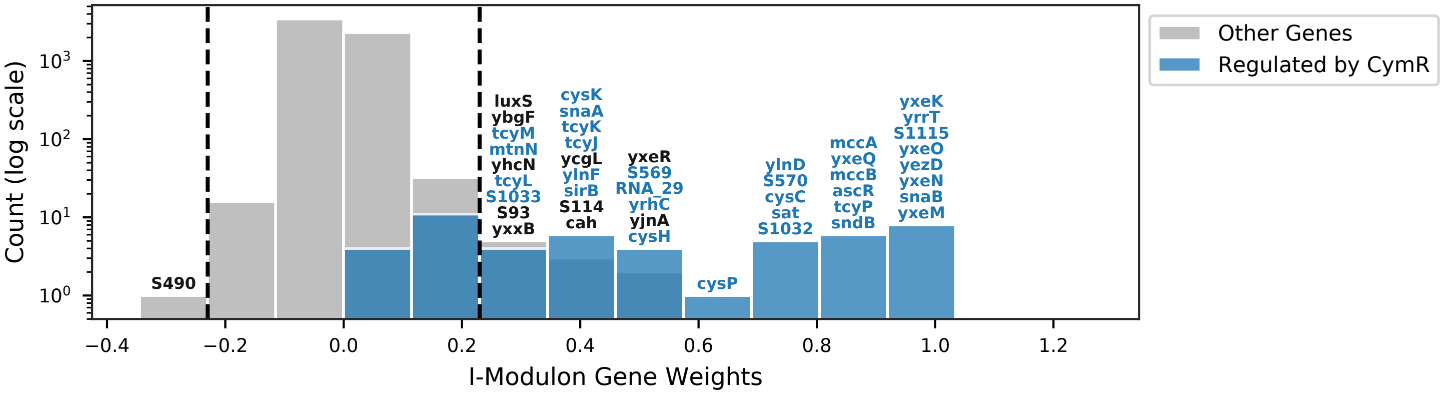


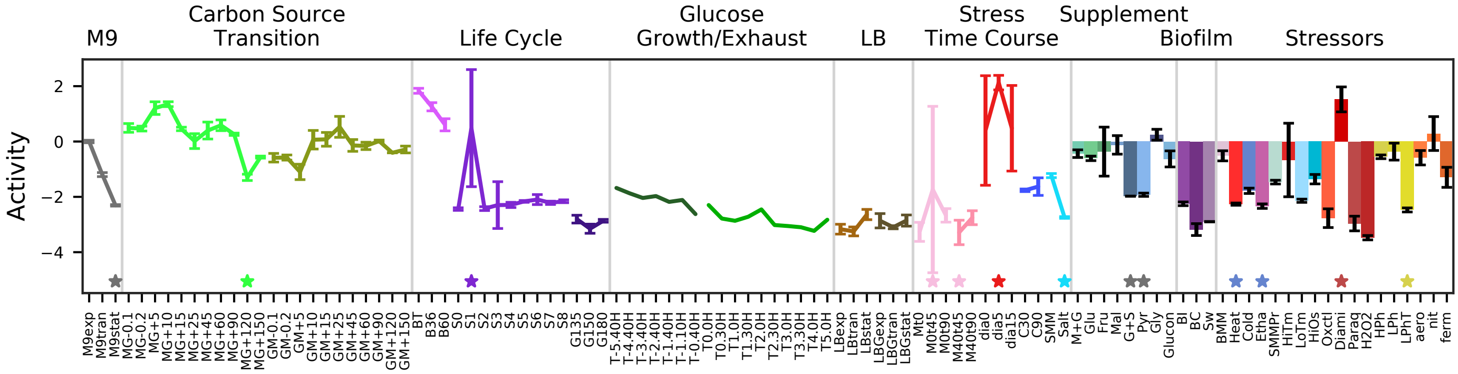


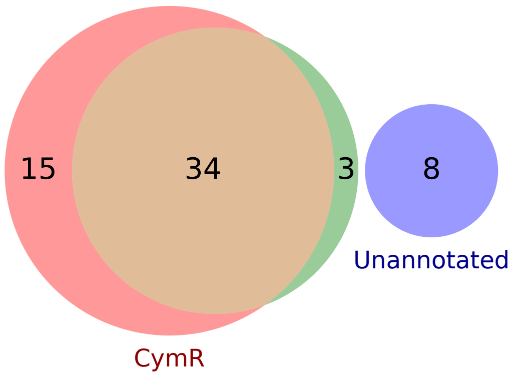


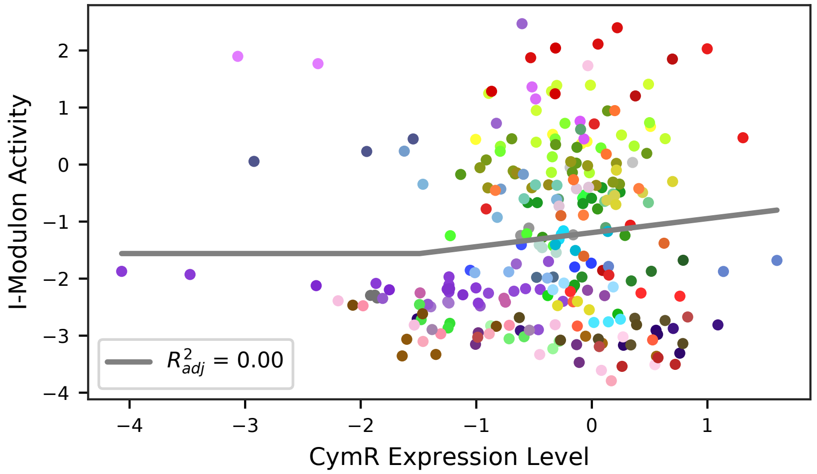

### 29 - Yrk – Putative Sulfur Carriers

Biological Function:
Putative sulfur carriers. High activity under diamide stress

No known regulator

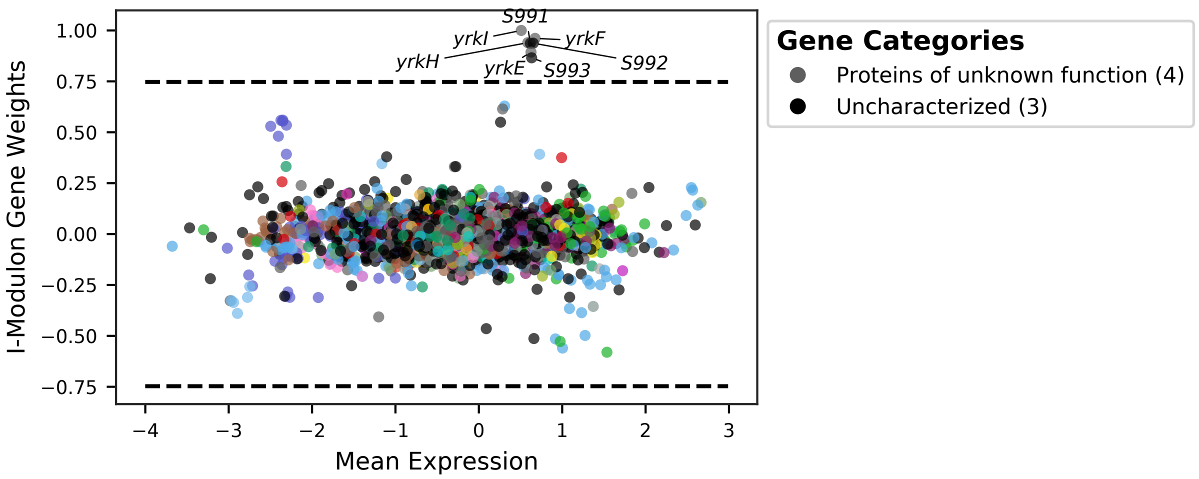


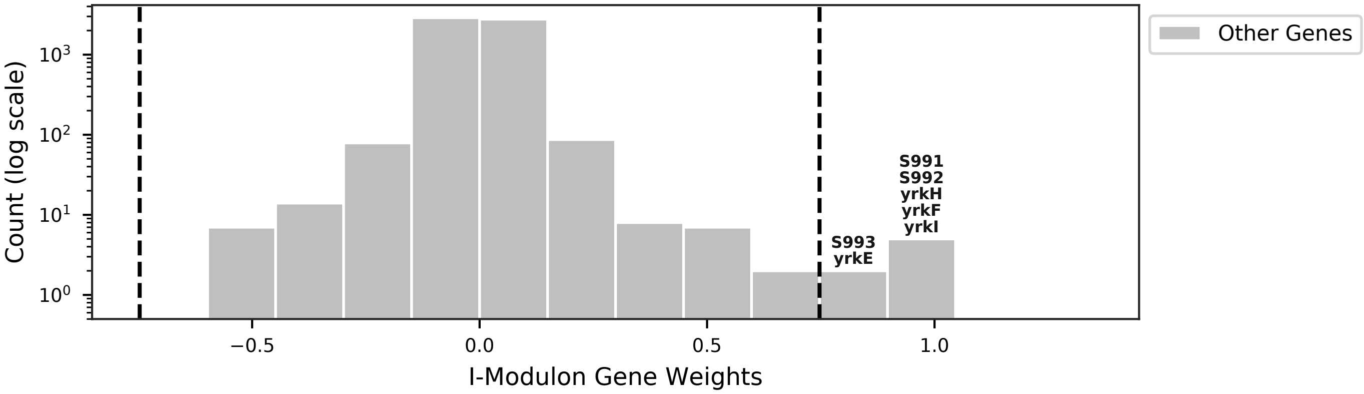


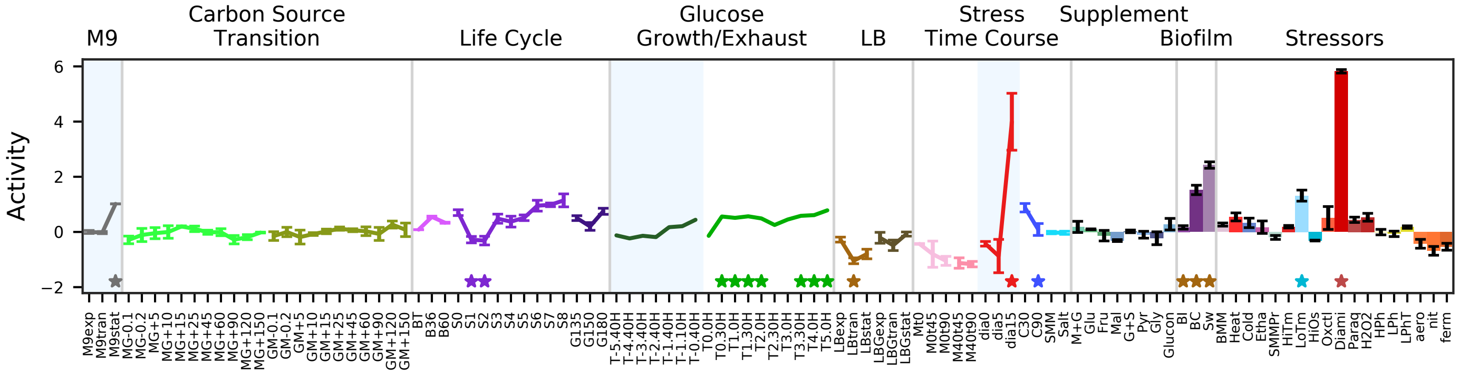


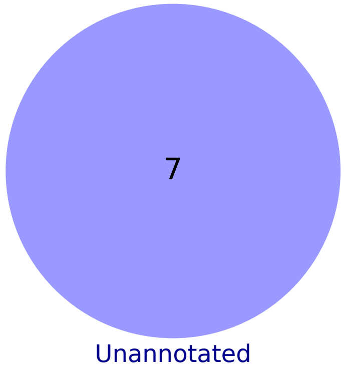

### 30 - ResD – Oxygen Limitation

Biological Function:
Anaerobic nitrate respiration, bacteriocin production, heme, cytochrome, and ATP synthesis, copper homeostasis

Enriched for known regulon:
ResD

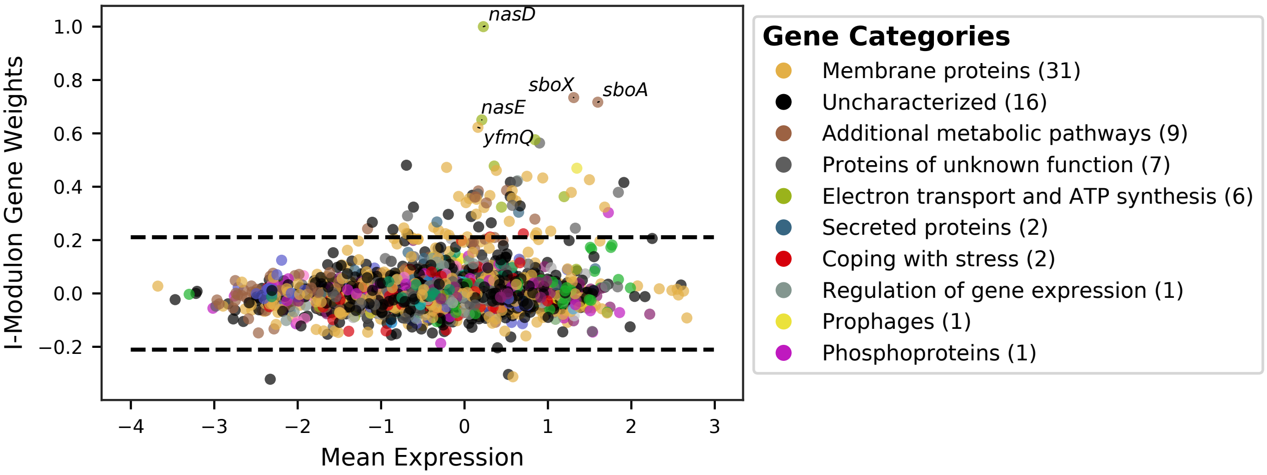


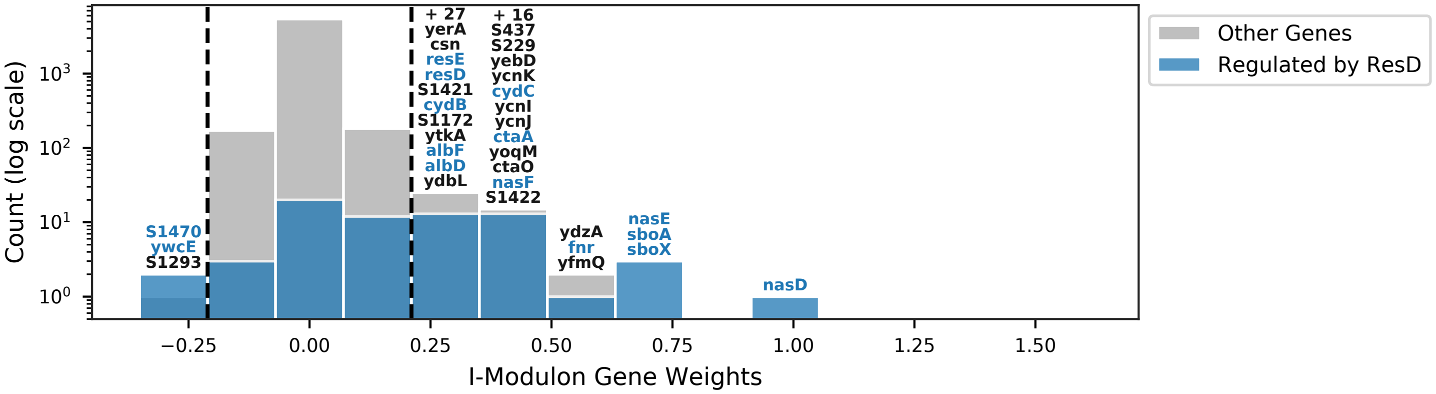


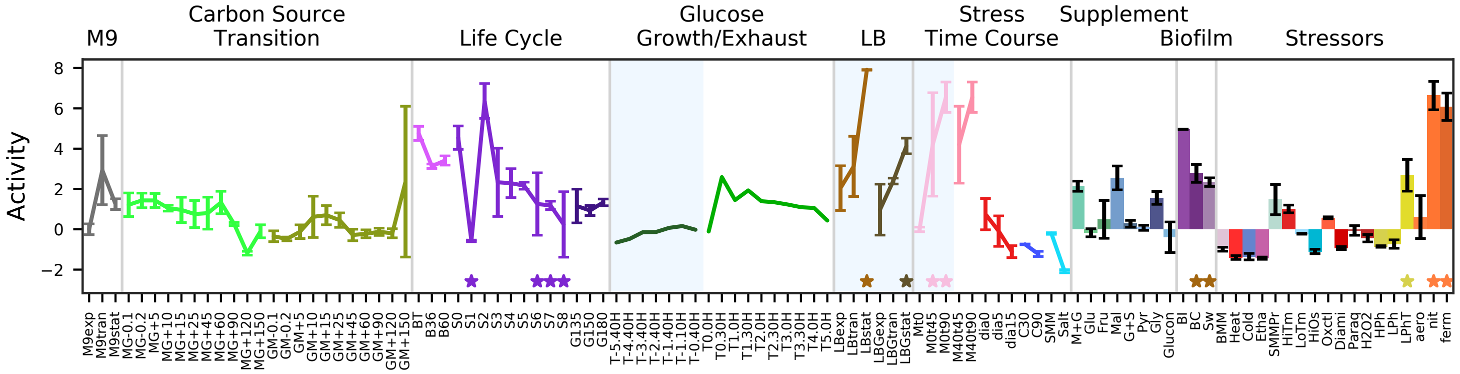


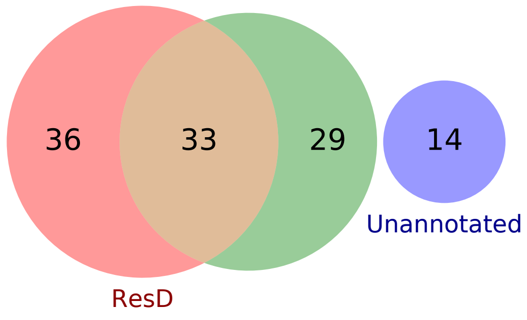


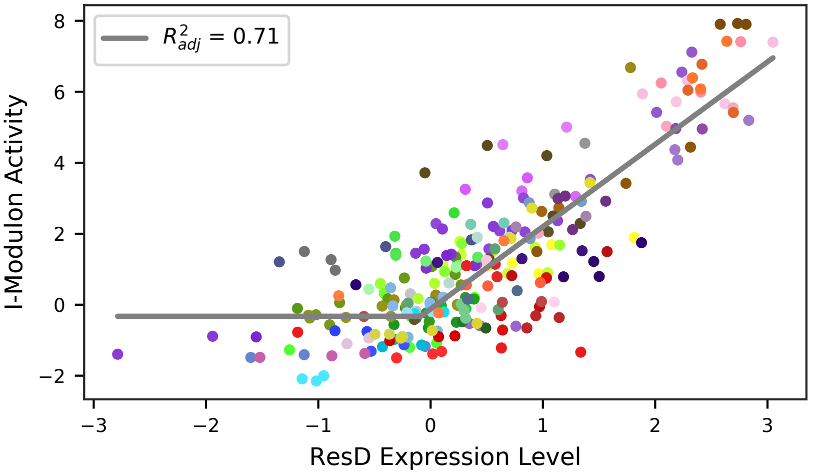

### 31 - Fnr – Nitrate Respiration

Biological Function:
Nitrate respiration and nitrite extrusion

Well-defined regulon:
Fnr + SigA

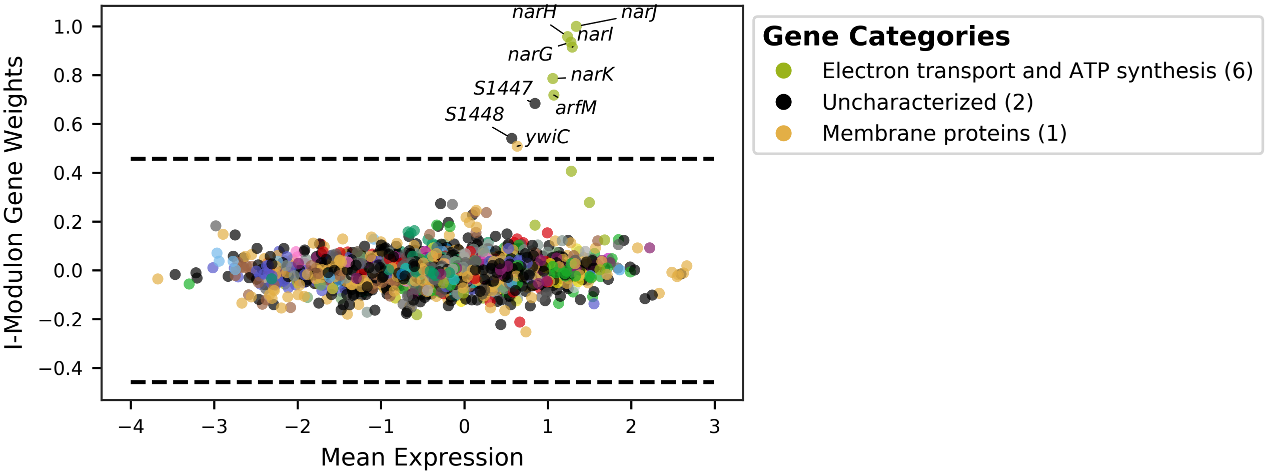


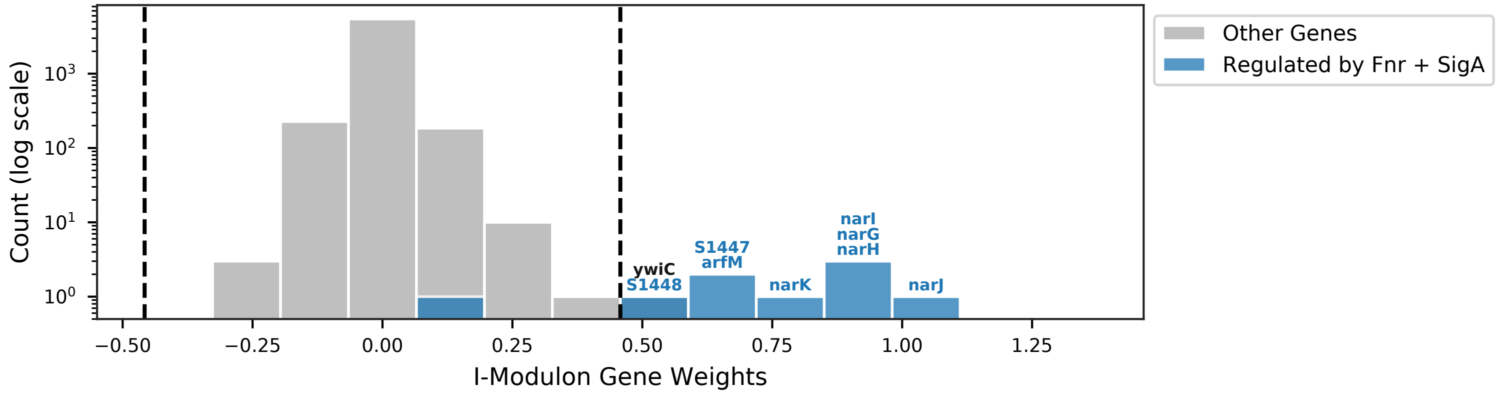


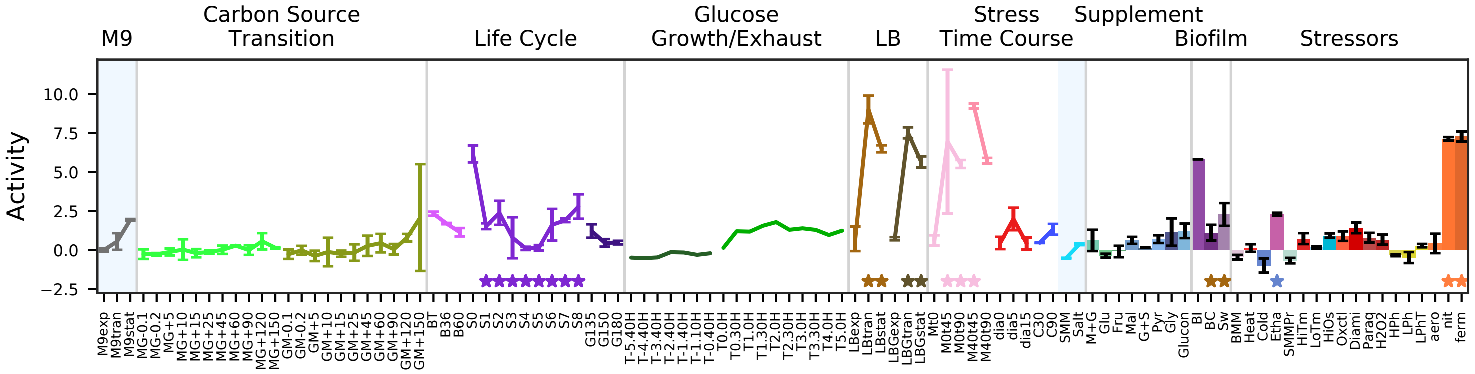


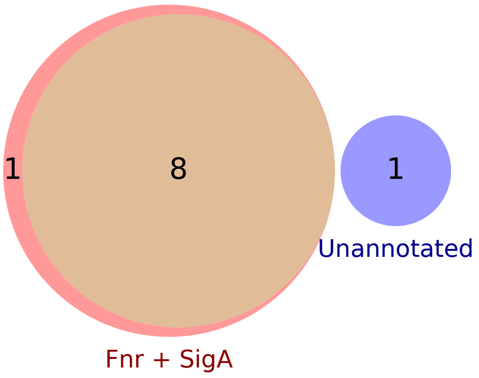


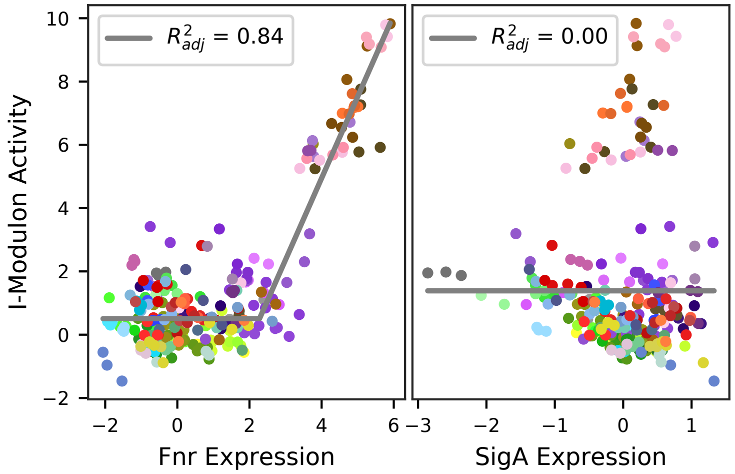

### 32 - Rex – Overflow Metabolism

Biological Function:
Overflow metabolism (LDH) and cytochrome BD oxidation  for respiration in anaerobic conditions

Subset of known regulon:
Rex

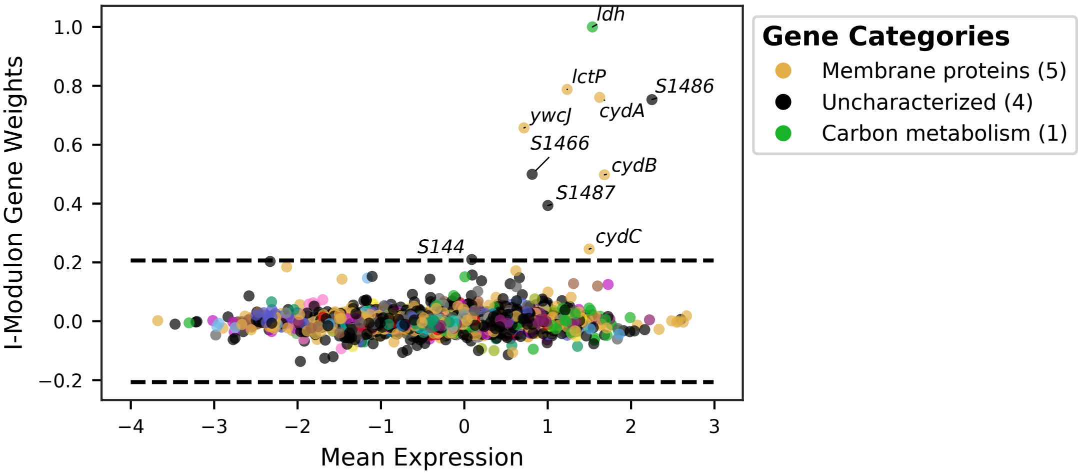


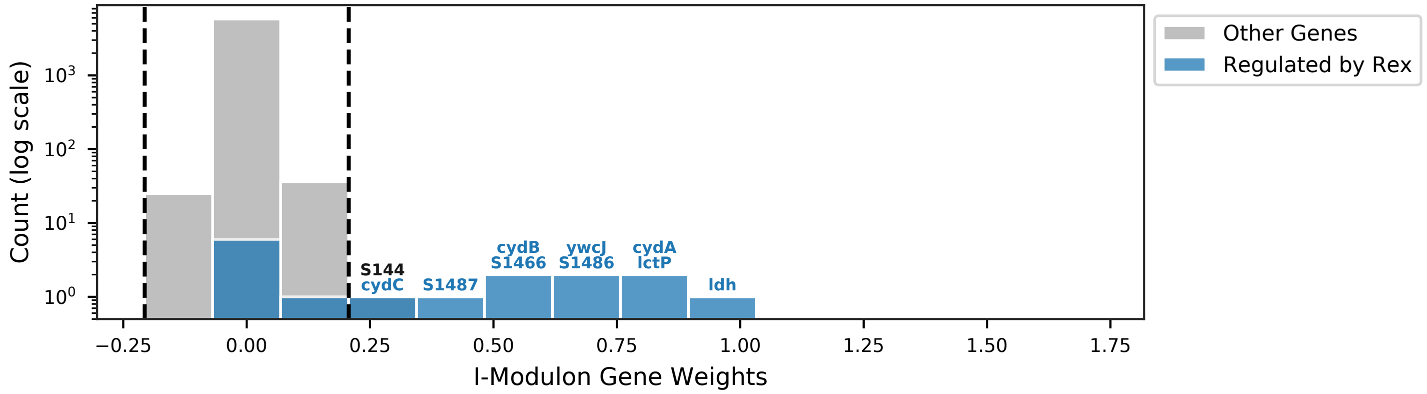


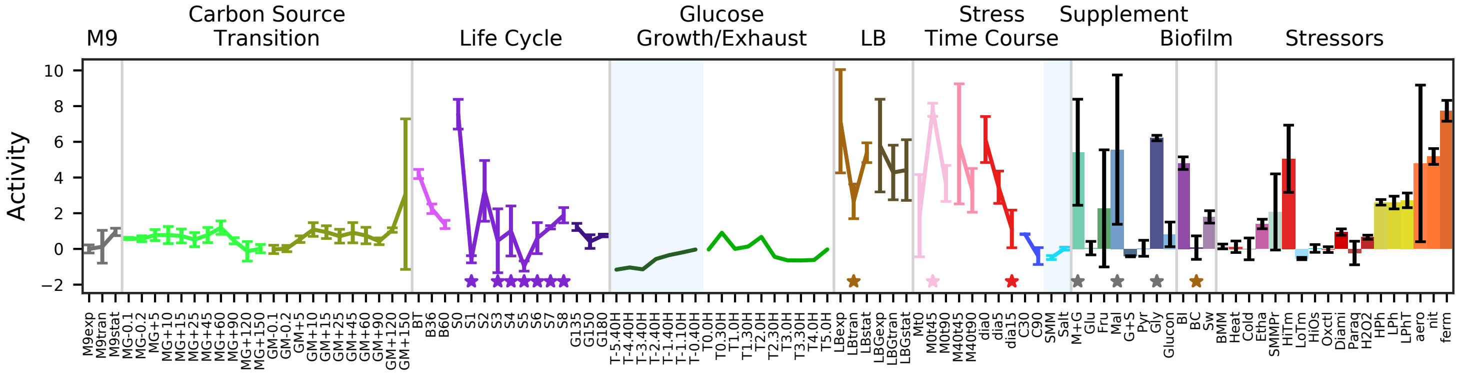


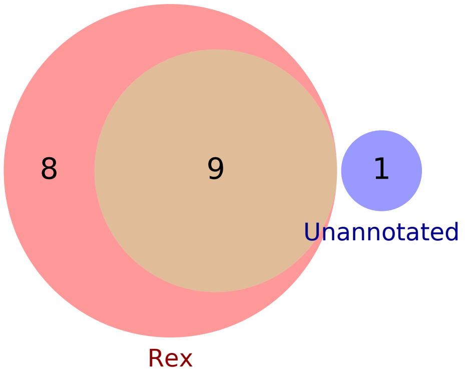


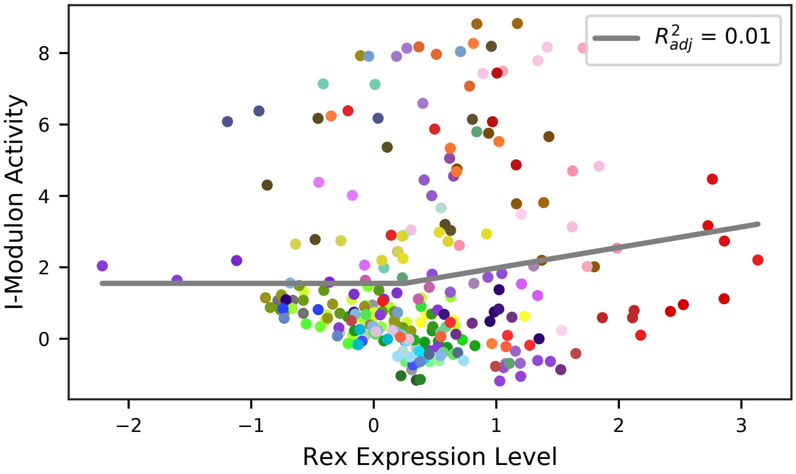

### 33 - LexA – DNA Repair

Biological Function:
SOS response for DNA protection and repair

Subset of known regulon:
LexA

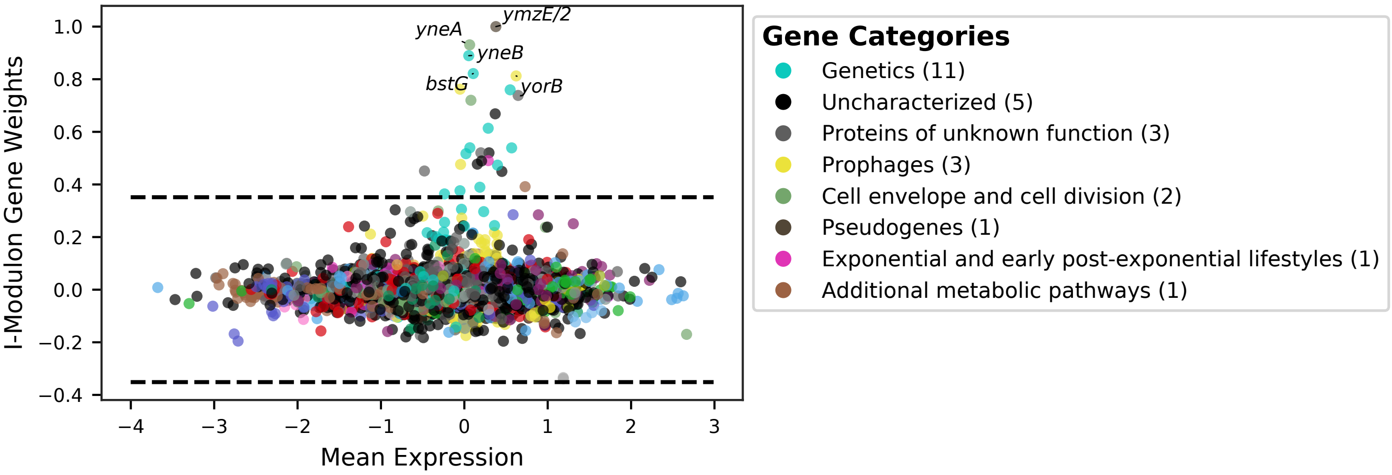


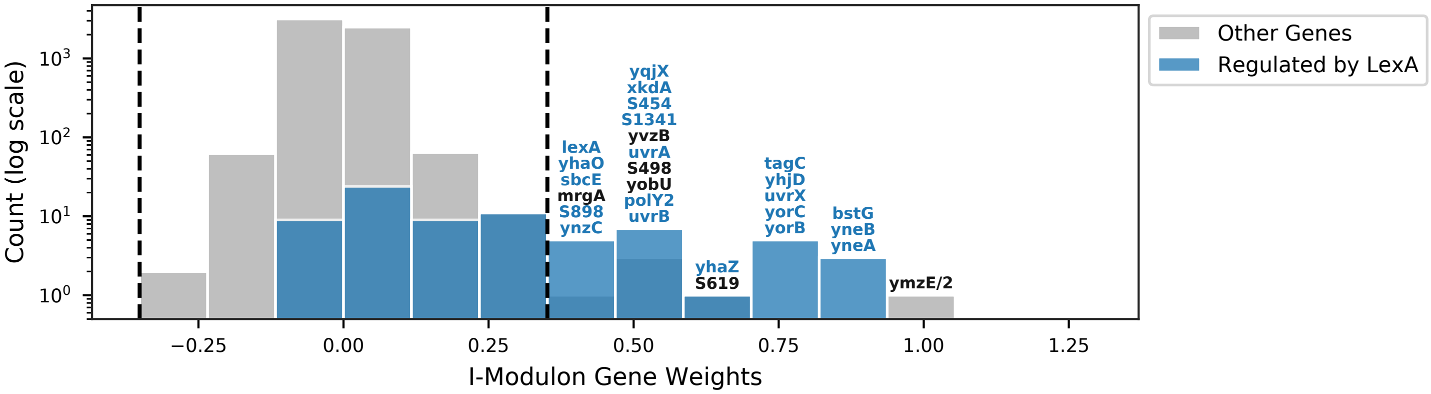


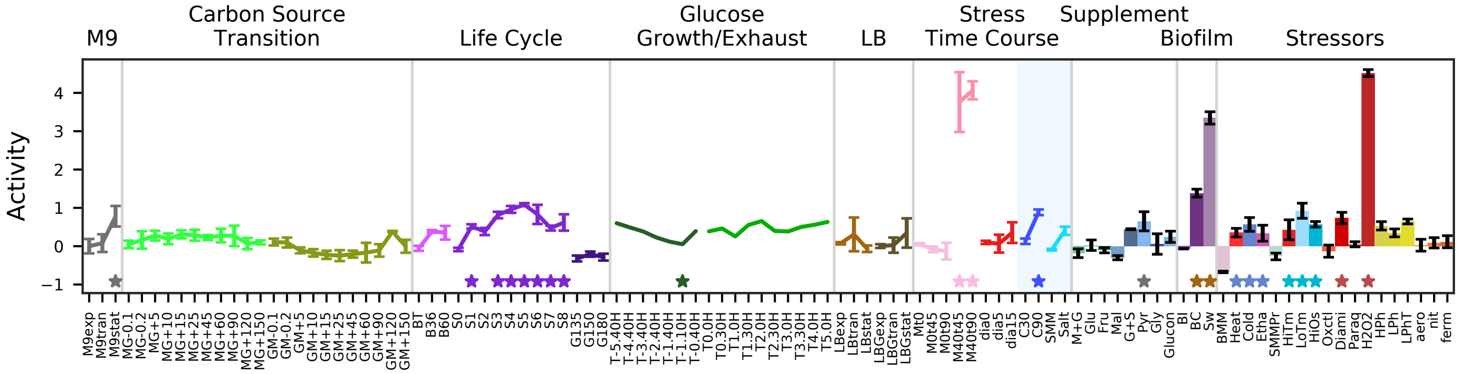


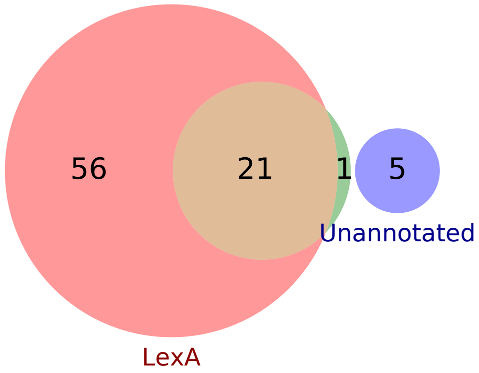


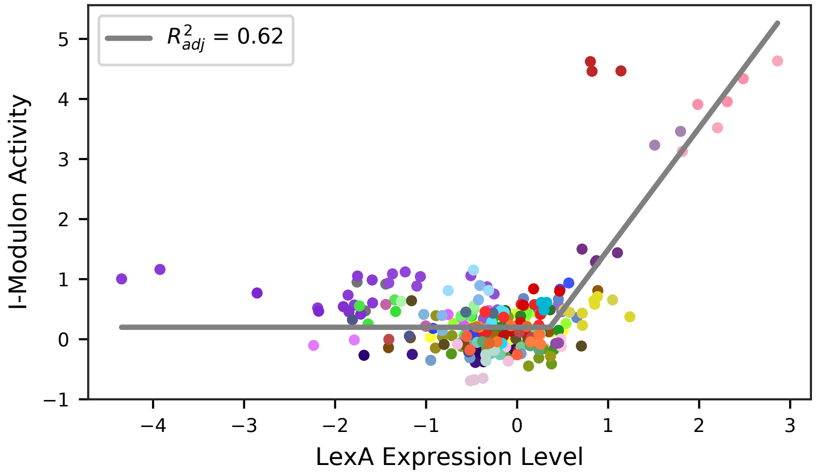

### 34 - SigW – Cell Wall Stress

Biological Function:
Adaptation to membrane active agents such as cefuroxime, antimicrobials from B. amyloliquefaciens, sdpC, and nisin. Control of membrane fluidity. Contains many unknown proteins.

Subset of known regulon:
SigW

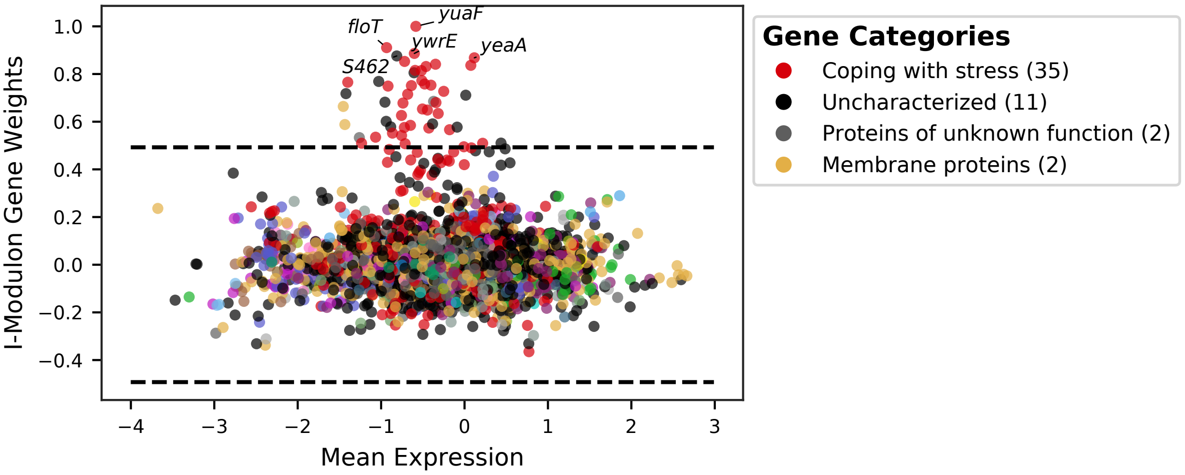


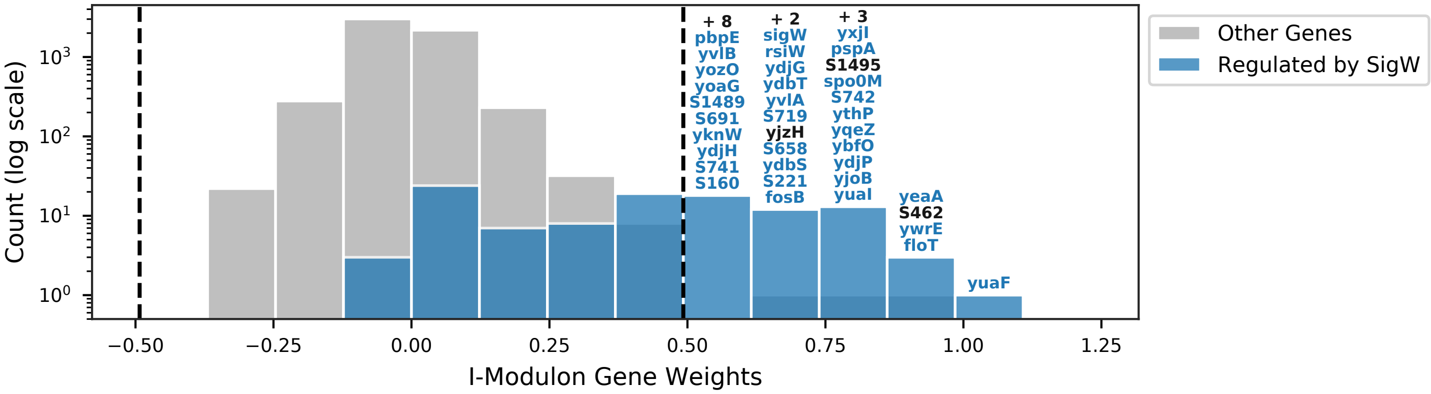


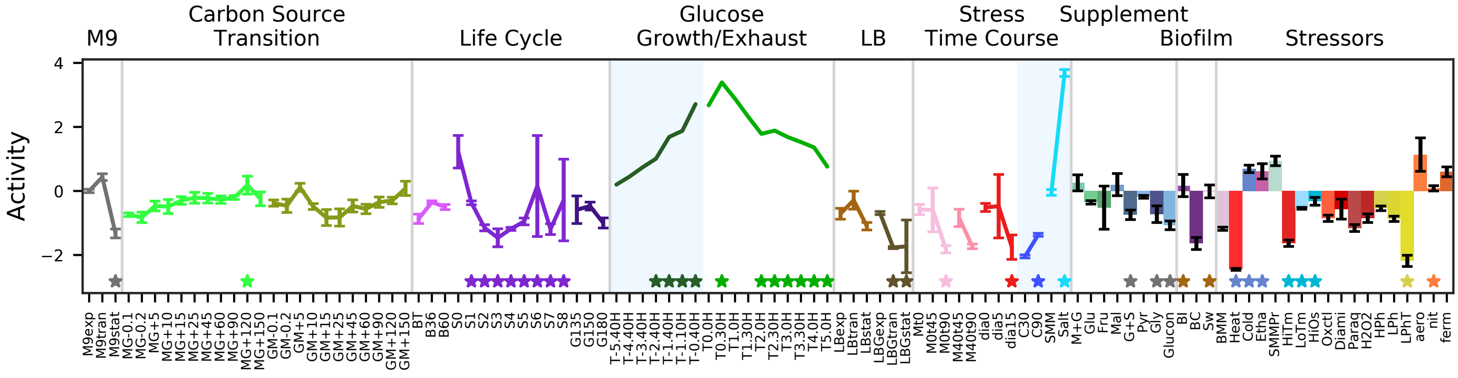


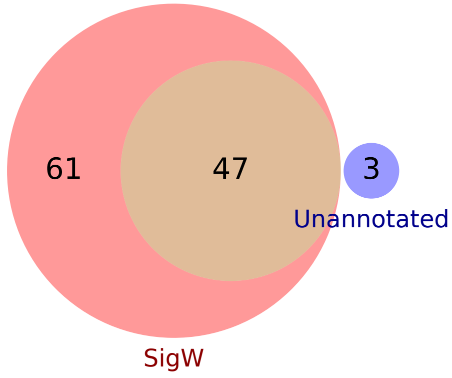


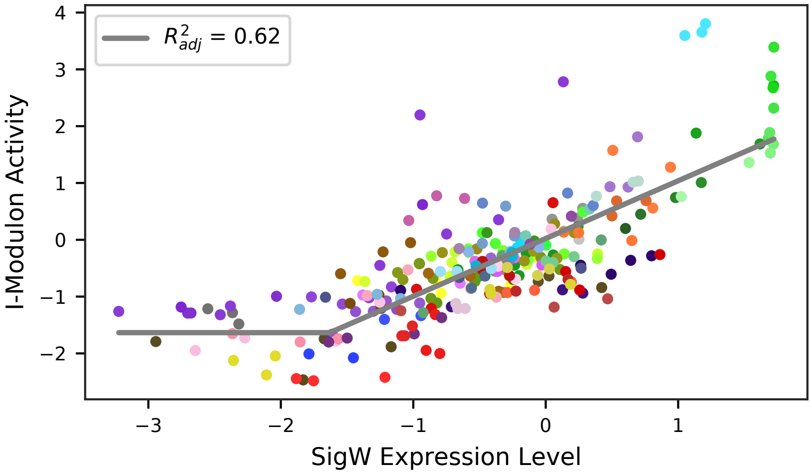

### 35 - SigM – Membrane Stress

Biological Function:
General stress proteins, alarmones, lipid carriers and flippases, DNA repair, inhibition of septation. Contains many UTRs and some unknown proteins.

Subset of known regulon:
SigM

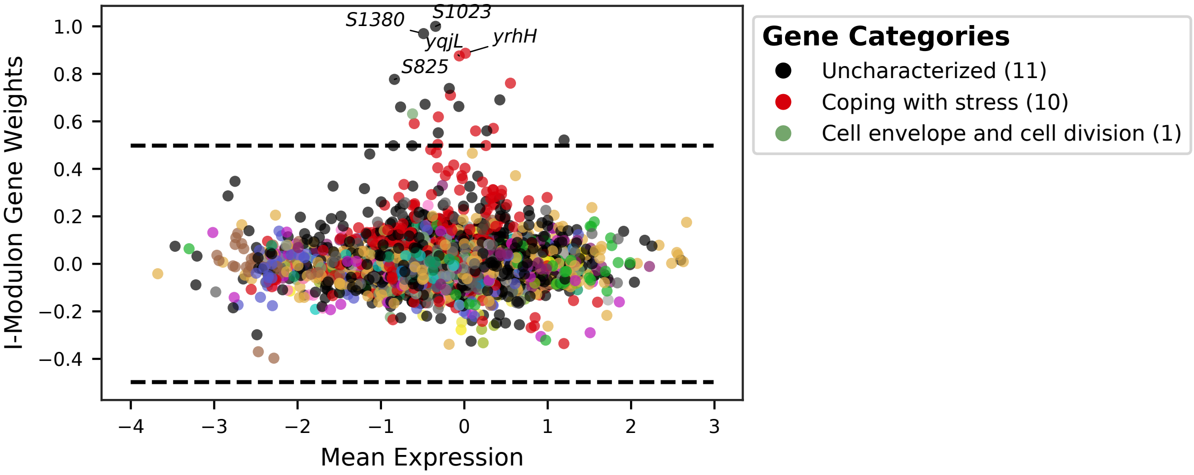


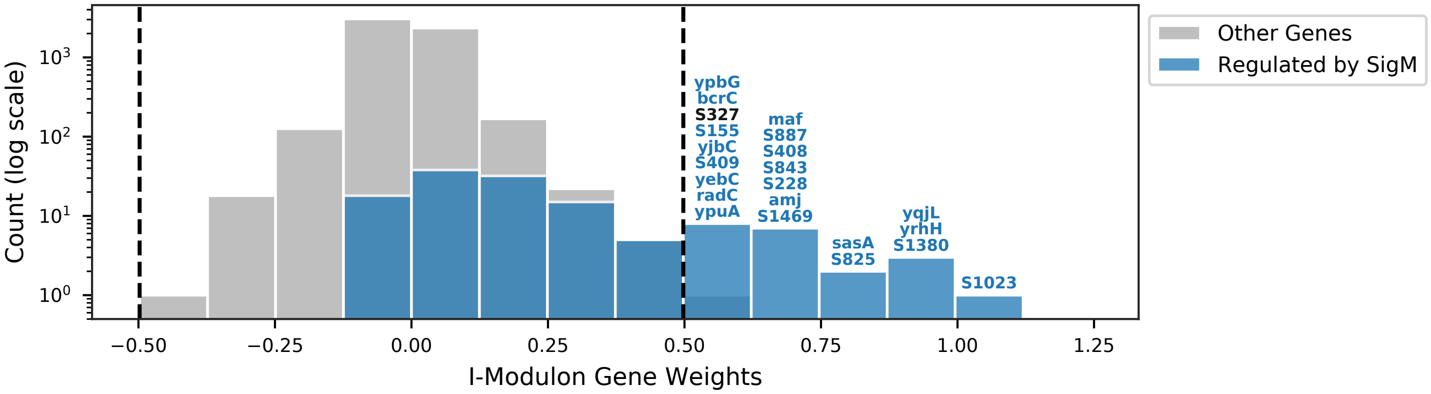


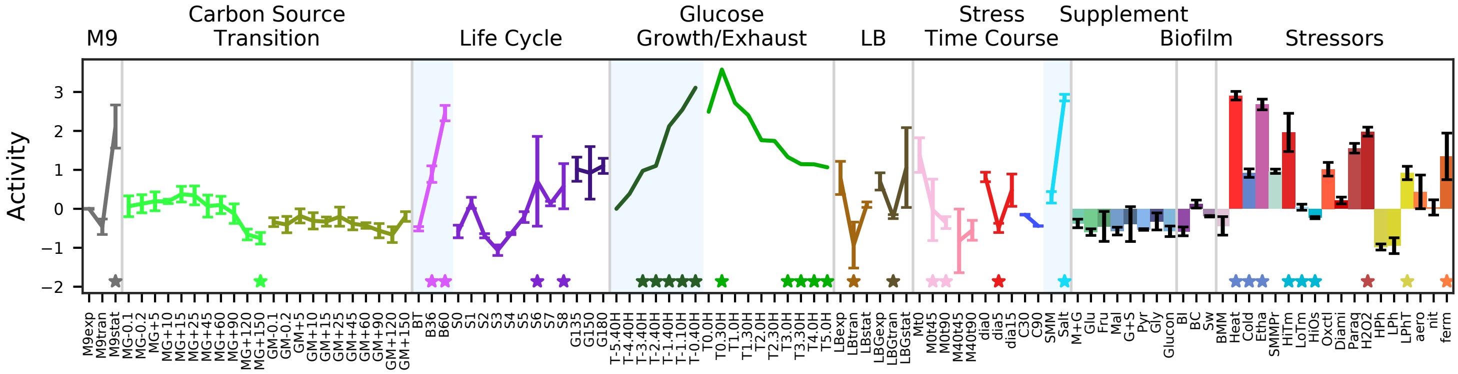


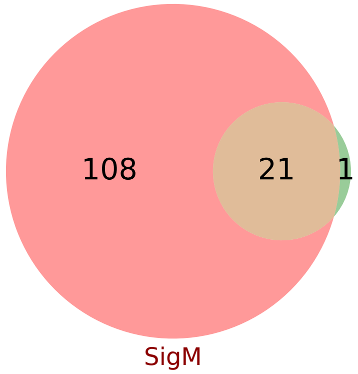


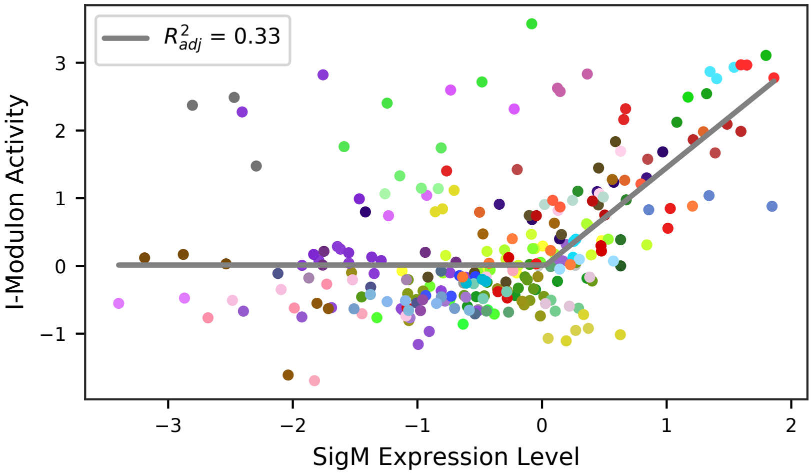

### 36 - Ybc – Uncharacterized Operon

Biological Function:
Contains one subunit of NADH dehydrogenase with several unknown genes. Categorized as prophage.
Responds to heat shock.

No known regulator

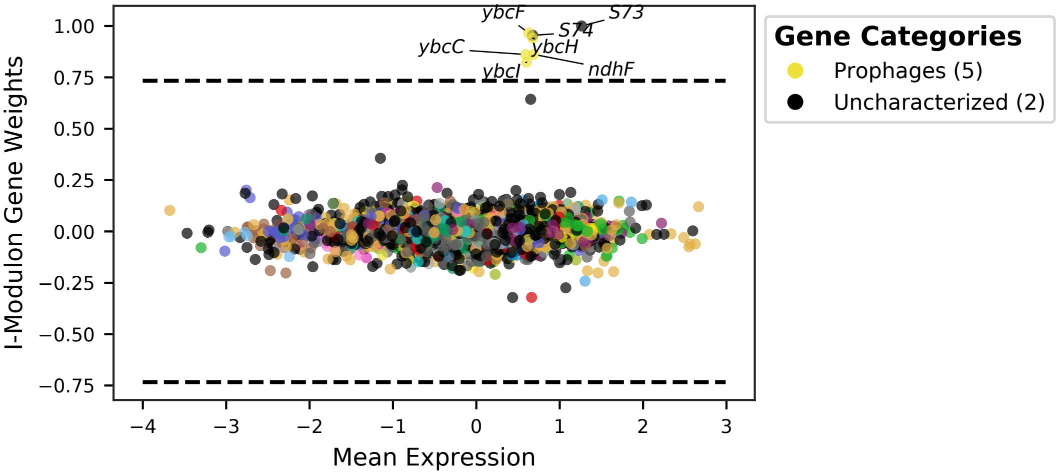


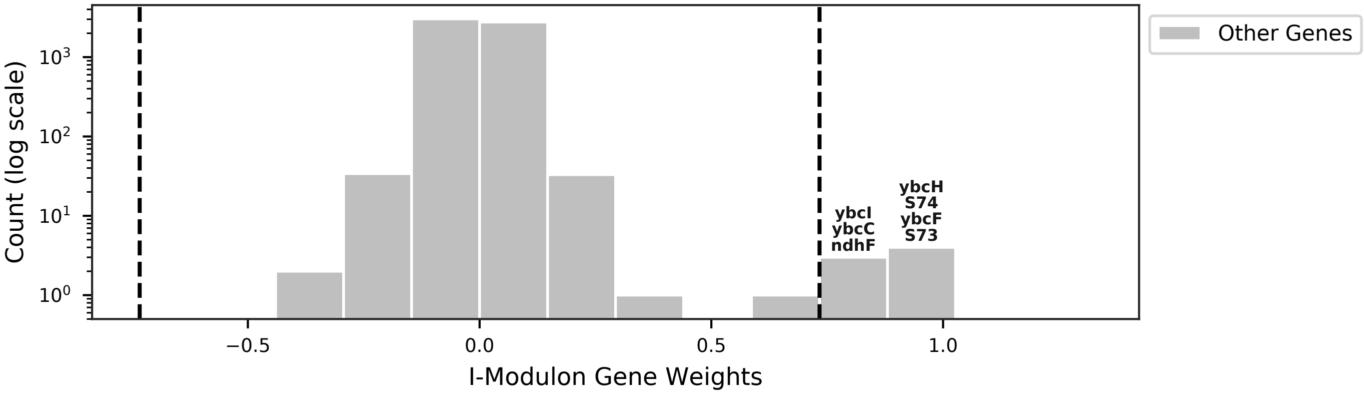


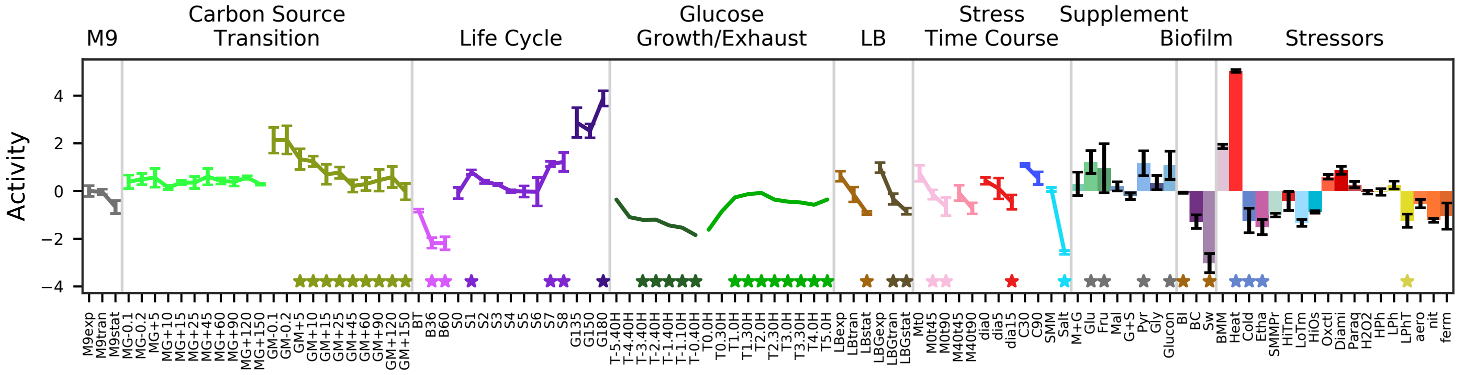


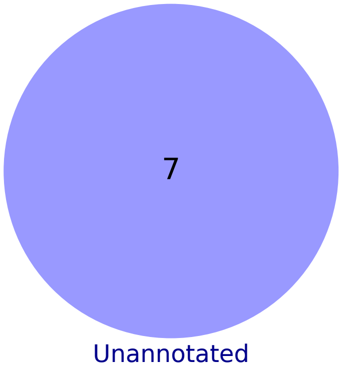

### 37 - SigB-1 – General Stress 1

Biological Function:
Survival of general stressors including ethanol, salt, paraquat, heat, and freeze-thaw

Subset of known regulon:
SigB

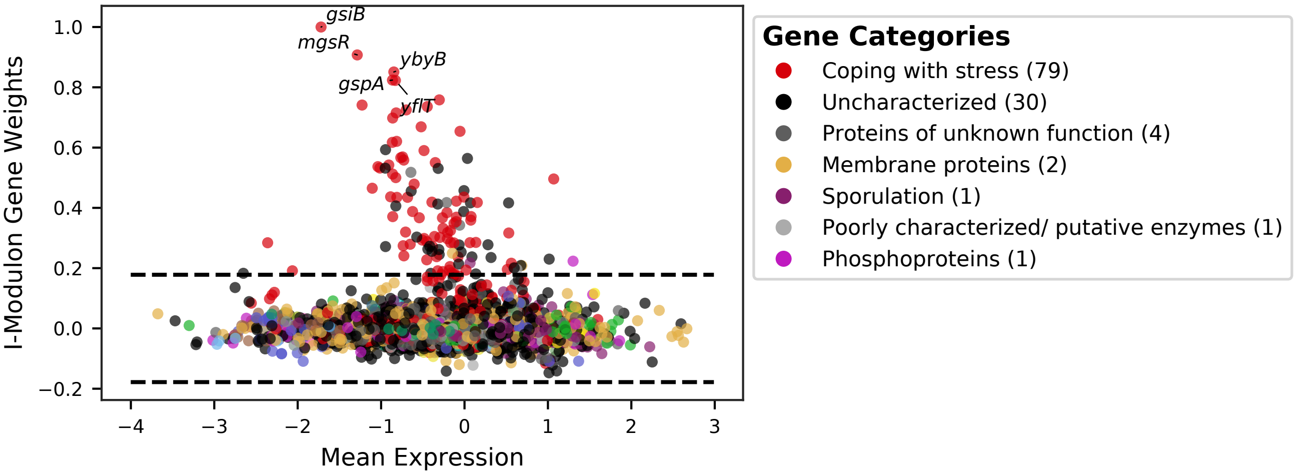


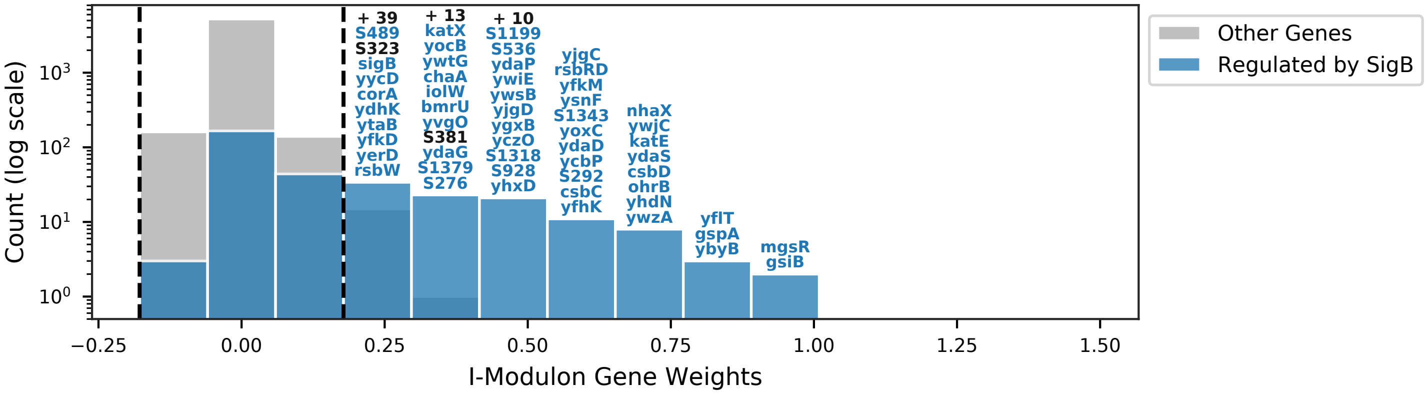


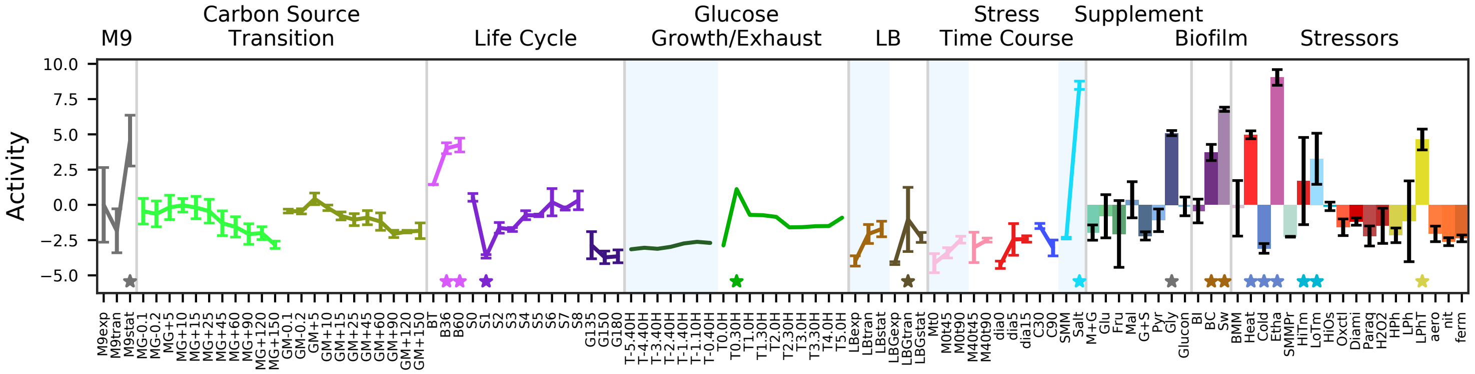


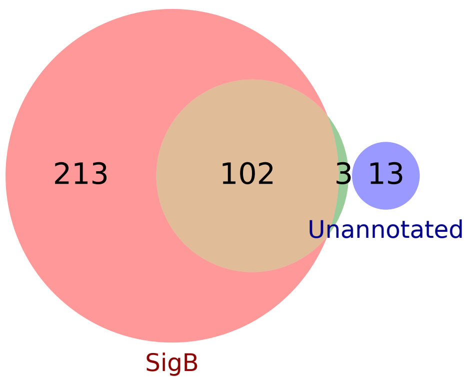


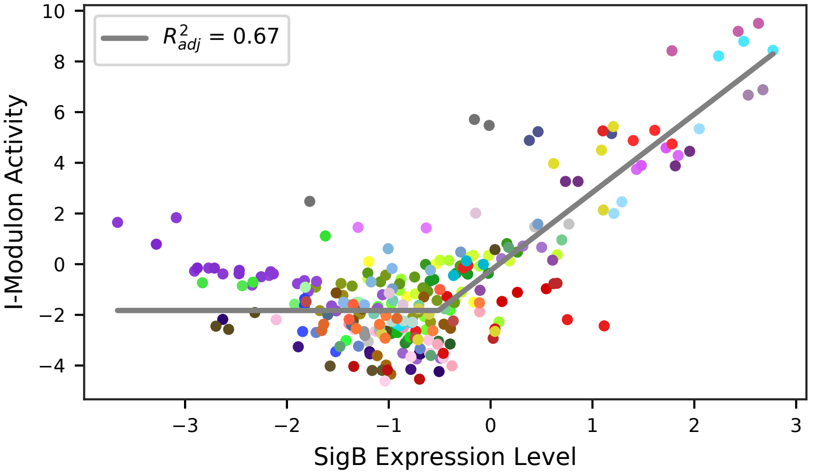

### 38 - SigB-2 – General Stress 2

Biological Function:
Likely accounting for microarray saturation in the ethanol condition of I-modulon SigB 1 by negatively weighting highly expressed ethanol genes

Enriched for known regulon:
SigB

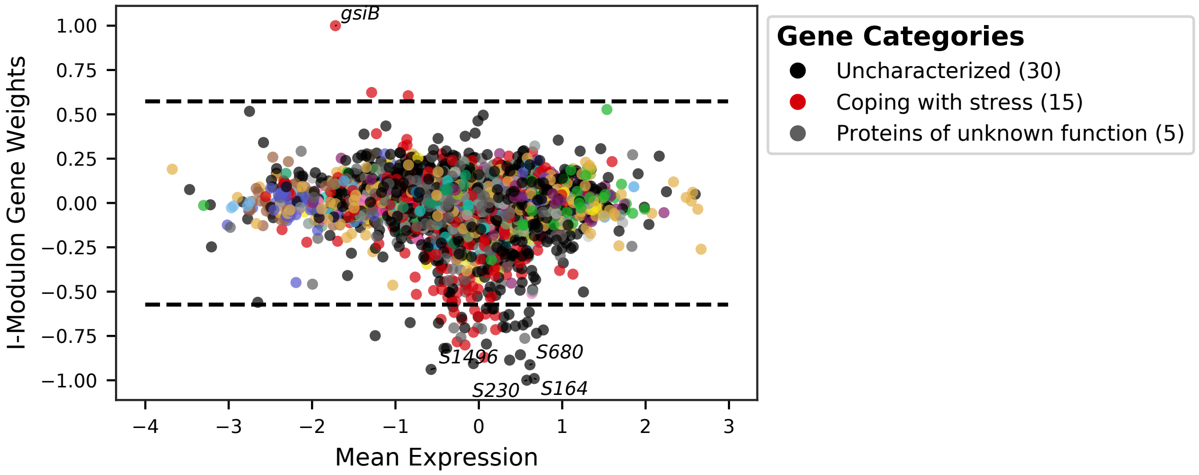


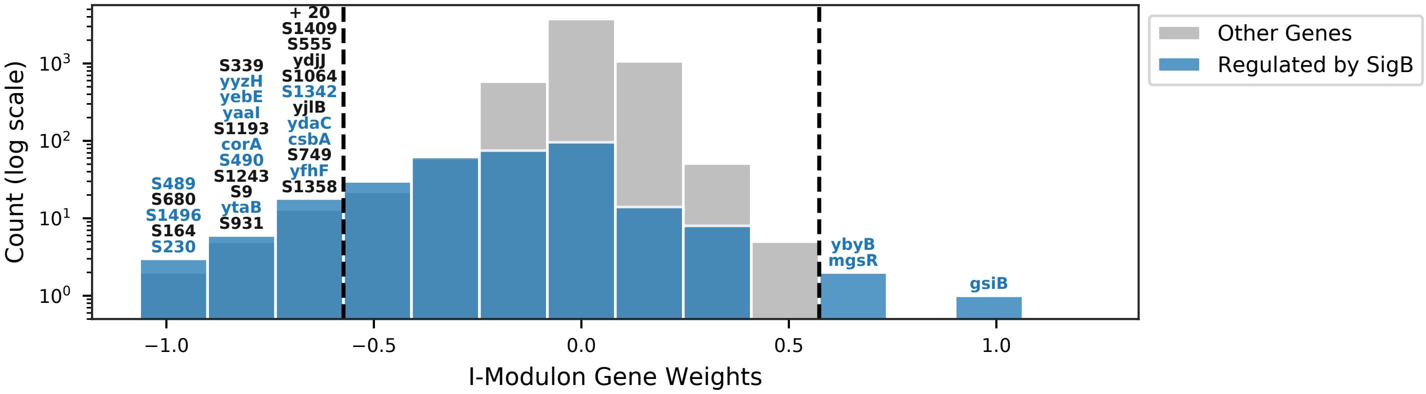


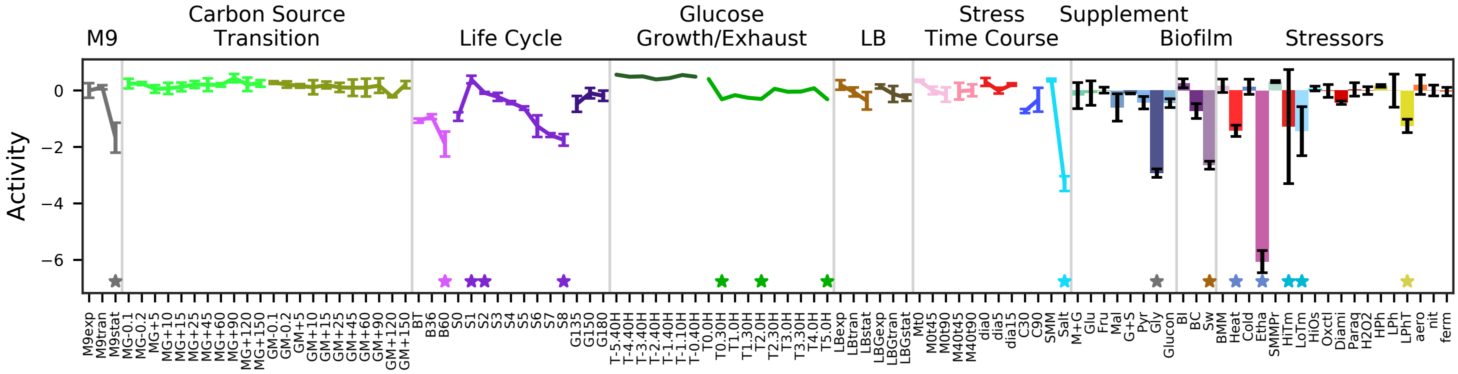


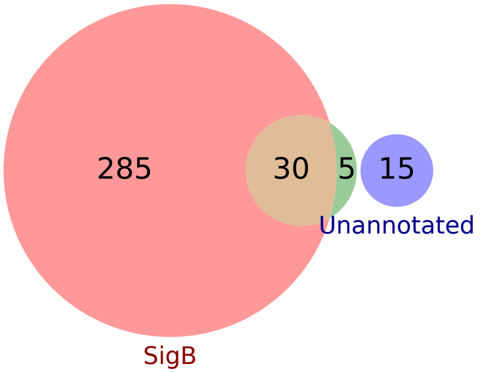


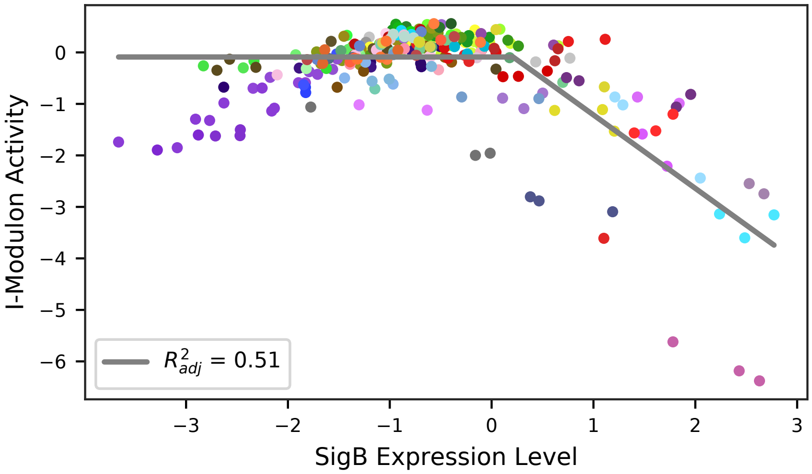

### 39 - StrRes – Translation

Biological Function:
Production of ribosomes, translation initiation factors, ADP formation, protein processing and secretory machinery, DNA replication, RNA polymerase

Subset of known regulon:
[stringent response] / [RplJ]

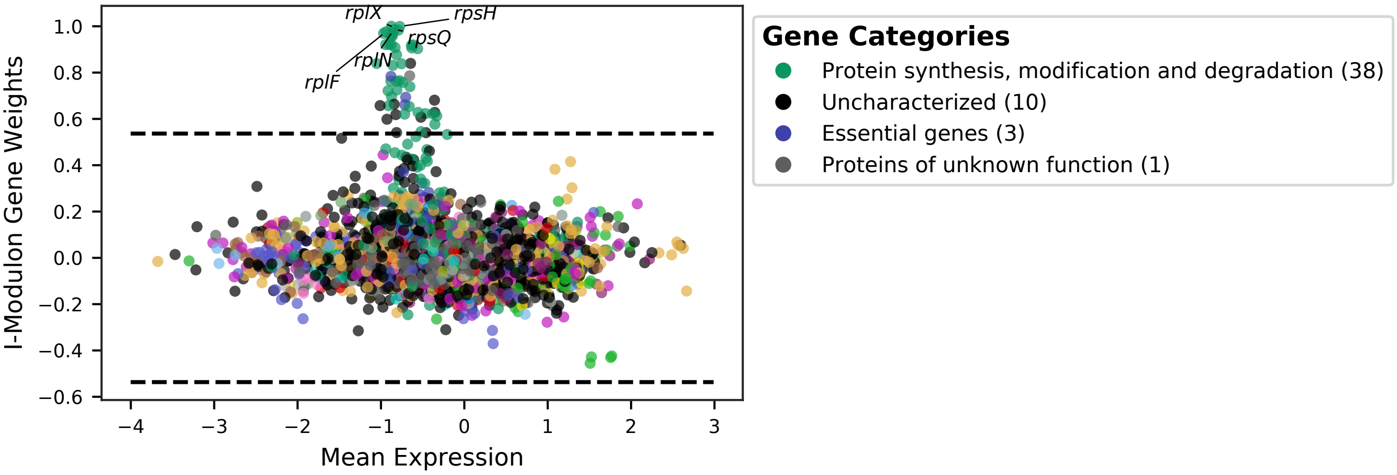


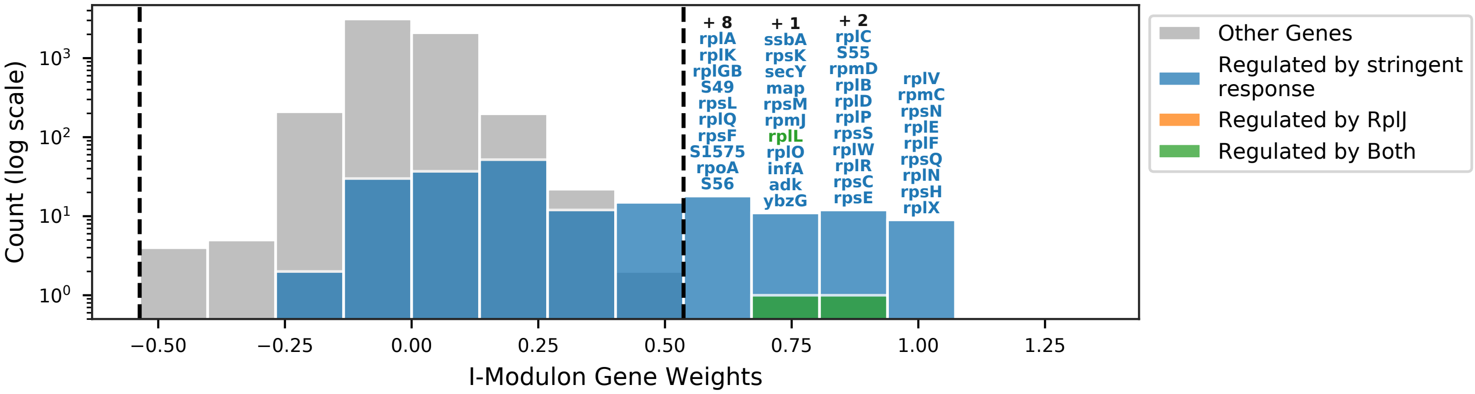


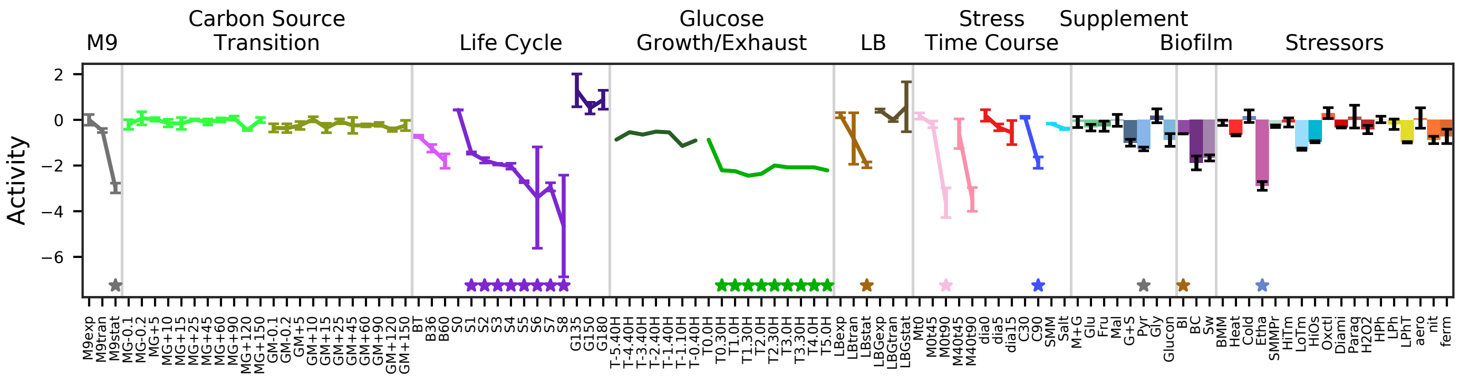


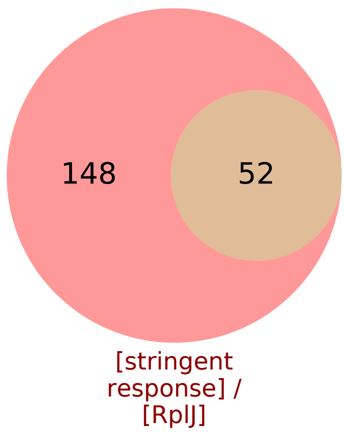


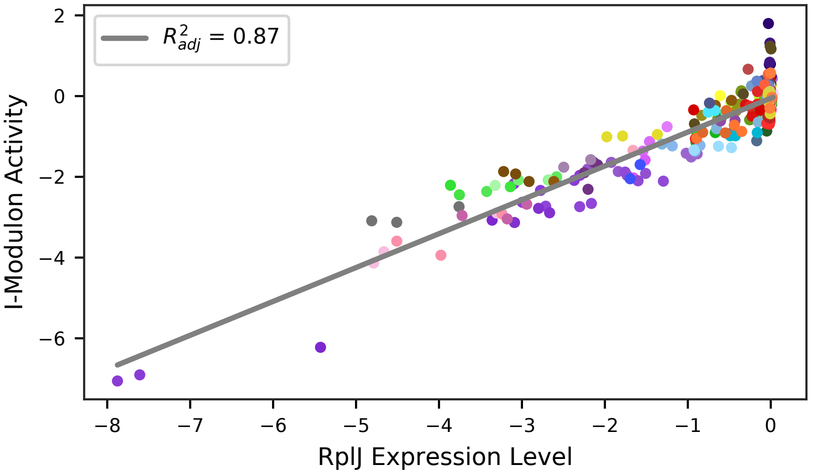

### 40 - SigD – Motility

Biological Function:
Production of flagella components and export machinery, chemotaxis receptors, use of extracellular energy (polyglutamic acid), regulation of swarming, sporulation inhibitors, cell wall turnover

Subset of known regulon:
SigD

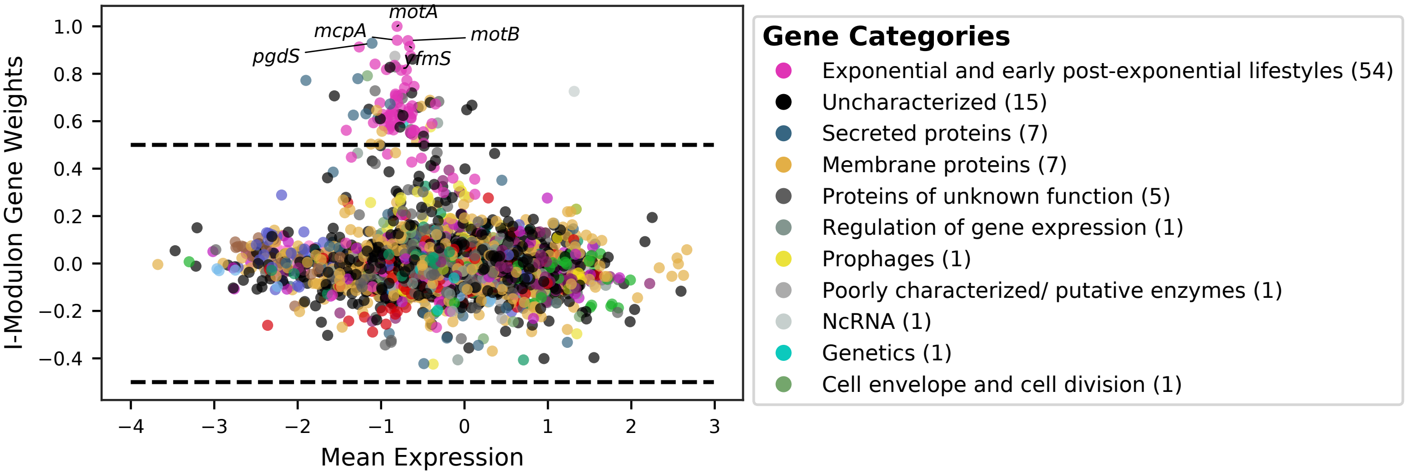


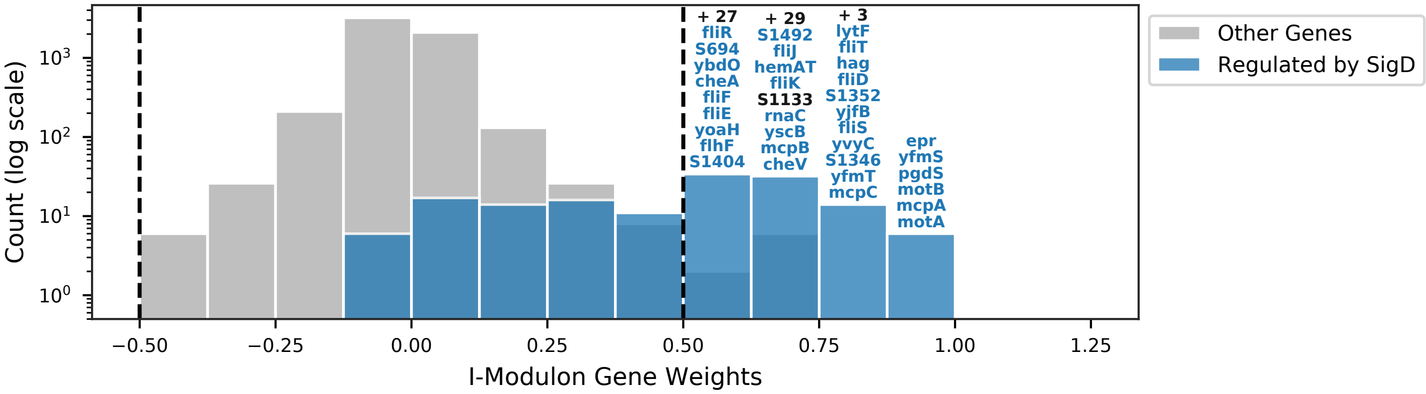


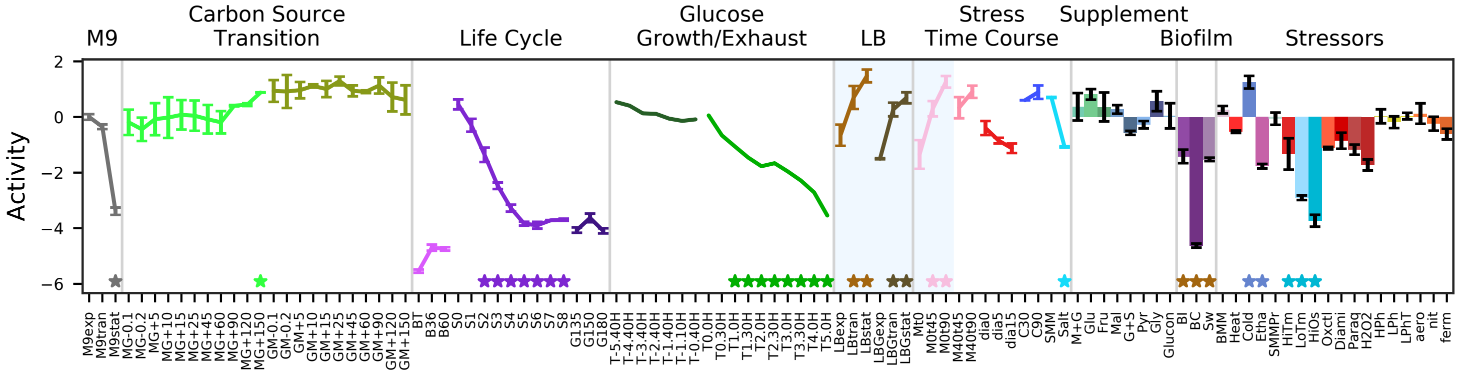


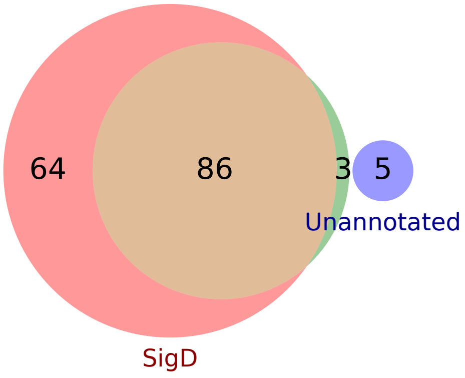


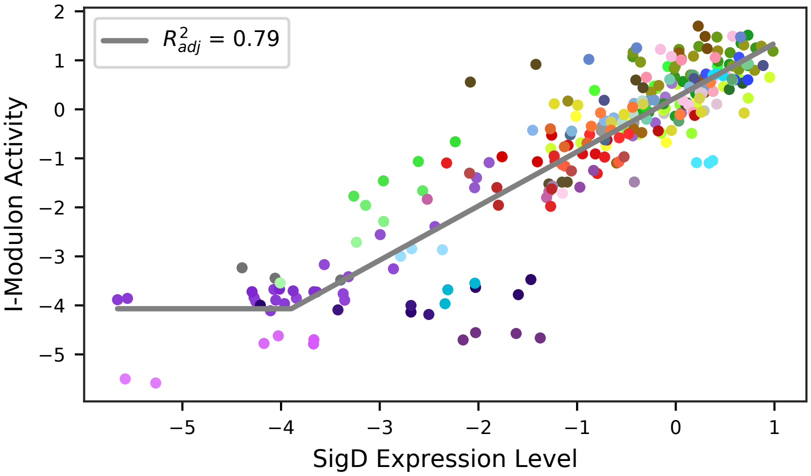

### 41 - Eps – Exopolymeric Substances

Biological Function:
Biosynthesis of tasA and extracellular polysaccharides, the major components of the biofilm

Well-defined regulon:
SigA + AbrB + RemA + SinR

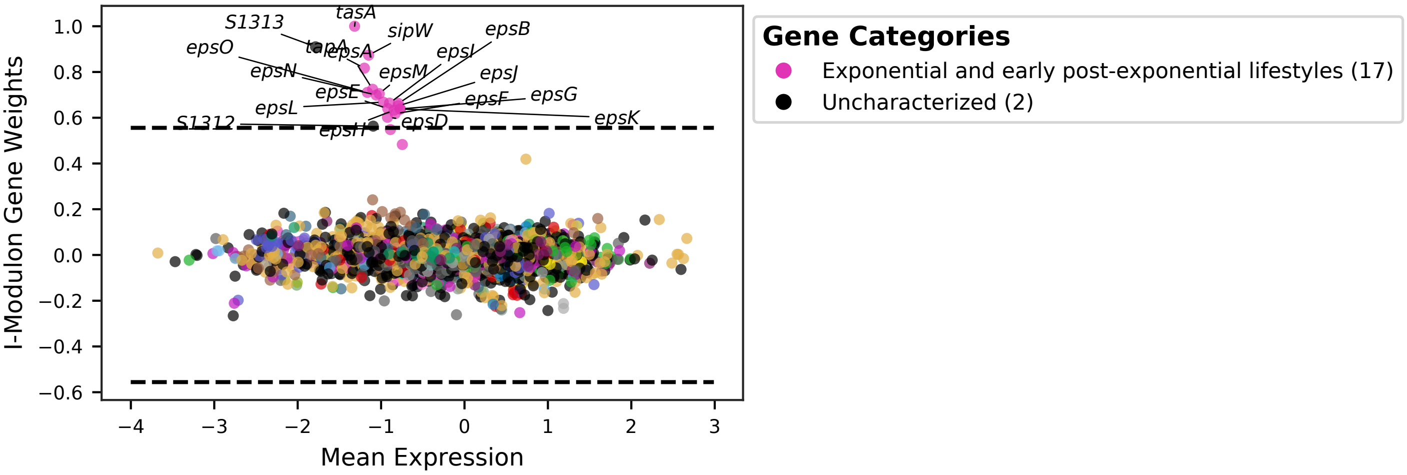


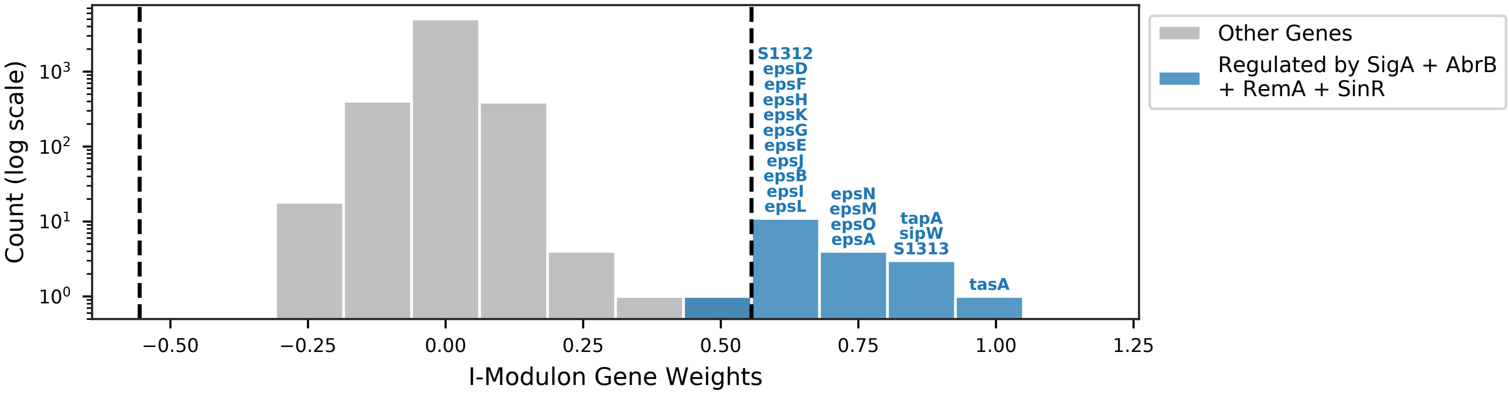


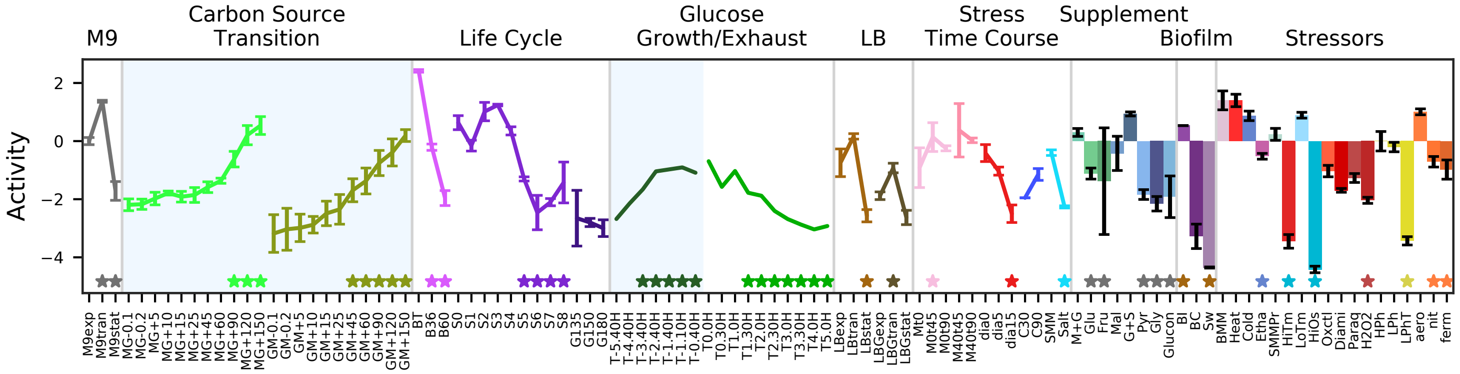


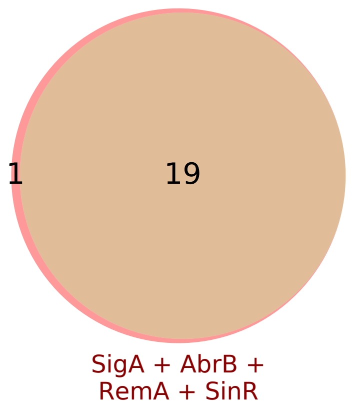


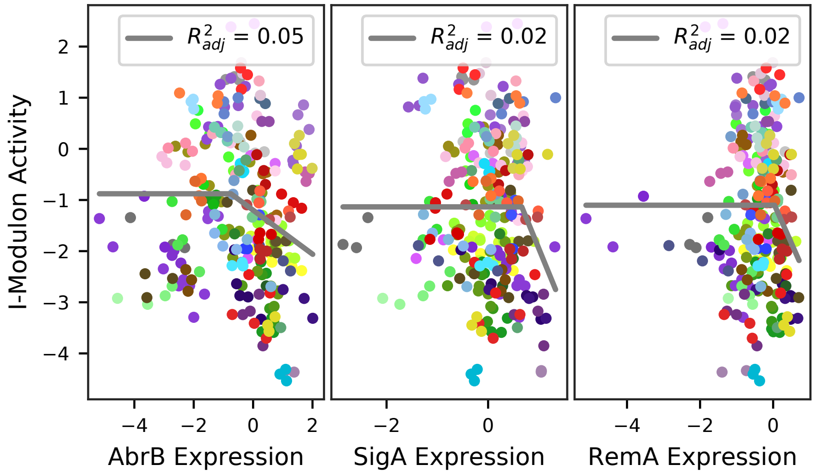

### 42 - ComA – Surfactants

Biological Function:
Activated during transition to stationary phase (quorum sensing), production of surfactant and exoenzymes to improve nutrient delivery in the biofilm, ComS for activation of competence

Well-defined regulon:
[ComA + CcpA + TnrA] / [Abh + SigA + ComA + Spx + PhoP + CodY + PerR]

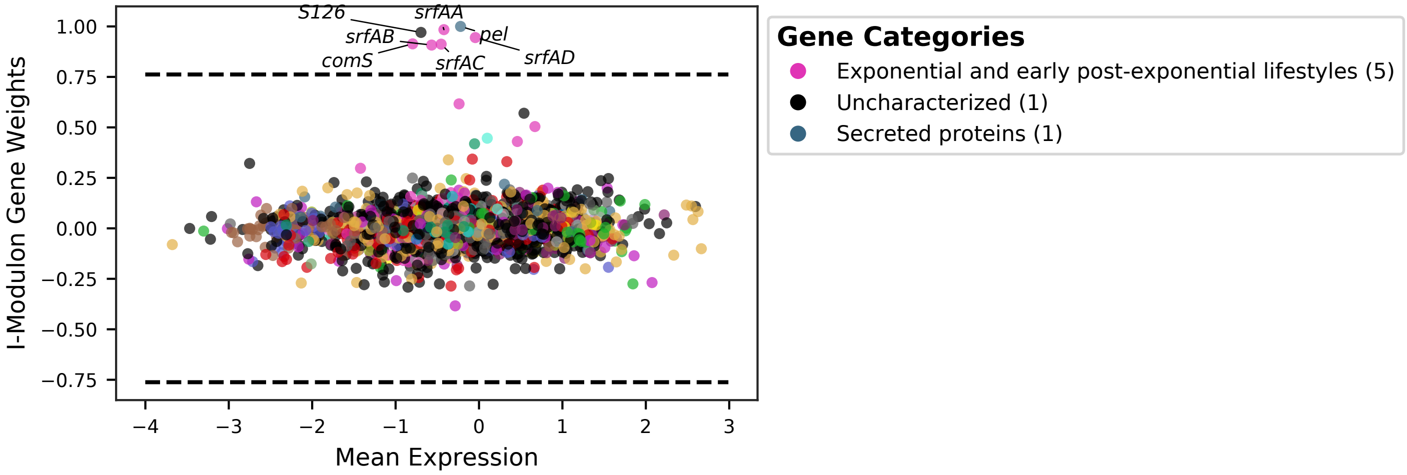


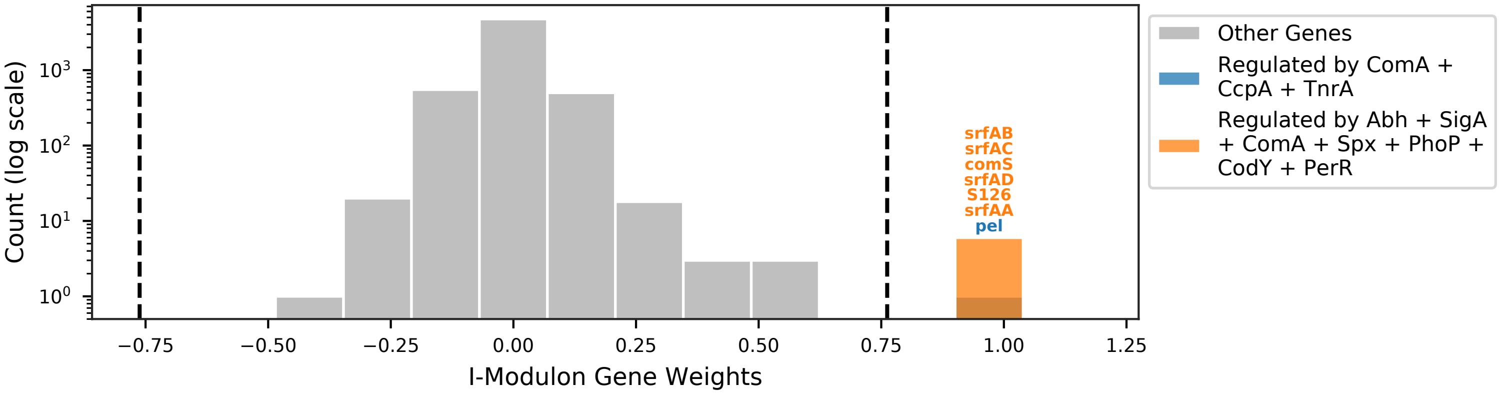


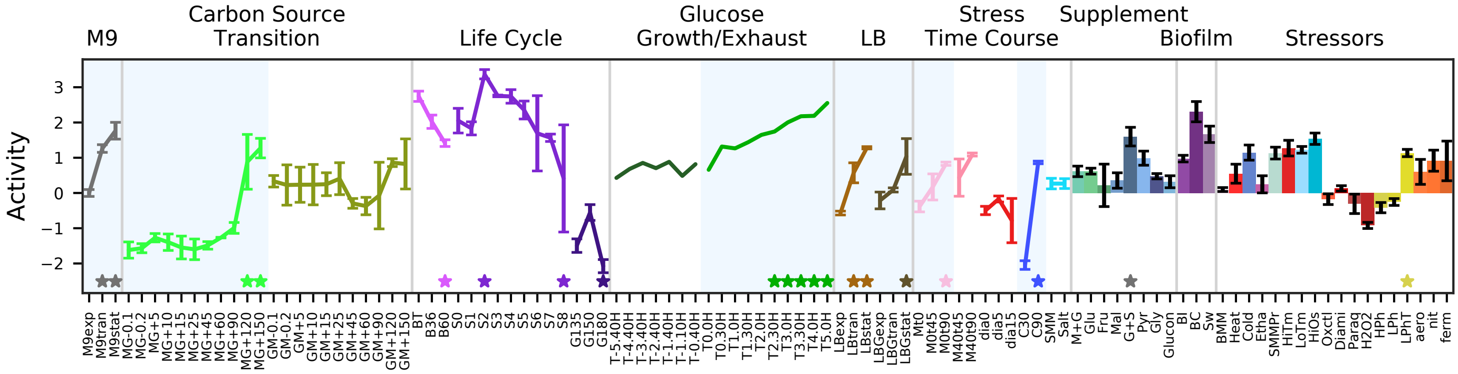


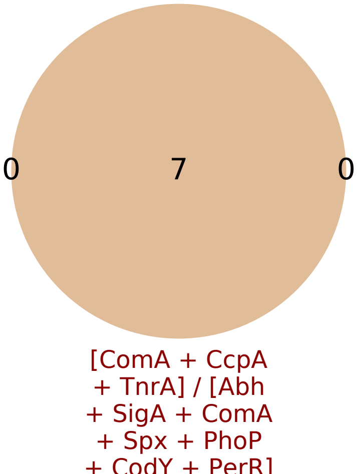


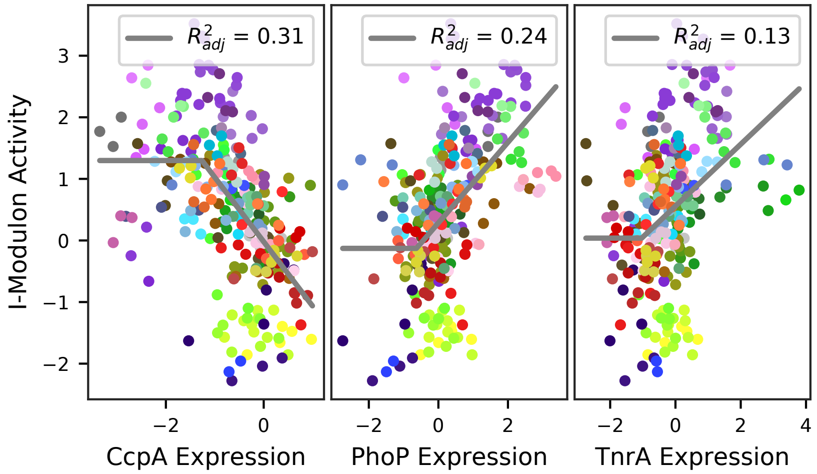

### 43 - DegU – Toxins and Colony Establishment

Biological Function:
Response to arrested flagellar motor. Secretion of yukE, a homolog of virulence factor EsxA. Production of antitoxins and putative toxin y-genes. Production of hydrophobic biofilm layer and lipoproteins for motility. Many unknown proteins.

Enriched for known regulon:
DegU

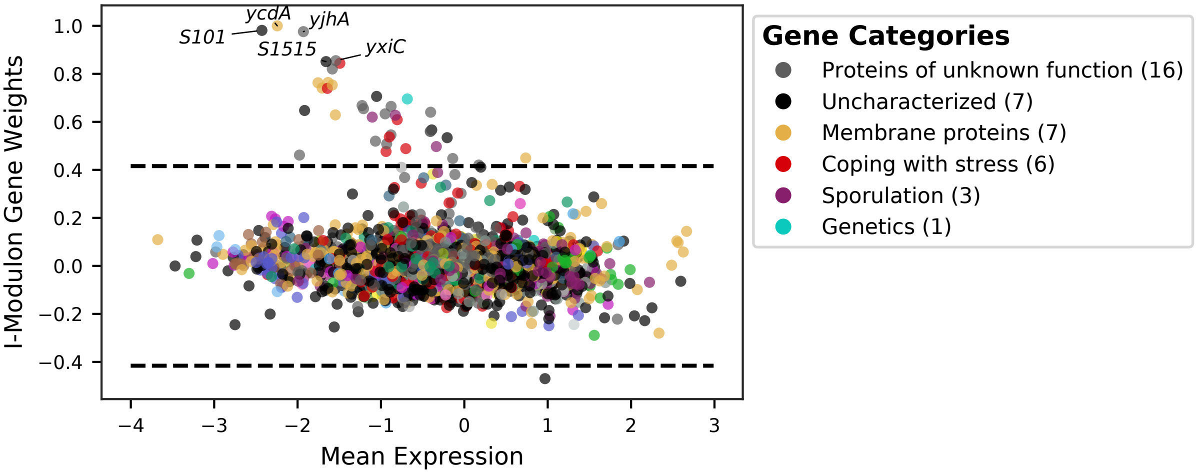


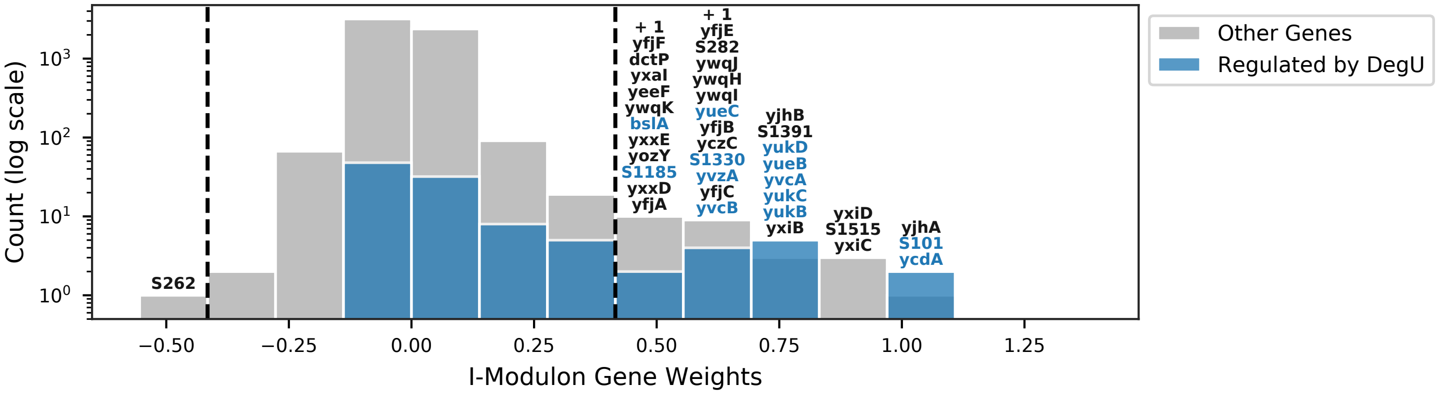


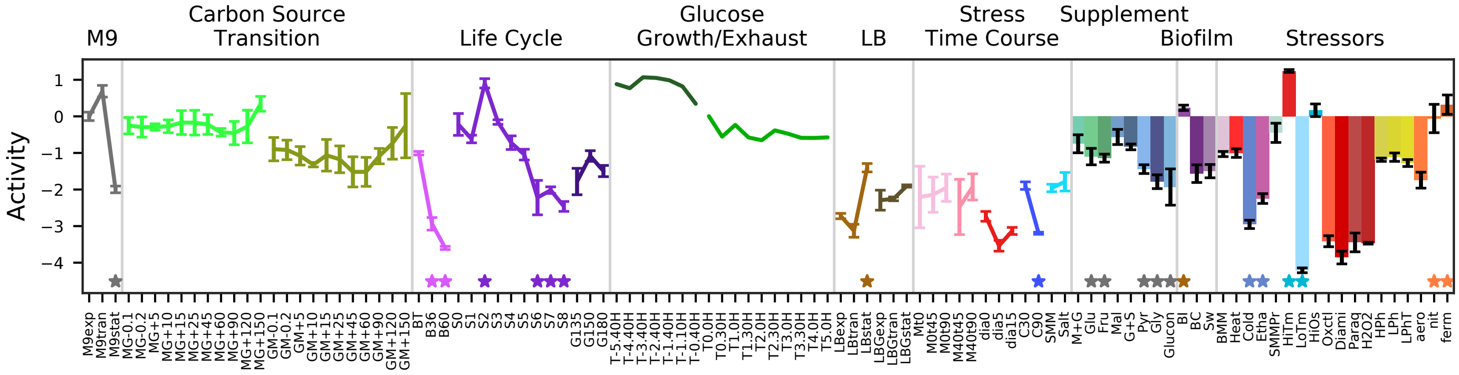


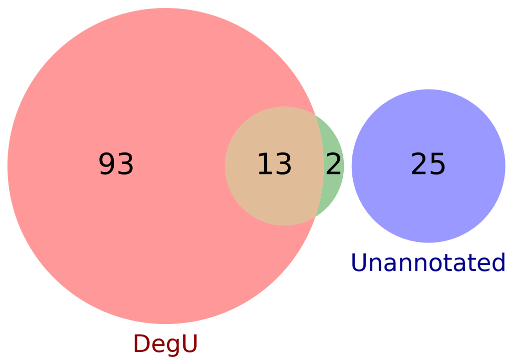


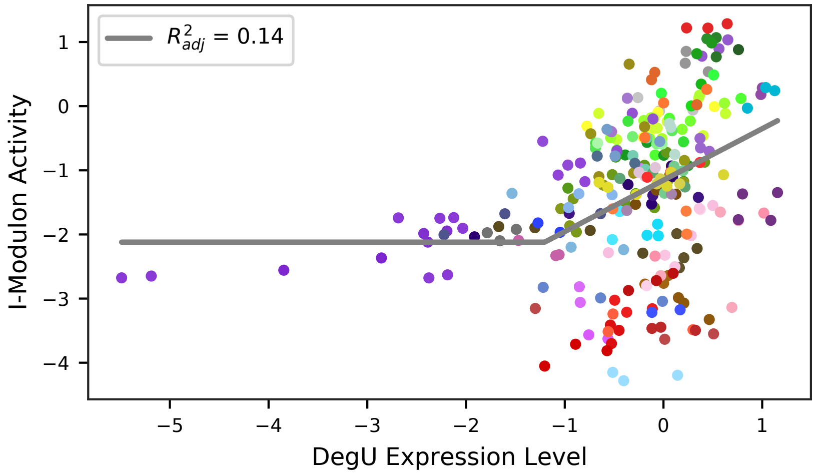

### 44 - AbrB – Transition State

Biological Function:
Various transition state roles: spore killing, antibiotic resistance, toxin production, metabolism, protein modification, and unknown proteins.

Subset of known regulon:
AbrB

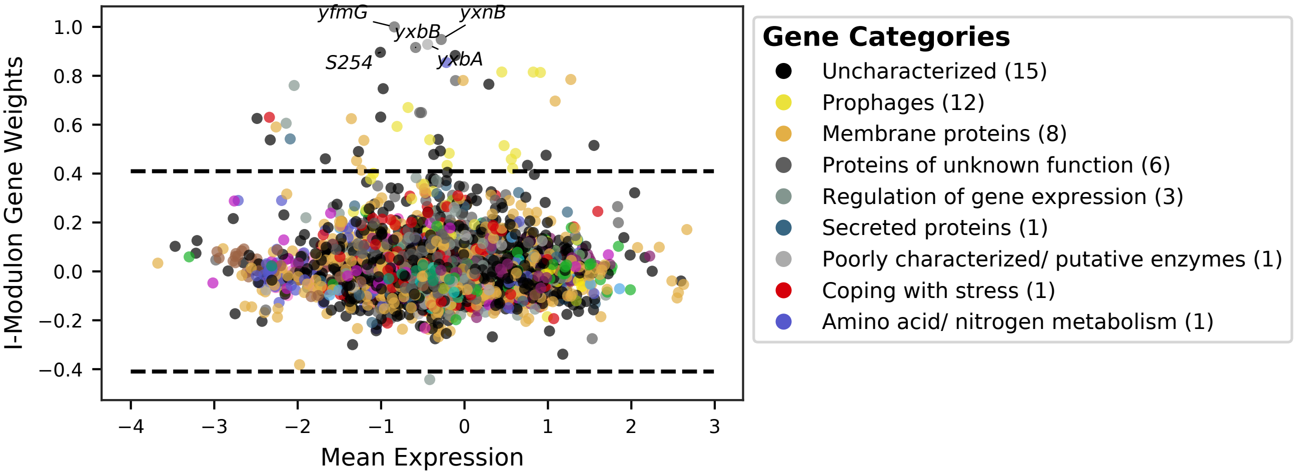


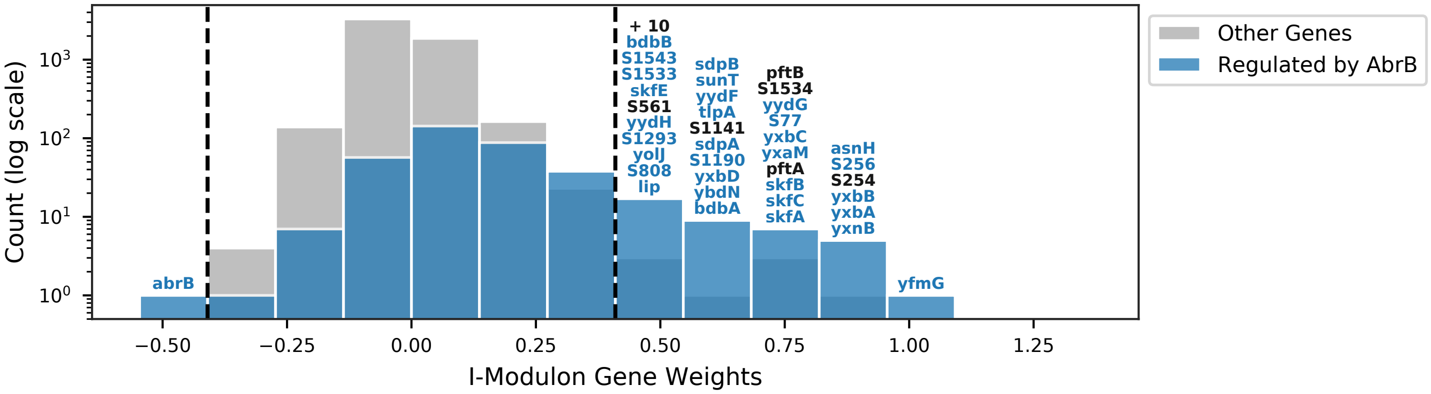


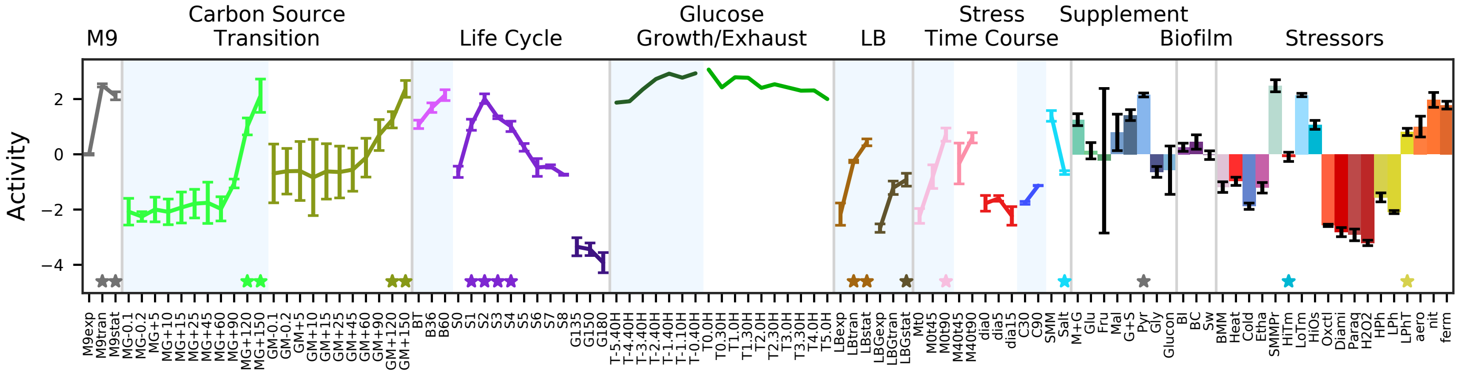


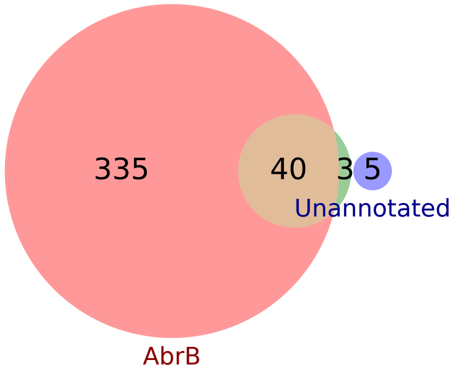


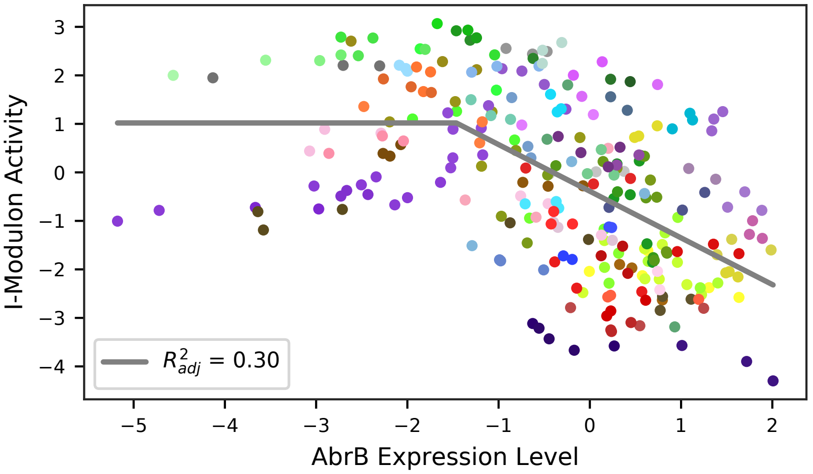

### 45 - Alb – Antilisterial Bacteriocin

Biological Function:
Produces and exports subtilosin, which kills anaerobic listeria bacteria

Well-defined regulon:
Rok + ResD + AbrB + SigA

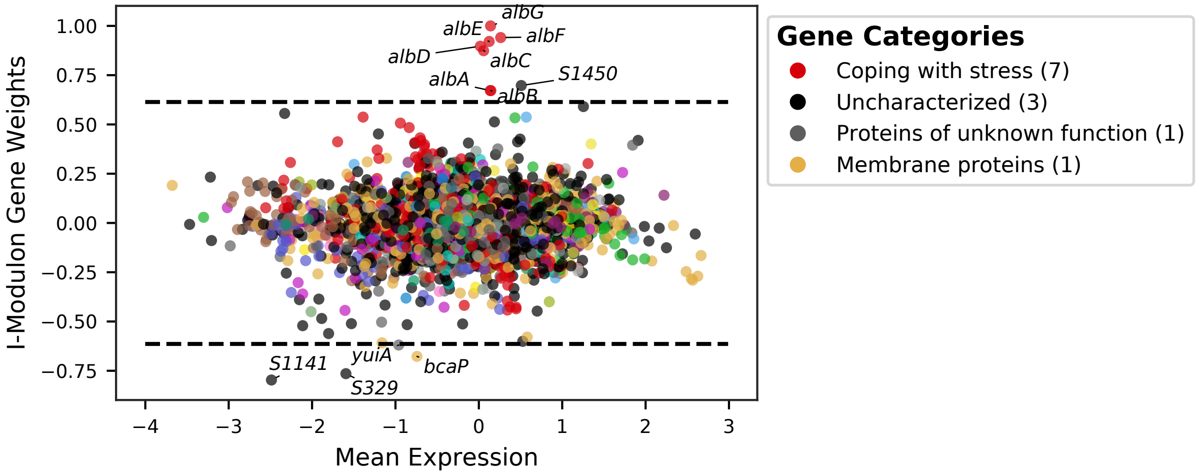


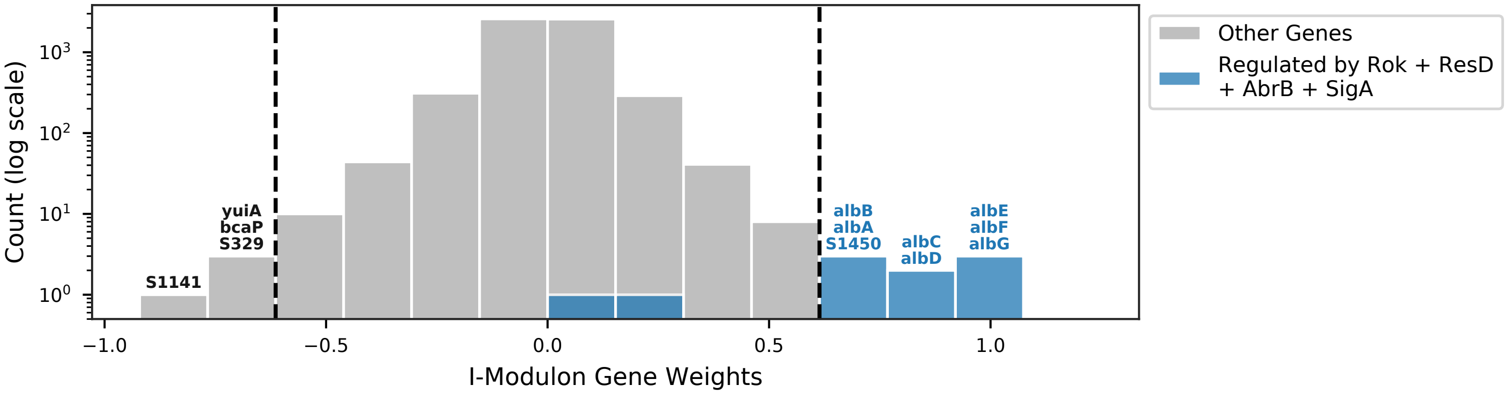


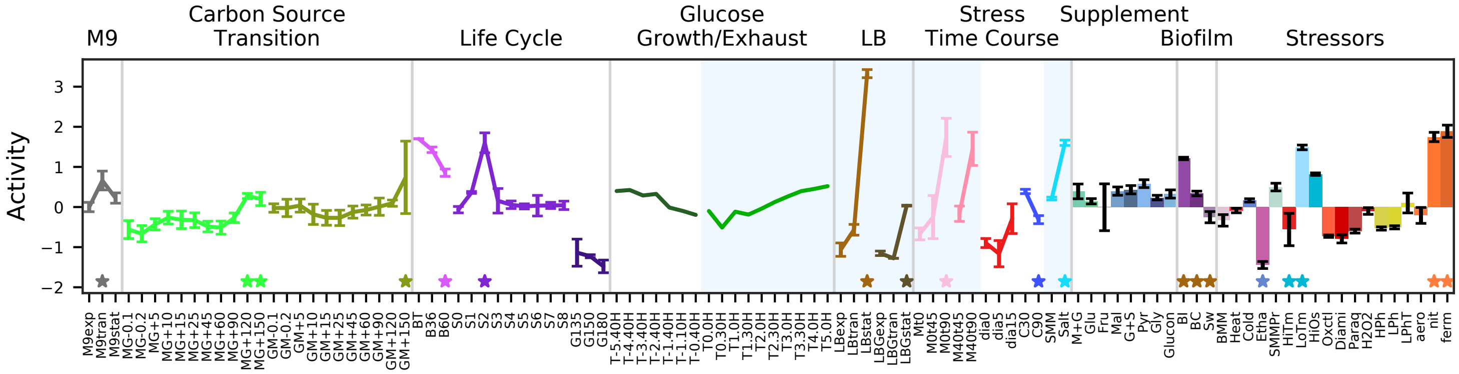


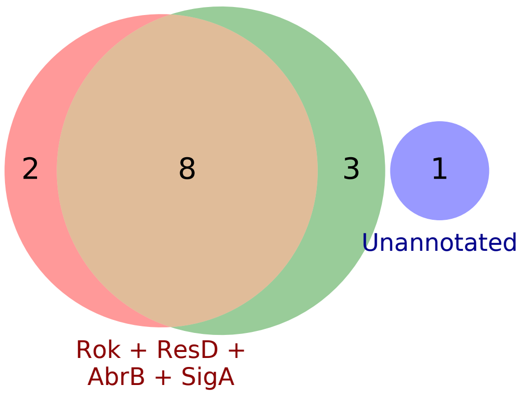


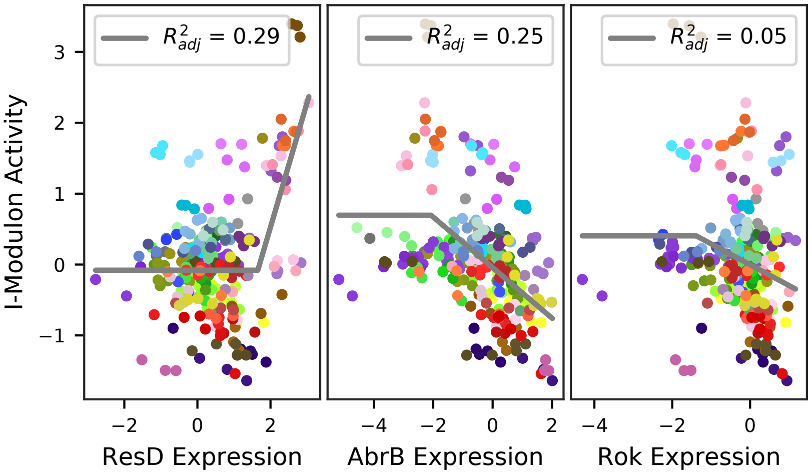

### 46 - WapA – Intercellular Competition

Biological Function:
Activation of WapA and its operon of unknown genes for interspecies toxin delivery

Contains unknown genes and known regulon:
SigA + YvrHb + DegU + WalR

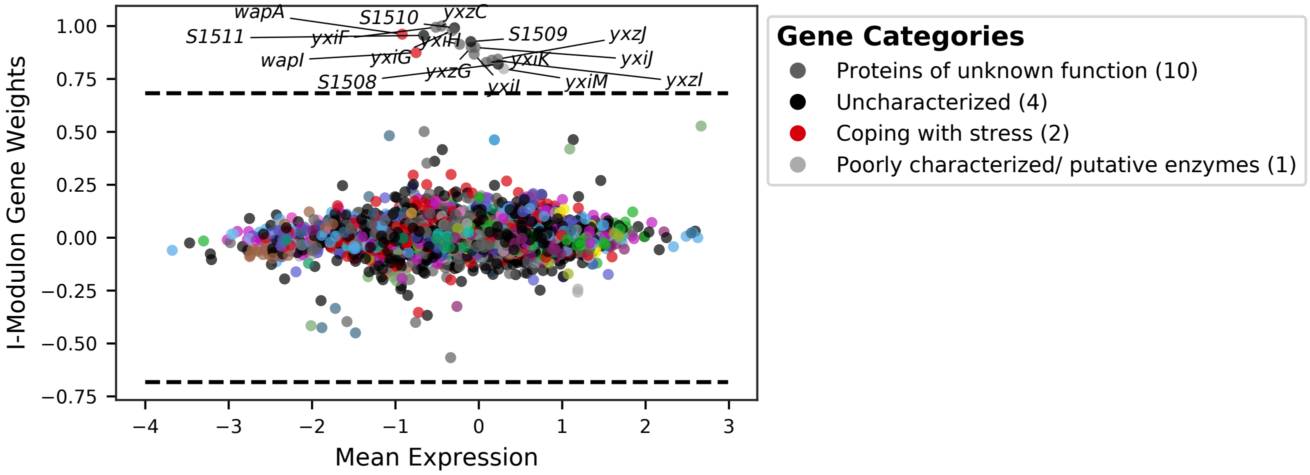


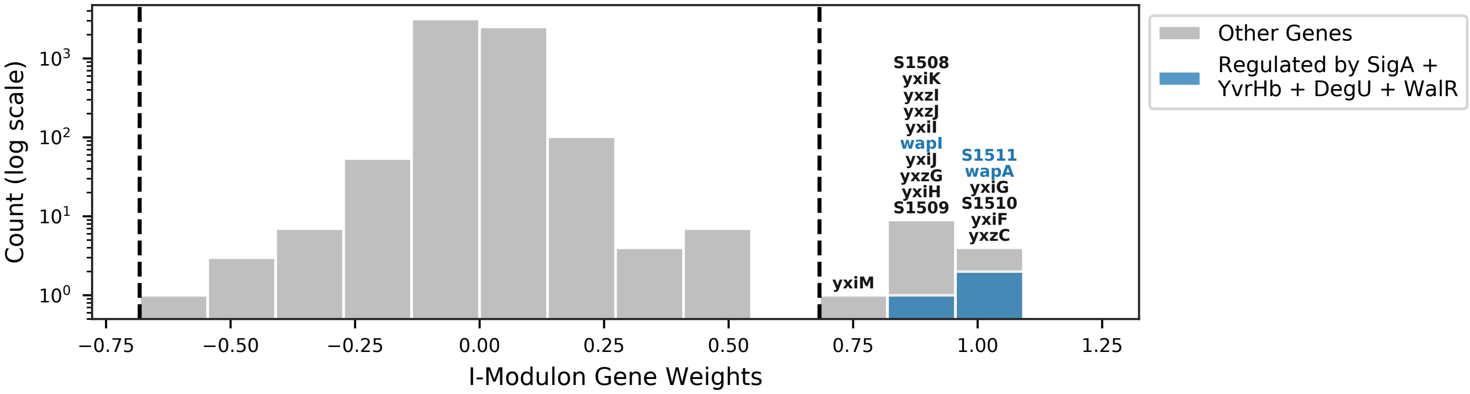


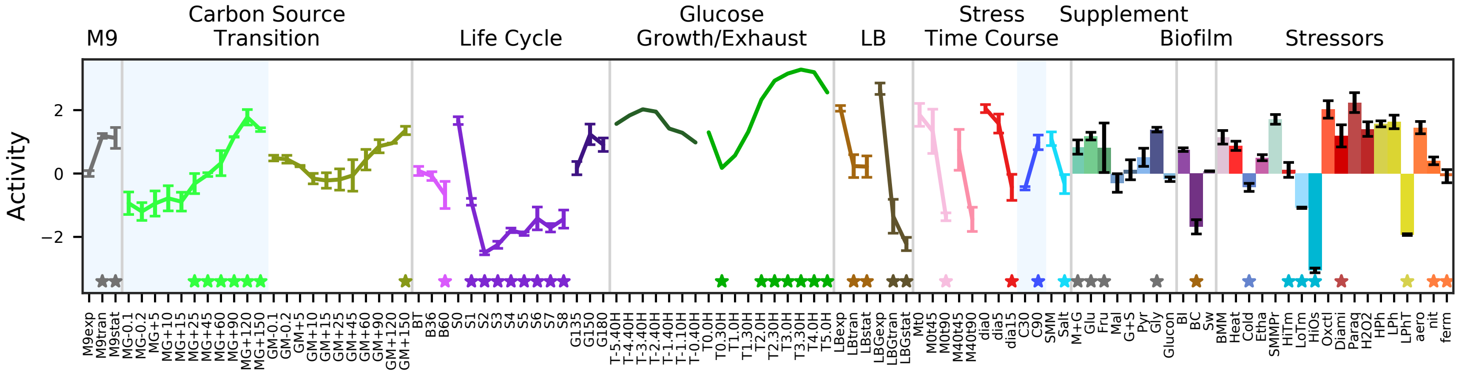


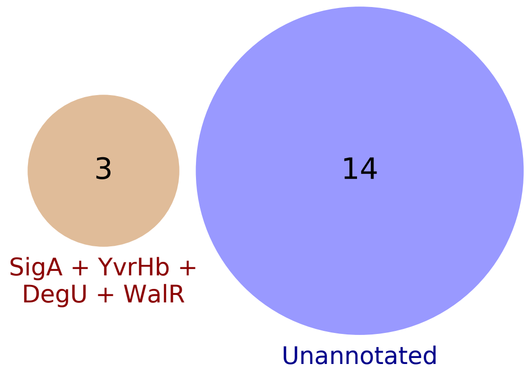


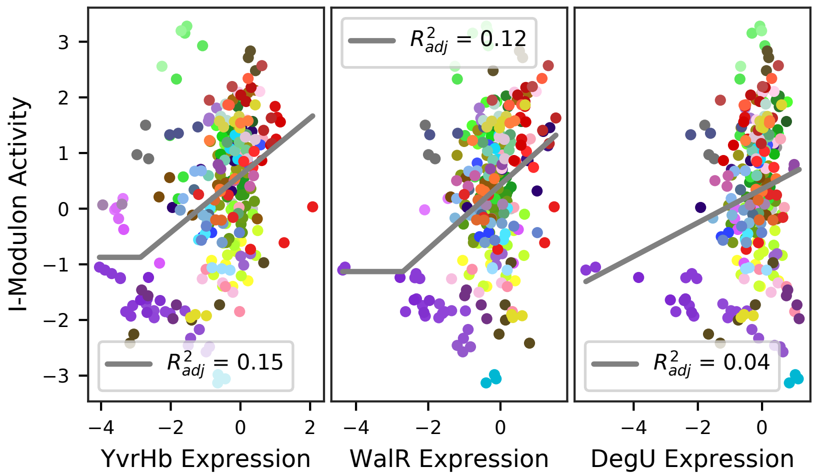

### 47 - ComK – Competence

Biological Function:
DNA transport, pseudopilus, recombination machinery, and regulatory proteins for competence

Enriched for known regulon:
ComK

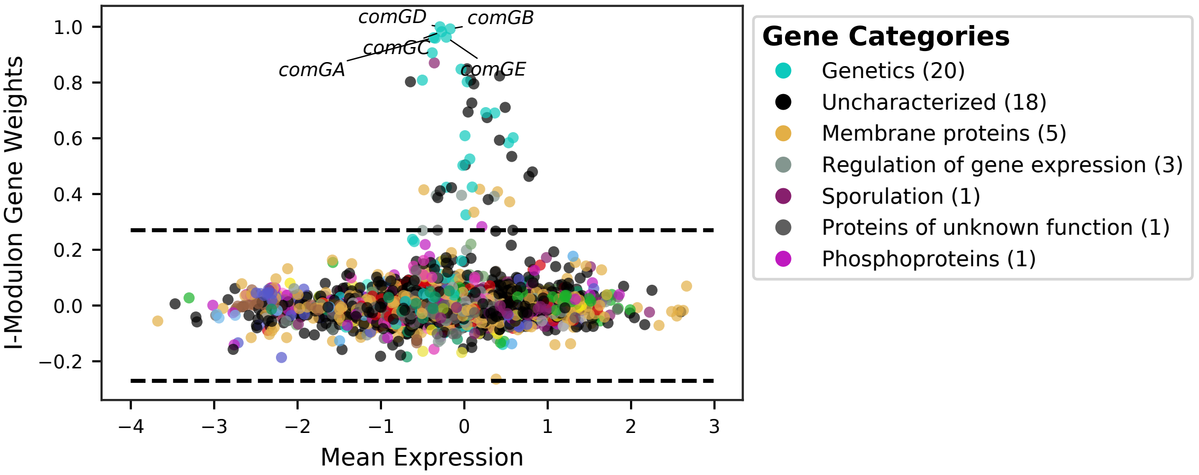


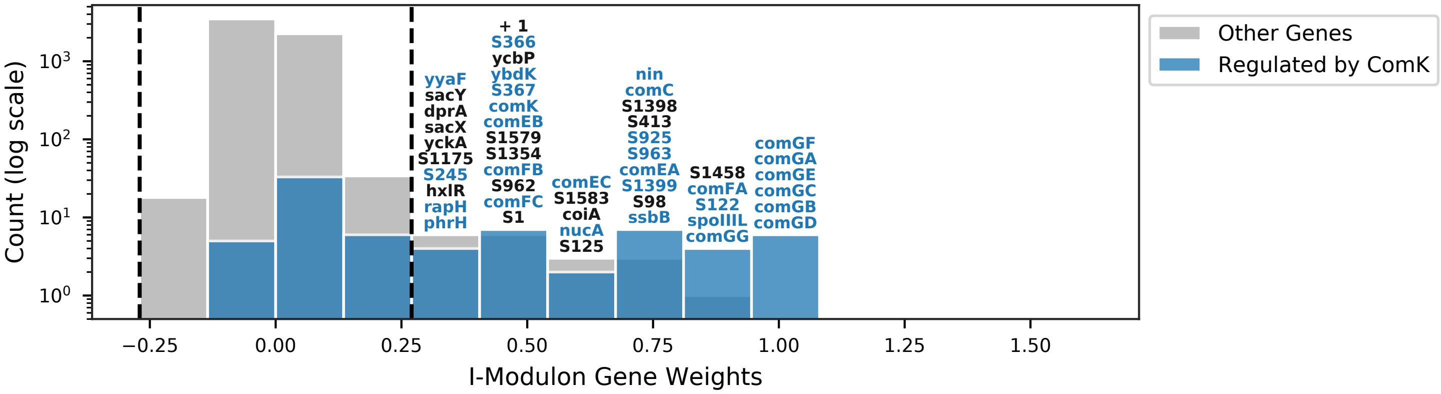


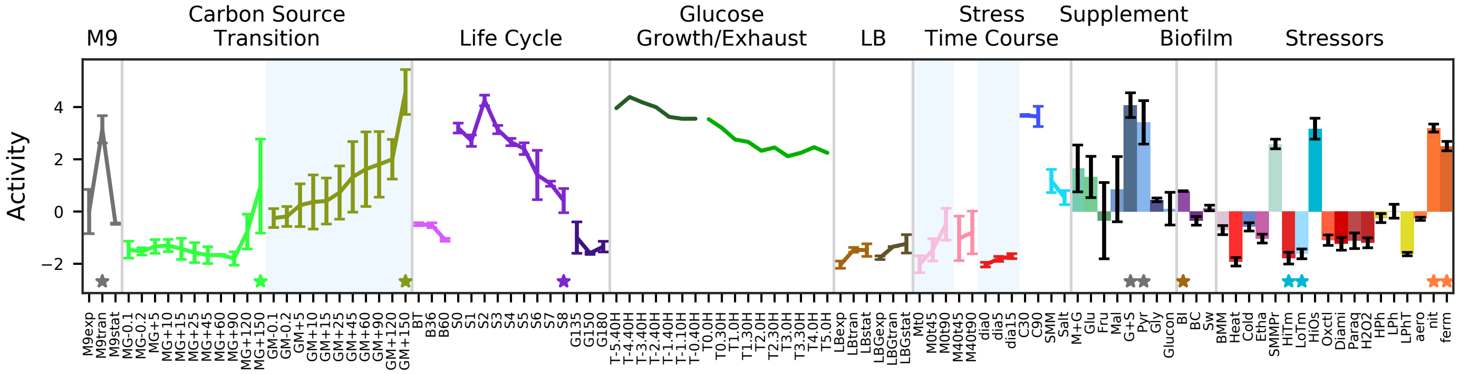


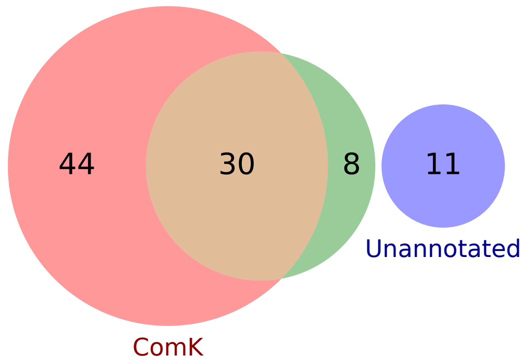


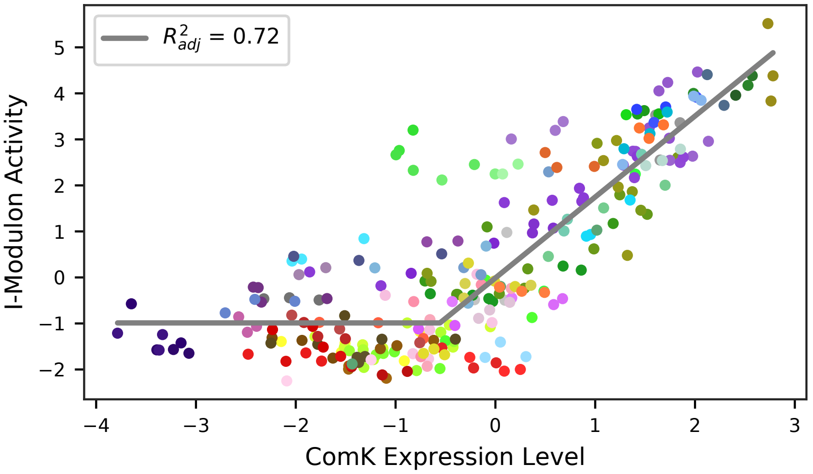

### 48 - Spo0A – Sporulation 1

Biological Function:
Initiation of sporulation through sigma factor, transcription factor, and regulatory protein expression, as well as septum formation and chromosome anchoring

Subset of known regulon:
[Spo0A] / [DnaA]

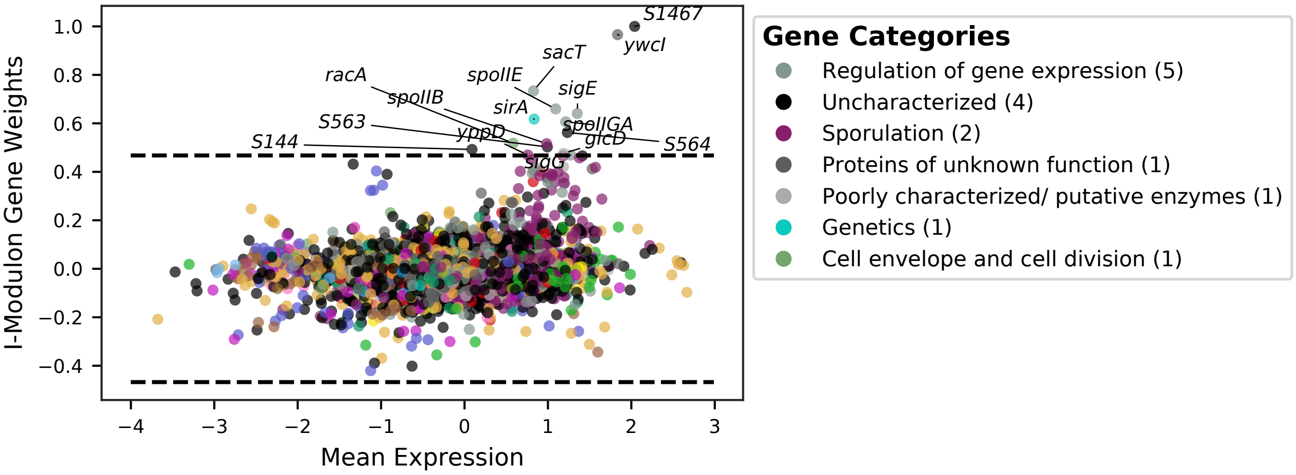


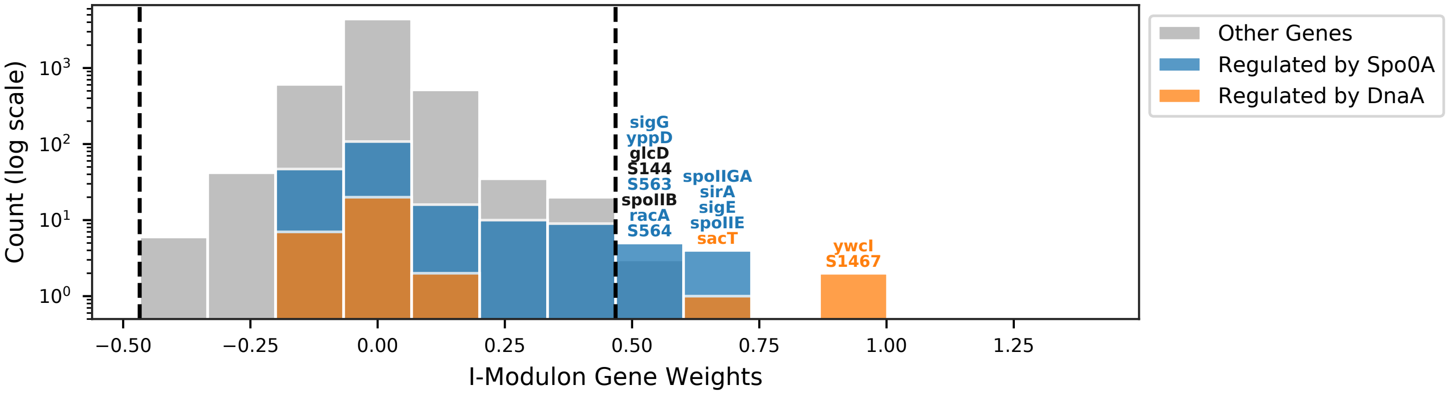


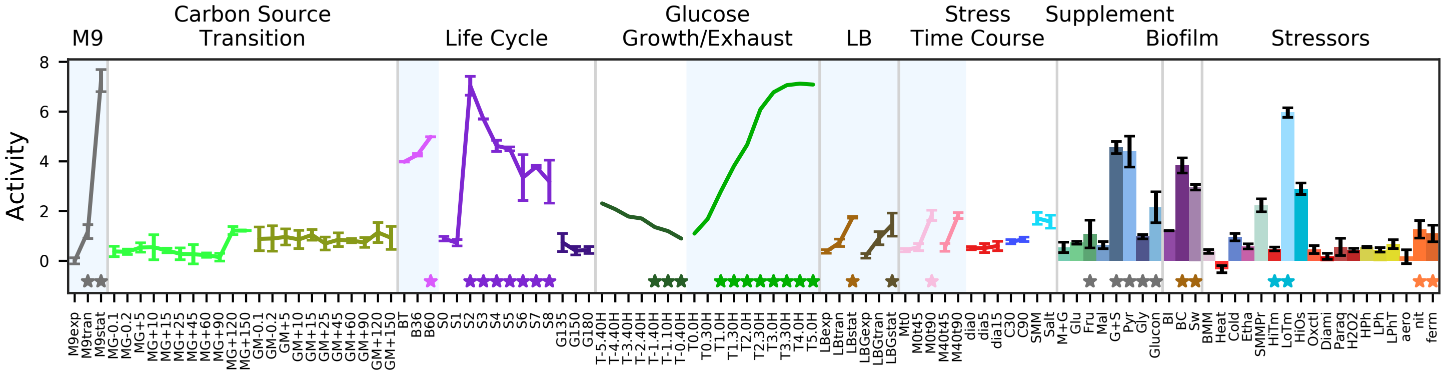


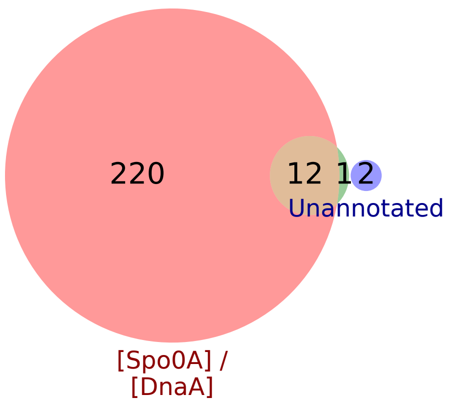


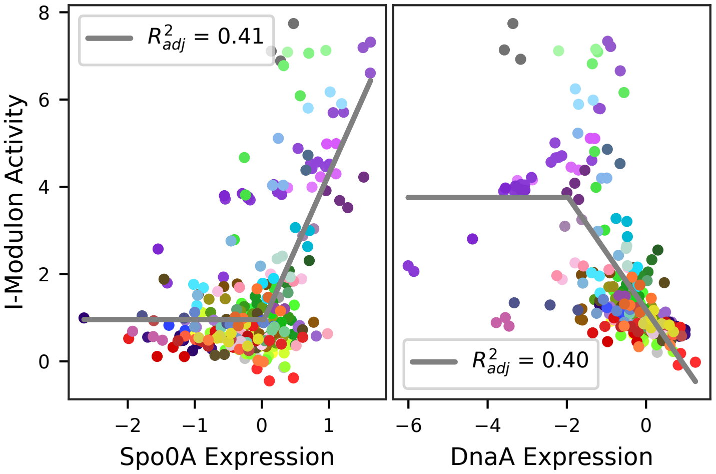

### 49 - SigE – Sporulation 2

Biological Function:
Spore maturation through core dehydration, nutrient transport and utilization, septum degradation, inner cortex synthesis, and regulator activation

Enriched for known regulon:
SigE

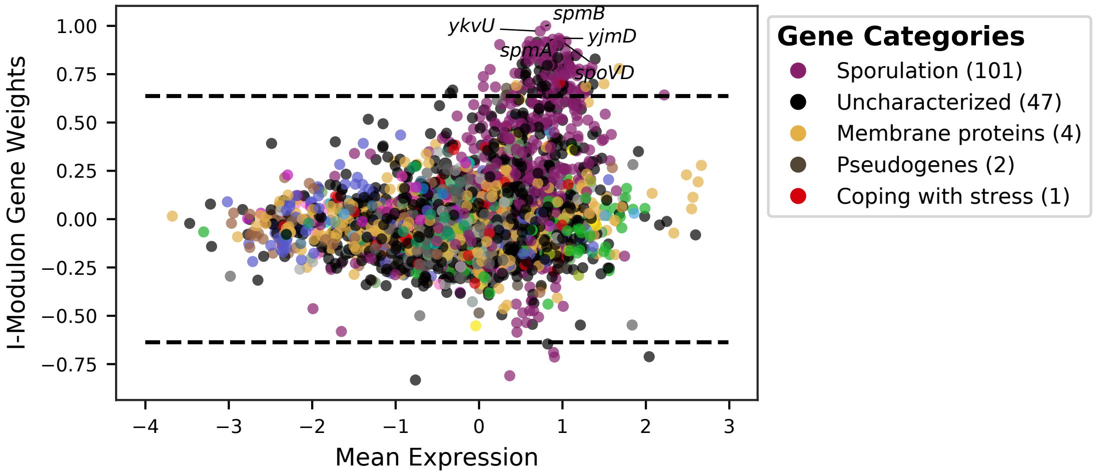


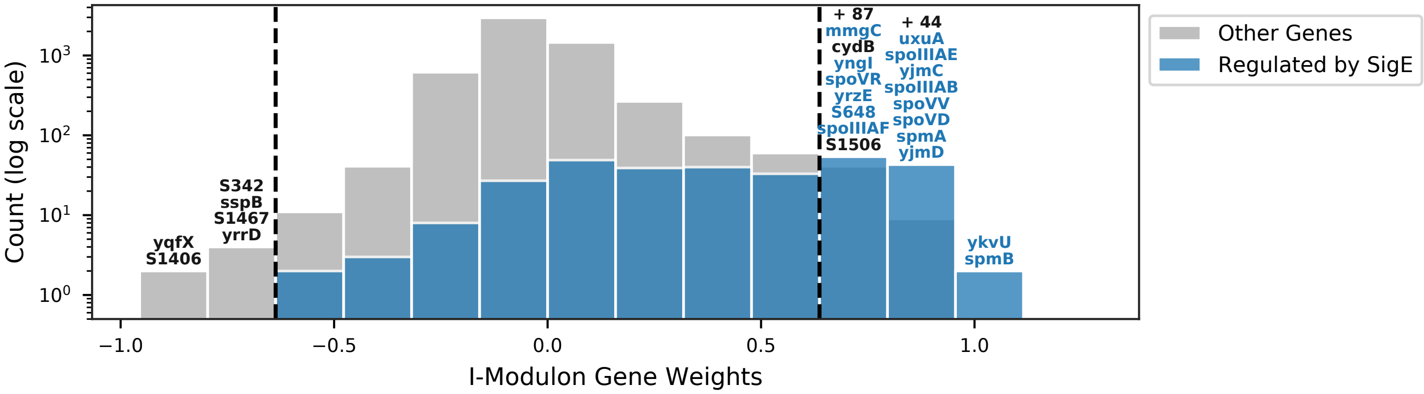


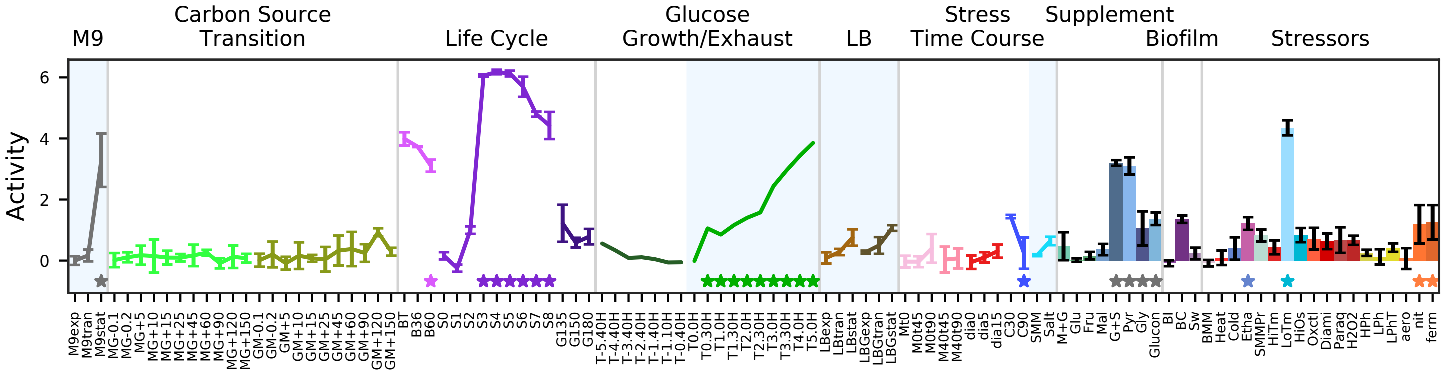


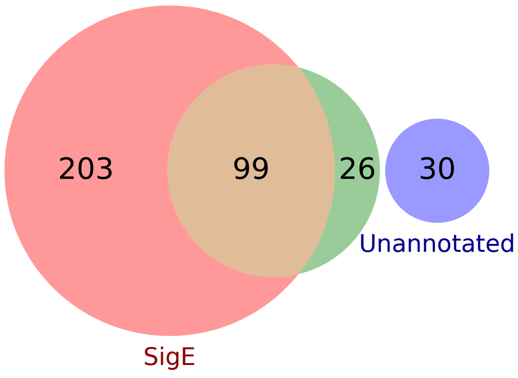


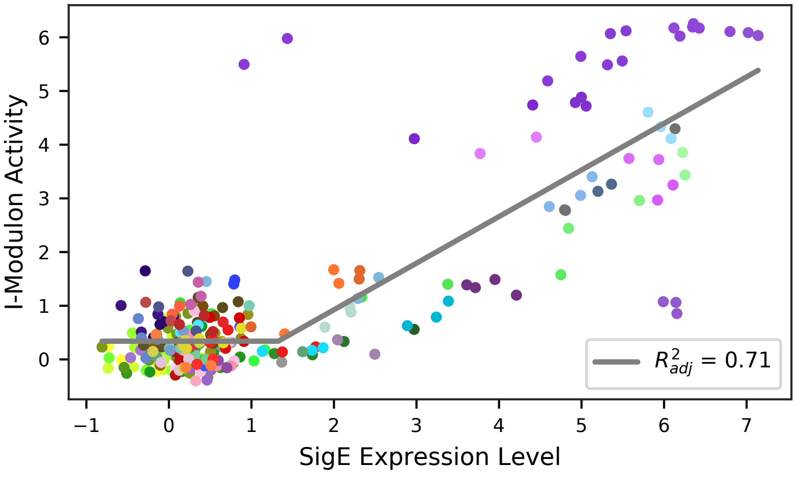

### 50 - SigE / G – Sporulation 3

Biological Function:
Spore coat production, transcription factor and regulator expression, spore DNA protection, spore stress resistance

Subset of known regulon:
[SigE] / [SigG]

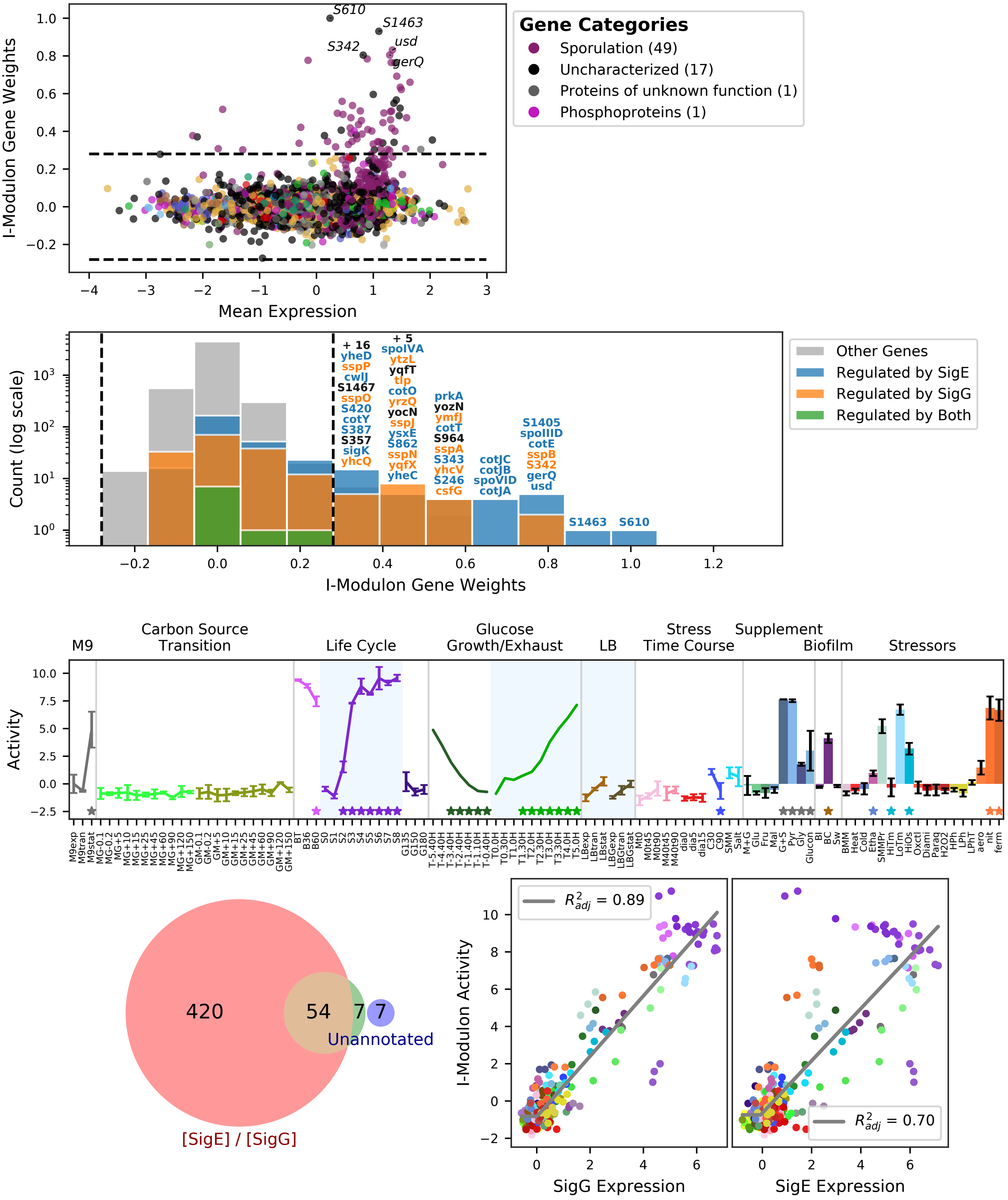

### 51 - SigG – Sporulation 4

Biological Function:
Calcium and nutrient delivery to the spore, endospore cortex maturation, lipoproteins, forespore-specific metabolic proteins, many unknown sporulation proteins, stress resistance, proteins required for germination

Enriched for known regulon:
SigG

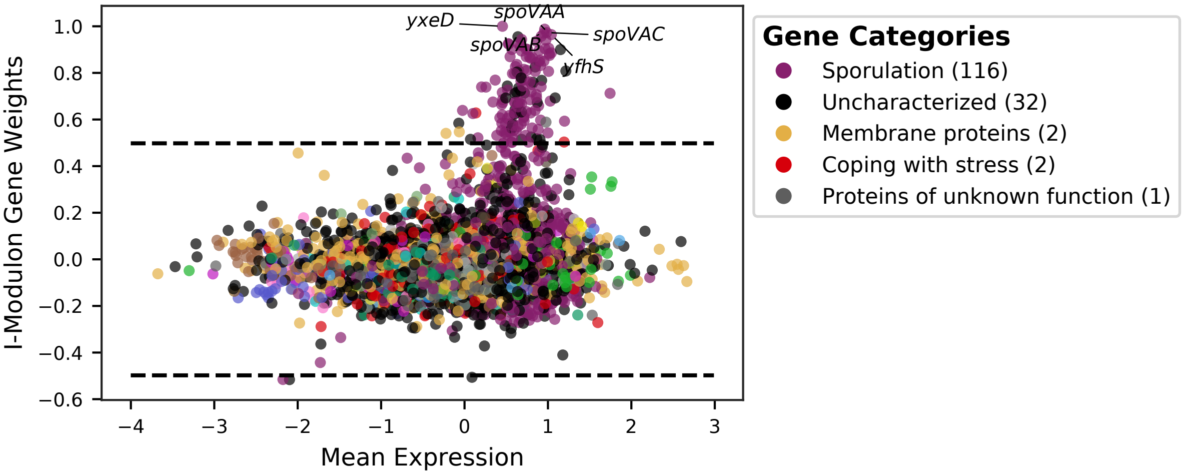


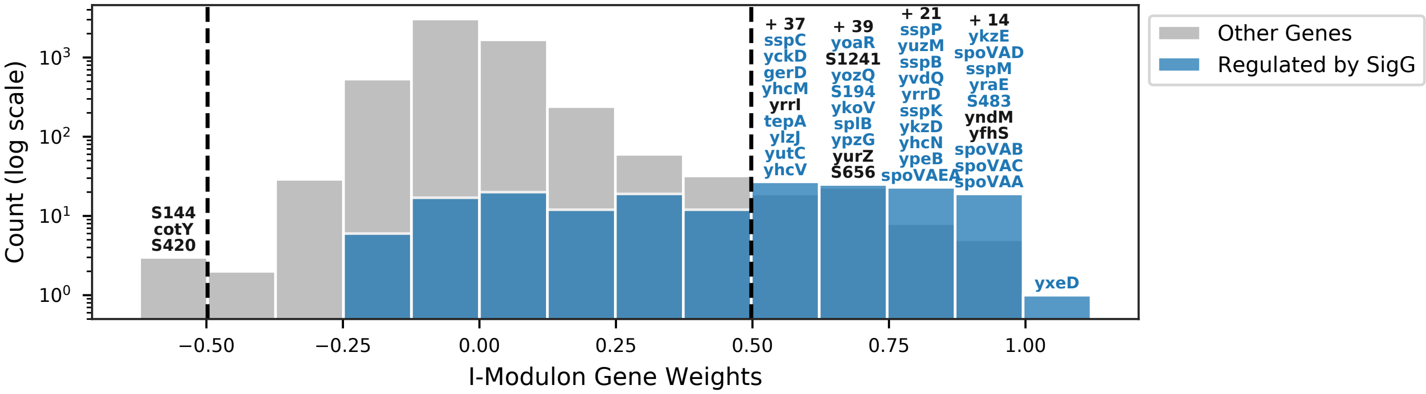


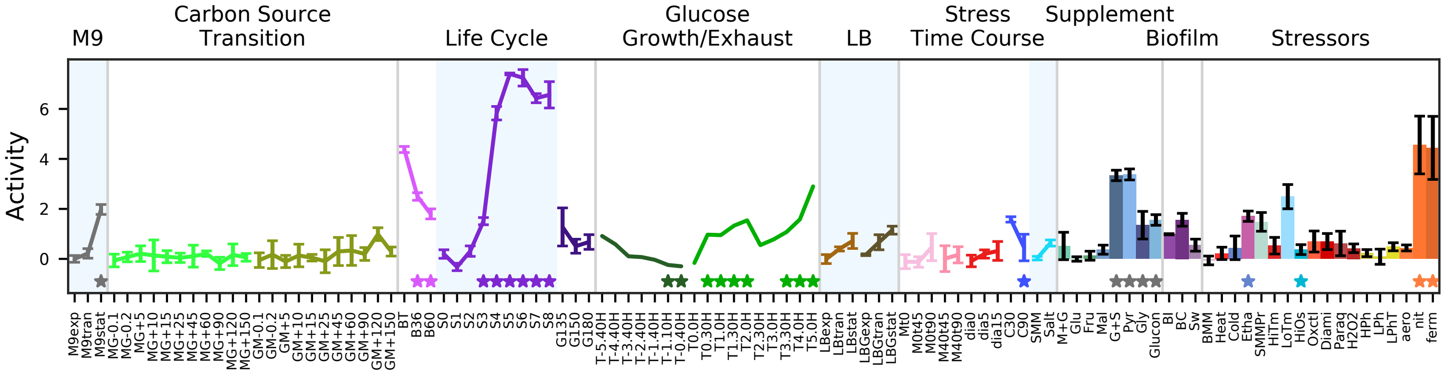


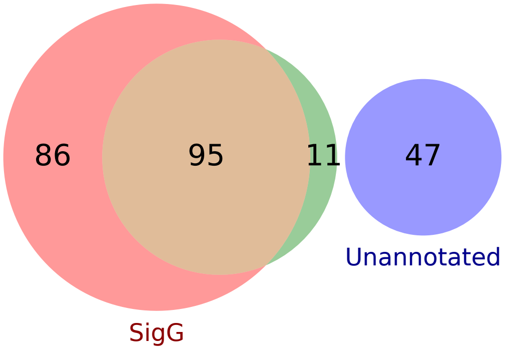


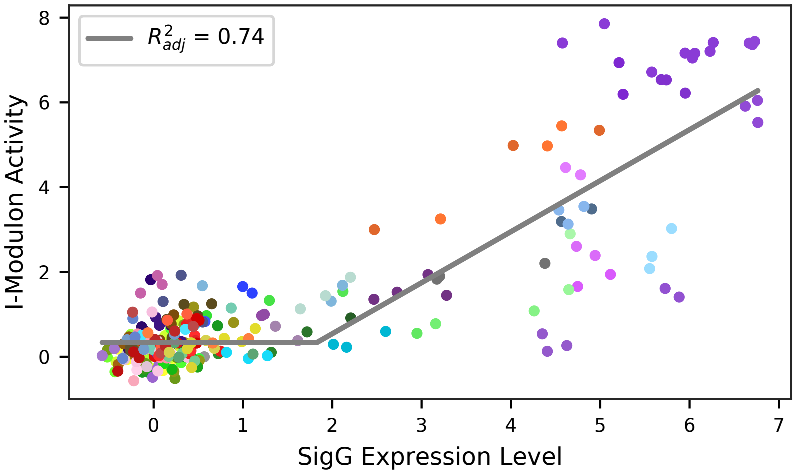

### 52 - SigK-1 – Sporulation 5

Biological Function:
Production of spore coat and crust proteins, L-sulfolactate production, spore coat cross-linking, stress resistance, many unknown proteins

Enriched for known regulon:
SigK

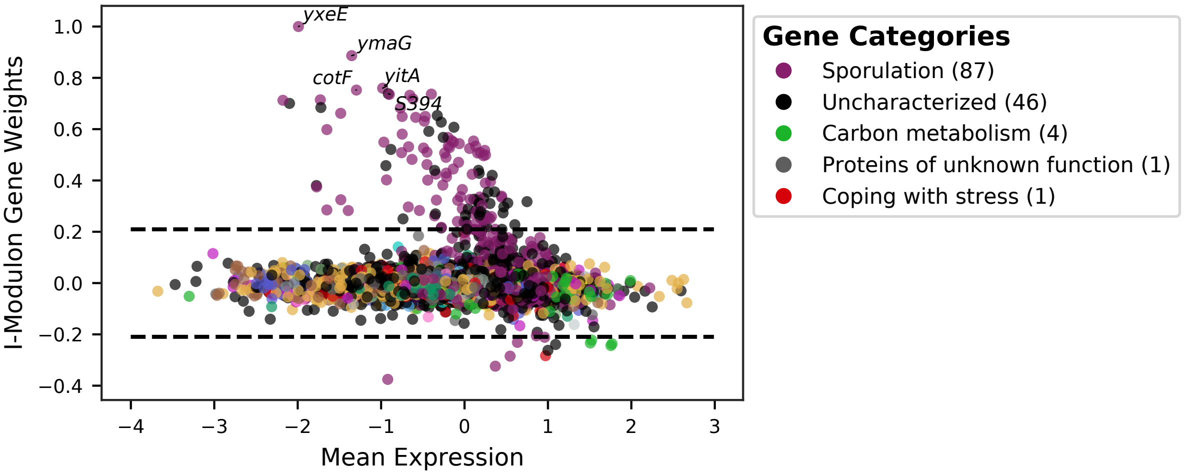


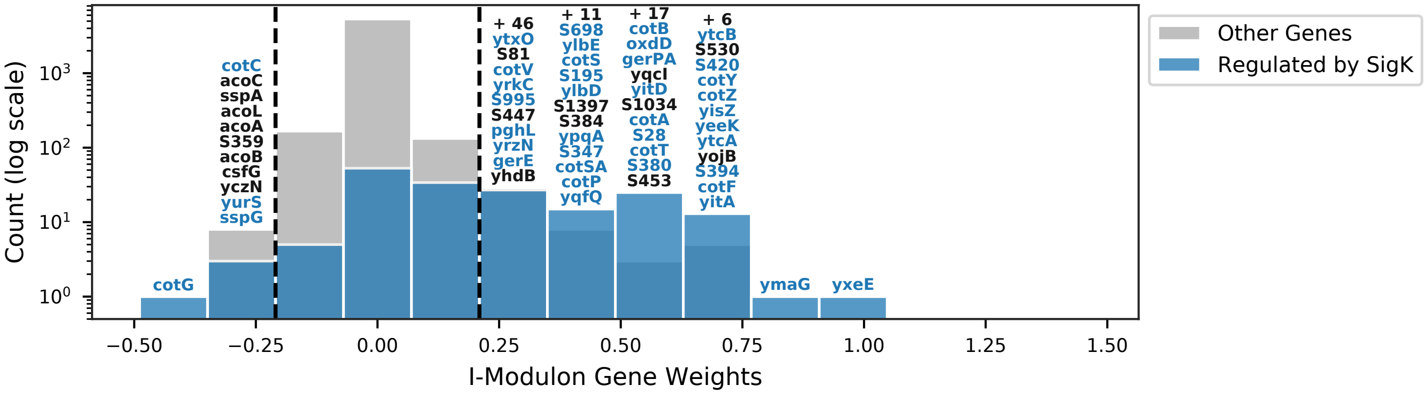


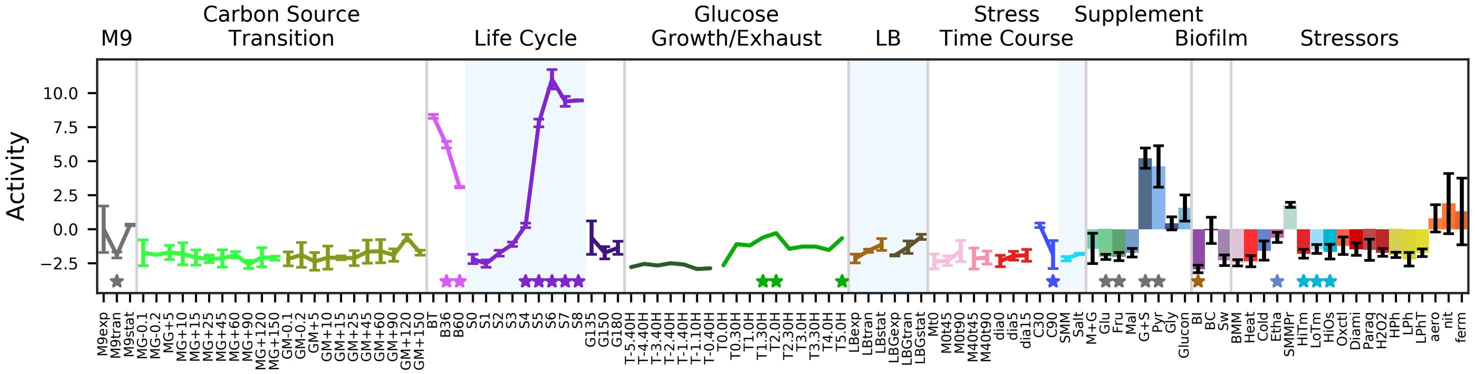


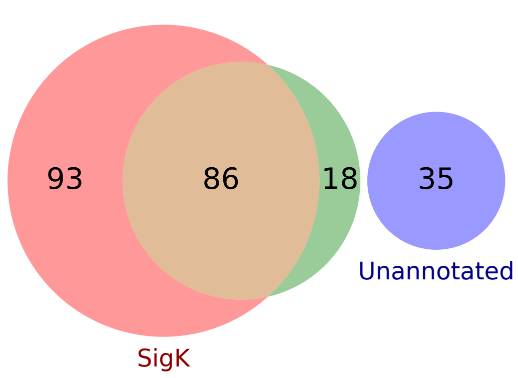


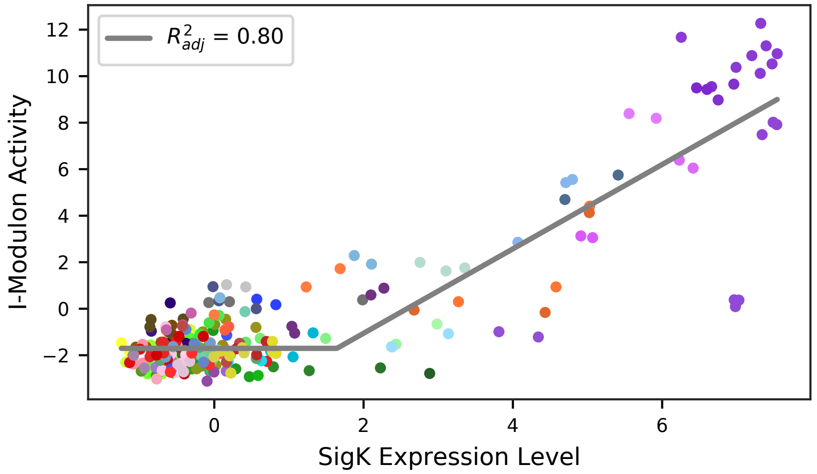

### 53 - SigK-2 – Sporulation 6

Biological Function:
Production of outermost spore coat and crust proteins, mother cell lysis

Enriched for known regulon:
SigK

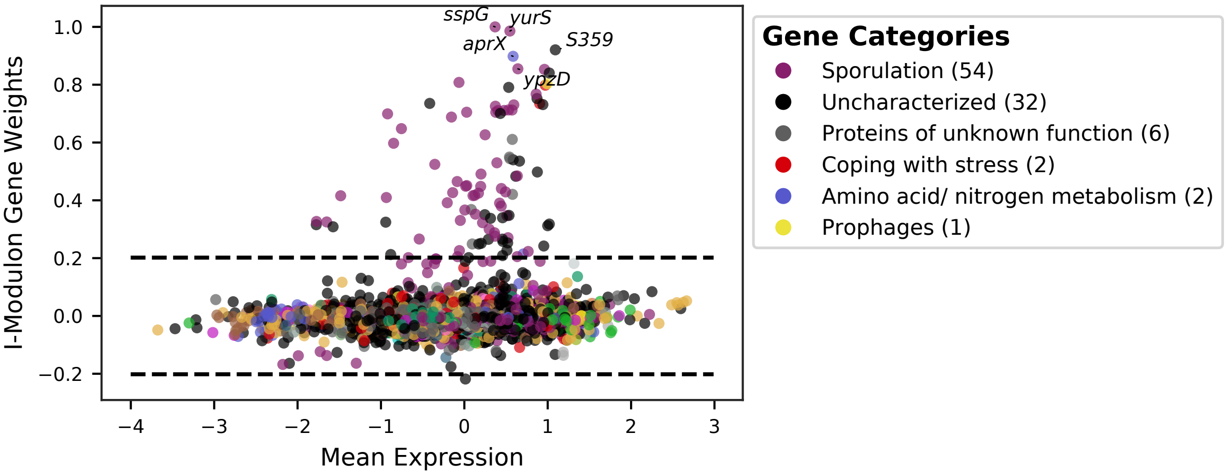


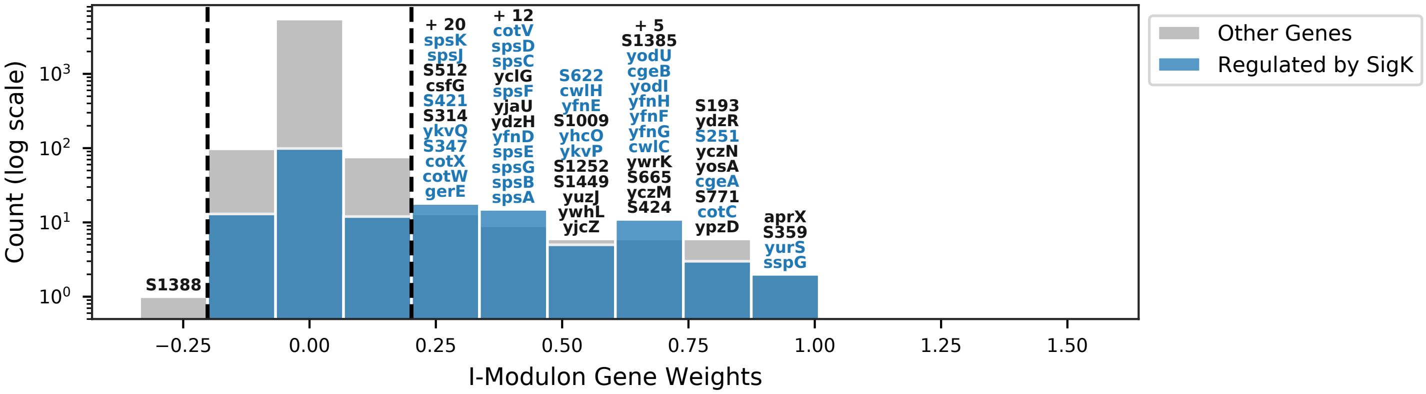


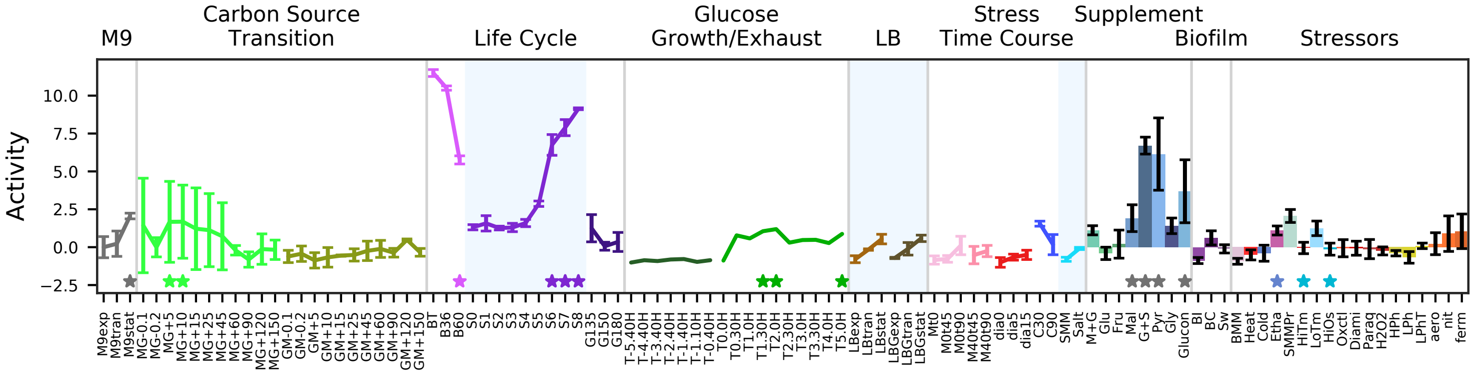


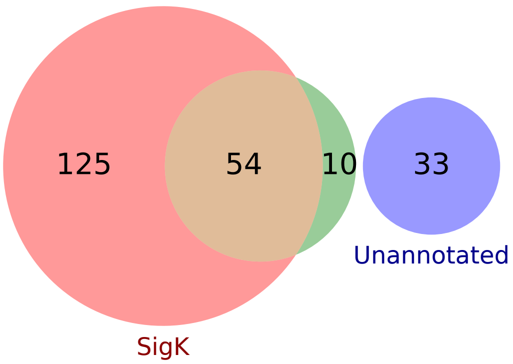


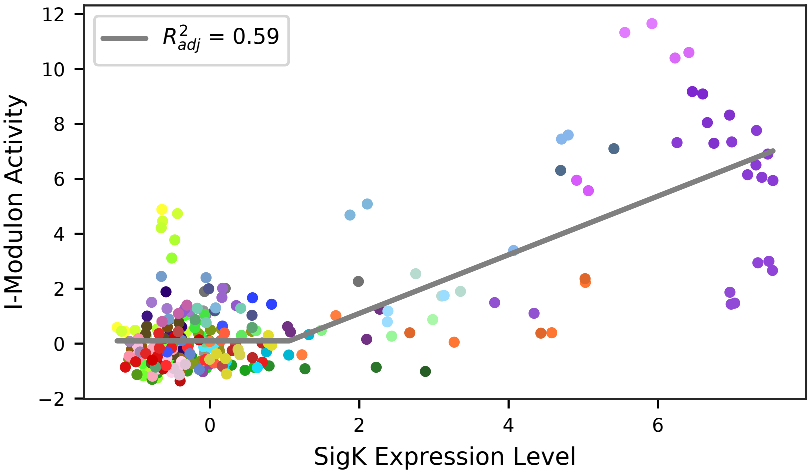

### 54 - GerE / GerR – Mid-late Spore Coat

Biological Function:
Production of spore coat and crust proteins. Omitted from sporulation analysis because all genes are contained by SigK1 & 2, and activity levels are very noisy in non-sporulating conditions.

Subset of known regulon:
[SigK + GerE] / [SigK + GerR]

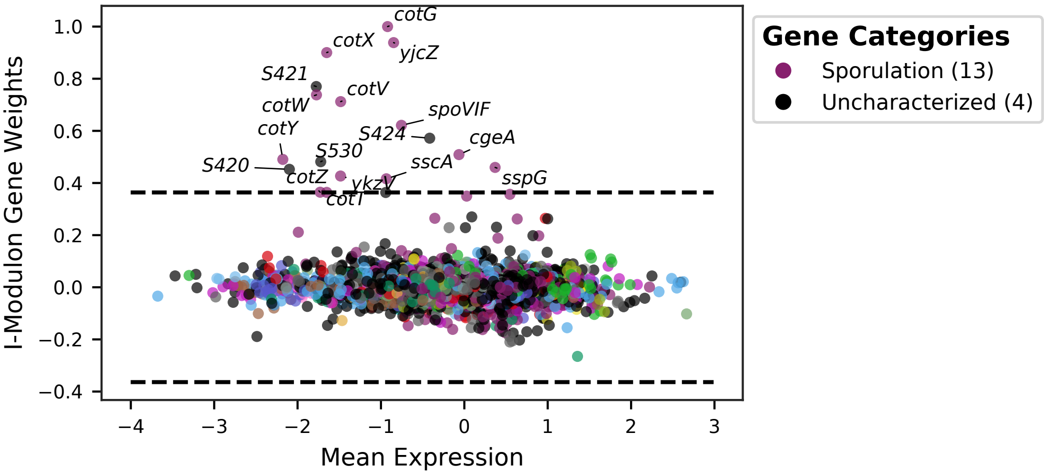


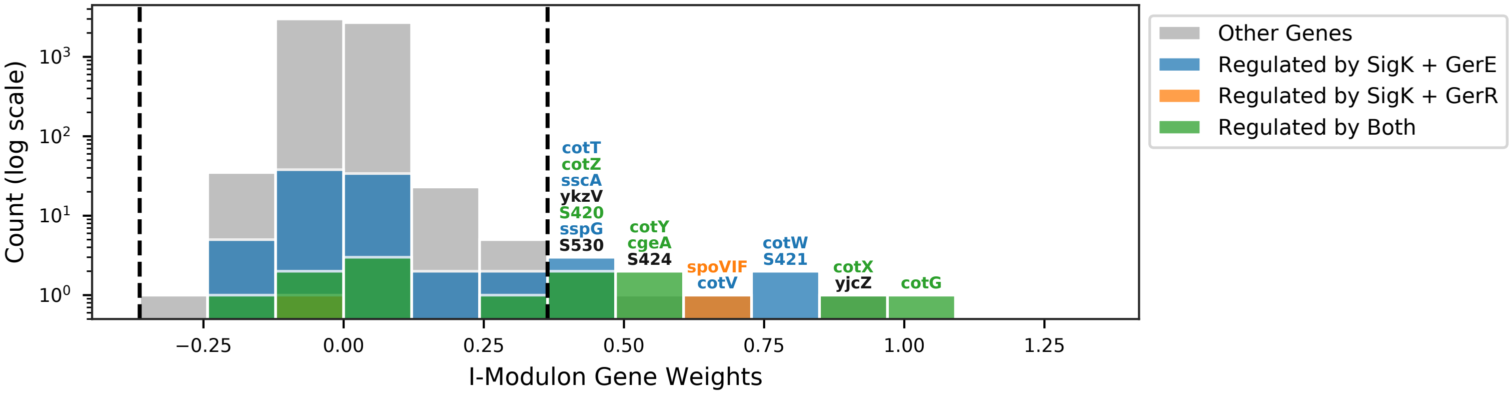


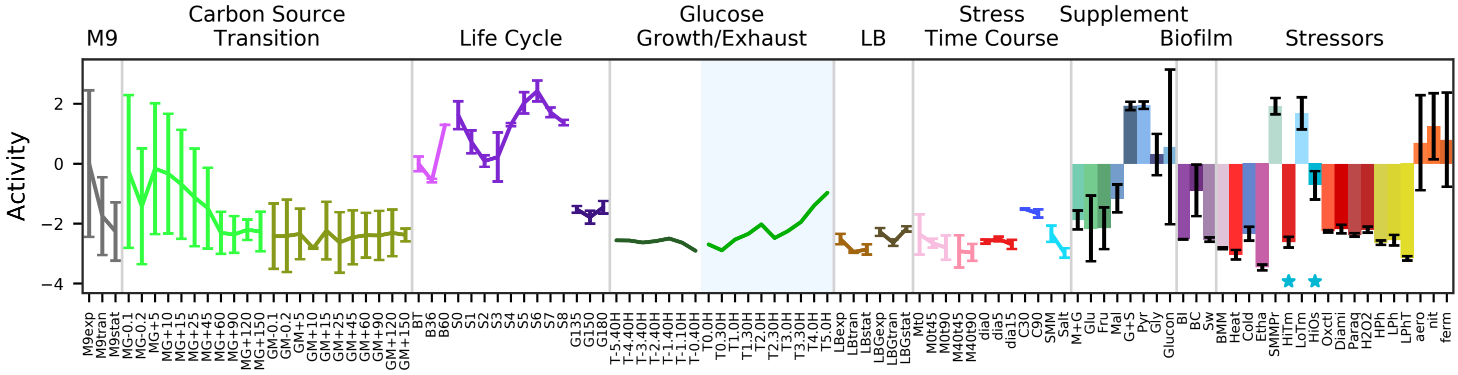


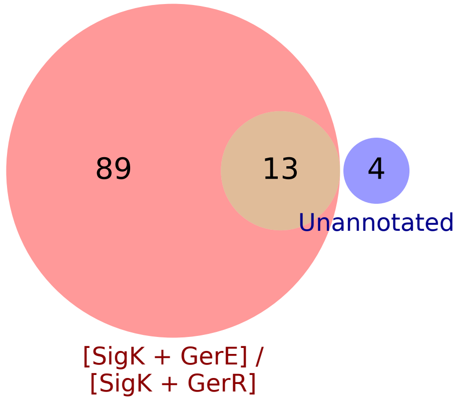


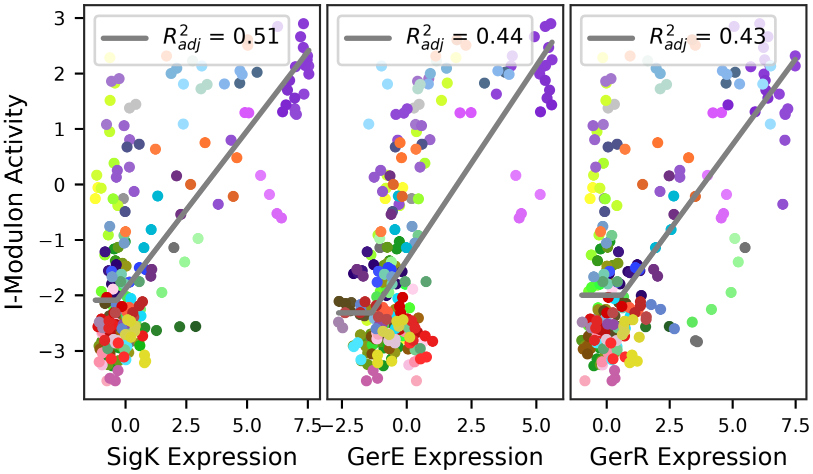

### 55 - Xpf – PBSX Prophage

Biological Function:
Prophage element that induces production of bacteriocin and cell lysis in response to DNA-damaging agents

Well-defined regulon:
Xpf

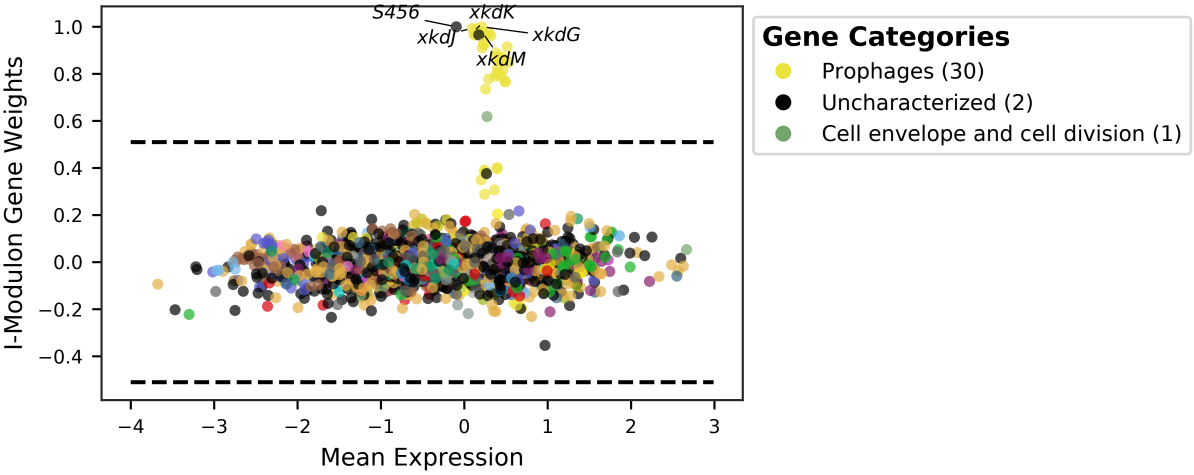


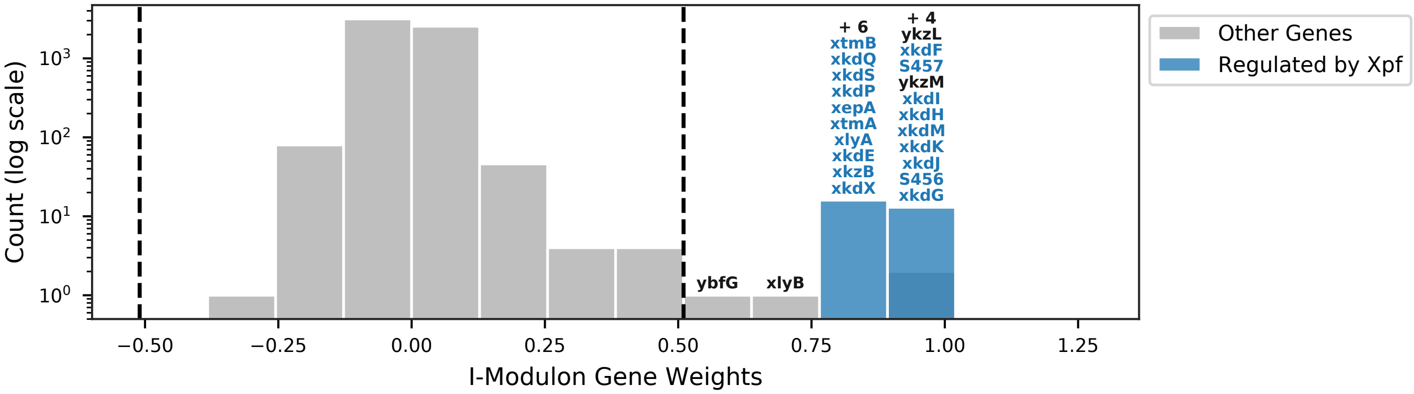


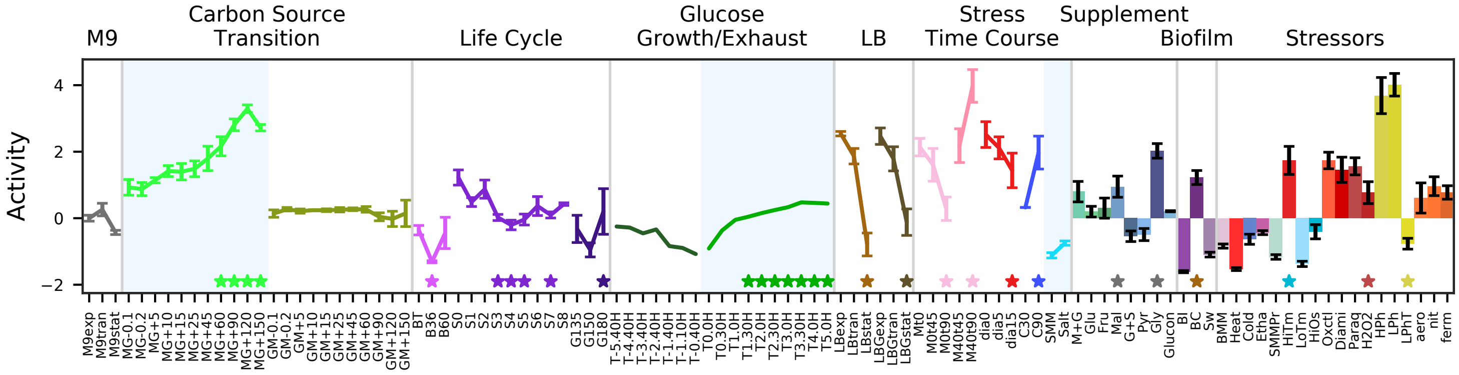


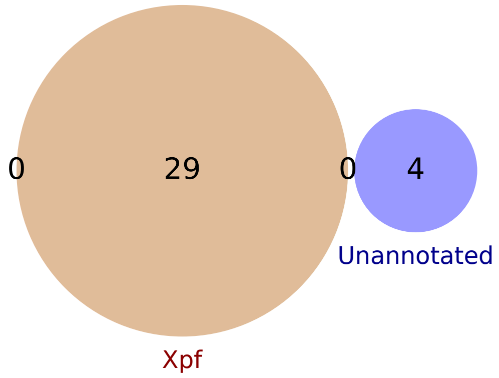


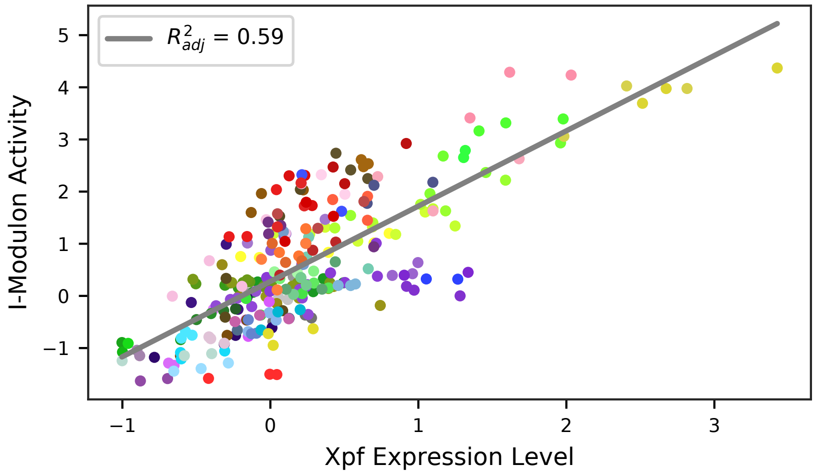

### 56 - Spβ-1 – SP-β Prophage 1

Biological Function:
Temperature-responsive SP-beta prophage. Response to stressors such as cold and ethanol. Subset of SPbeta Prophage 2.

No known regulator

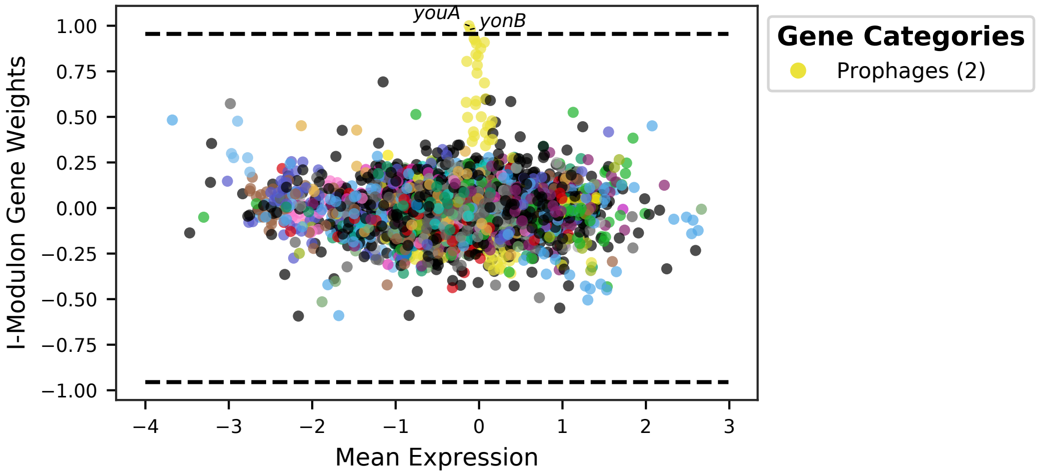


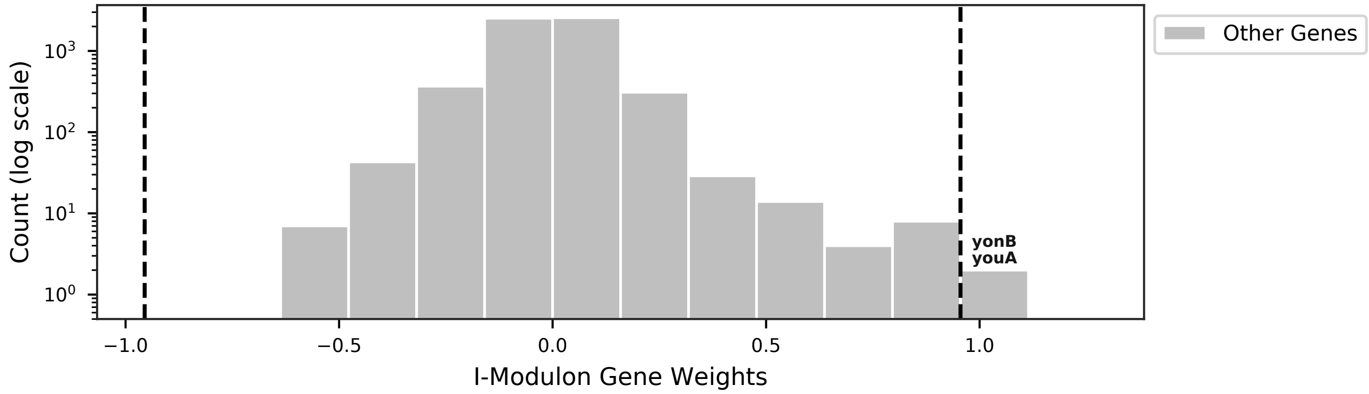


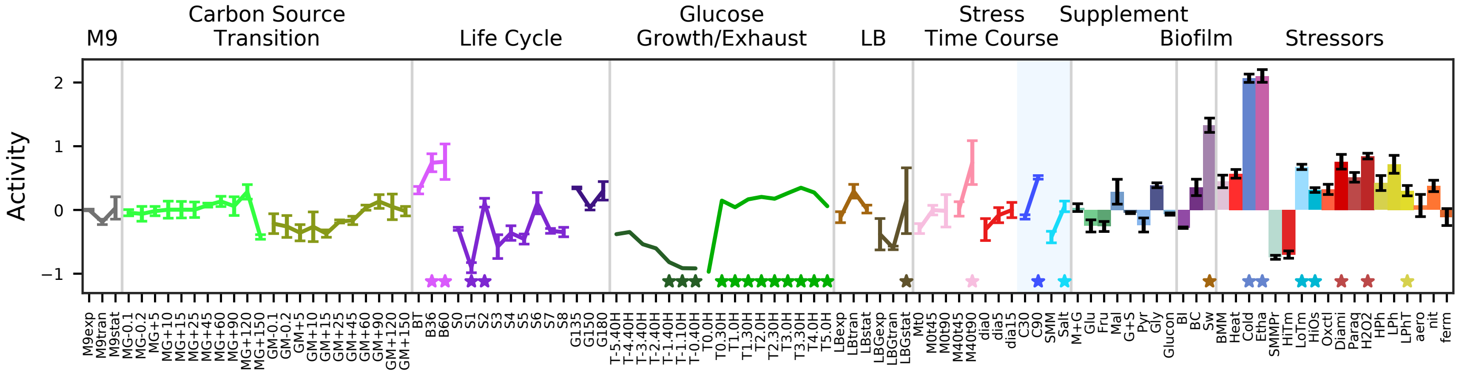


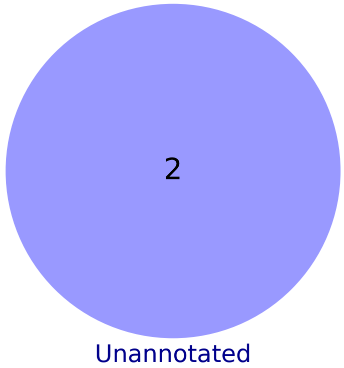

### 57 - Spβ-2 – SP-β Prophage 2

Biological Function:
Temperature-responsive SP-beta prophage. Prophage, unknown, and putative metabolic genes. Response to mitomycin.

Contains unknown genes and known regulon:
CsoR + SigA + SigK + SigE

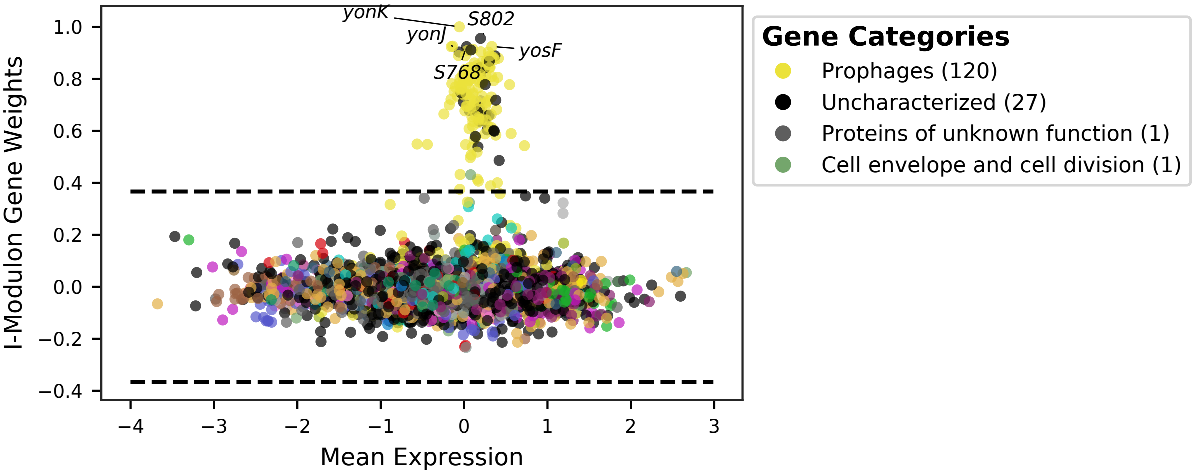


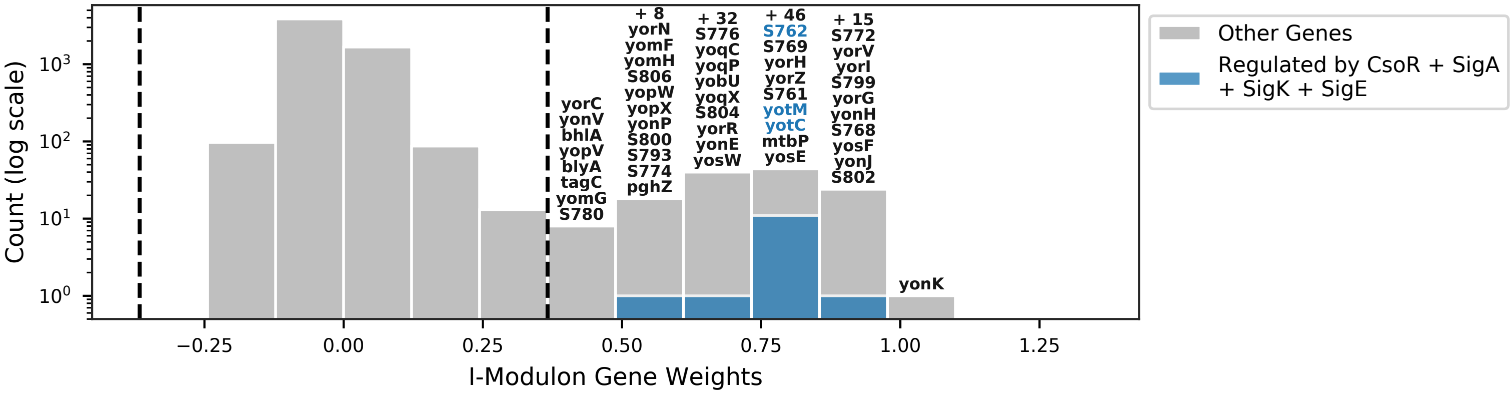


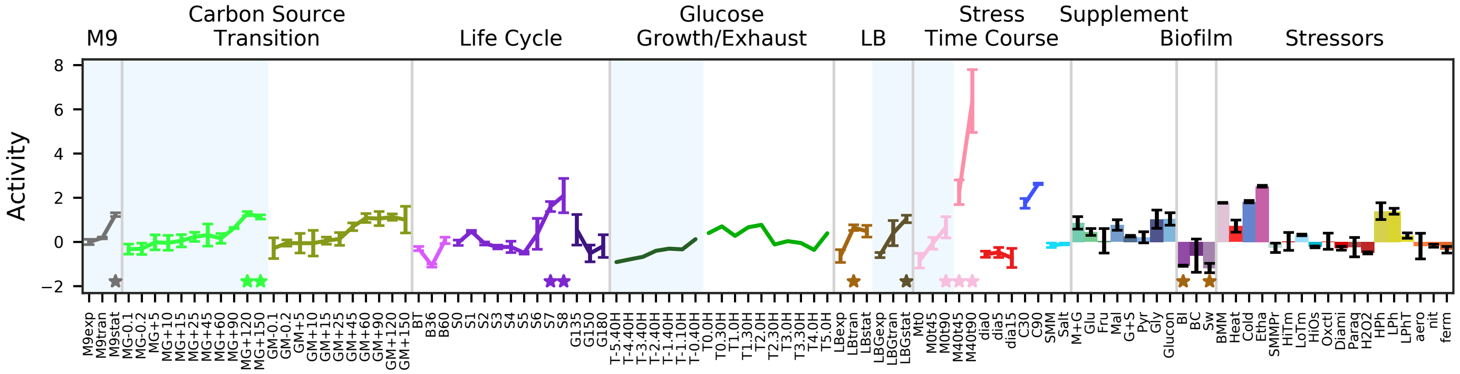


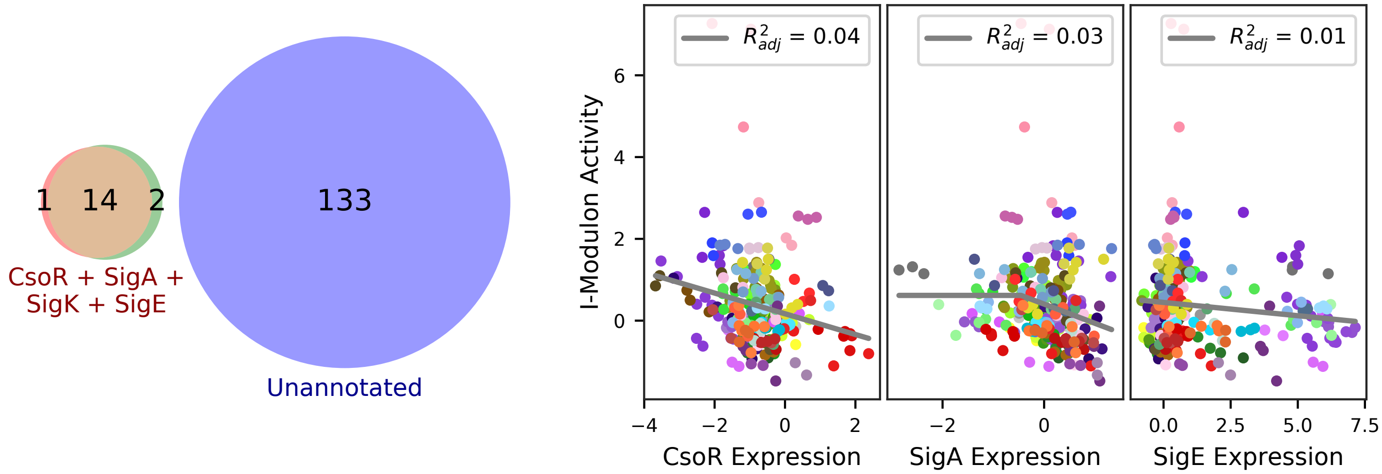

### 58 - ImmR – ICEBs1

Biological Function:
Mobile genetic element that participates in horizontal gene transfer

Well-defined regulon:
ImmR

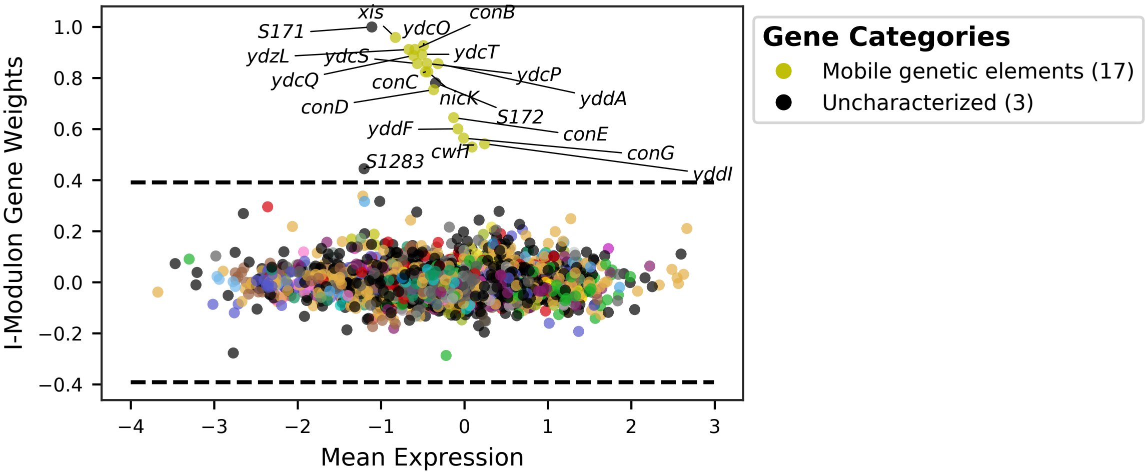


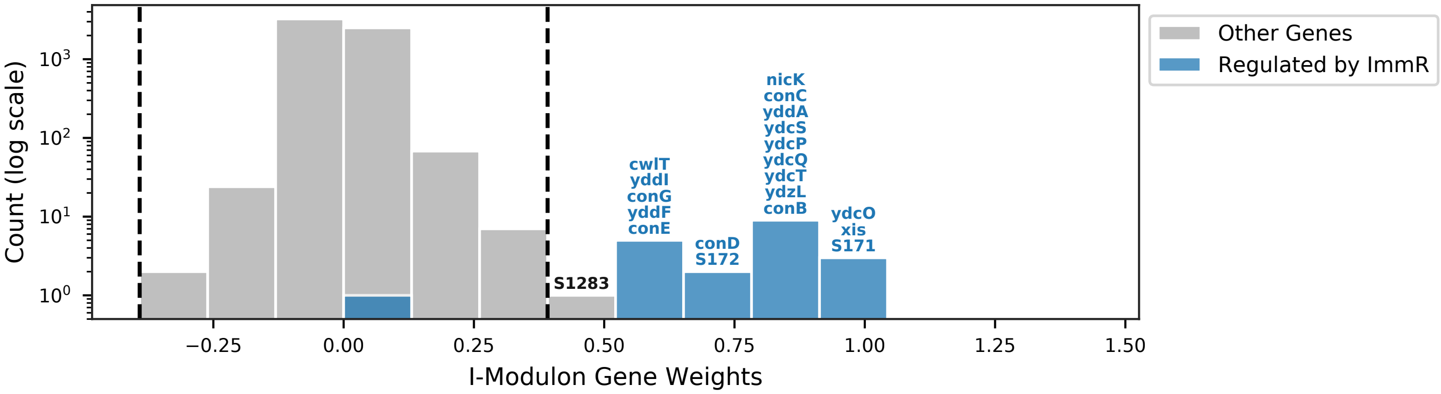


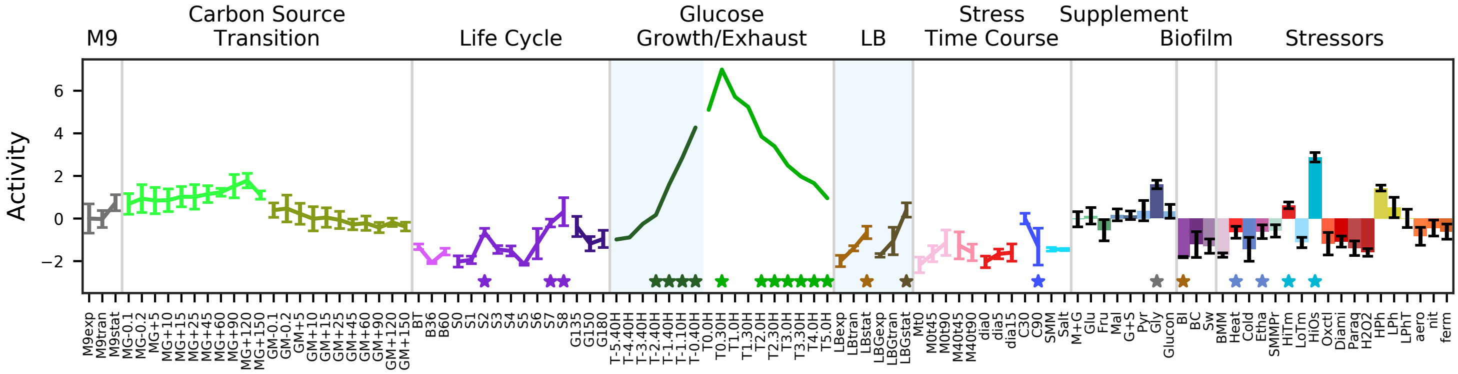


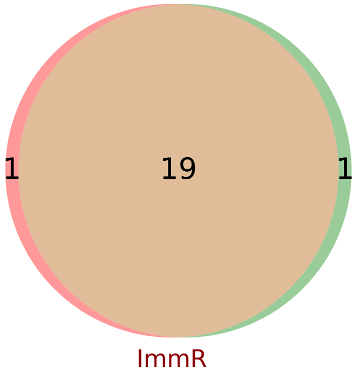


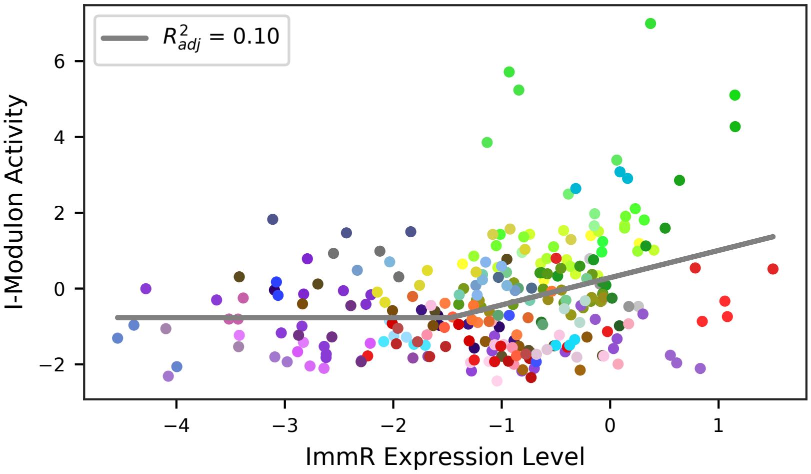

### 59 - SknR – Skin Element

Biological Function:
Prophage element that disrupts the SigK gene but also plays roles in preventing early germination and regulating proliferation. Unknown and toxin resistance genes.

Enriched for known regulon:
SknR

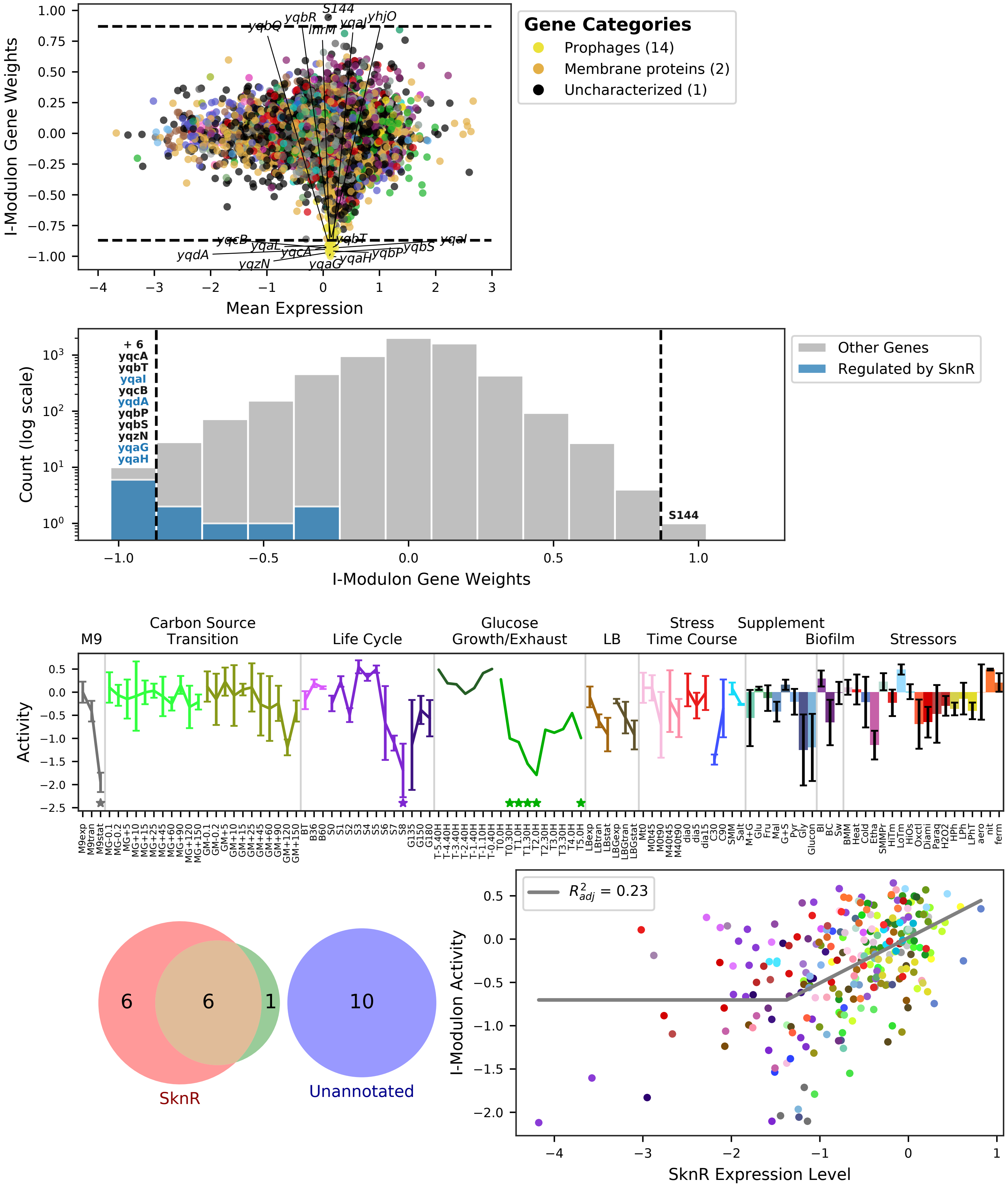

### 60 - IolR / MelR – Inositol, Melibiose, and Acetoin

Biological Function:
Uptake and utilization of inositol, melibiose/raffinose, and acetoin

Well-defined regulon:
[IolR] / [MelR] / [AcoR]

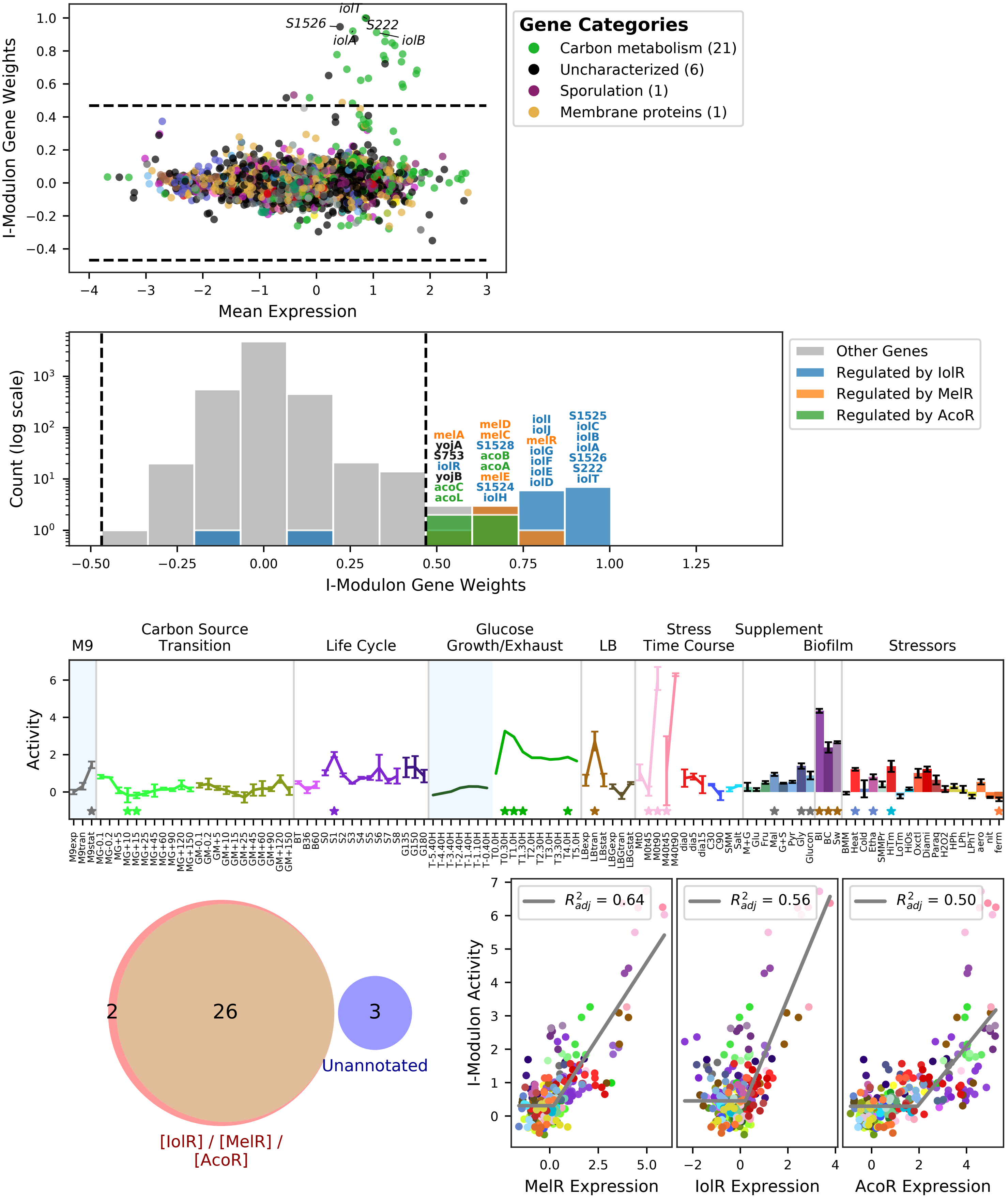

### 61 - NadR / BirA – NAD and Biotin

Biological Function:
Biosynthesis of NAD and biotin, plus safA & coxA sporulation proteins and repression of manganese uptake

Well-defined regulon:
[NadR] / [BirA]

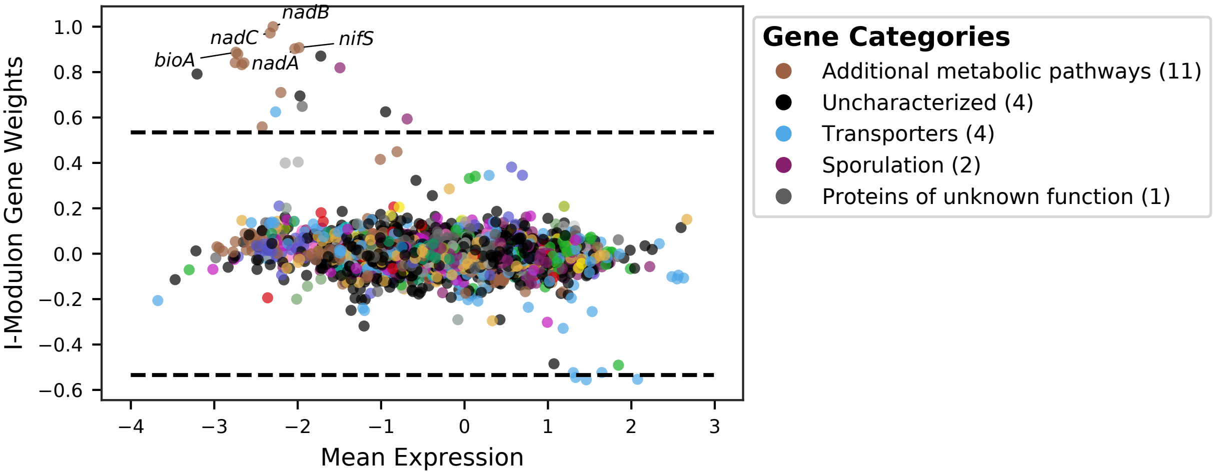


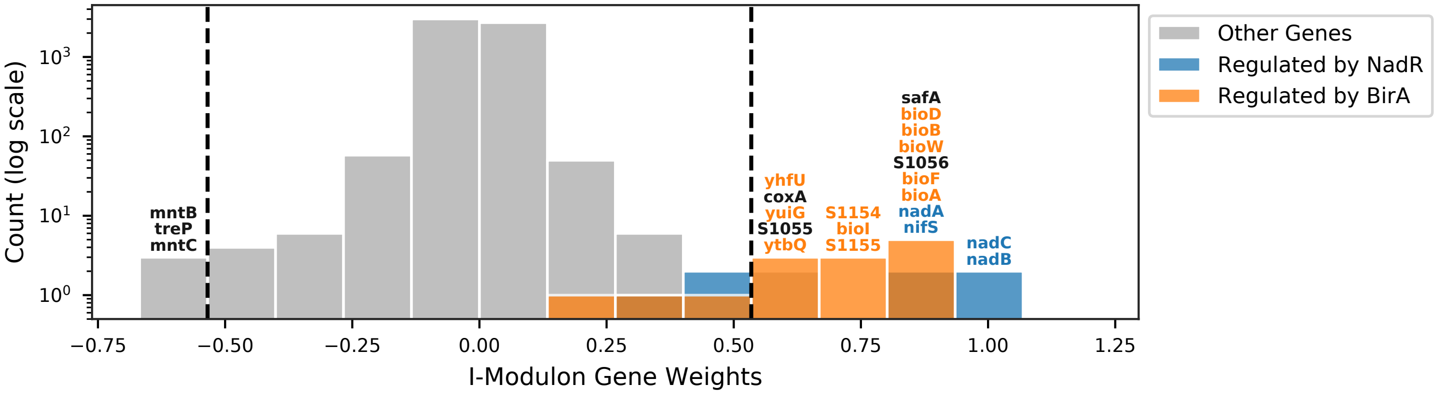


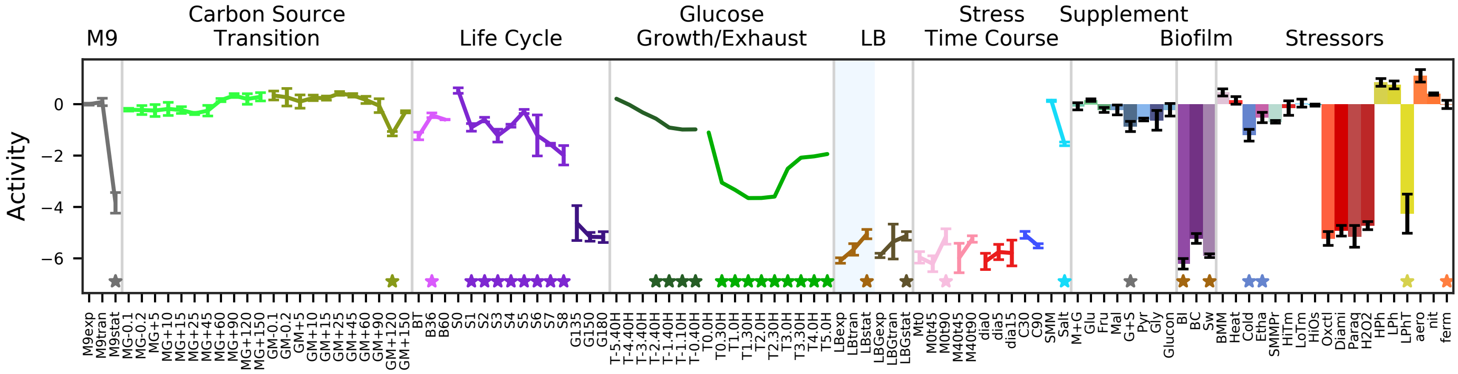


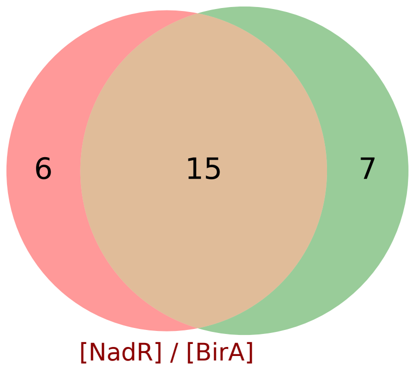


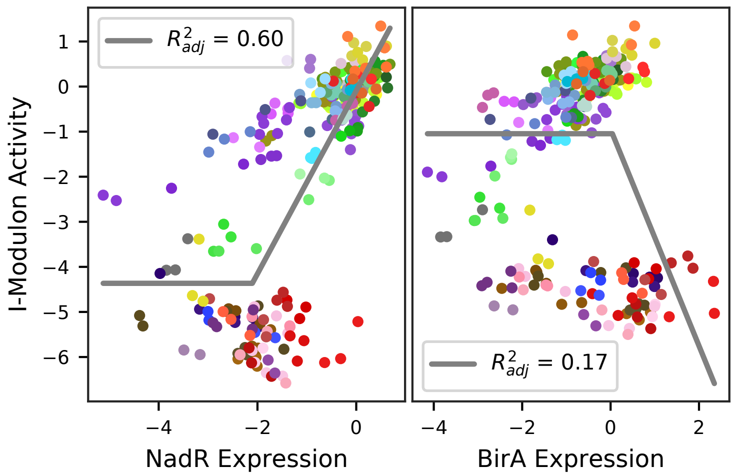

### 62 - MtlR / AnsR – Mannitol, Asparagine, and Aspartate

Biological Function:
Uptake and utilization of mannitol, degradation of asparagine and aspartate

Well-defined regulon:
[MtlR] / [AnsR + SigA]

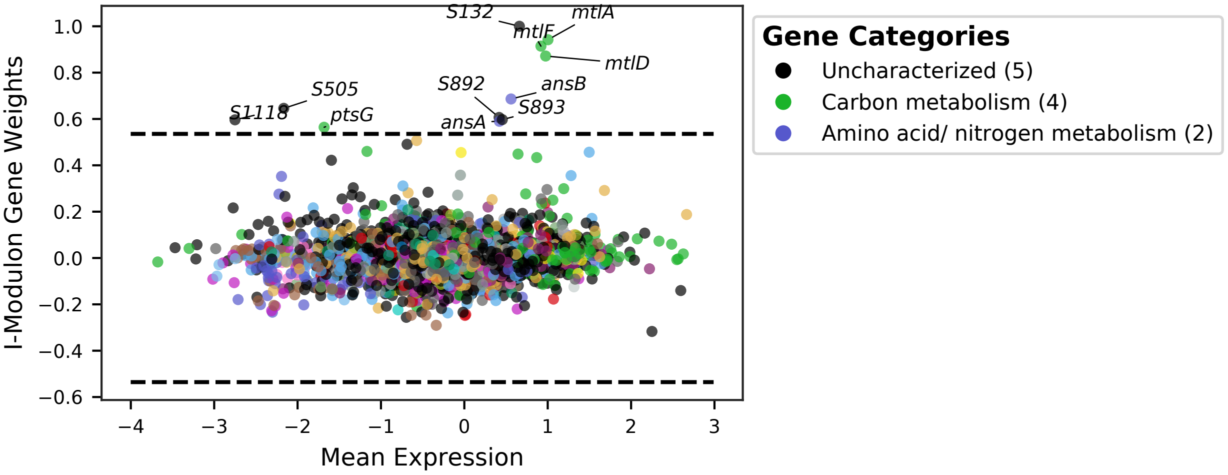


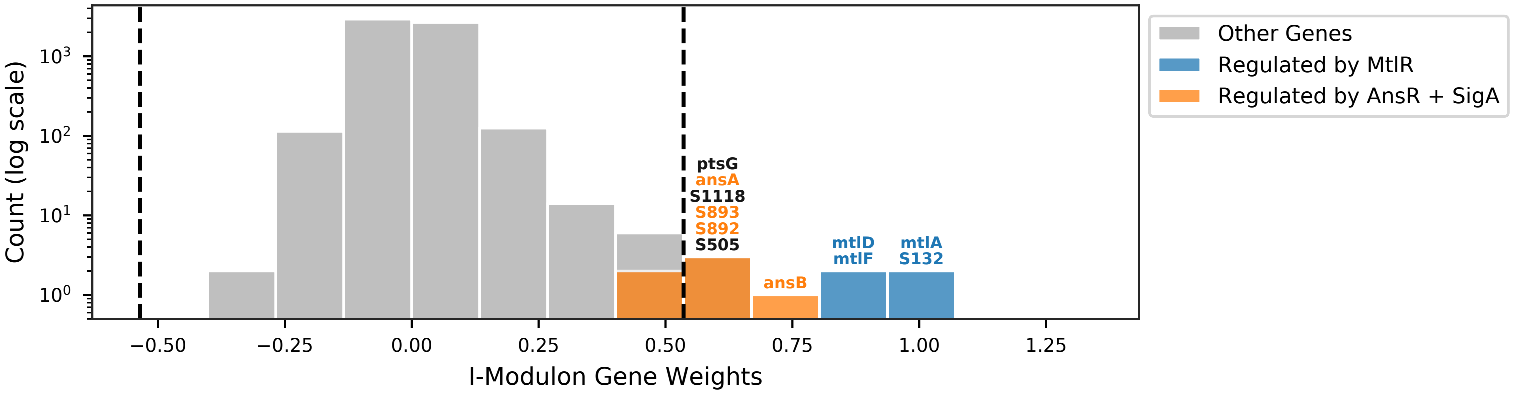


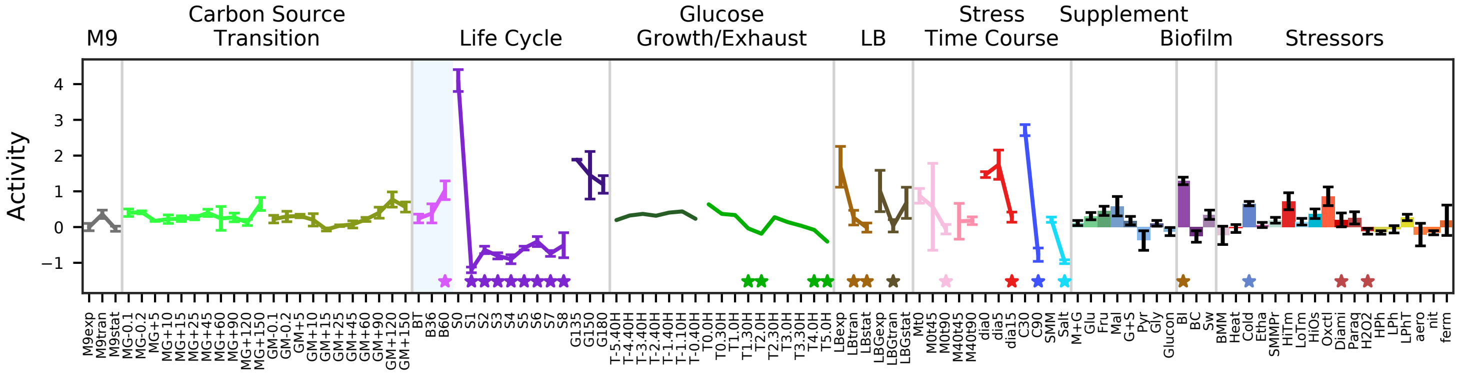


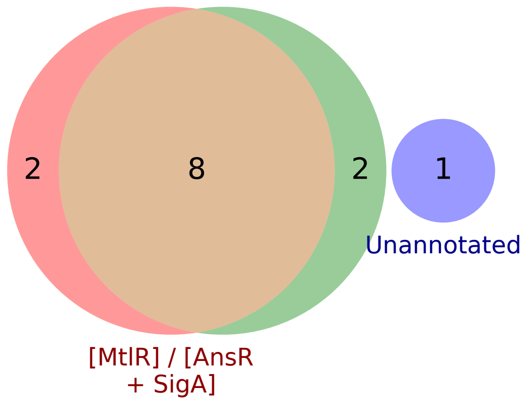


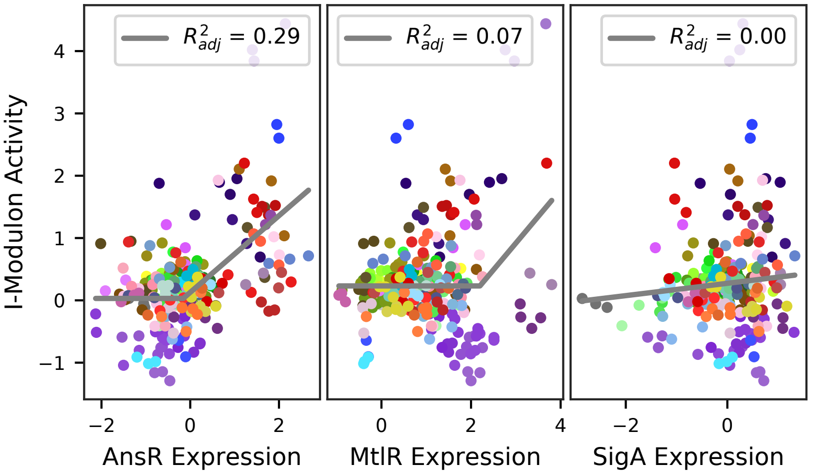

### 63 - TnrA / PucR – Nitrogen Limitation

Biological Function:
Ammonium uptake, utilization of nitrate and xanthine. Suggests nitrate limitation in the BT condition

Subset of known regulon:
[SigA + TnrA] / [SigA + PucR]

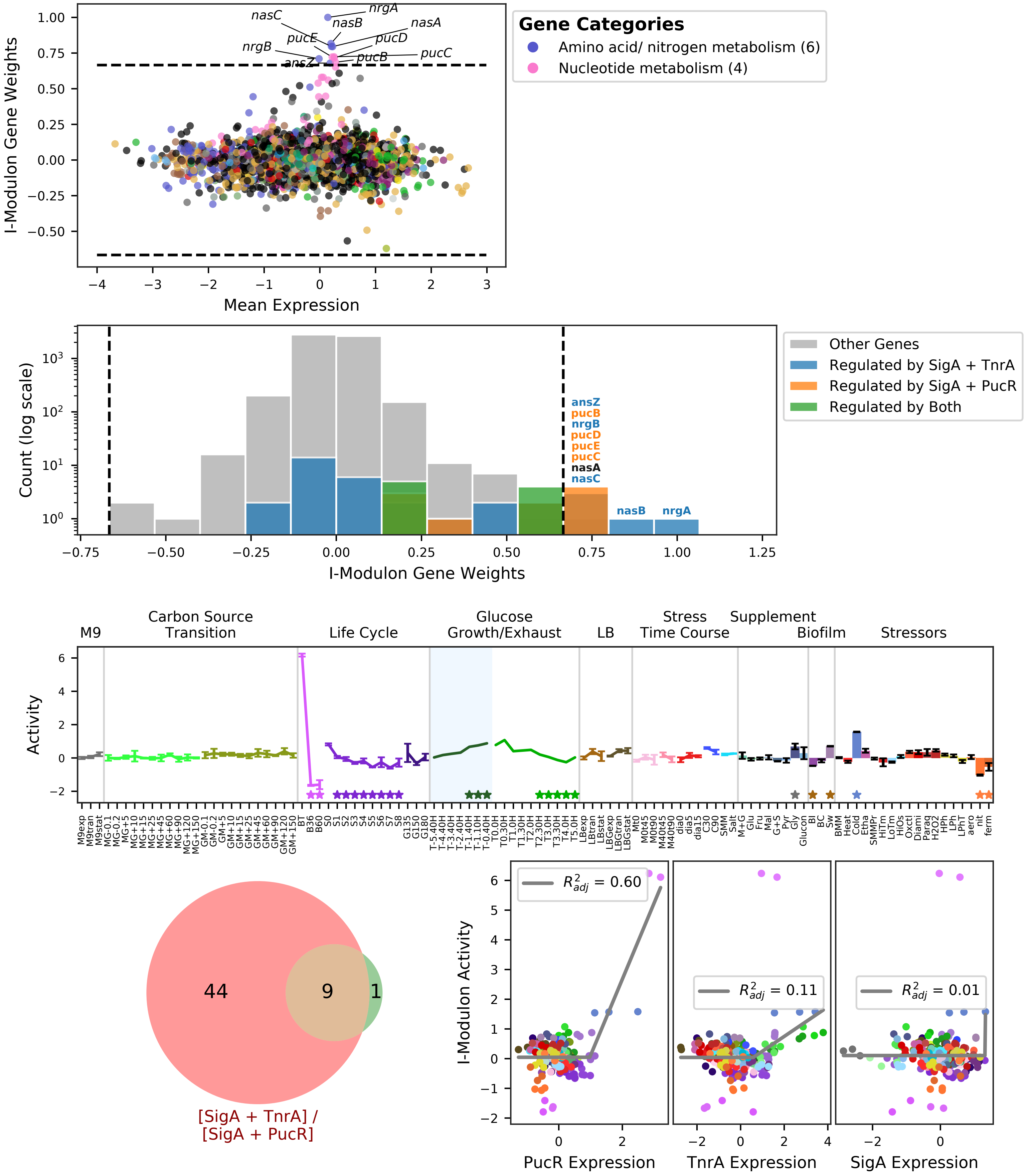

### 64 - CymR / NsrR – Sulfur and Nitrate Respiration

Biological Function:
Import and utilization of sulfonate and cystine for cysteine production. Nitrite reduction

Subset of known regulon:
[SigA + CymR] / [ResD + SigA + NsrR]

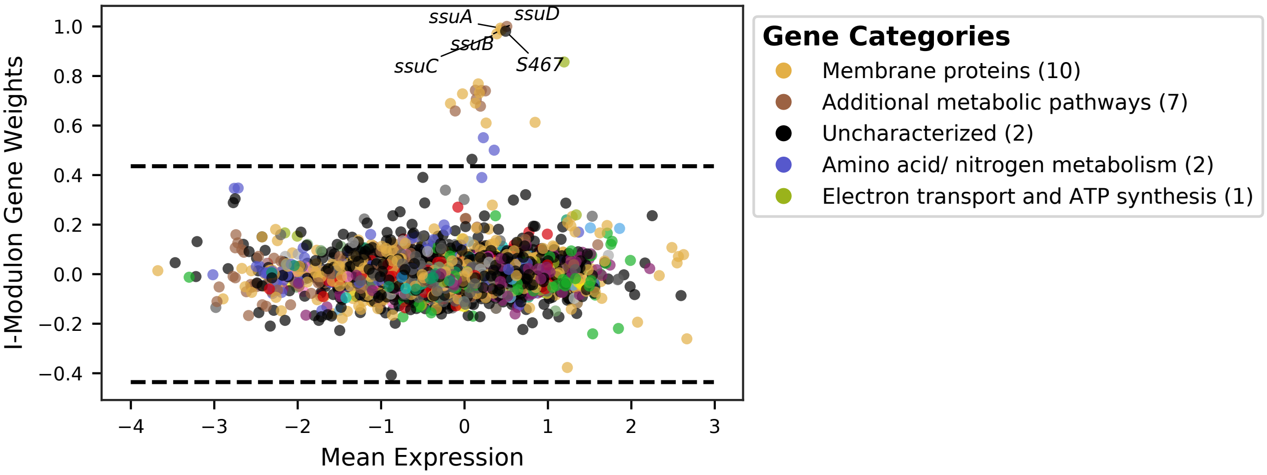


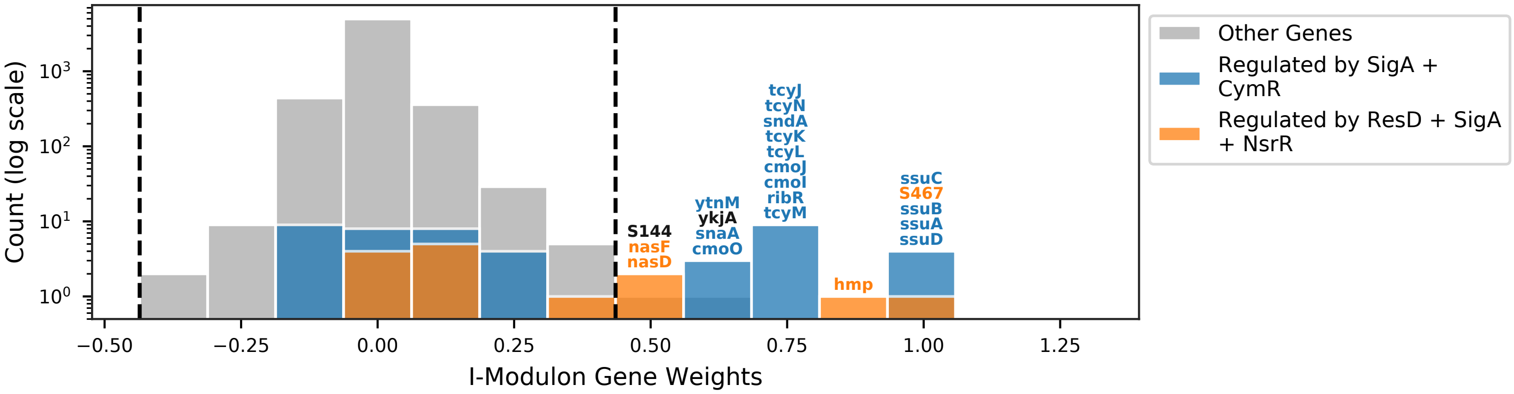


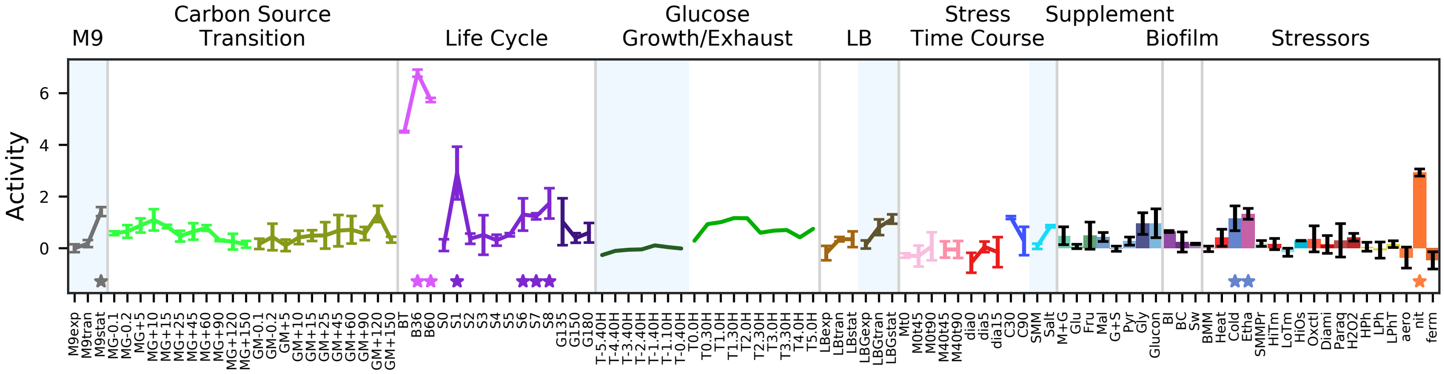


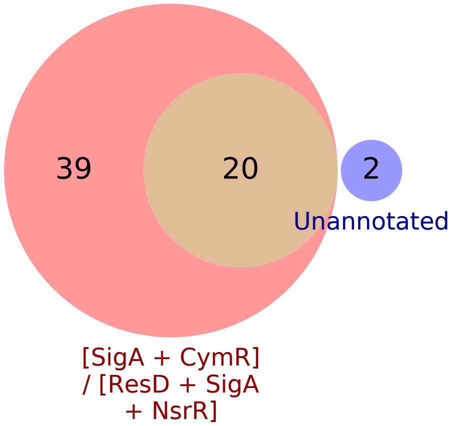


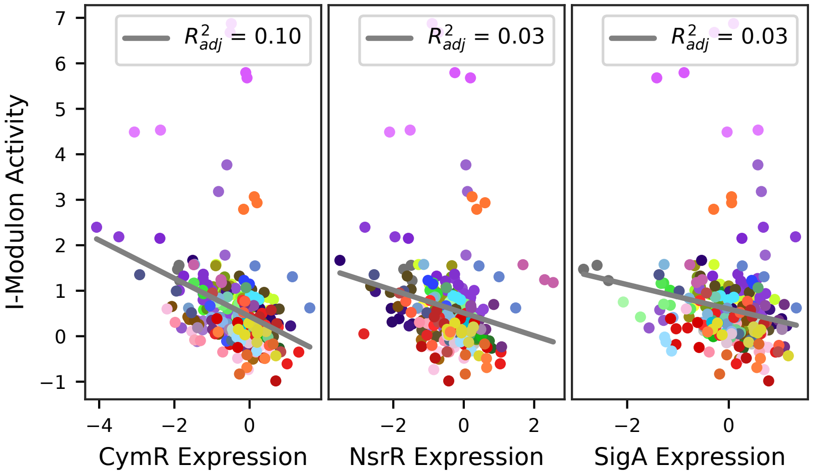

### 65 - ArsR+ – Diamide Stress

Biological Function:
Detoxification in response to: oxidative protein damage, arsenic, catechol, copper, toxic quinones

Enriched for known regulon:
[ArsR] / [YodB] / [SigA + CtsR] / [SigA + CsoR]

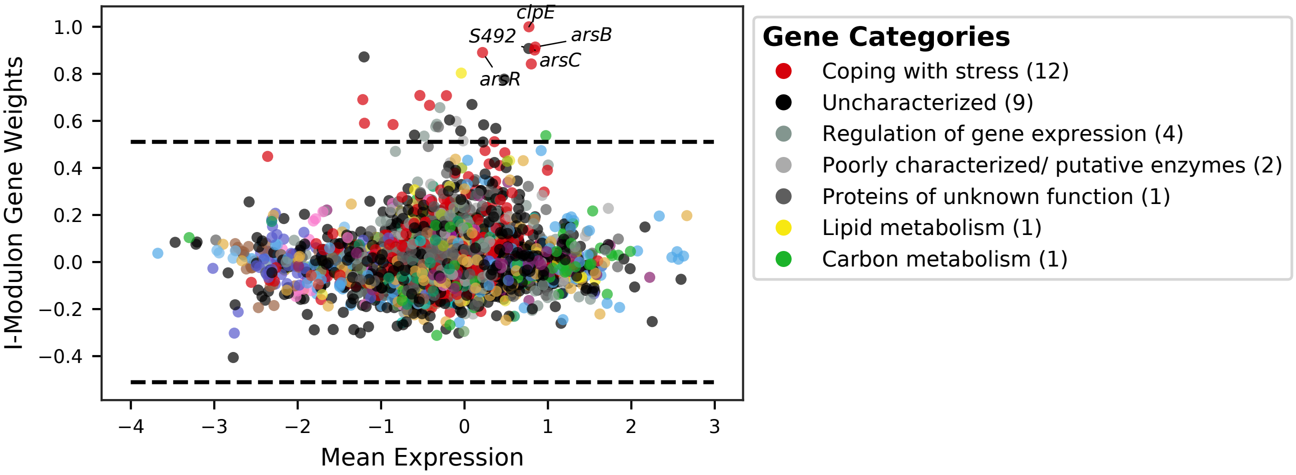


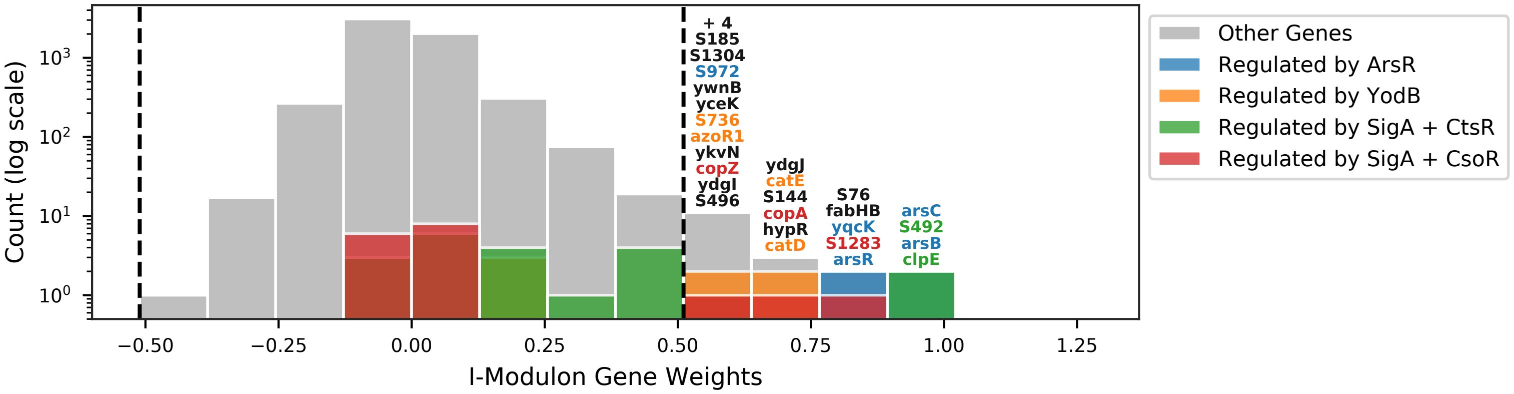


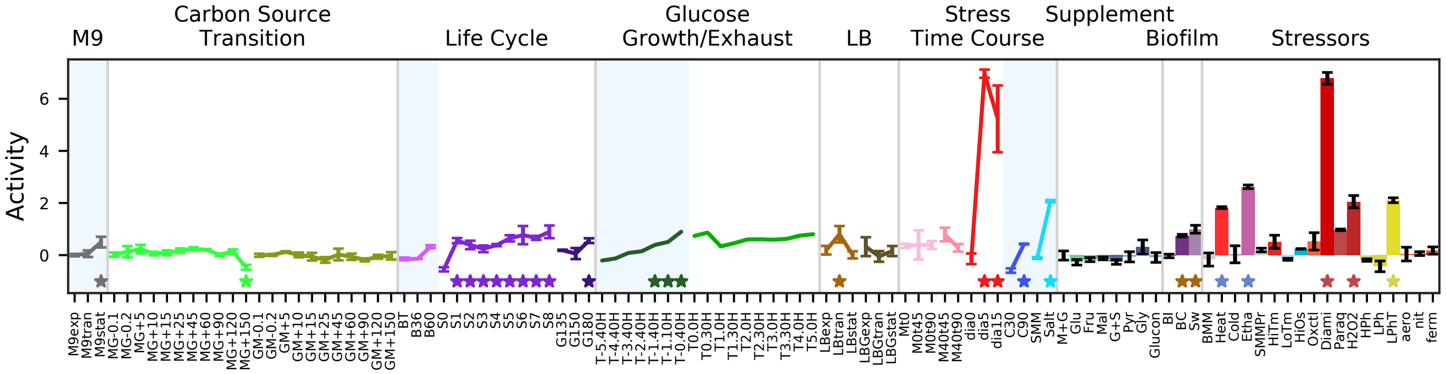


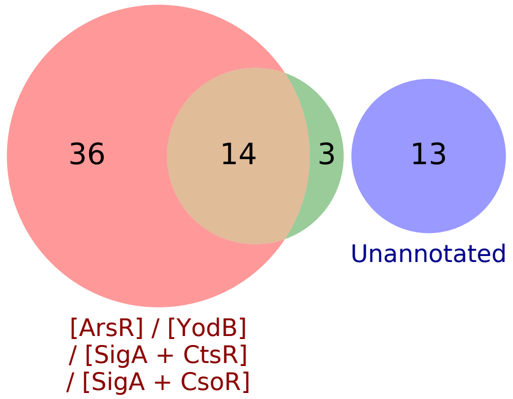


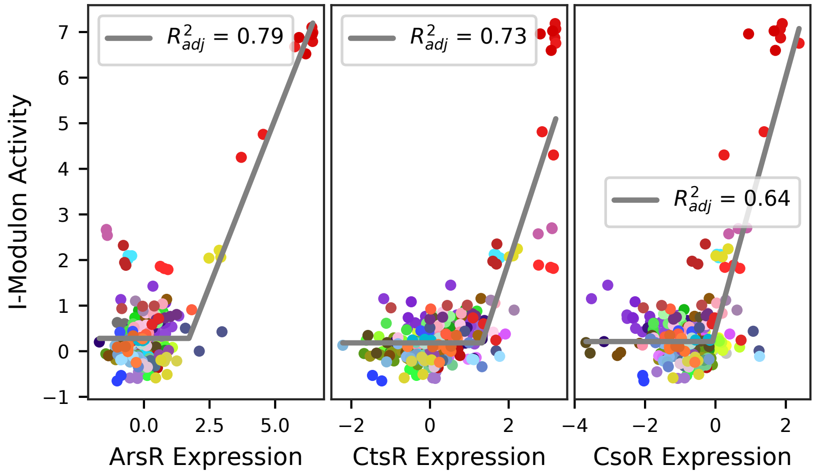

### 66 - Uncharacterized 1

Biological Function:
Contains a SigE/SigG inhibitor, spore coat proteins, translational proteins, oxidative stress response, many RNAs

No known regulator

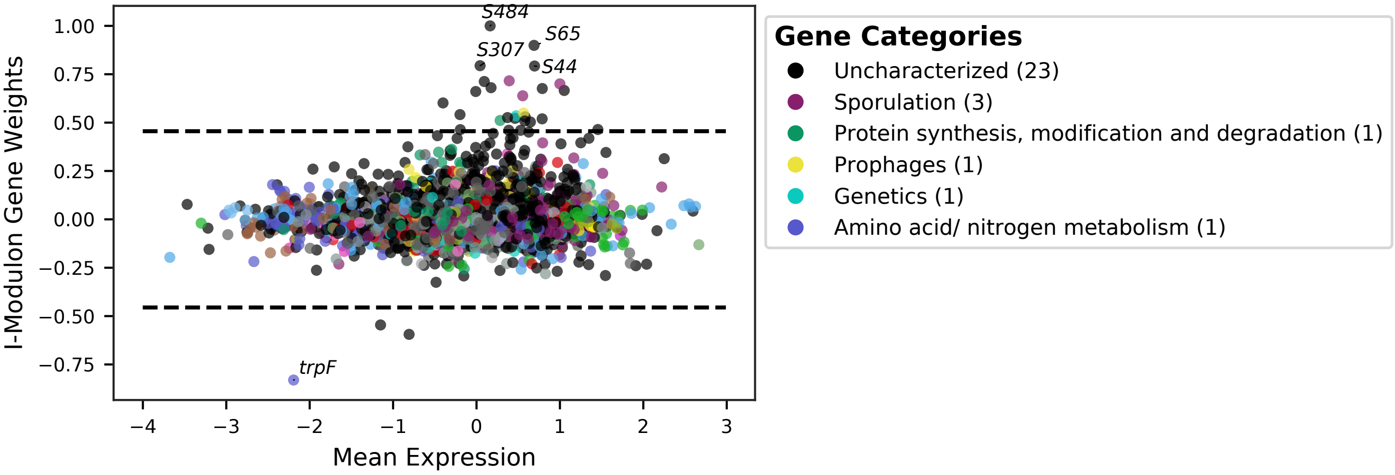


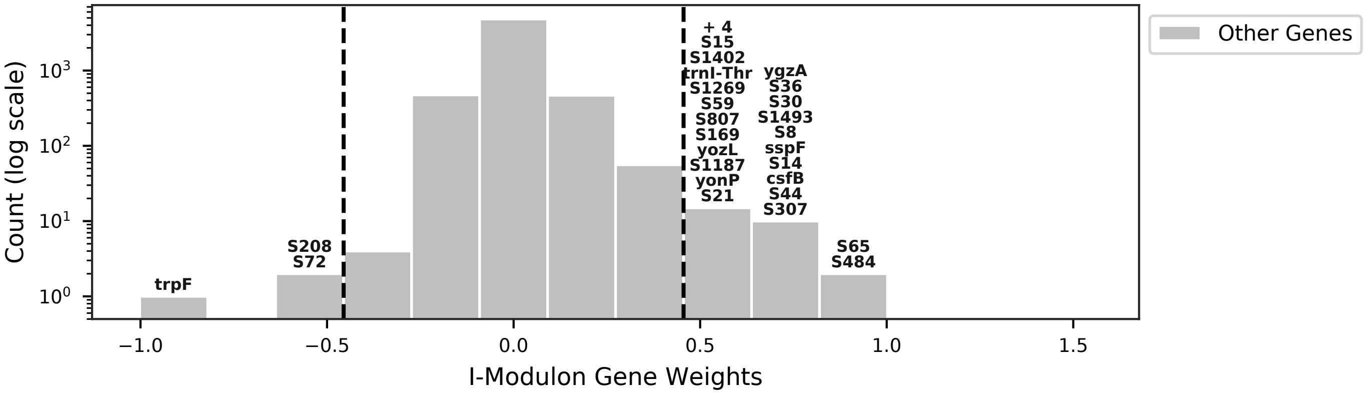


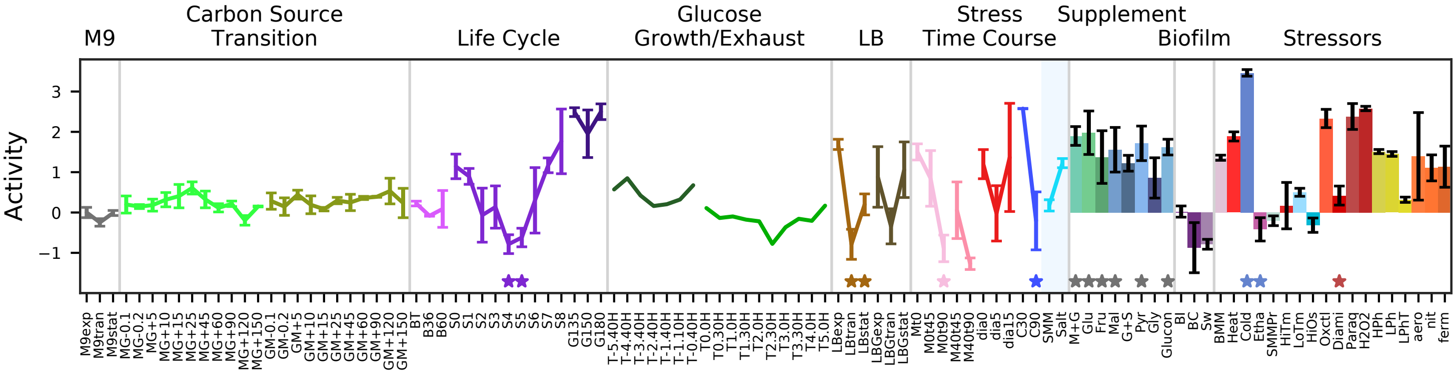


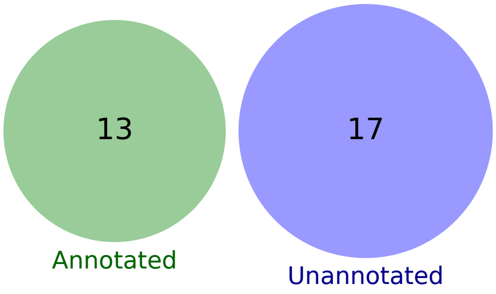

### 67 - Uncharacterized 2

Biological Function:
Contains cold shock, sporulation onset, toxin, amino acid homeostasis, translataional, other proteins, many RNAs

No known regulator

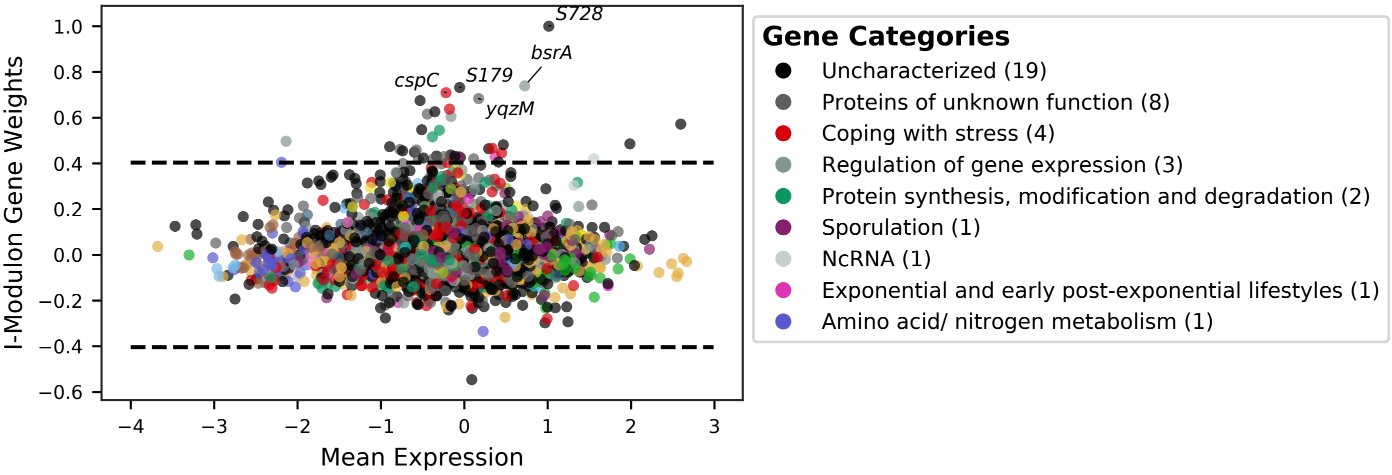


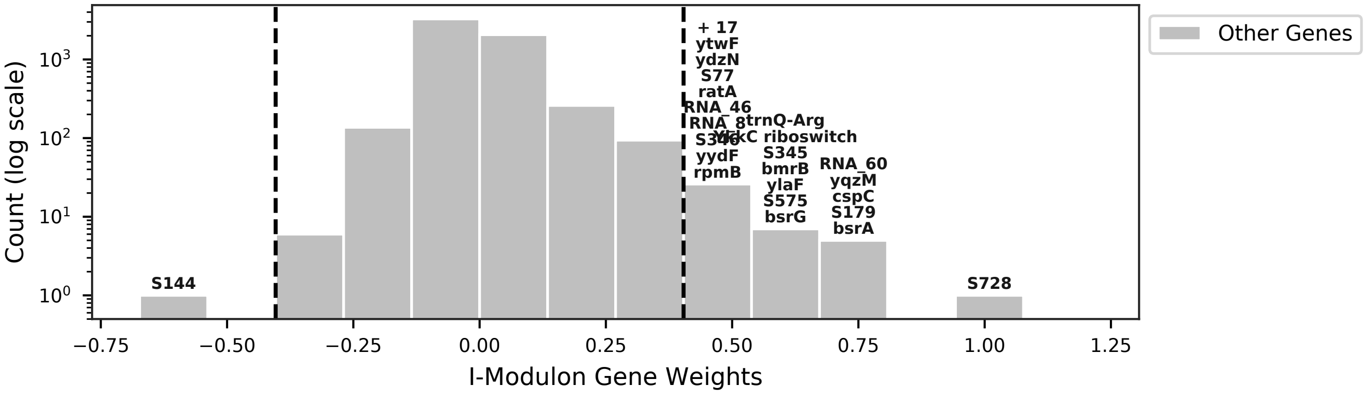


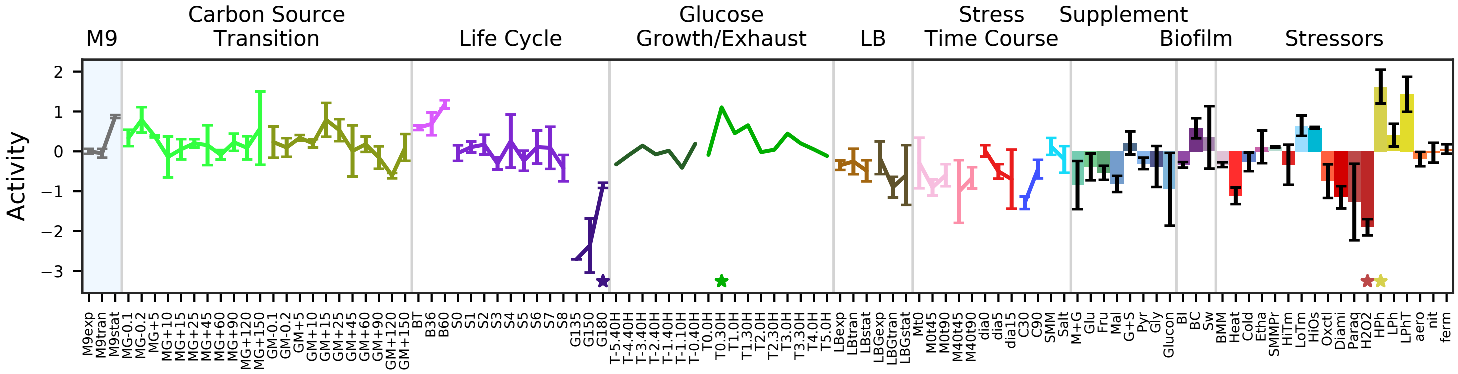


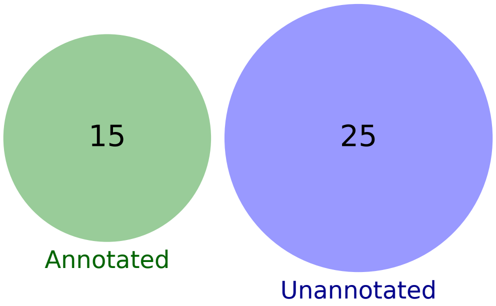

### 68 - Uncharacterized 3

Biological Function:
Contains many poorly characterized RNAs with noisy activity but clear time correlations, especially in sporulation. May be indicative of overall decreased translational activity

No known regulator

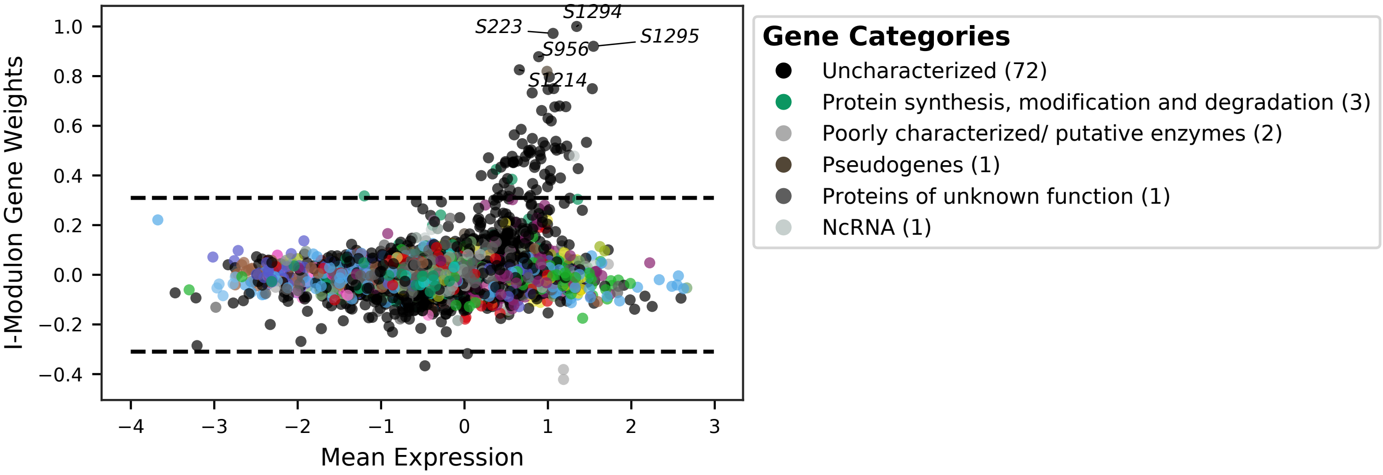


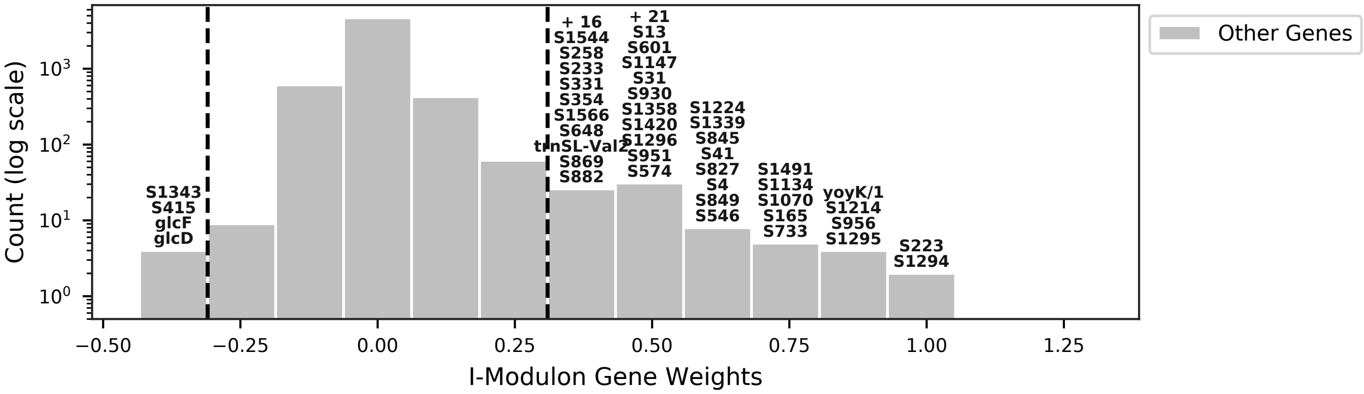


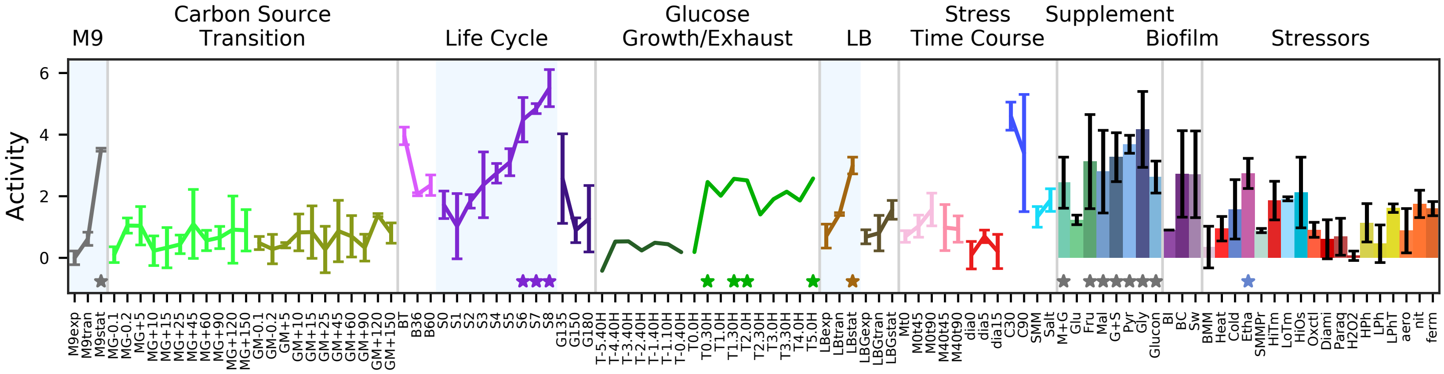


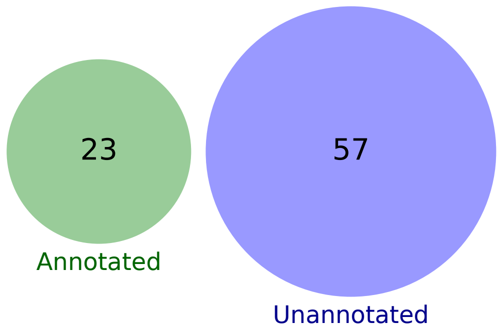

### 69 - Uncharacterized 4

Biological Function:
Contains many poorly characterized RNAs with noisy activity

No known regulator

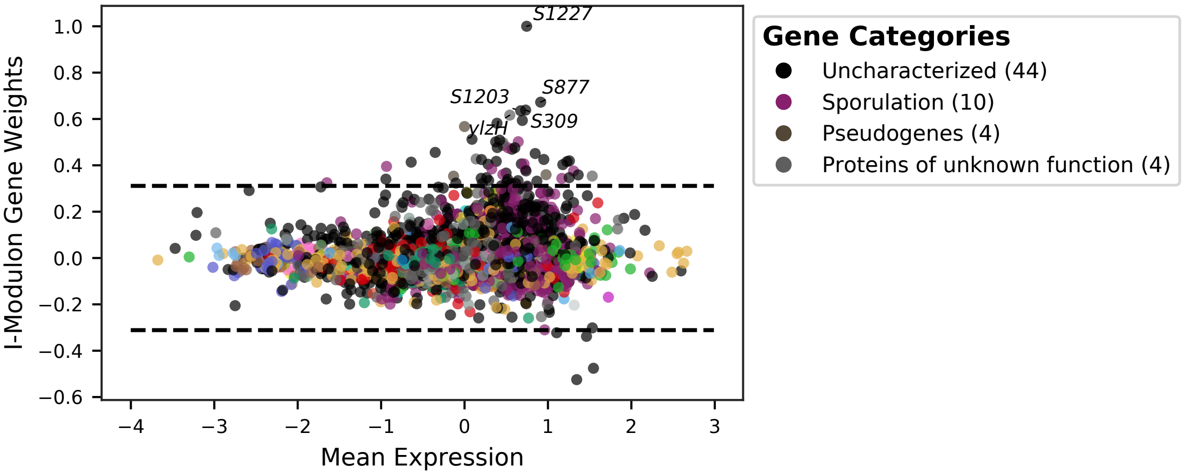


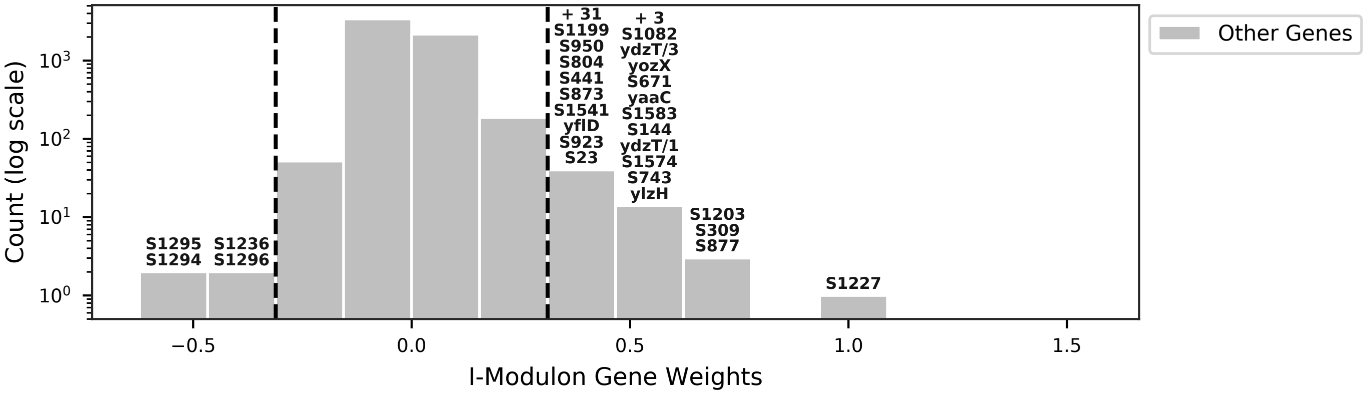


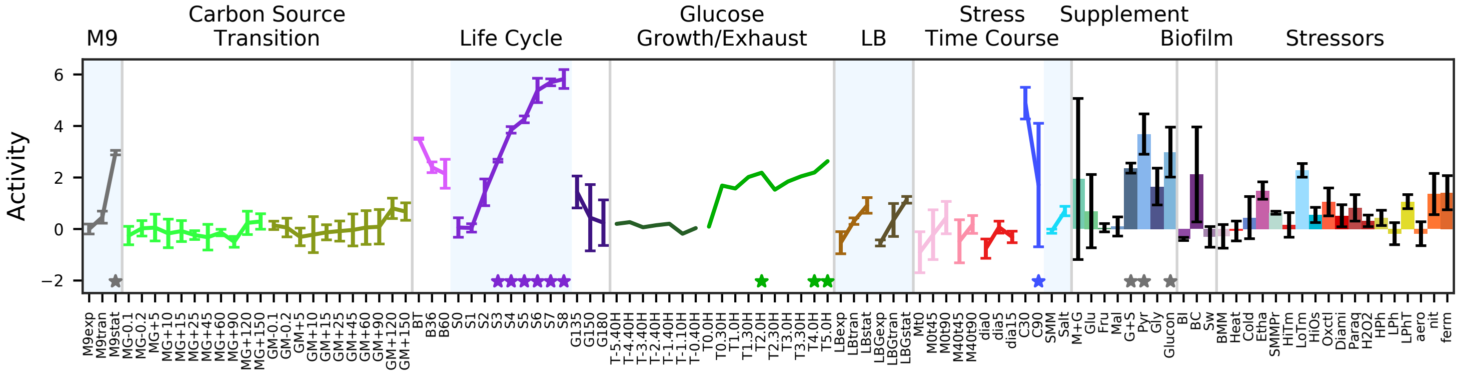


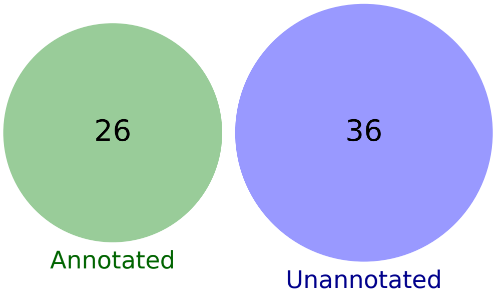

### 70 - Uncharacterized 5

Biological Function:
Contains many poorly characterized RNAs with noisy activity

No known regulator

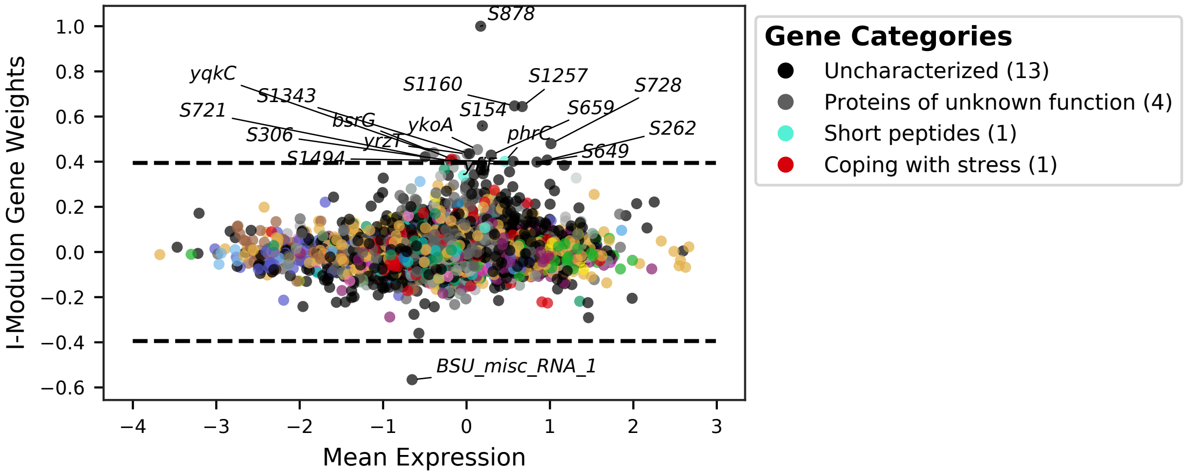


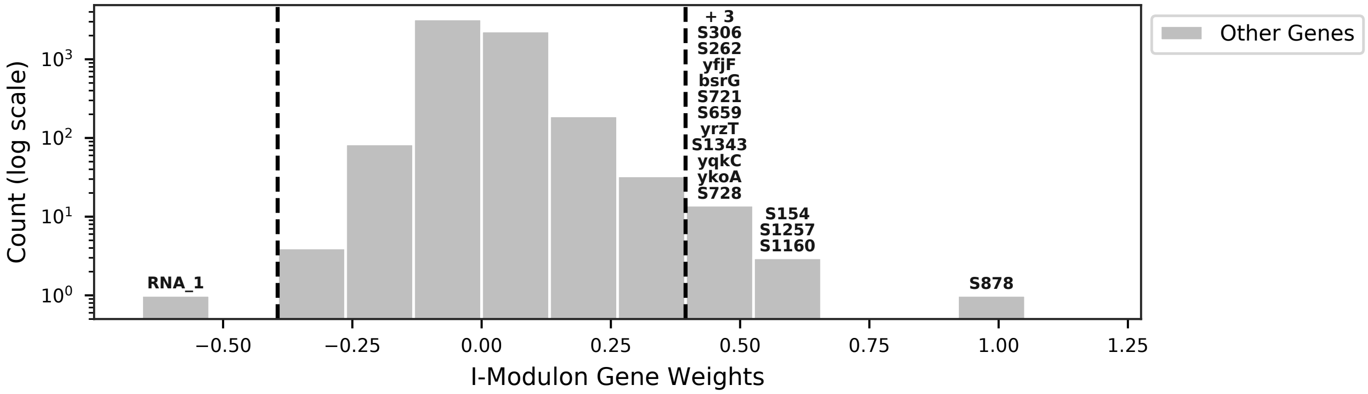


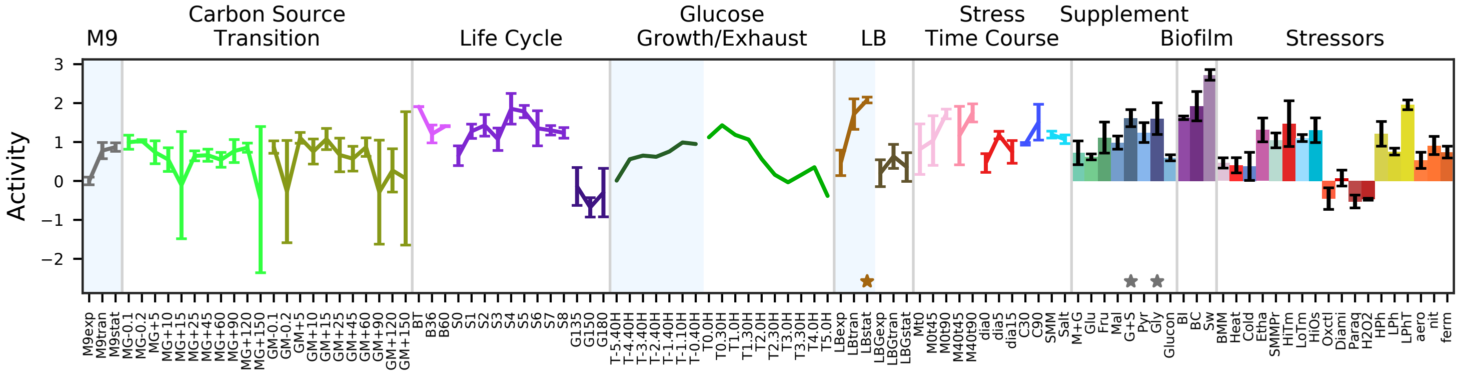


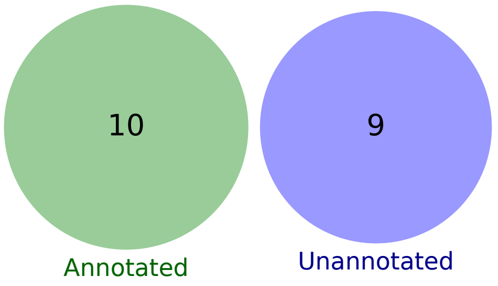

### 71 - Uncharacterized 6

Biological Function:
5’ ends of tRNAs and independent transcripts

No known regulator

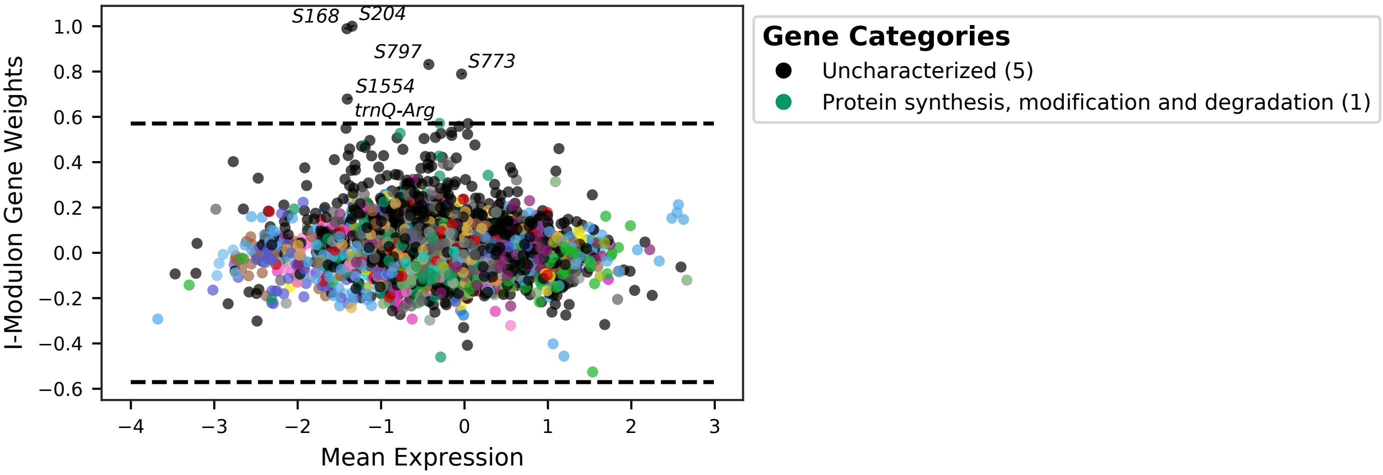


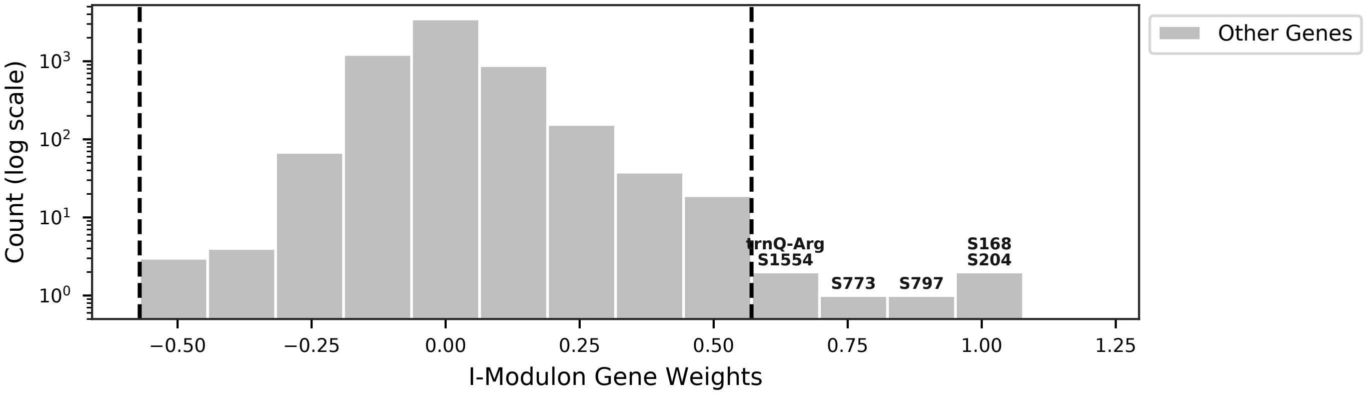


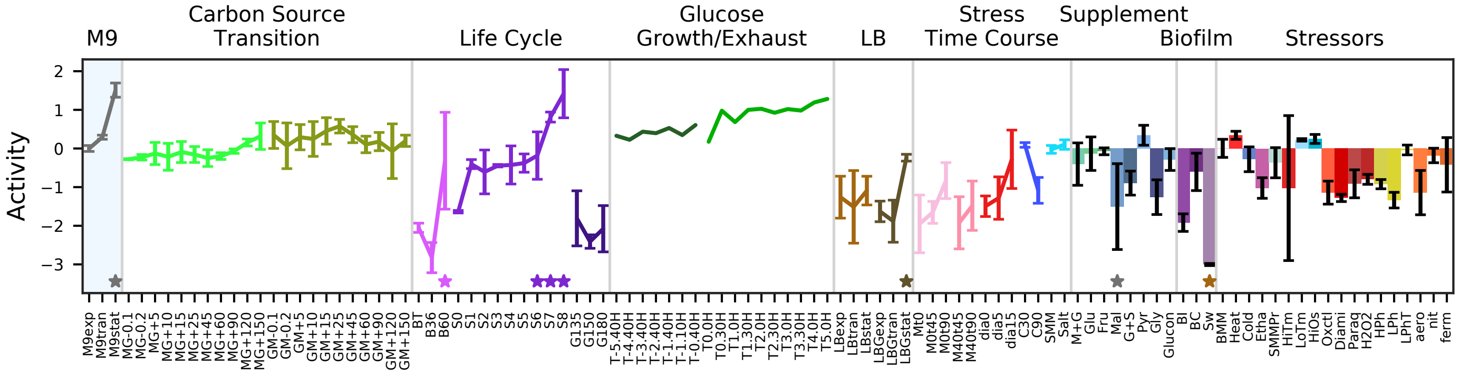


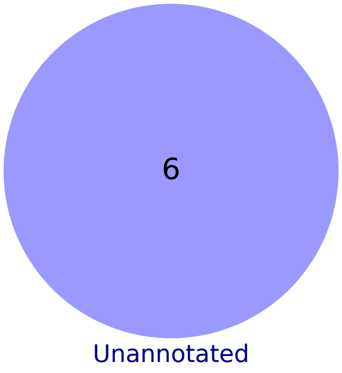

### 72 - Noise – S144

Biological Function:
Likely accounting for noisey RNA S144, an independent transcript

No known regulator

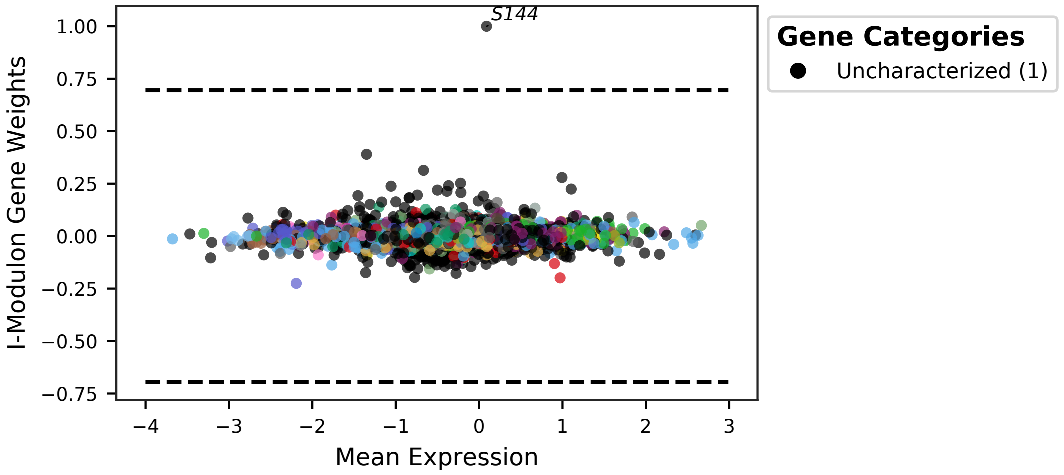


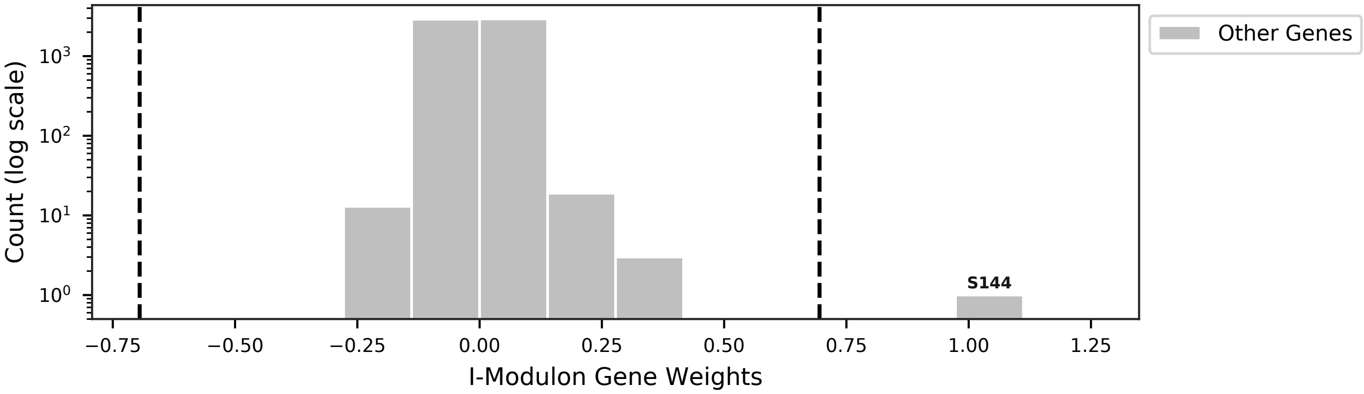


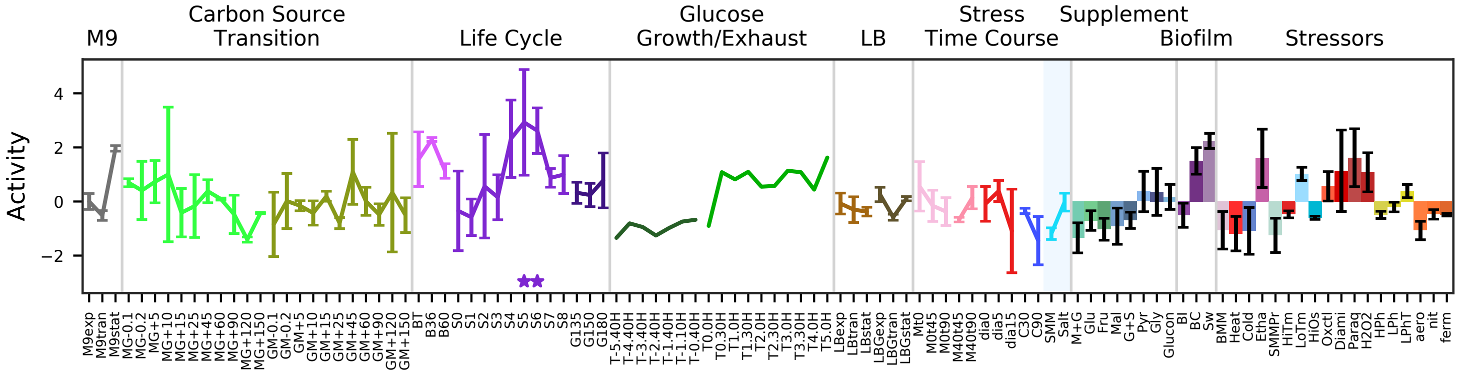


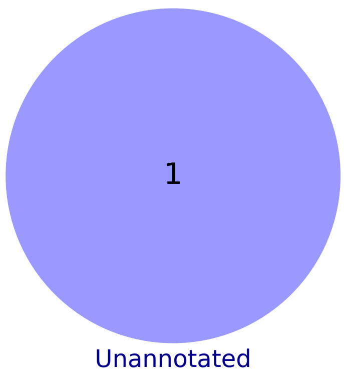

### 73 - Noise – S415

Biological Function:
Noisy independent transcript S415

No known regulator

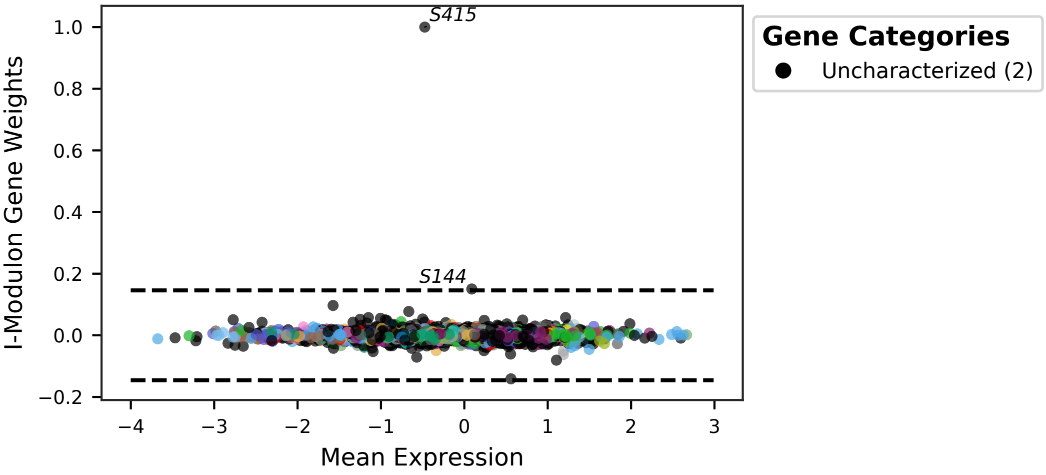


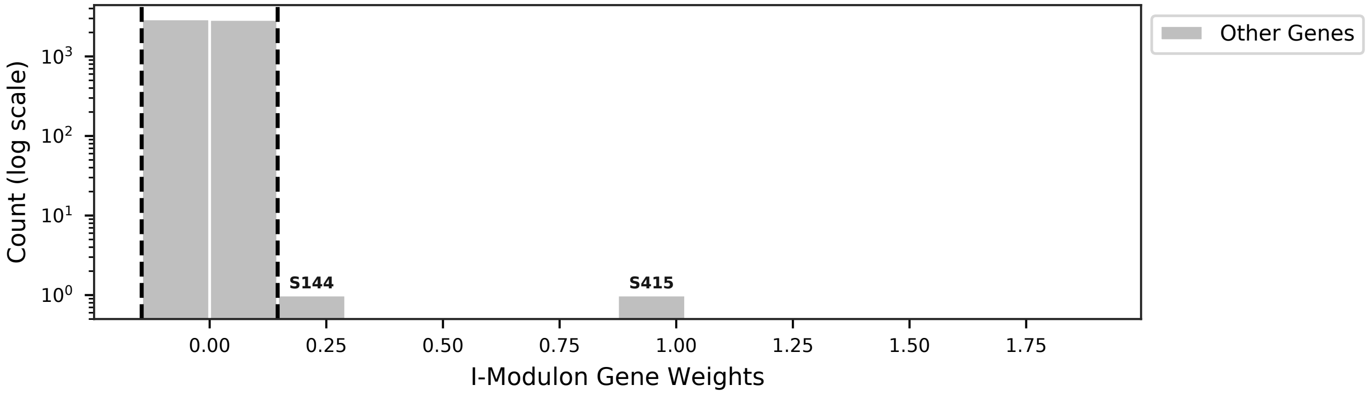


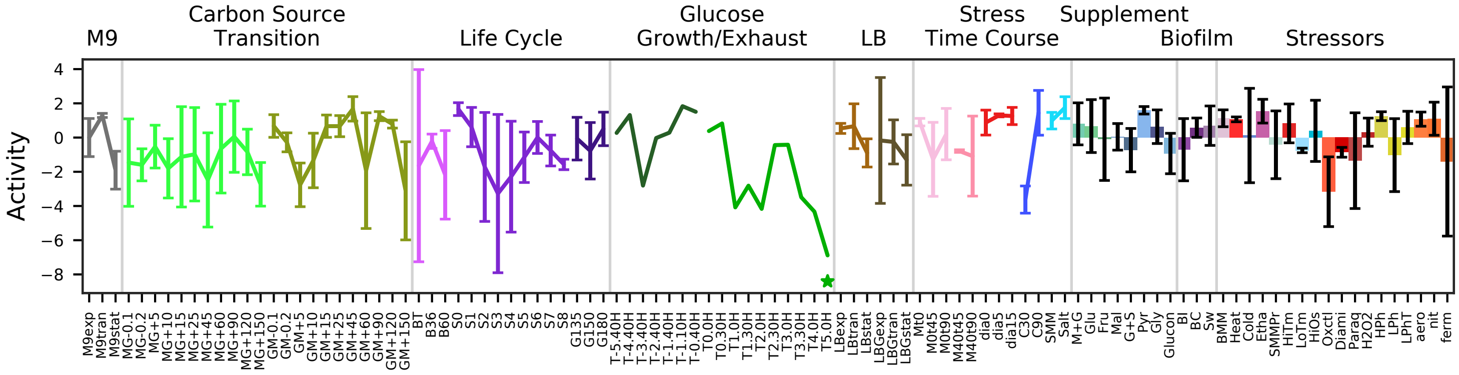


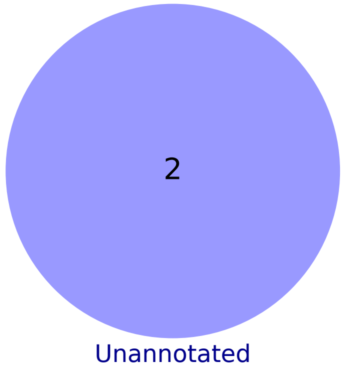

### 74 - Noise – S1235

Biological Function:
Independent transcript S1235, behaves similar to late sporulation I-modulons

No known regulator

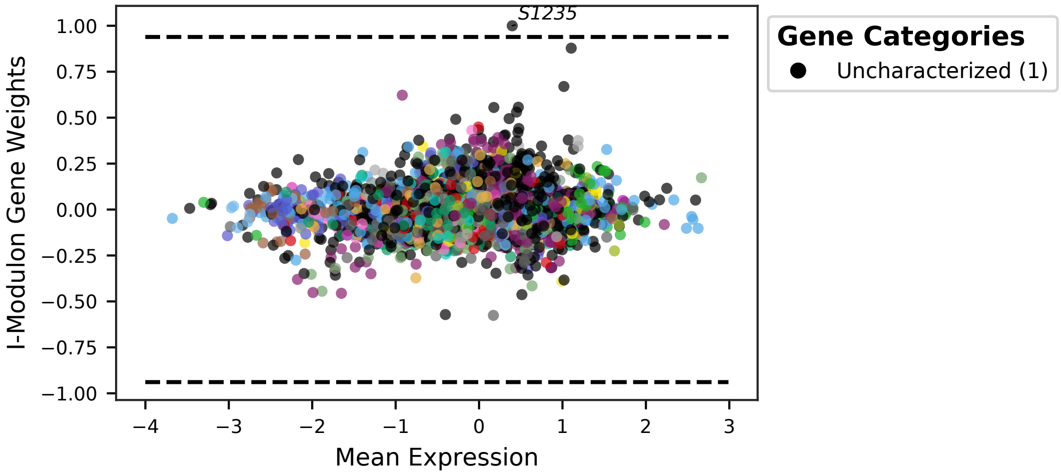


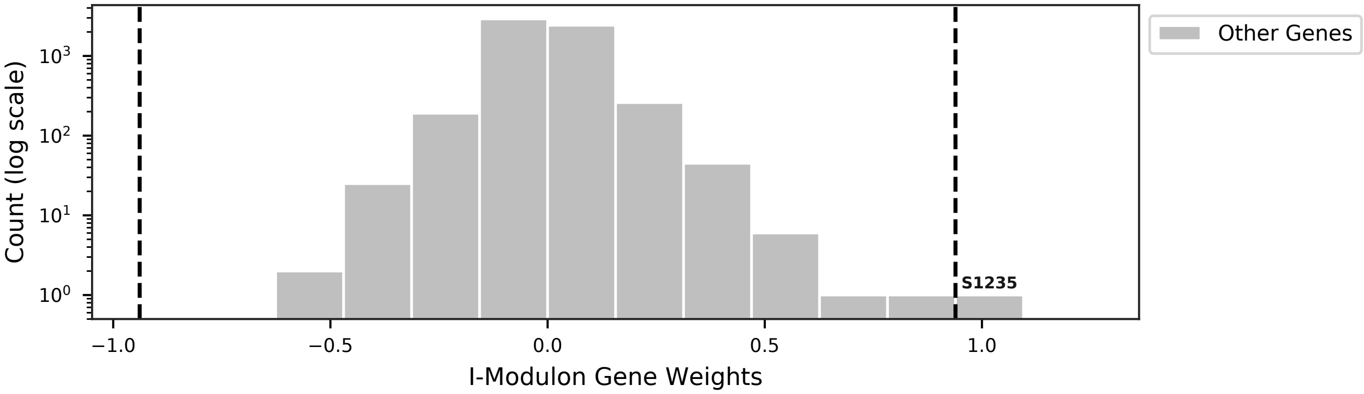


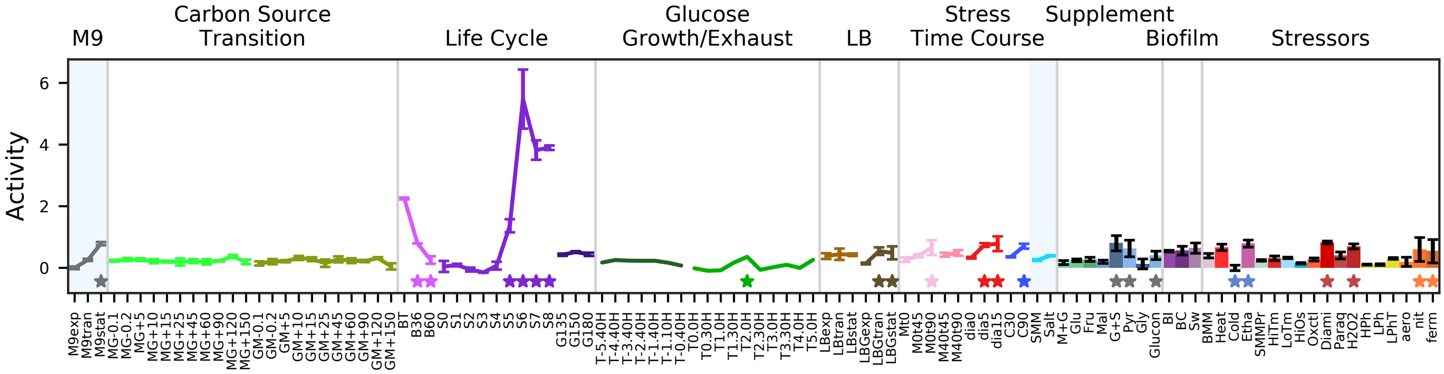


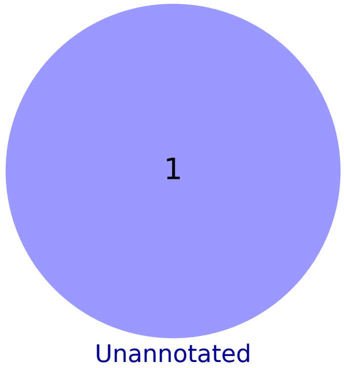

### 75 - Noise – S1210

Biological Function:
Noisy independent transcript S1210

No known regulator

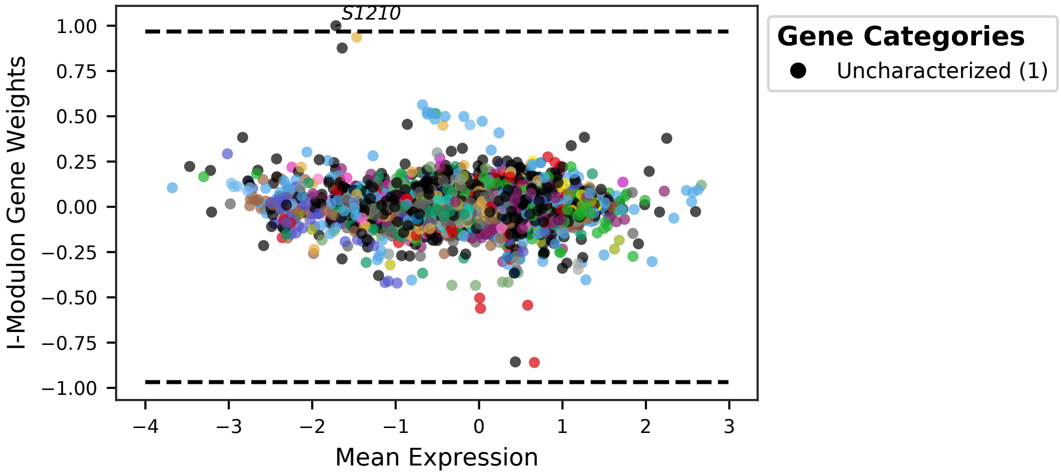


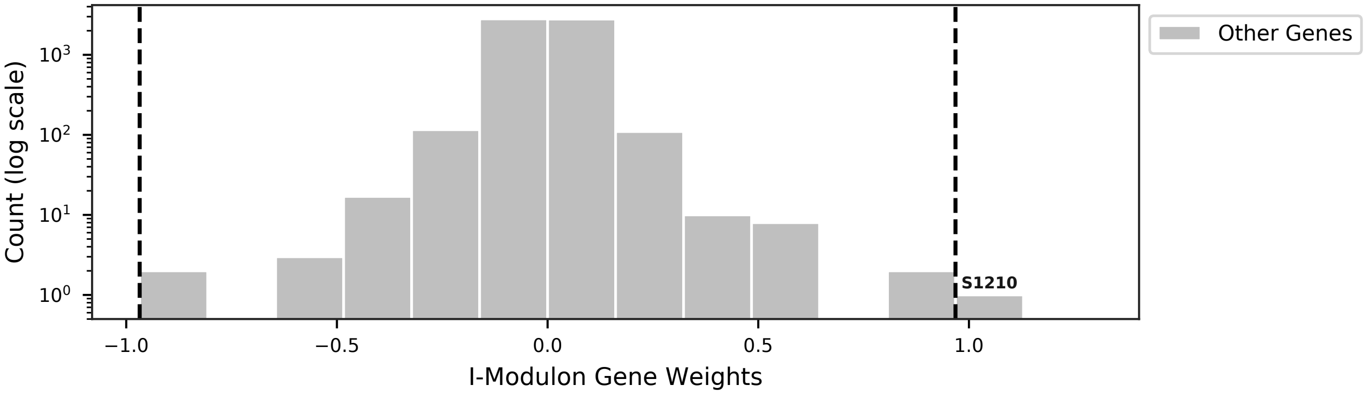


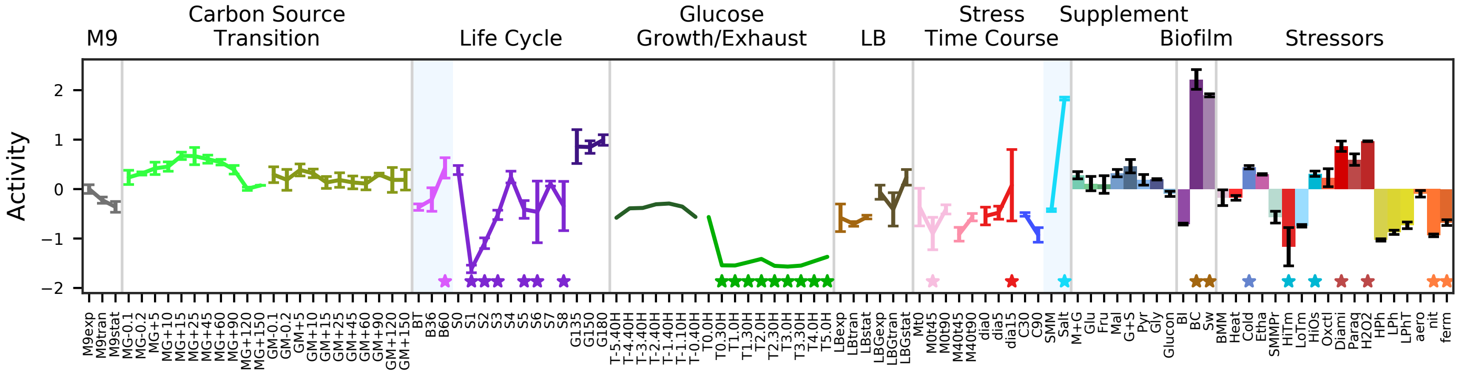


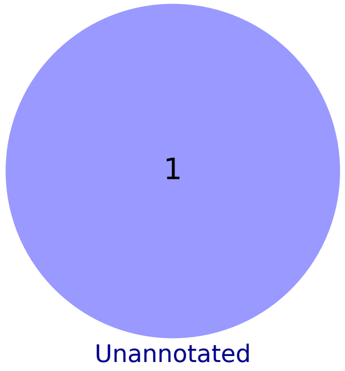

### 76 - Noise – Carbon Source Transition 1

Biological Function:
Likely accounting for noise in the carbon source transition experiments

No known regulator

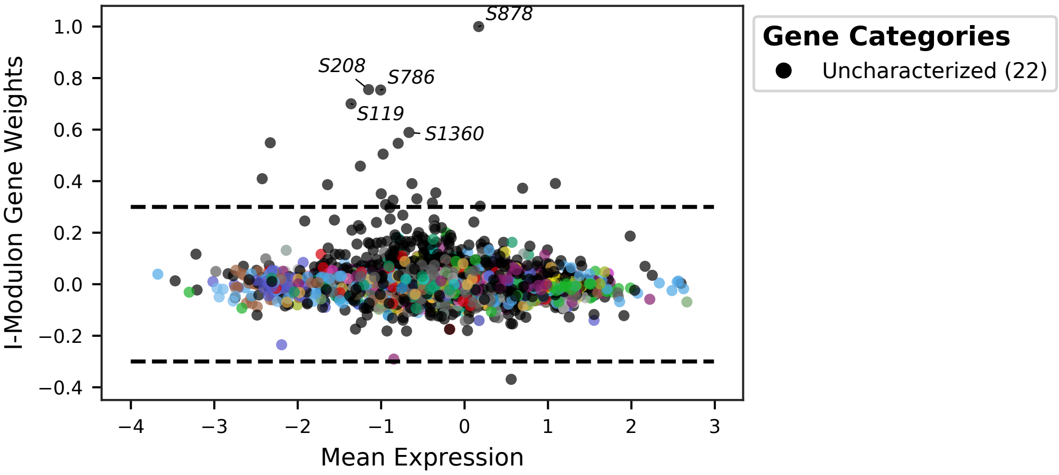


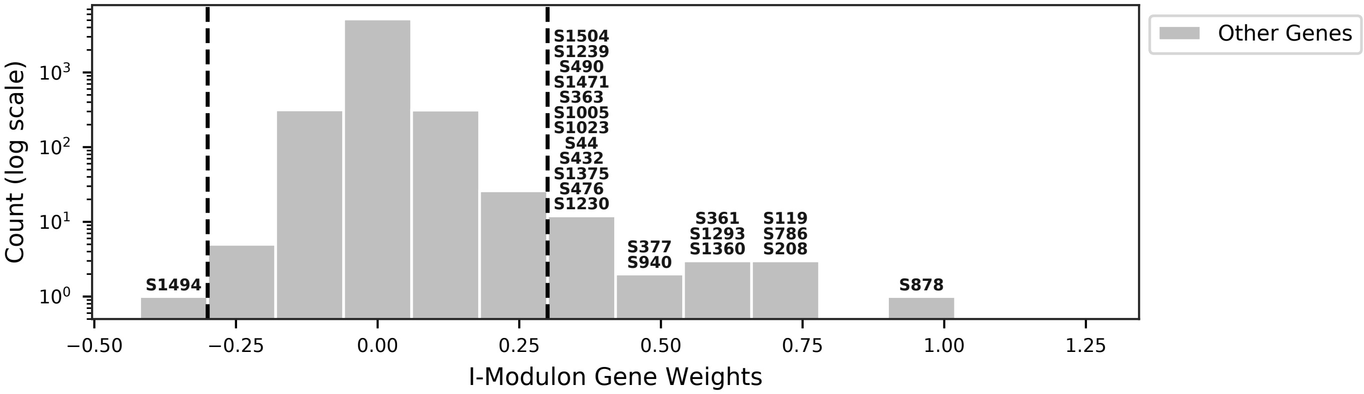


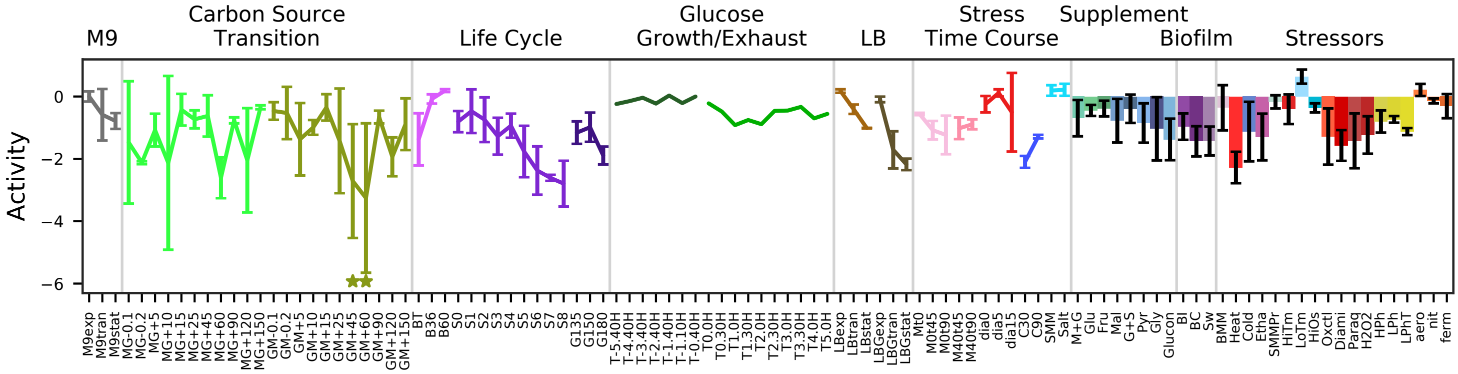


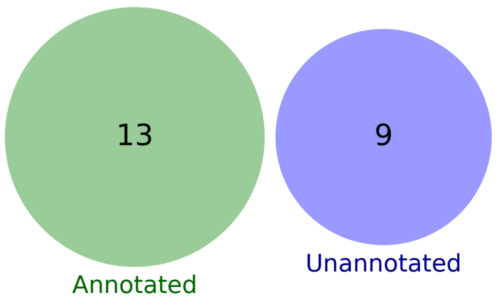

### 77 - Noise – Carbon Source Transition 2

Biological Function:
Likely accounting for noise in carbon source transition and gluconnate experiments

No known regulator

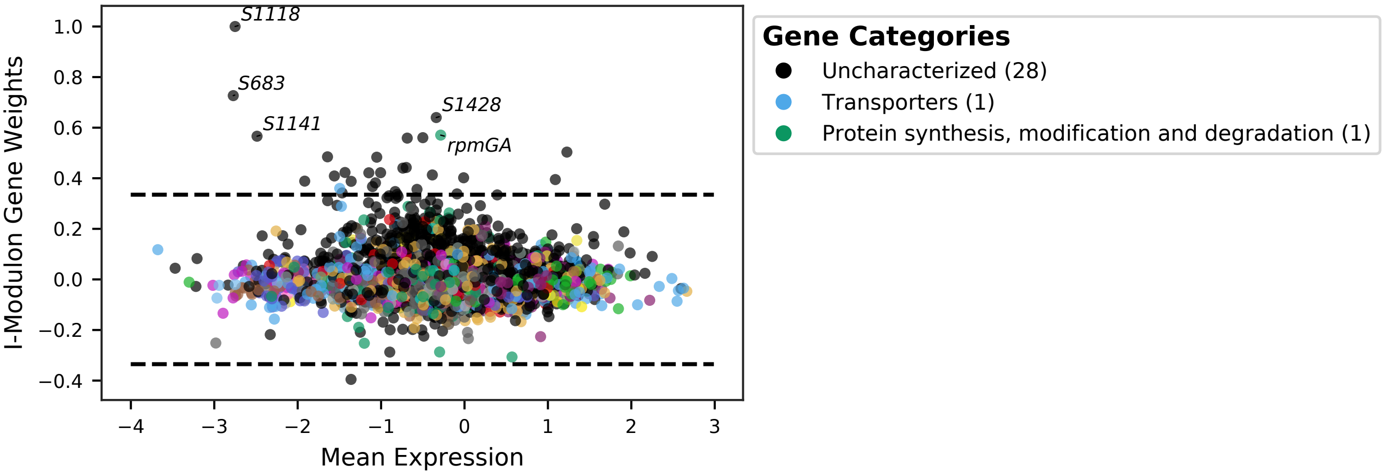


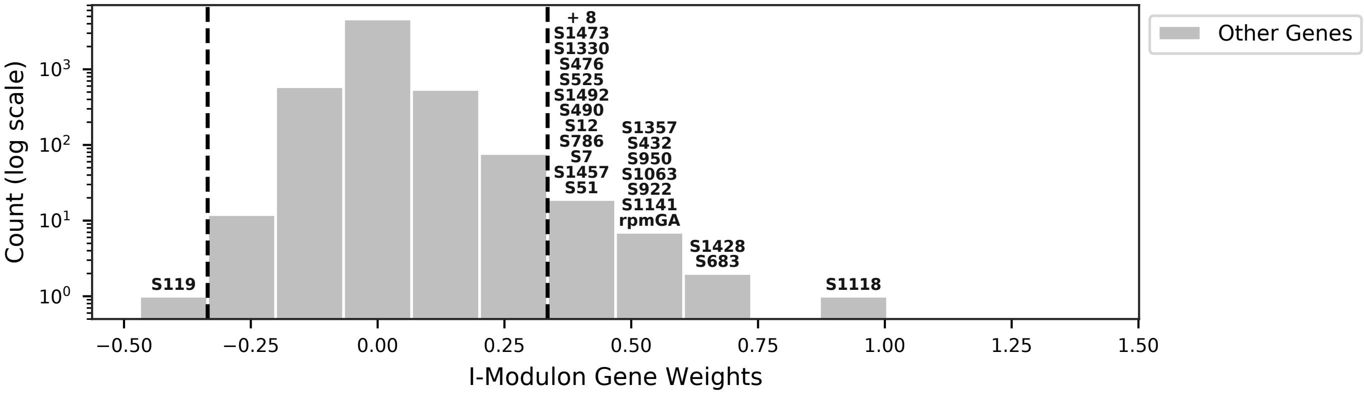


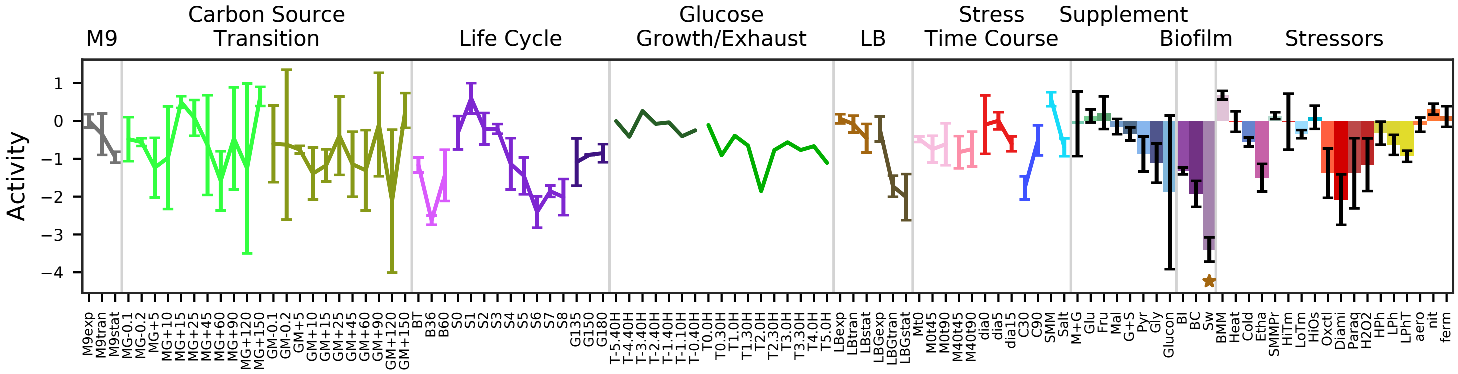


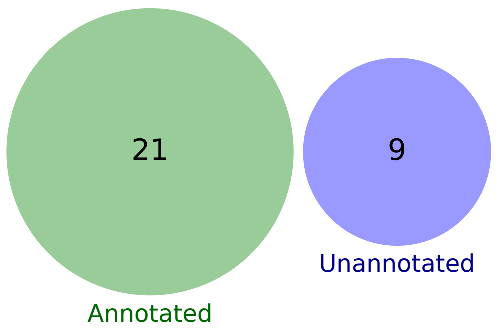

### 78 - Noise – LBGexp_1

Biological Function:
Likely accounting for noise in the first sample of LBG_exp. Top few RNAs are 5' UTRs of glucose metabolism genes

No known regulator

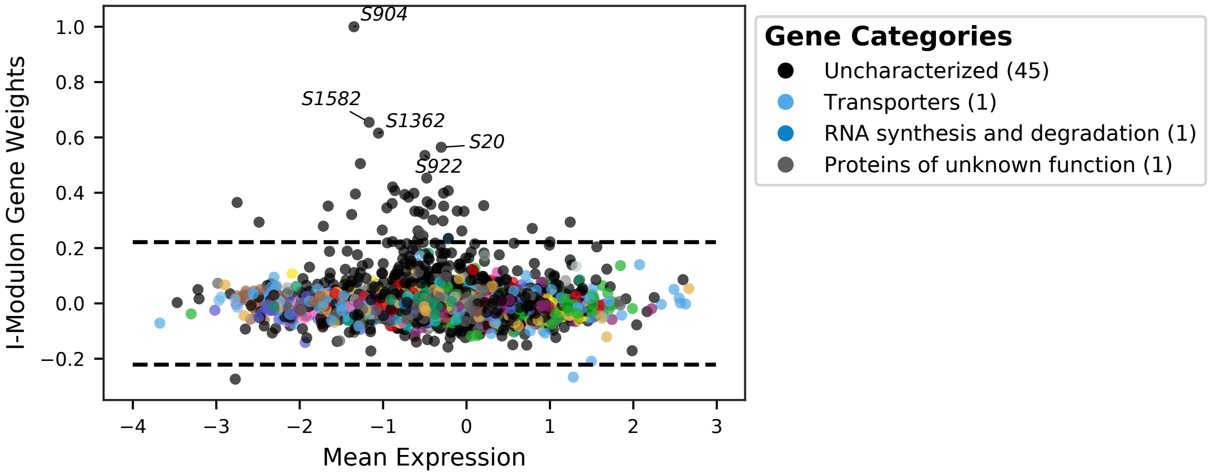


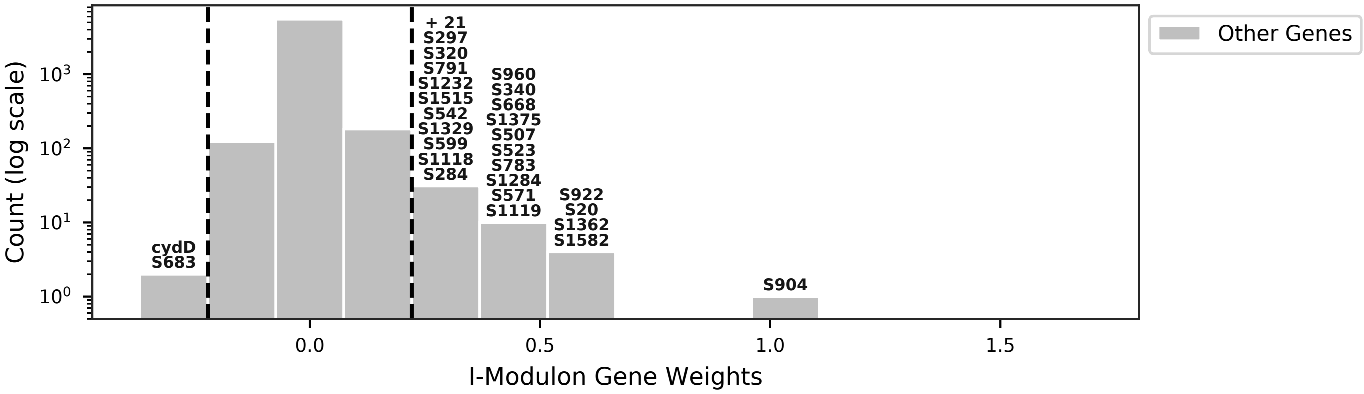


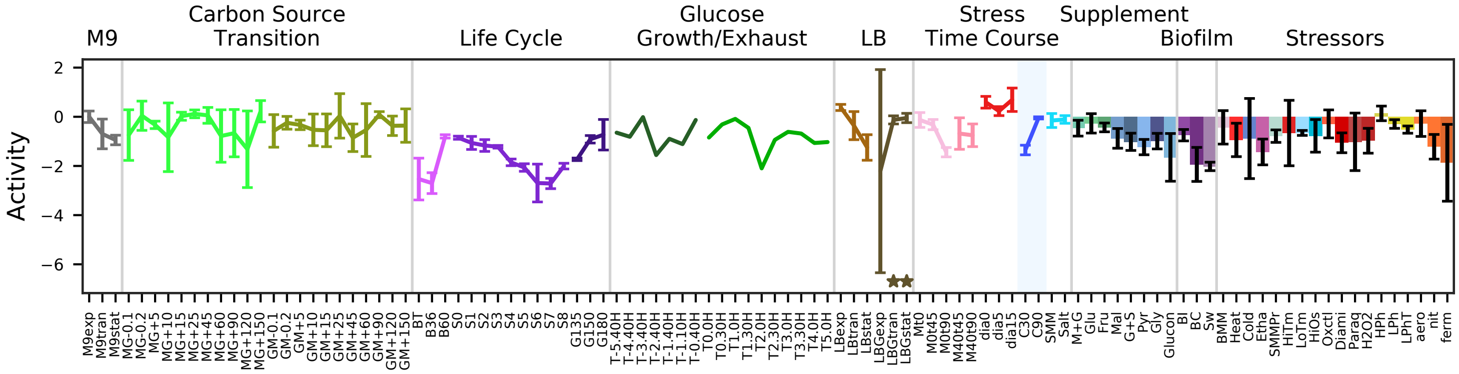


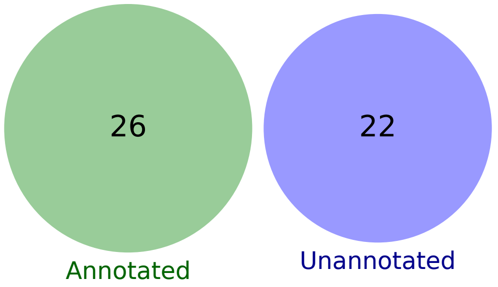

### 79 - Noise – Diamide 15

Biological Function:
Likely accounting for noise in the diamide after 15 minutes and LBG exponential conditions

No known regulator

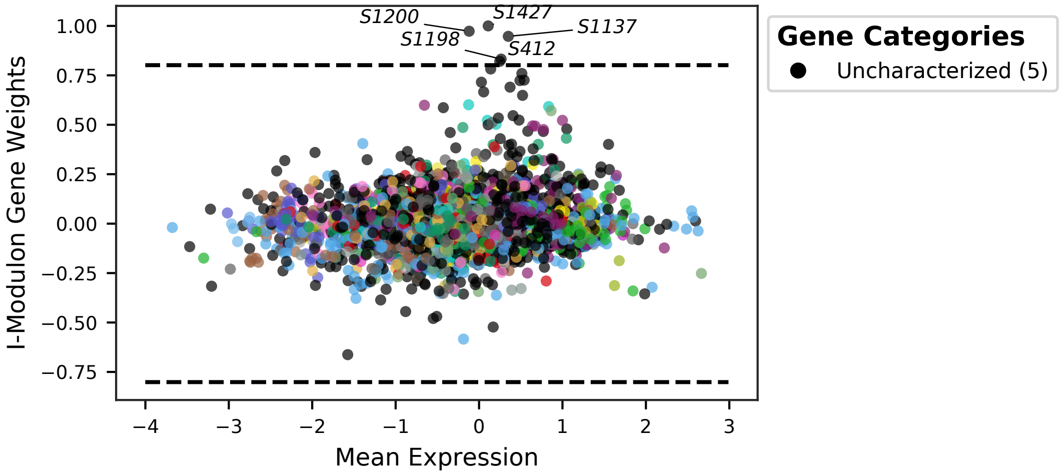


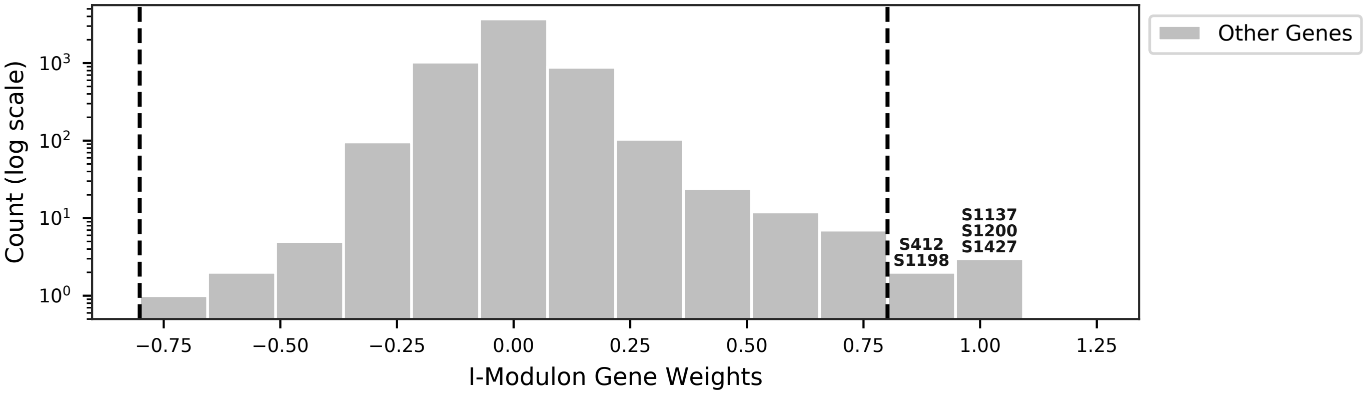


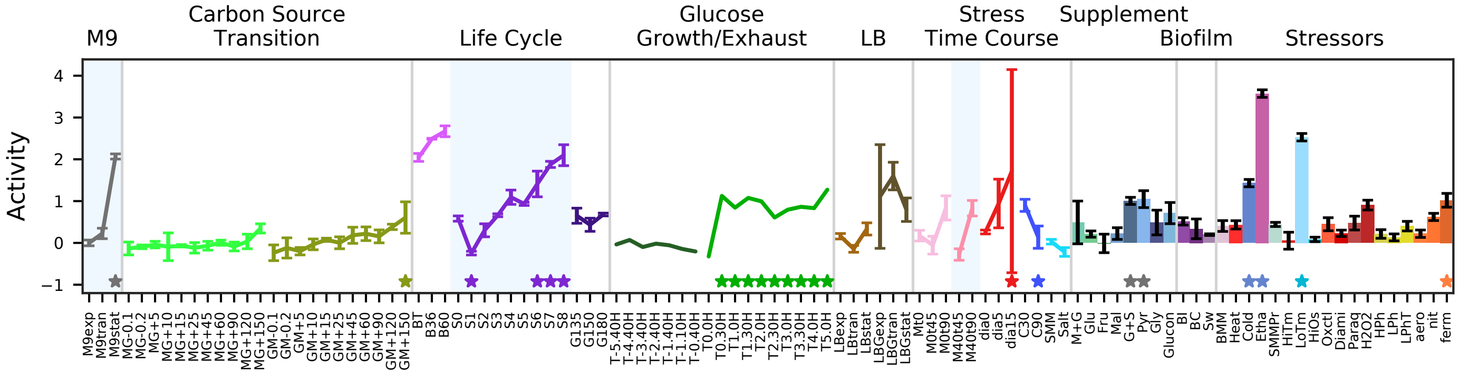


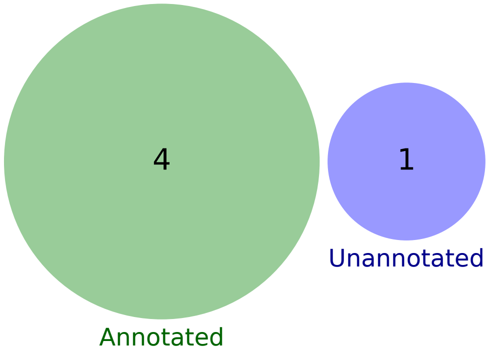

### 80 - Empty 1

Biological Function:
Contains no genes

No known regulator

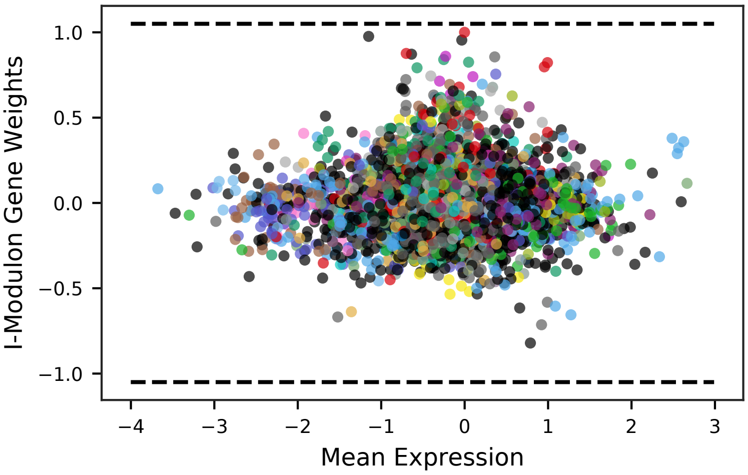


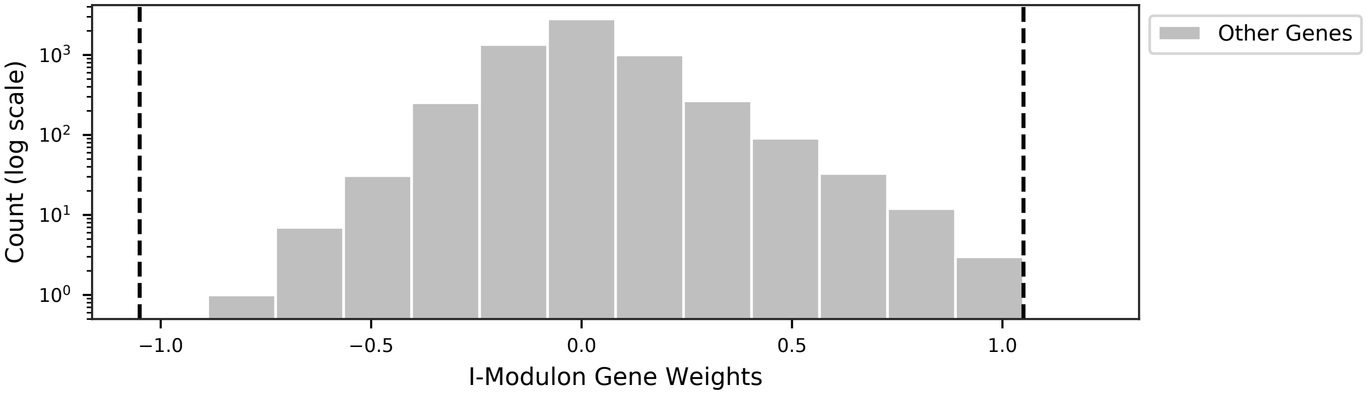


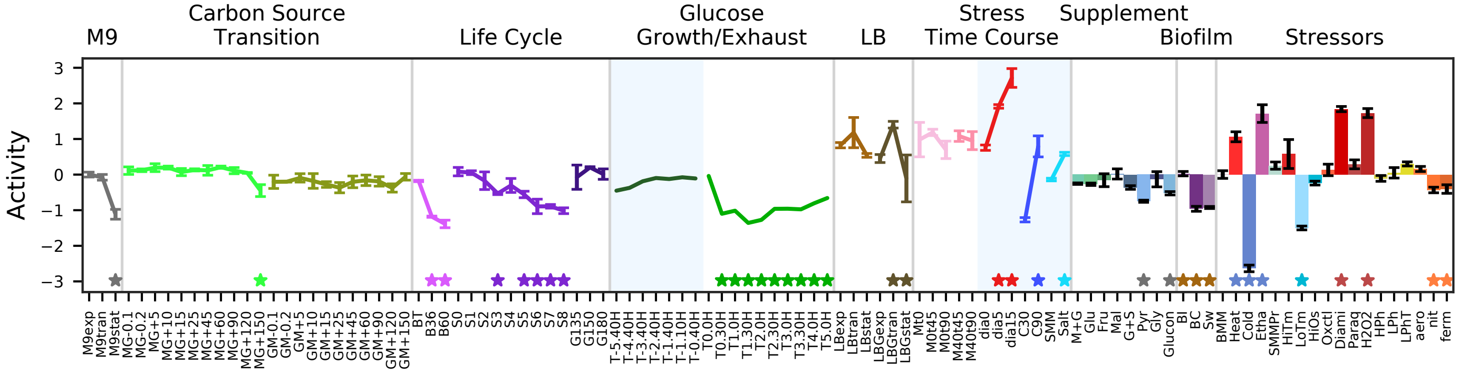

### 81 - Empty 2

Biological Function:
Contains no genes

No known regulator

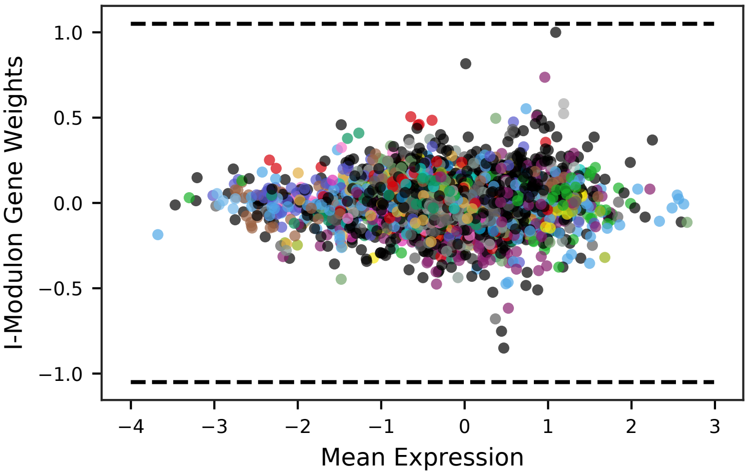


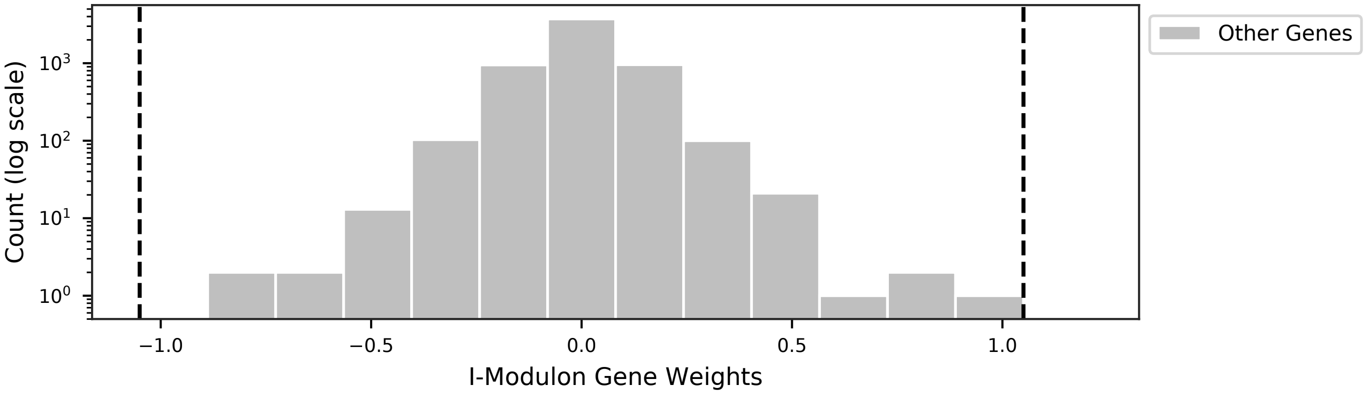


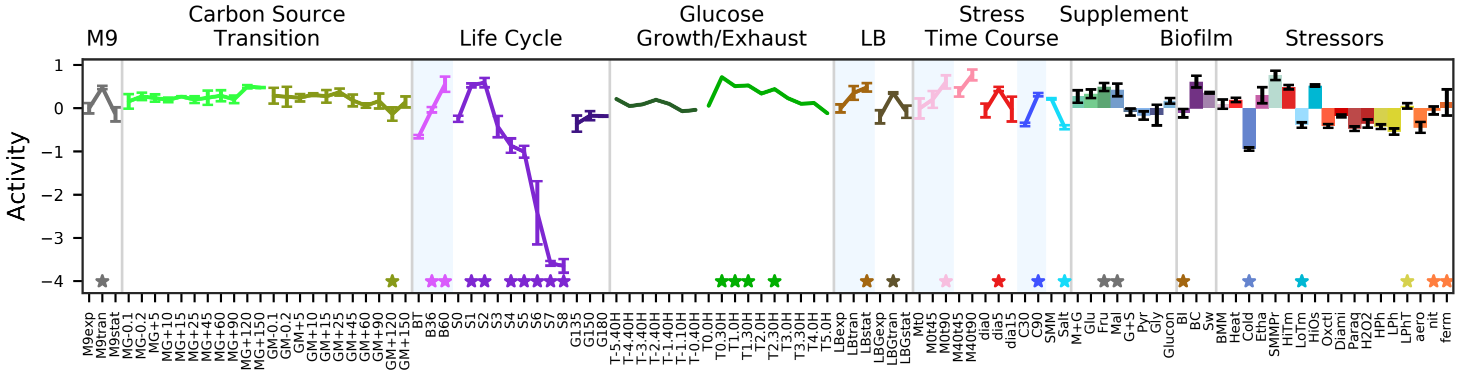

### 82 - Empty 3

Biological Function:
Contains no genes

No known regulator

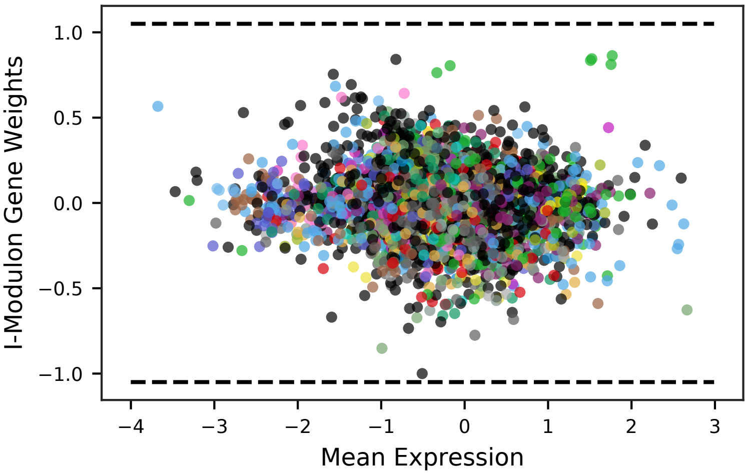


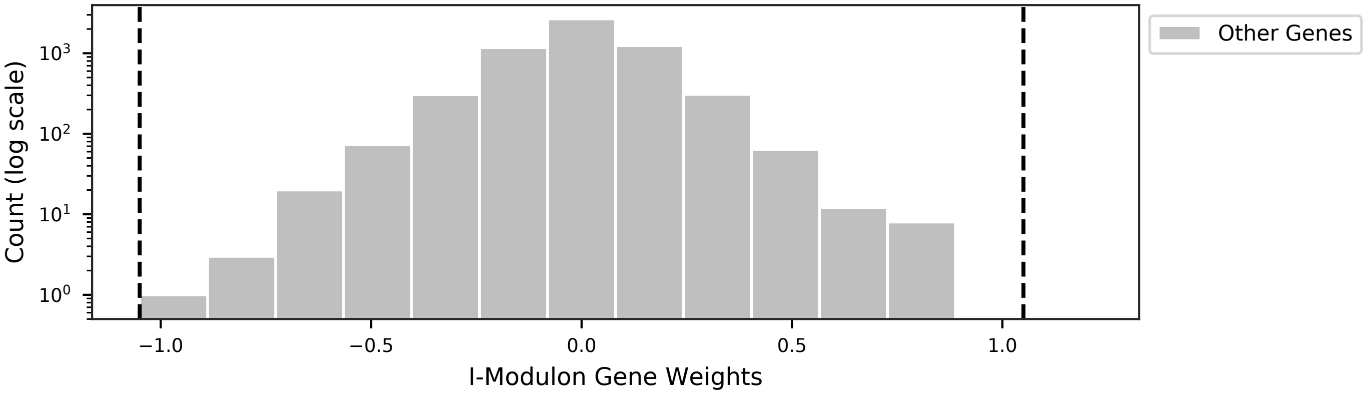


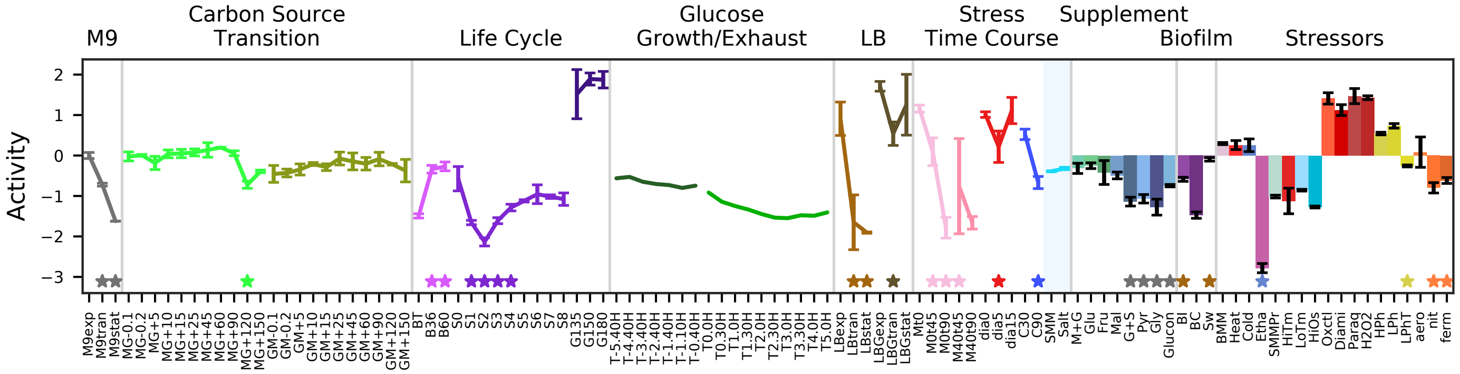
